# Structural basis of the two-photon photoactivation mechanism of orange carotenoid protein

**DOI:** 10.64898/2026.04.21.718960

**Authors:** Rory Munro, Elena A. Andreeva, Elisabeth Hartmann, Quentin Goor, Hosni El Zein, Stanislaw Nizinski, Adjélé Wilson, Elke De Zitter, Gregory Effantin, Nicolas Coquelle, Ninon Zala, Martin V. Appleby, Shira Bar-Zvi, Camila Bacellar, Emma Beale, Emmanuelle Bignon, Bernhard Brutscher, Martin Byrdin, Claudio Cirelli, Florian Dworkowski, Lutz Foucar, Guillaume Gotthard, Alexander Gorel, Marie Luise Grünbein, Mario Hilpert, Philip J.M. Johnson, Marco Kloos, Gregor Knopp, Karol Nass, Gabriela Nass Kovacs, Dmitry Ozerov, Christopher J. Milne, Gotard Burdziński, Christophe Chipot, Yasaman Karami, François Dehez, Martin Weik, R. Bruce Doak, Robert L. Shoeman, Giorgio Schirò, Michel Sliwa, Diana Kirilovsky, Ilme Schlichting, Jacques-Philippe Colletier

## Abstract

Cyanobacteria have produced Earth’s oxygen for 2.4 billion years by adapting to fluctuating irradiance. This adaptation relies on orange carotenoid protein (OCP), which mediates light-intensity– dependent photoprotective energy dissipation using a unique two-photon absorption mechanism. Photon absorption by ground-state OCP (OCP^O^) generates a metastable intermediate (OCP^1hν^) that either relaxes thermally or, upon absorption of a second photon within ∼1 s, converts to the active photoprotective state (OCP^R^). By integrating static and time-resolved crystallography, cryo-EM, computation, spectroscopy and biochemistry, we assign the structure of OCP^1hν^, establish its functional relevance and capture structural snapshots along the OCP^O^→OCP^1hν^ and OCP^1hν^→OCP^R^ photochemical pathways. We elucidate the molecular mechanism of OCP, which serves as a unique biological circuit breaker protecting the photosynthetic machinery from high light flux.

## Introduction

For survival, cyanobacteria need to dynamically adjust photosynthesis and photoprotection to changing sunlight intensities. Central to this adaptation is the orange carotenoid protein (OCP)^1–3^ which binds keto-carotenoid pigments like canthaxanthin (CAN)^4^, echinenone (ECN)^5^ and 3’-hydroxy-echinenone. Under low irradiance, OCP remains in its inactive orange state (OCP^O^). However, under continuous exposure to intense blue-green light, OCP^O^ converts to an active red form (OCP^R^)^4,6,7^, that binds to phycobilisome complexes^2,7,8^ —the primary light-harvesting antennas in cyanobacteria. Upon this interaction, the excess energy is dissipated, preventing the formation of harmful singlet oxygen radicals and protecting the photosynthetic machinery. Since this comes at the price of a reduced photosynthetic efficiency, the formation of OCP^R^ is finely tuned by light intensity.

In ECN-functionalized OCP, this tuning is achieved by a sequential two-photon absorption mechanism^9–13^, where absorption of a first photon generates a metastable OCP^1hv^ intermediate (with a similar absorption spectrum as OCP^O^), that needs to absorb a second photon after a roughly one second delay to complete activation to OCP^R^, otherwise it reverts thermally to OCP^O^ (Fig. 1A). In contrast to a one-photon absorption mechanism, this mechanism results in a non-linear response with respect to light intensity, activating OCP^R^-driven photoprotective mechanisms only above a specific light intensity threshold and only if the light persists long enough. Thus, ECN-OCP acts as a circuit-breaker, disrupting photosynthesis only above a certain intensity threshold. By contrast, the photoactivation of CAN-OCP has been proposed to occur via a one-photon absorption mechanism^13^ (Fig. 1B), whereby an intermediate equivalent to ECN-OCP^1hv^ would decay thermally to OCP^R^, effectively avoiding the need for a second light-induced step (Fig. 1B). Structurally, it is known that the transition from OCP^O^ to OCP^R^ entails a 12 Å migration of the carotenoid from the interface between the N- and C- terminal domains (NTD, CTD) into the NTD, and a separation of the two domains^4,14^ (Fig. 2A). However, the structure of the OCP^1hν^ intermediate is unknown, as are the pathways from OCP^O^ to OCP^1hν^ and from OCP^1hν^ to OCP^R^. We addressed this knowledge gap by combining mutagenesis, spectroscopy, cryo-electron microscopy (cryo-EM) as well as static and time-resolved room-temperature (RT) crystallography on CAN- and ECN-functionalised OCP from *Planktothrix aghardii*. We identify the structure of the OCP^1hv^ intermediate and show that transition to this conformation is essential to elicit formation of OCP^R^. By performing crystallographic pump probe experiments starting from both OCP^O^ and the OCP^1hv^ intermediate, we map sub-ps to µs structural changes *en route* to OCP^1hv^ and OCP^R^, respectively. We also characterize the crystallographic endpoint of the OCP^1hv^ to OCP^R^ transition using serial crystallography after prolonged illumination. Together, our findings provide the structural basis for the unique two-photon absorption mechanism of OCP.

**Figure 1:**
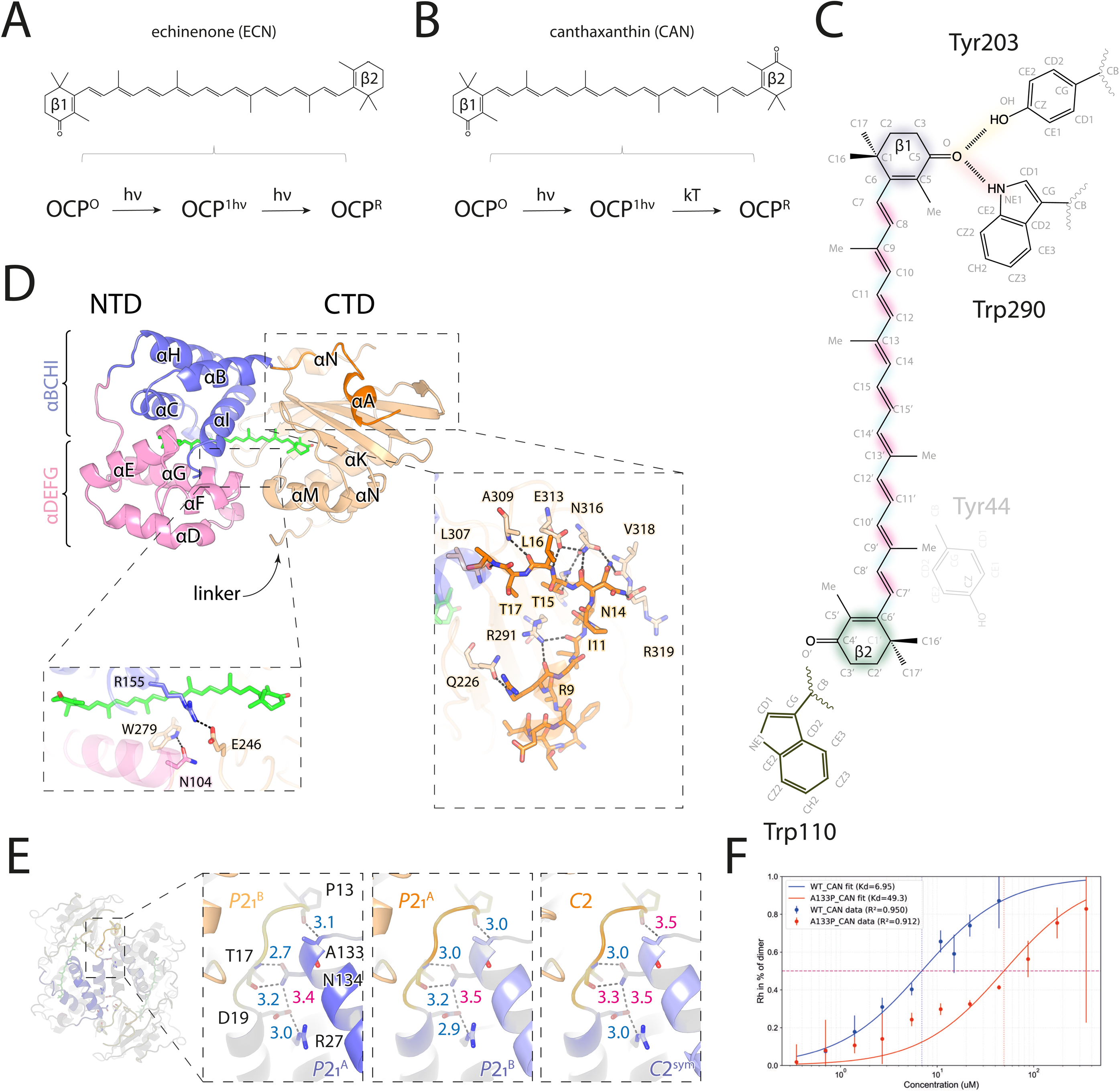
General features of the OCP photosensor. (A) It has been proposed that ECN-OCP photoactivates via a strict two-photon absorption mechanism^13^, while CAN-OCP (B) photoactivates via a single photon absorption mechanism^13^, where the OCP^1hν^ intermediate can thermally reach OCP^R^. In both cases, OCP^R^ reverts thermally to OCP^O^. (C) Nomenclature used to refer to CAN atoms, dihedral angles, and overview of the main interactions of CAN and the OCP scaffold. The keto-group at the β1 ionone ring H-bonds to residues Y203 and W290 in the C-terminal domain (CTD); the β2 ionone ring stacks between the aromatic rings of Y44 and W110 in the N-terminal domain (NTD). (D) Architecture of the OCP photosensor, highlighting the (i) fully-helical nature of the N-terminal domain (NTD) composed of two four-helix bundles (αBCHI and αDEFG, in slate and pink, respectively) sandwiching the carotenoid at their interface and connected by long αCD and αGH loops; (ii) the mixed α/β nature of the CTD, onto which the αA helix attaches; and (iii) the main tether between the two domains, viz. the R155—E246 and N104—W279 salt-bridge and H-bond, respectively. (E) Clade 1 OCP such as the *Planktothrix agardhii* and *Synechocystis* PCC6803 OCP form dimers, through a high-conserved interface featuring the D19—R27 salt-bridge and three H-bonds, including the P13—A133 H-bonds connecting the αA of a monomer with the αGH loop of the second monomer. (F) Absence of the P13—A133 H-bond decreases the affinity of the *Planktothrix agardhii* OCP dimer ∼8-fold.

**Figure 2:**
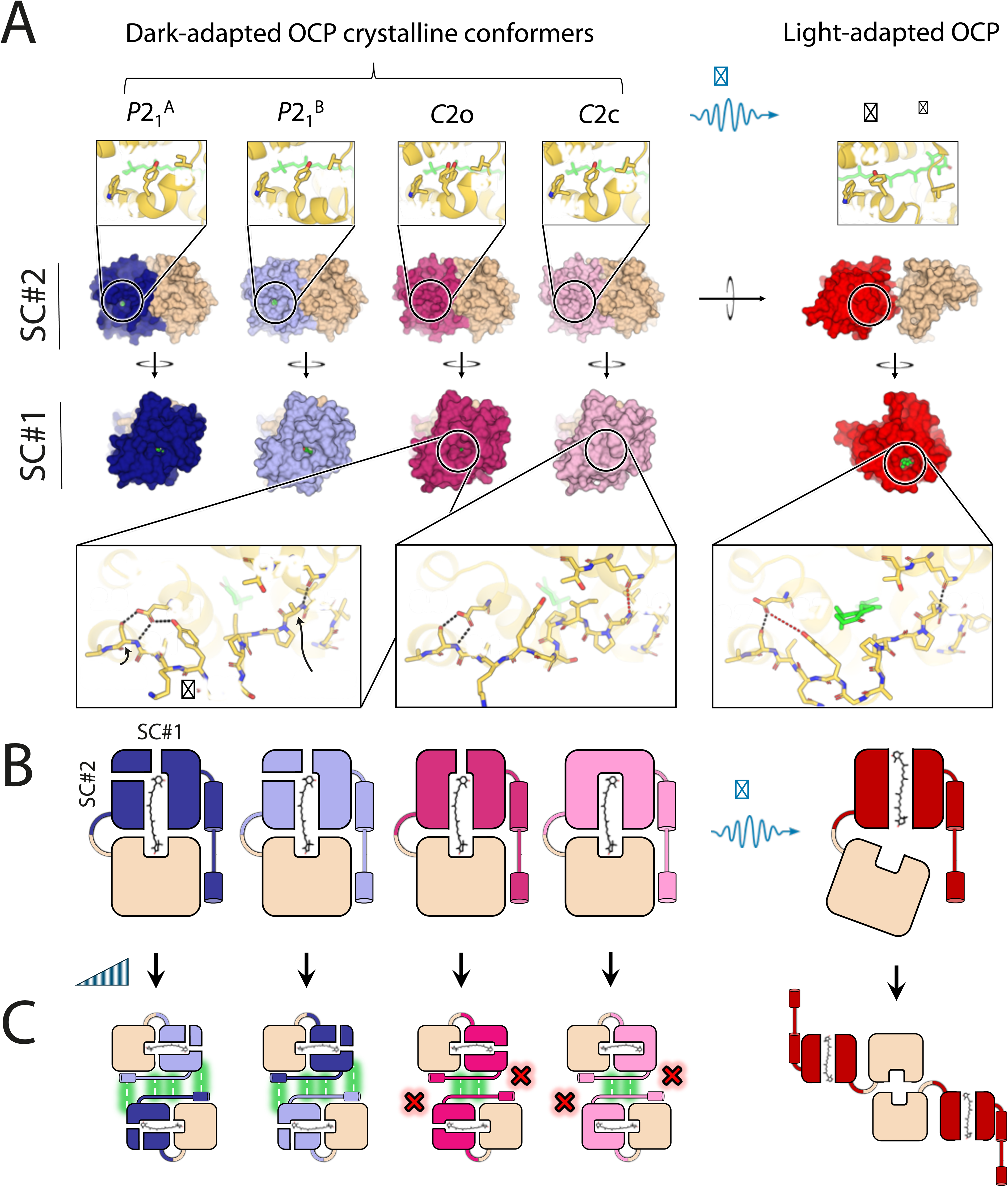
**The four-protomer problem. (**A) Dark adapted *Planktothrix agardhii* OCP can adopt four distinct sub-states in two crystal forms, depending on pH, relative humidity and illumination conditions. The conformational sub-states (colored by domain, with the CTD in beige and the NTD in different shades of blue (P2_1_ chains) or pink (C2 conformers) differ in the conformation of their αCD and αGH loops, and therefore in the shielding of the β2-ionone ring from solvent within the carotenoid tunnel. Solvent channel #2 (SC#2), bordered by the αC helix and the αCD loop, is open in the P2_1_^A^ and P2 ^B^ conformers, but closed in C2 conformers. These changes, which yield C2o, are required to allow further progression to the C2c state, characterized by additional closure of SC#1, upon folding of the αGH loop into a 3^10^ helix, and thus complete shielding of the β2-ionone ring from solvent. SC#1 must be open for the carotenoid to translocate along the tunnel, inducing domain dissociation and yielding the OCP^R^ state (red). (B) Schematic representation of the various OCP dark states identified in our crystals (P2_1_ ^A^, P2_1_ ^B^, C2o and C2c) and of the photo-activated OCP^R^ state, characterized by cryo-EM (PDB: 7SC9) in previous work. (C) All OCP dark-states form dimers upon an increase of concentration; however, the C2o and C2c conformers lack the intermolecular P13—A133 H-bond tethering the αGH loop of one monomer to the αA helix of the other in the dimer, offering conformation “freedom” of the loop. OCP^R^ is also able to form dimers, supported by a different interface, unavailable in the dark states as it is otherwise occupied by the carotenoid.

## Results

### Structural characterization of the OCP dark states

A mechanistic analysis of structural changes occurring during a reaction demands a well-defined initial state. For OCP, identifying this starting point was not trivial since several dark state conformations have been described^4,6,14–20^ (Supplementary text 1). The general features of the OCP^O^ protomer are the H-bonding of the 4-keto oxygen of the β1-ionone ring of the pigment to Y203 and W290 in the mixed α/β CTD (residue numbering according to *Planktothrix aghardii* OCP), and the sandwiching of the β2-ionone ring between the aromatic side chains of Y44 and W110 in the NTD (Fig. 1C). The latter residues are part of two four-helix bundles (αBCHI and αDEFG) lining the carotenoid tunnel in the fully α-helical NTD (Fig. 1D, Supplementary Fig. 1). The OCP^O^ structure is further stabilized by protein interactions, including the conserved N104–W279 and R155–E246 H-bond and salt-bridge, respectively, as well as the multiple H-bonds attaching the N-terminal αA-helix on the face of the β-sheet opposed to Y203 and W290 (Fig. 1D). These interactions are conserved in all known OCP structures. Clade 1 OCPs^21^, such as those from *Planktothrix aghardii* and *Synechocystis* PCC6803, additionally form dimers (Kd ∼ 8 and 14 µM, respectively^10^), which is reflected in their two crystal forms (Fig. 1E). The P2_1_ space group of the *Planktothrix aghardii* OCP (hereafter referred to as OCP) features a biological dimer in the asymmetric unit, with constitutive monomers referred to as P2_1_^A^ and P2_1_^B^. By contrast, the C2 asymmetric unit contains a monomer, from which a similar, perfectly-symmetric biological dimer can be derived by crystallographic operations. This dimer is unique in that the highly conserved P13^A/B^(O)/A133^B/A^(N) intermolecular H-bond, connecting the αA helix of one protomer to the αGH loop of the other protomer, is missing (distance > 3.2 Å; Fig. 1E). The intermolecular T17^A/B^(N)–N134 ^B/A^(OD1) and T17^A/B^(O)–N134 ^B/A^(ND2) H-bonds and the D19^A/B^(OD1)–R27^B/A^(NH2) salt bridge are nevertheless conserved (Fig. 1E).

The P2₁^A^, P2₁^B^, and C2 protomers share similar overall structures (Fig 2A and Supplementary table 1), however their αC helices and inter-bundle loops (αGH and αCD) adopt distinct conformations (Supplementary Figs. S1, S2 and S3), shaping the carotenoid tunnel and modulating solvent access through channels 1 and 2 (SC#1 and SC#2, see Supplementary text 1). The P2₁^B^ protomer pairs the long, straight αC helix seen in most OCP structures with a coiled αCD loop where V53 lines the carotenoid tunnel wall (Supplementary Figs. S1, S2 and S3). Contrastingly, the P2₁^A^ and C2 protomers share a kinked αC helix, which instead positions M47 and I51 along the carotenoid tunnel wall upon folding of the N-terminal part of the αCD loop into a 3_10_ helix (Supplementary Figs. S1, S2 and S3). This conformational change strengthens interdomain contacts, substituting the A54(N)–W279(O) H-bond with T52(OG1)–P278(O) and T52(N)–W279(O) (Supplementary Fig. S2). The positioning of M47 and I51 nevertheless differs in the P2_1_^A^ and the C2 chains, altering the Y44–W41 distance that regulates access through solvent channel 2 (SC#2) (Fig. 2A and Supplementary Figs. S2 and S3). As a result, SC#2 is closed in the C2 chain (ring-to-ring distance ∼6.9–7.1 Å, depending on Y44(OH)–E115(OE1) H-bonding) but open in the P2₁^A^ (7.1 Å) and P2₁^B^ chains (7.4 Å) (Fig. 2A, B and Supplementary Figs. S1, S2 and S3). Differences between the three protomers raise the question as to which best represents the structure of OCPᴼ in solution, *i.e.* unperturbed by crystal packing. We obtained this insight by cryo-electron microscopic analysis of a constitutively-dimeric mutant with a disulfide bond at the dimer interface (A23C; Supplementary Fig. S4 and S5, Supplementary table 2, Supplementary text 1 and Materials and Methods). We found that the cryo-EM structure is most similar to the P2 ^B^ conformation (Supplementary Fig. S4).

Relationships between the various crystal forms were further investigated by RT crystallography. We found that while the P2_1_ and C2 crystal forms appear spontaneously in crystallization setups, the P2_1_ form can also be transformed into the C2 form by (controlled) dehydration (Supplementary Fig. 6A, B, Supplementary table 3, Supplementary text 1 and Materials and Methods), with higher yield when increasing the pH from 5.0 (crystallization conditions) to 7.5 (as in biochemistry studies). Unexpectedly, depending on the dehydration conditions, a new conformation of the C2 structure appears, that is characterized by a conformational change of the αGH loop, resulting in a closing of the carotenoid tunnel at its NTD end (SC#1) (Fig. 2A, B). This conformation—which cannot be accessed in the P2_1_ crystals regardless of experimental conditions (Supplementary Fig. 6C and Supplementary text 1) — blocks the path taken by the carotenoid to reach the OCP^R^ conformation (Fig. 2A and Supplementary Figs. 6D and 7). We refer to this structure as the ‘closed’ C2 (C2c) conformer in contrast to the ‘open’ C2 (C2o) structure in which only SC#2 is closed (Fig. 2A). The defining features of the C2c conformer are ruptures of the conserved intramolecular Q79(OE1)–A123(N) and D35(OD1)–Y129(OH) H-bonds, and the folding of the αGH loop into a 3_10_-helix, which borders SC#1 (Figure 2A, B and Supplementary table 4). This conformational rearrangement exchanges I125 for P126 as the residue laying atop the W41 side chain, and “plugs” P126 into the carotenoid tunnel (CB is 4.0 Å from the carotenoid C06), thereby closing SC#1 (Supplementary Fig. S7 and Supplementary table 4). The associated rupture of the D35(OD1)–Y129(OH) H-bond affords a 180° change in Y129(χ1) that results in placement of its aromatic ring behind P126, double-locking SC#1 (Figure 2A, B, Supplementary Fig. S7 and Supplementary table 4). Since P126 and Y129 were shown previously to be critical for OCP^R^ stability^4^, we asked about the functional relevance of the C2c conformation, specifically whether it might correspond to a persistant population of OCP^1hv^ intermediate in CAN-OCP.

#### Structural assignment of the OCP^1hv^ intermediate to C2c

An implication of the hypothesis that the C2c conformer structurally corresponds to the OCP^1hv^ intermediate is that it should accumulate upon illumination of C2o. We tested this prediction by illuminating C2 macrocrystals in the C2o conformer (100% occupancy) with 537-nm light for different times and determined the resulting structures by scalar structure factor amplitude extrapolation^22,23^

(Supplementary text 2 and Supplementary table 1). We found that irrespective of the functionalizing carotenoid (CAN or ECN), the C2c conformer appears upon 1 min illumination (Figure 3A and Supplementary Figs. S8 and S9), but its occupancy decreases upon prolonged illumination (Supplementary text 2). Thus, the unexpected closing of the carotenoid tunnel (via SC#1) in the C2c conformer is not only light-inducible, but transient, temporarily shielding the β2 ionone ring from the solvent before opening again – as required for the translocation of the carotenoid into the NTD^4^ in OCP^R^ (Fig 2A,B). This result supports the notion that the C2c conformer represents a functional intermediate along the OCP photocycle.

**Figure 3:**
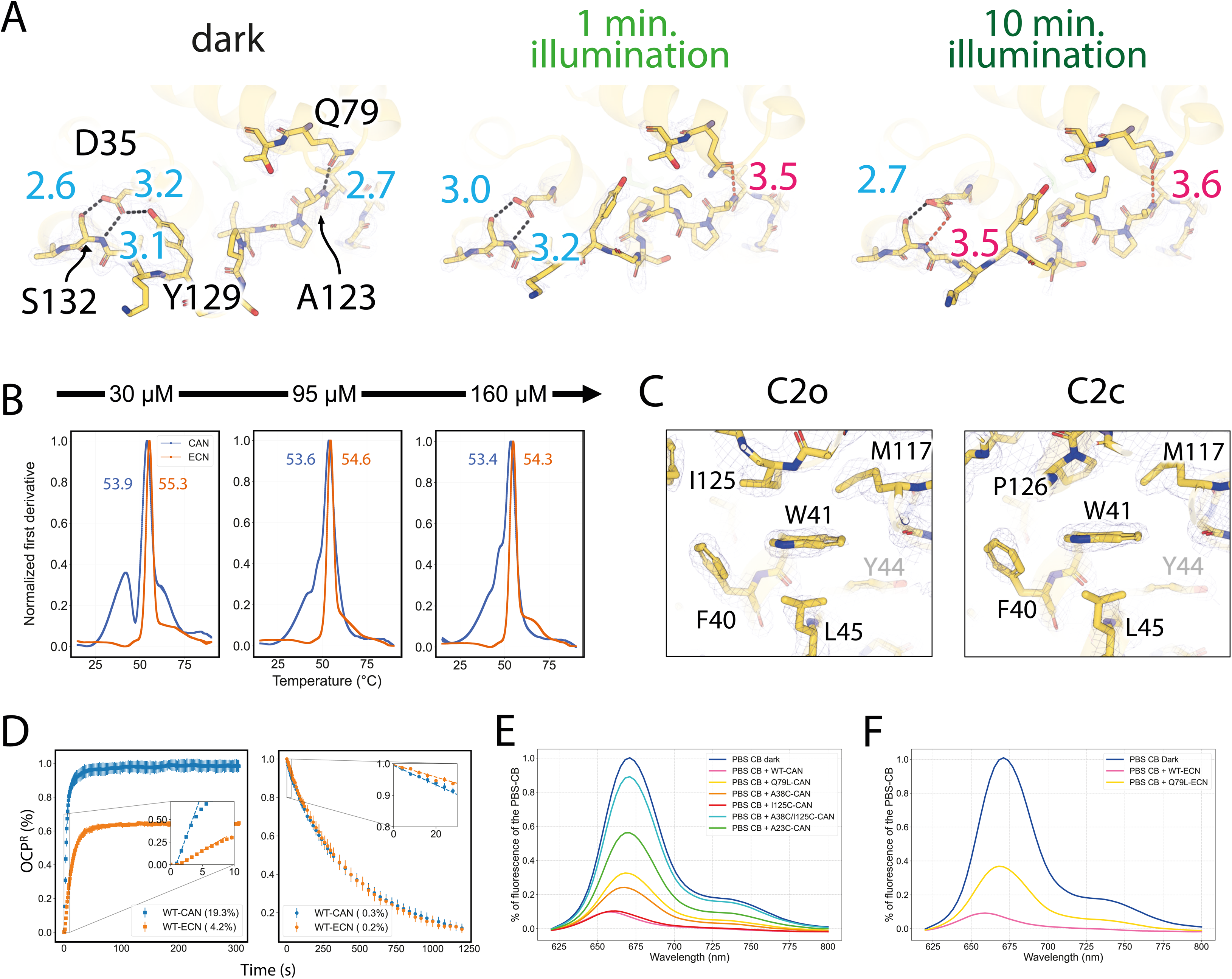
The C2c conformer is an obligate photo-intermediate of which a persistent population exists in CAN- OCP. (A) The C2c conformer is observed upon ≥ 1 min illumination of C2o macrocrystals. Numbers indicate interatomic distances up to (blue) and above (magenta) 3.2 Å, respectively. (B) The normalized first derivative of temperature-controlled scanning fluorimetry (TCSF) data reveals the concentration-dependent transition between two OCP conformers, in CAN-OCP (blue). This transition, which is not visible in the ECN-OCP (orange) nor CAN-A23C (Supplementary fig. S19), could be assigned to structural changes occurring in the vicinity of W41 (Supplementary fig. S11). As the W41 environment changes upon the C2c to C2o transition, favored by dimerization upon formation of the P13—A133 H-bond (Fig. 1E, F, Supplementary fig. S19), we assign the concentration-dependent TCSF peak to the partial unfolding of the closed C2c conformer, representing the OCP^1hν^ intermediate, into a more stable P2_1_^B^/P2_1_^A^/C2o conformer(s), representative of OCP^O^. This refolding occurs a few degrees before denaturation sets in, suggesting a high thermal barrier between the two states. (C) Local environment around αC residue Trp41 in the C2o and C2c state. The aromatic ring of W41 serves as a sliding base for the αGH loop, which either positions the I125 (C2o) or P126 (C2c) side chain over it. (D) The higher photoactivation rate of CAN-OCP could originate from the presence of a persistent population of the OCP^1hν^ intermediate, allowing formation of OCP^R^ by parallel single-photon and two-photon absorption mechanisms. We note that unlike data published earlier ^18^, the thermal recovery of CAN- and ECN-OCP is similar. This is due to shortening of the his-tag placed at the N-terminus of the protein in the present work. (E, F) Quenching of the core base of the phycobilisome CAN- (E) and ECN-(F) functionalized wild-type *Planktothrix aghardii* OCP and variants designed to test mechanistic hypotheses. The A38C-I125C mutant, covalently locked in the opened state, is not functional.

To probe the functional role of the C2c conformer, we designed mutants that preclude formation or stabilize this structure via disulfide links or H-bond modifications, respectively, determined their structures by RT crystallography (Supplementary table 5), and assessed their properties using temperature-controlled scanning-fluorimetry (TCSF) and spectroscopy (Fig. 3B-F and Supplementary Table 6). Both ECN and CAN complexes were investigated as detailed in Supplementary figs S10-S16, Supplementary texts 3 and 4, and Supplementary table 6. In TCSF experiments, only CAN-OCP showed a temperature and concentration-dependent peak (∼40° C for the monomer, ∼47° C for the dimer; Fig. 3B, Supplementary fig. S10, Supplementary Table 6, and Supplementary text 4), that the W41F variant links to structural rearrangements around W41 (Supplementary fig. S11). Given that the local environment of W41 markedly differs in the C2o and C2c conformers (proximity to I125 or P126, respectively) (Fig. 3C, Supplementary fig. S7 and Supplementary table 4), we attribute this signal to the ‘melting’ of the C2c conformation. The absence of this feature in ECN-OCP strongly supports the hypothesis that the C2c conformer corresponds to the metastable OCP^1hv^ intermediate, which is not present in ECN-OCP^13^ (Fig. 3D), explaining why the latter photoactivates via a strict two-photon mechanism^13^ (Fig. 3D and Supplementary Table 6). Demonstrative of the functional relevance of the C2c conformer is the observation that both CAN- and ECN-OCPs are inactive (Supplementary fig. S12A-C) – and therefore uncapable of quenching the PBS (Fig. 3E, F) – when the C2c formation is prevented by a disulfide bond in the A38C–I125C mutant, structurally locking SC#1 in the open conformation (P2_1_^B^, P2_1_^A^, C2o) irrespective of the illumination or dehydration conditions (Supplementary figs. S12A and S13, and Supplementary table 5). We investigated the importance of (temporarily) shielding the carotenoid from the solvent by engineering the A38C-I125C-T80W mutant, also unable to reach the C2c state but whose SC#1 channel can close with the aromatic side chain acting as a “surrogate” door. Photoactivity was partially restored in this protein, demonstrating the importance of shielding the carotenoid from the bulk to access the OCP^R^ state, thereby underpinning the requirement that the C2c state forms (Supplementary figs. S12 and S13 and Supplementary tables 4 and 6). Accordingly, destabilizing the SC#1-open form (P2_1_^B^, P2_1_^A^, C2o) by introduction of the Q79L or D35T mutations (Supplementary figs. S14A-H and S15) does not abolish photoactivity or PBS quenching (Fig. 3E,F). The initial photoactivation rate is similarly reduced in both mutants (Supplementary figs. S14C, G and Supplementary table 6), confirming the functional role of the Q79–A123 and D35–Y129 H-bonds. However, their recovery rates differ (Supplementary figs. S14D, H and Supplementary table 6), consistent with the respective presence or absence of these H-bonds in OCP^R^.

The recovery is hardly changed in the D35T variant (Supplementary fig. S14H), consistent with switching of the D35—Y129 H-bond present in OCP^O^ by the E34—Y129 H-bond in OCP^R^ (Fig. 2A). By contrast, the Q79L mutant exhibits a significantly increased recovery rate, consistent with the presence of the Q79—A123 H-bond in both OCP^O^ and OCP^R^ and the expected negative impact on OCP^R^ stability. Molecular dynamics simulations additionally show increased stability of the C2c conformer in the Q79L mutant compared to wild-type (Supplementary text 1 and Supplementary figs. S16-18), further explaining the dramatically increased recovery rate (Supplementary fig. S14D and Supplementary table 6). We last investigated the importance of the P13^A/B^(O)–A133^B/A^(N) hydrogen bond, broken in the C2o and C2c conformers (Figs. 1E and 2C), by engineering an A133P mutant, which showed a ∼10-fold reduction in dimer affinity (Fig. 1F). The mutation has little effect on photoactivation, recovery, or the dark-state crystal or cryo-EM structures (Supplementary fig. S14I-L and Supplementary table 2).

Altogether, these results underscore the functional relevance of the C2c conformer and support its assignment to the OCP^1hv^ intermediate. They also firmly support that at RT, CAN-OCP populates both the OCP^O^ state and the OCP^1hv^ intermediate (Supplementary texts 1, 2, 3 and 4).

#### Observing photo-triggered changes in CAN-OCP by time-resolved crystallography

Because our crystal structures represent either OCP^O^ isoforms (P2_1_^A^, P2_1_^B^, C2o) or the OCP^1hν^ intermediate (C2c) (Fig. 2A), they provide starting points to obtain atomic information on how excitation of the dark-state(s) yield(s) the OCP^1hν^ intermediate (P2_1_ crystals), as well as how excitation of the latter yields the OCP^R^ state (C2 crystals). Thus, we performed time-resolved serial femtosecond crystallography (TR-SFX) experiments using a mixture of the two crystal forms to probe photo-triggered structural changes occurring upon photoexcitation of OCP^O^ and the OCP^1hν^ intermediate, respectively (Supplementary figs. S20-S23 and Supplementary text 5). The experiments were performed at SwissFEL using a pump-probe scheme. For injection into the XFEL beam, OCP_CAN_ microcrystals were embedded in lipidic cubic phase (LCP) (Supplementary figs. S20A, B), resulting in a red shift of their absorption spectrum (Supplementary fig. S21D). Nevertheless, no structural changes occur in the P2_1_ and C2 protomers, compared to equivalent structures solved in absence of LCP (Supplementary fig. 24). Moreover, the photodynamics are the same in solution and microcrystalline OCP (Supplementary figs. 21-23 and Supplementary text 6).

The time delays between a 535 nm pump and an X-ray probe pulse were chosen according to the lifetimes of the photocycle intermediates derived by transient spectroscopy. It has been shown that following photoexcitation of the carotenoid to the optically allowed S_2_ state (lifetime ∼150 fs), it decays to one of three excited-states: a sub-ps lived intramolecular charge-transfer (ICT) state (lifetime ∼0.5 ps), a ps-lived mixed S_1_/ICT state called S_1_ (lifetime ∼2.5 ps)^6^, and a state referred to as S* characterized by a lifetime of several ps (∼7.5 ps) and presumed to display a distorted geometry^24^. It is debated whether these excited states form in parallel or serially, and thus which of them is the precursor of P_1_ (lifetime ∼50 ns) – the first of several red-shifted intermediate(s) (P_2_ ± P_2’_, P_3_, P_N_, P_M_, P_X_) on the path to OCP^R^ ^18,24,25^ that all share the characteristic of broken H-bonds between the carotenoid and the protein^26,27^. In a first experiment, focused on short time delays, diffraction data were collected at 0.3, 1.5, 10 and 100 ps after optical-excitation with a 85 fs pulse (1.5 mJ/cm^2^, 18 GW/cm^2^, Supplementary tables 7 and 8, Supplementary text 5, and Materials and Methods). The use of such a low power density, corresponding to 0.4 absorbed photon per carotenoid on average, avoids formation of a long-lived, off-pathway radical-state^9^ (Supplementary figs. S21-S23, Supplementary text 6, and Materials and Methods). In a second TR-SFX experiment, diffraction data were collected 10 ns and 1 µs after photoexcitation using 3.7 ns optical pulses at 17 mJ/cm^2^ (4.6 MW/cm^2^), corresponding to a maximum of nominally five absorbed photons per chromophore (Supplementary tables 7 and 8, Supplementary text 5, and Materials and Methods). Electron density maps calculated from extrapolated structure factor amplitudes were interpreted with a single structure, assuming that at all probed time-delays, the state with the highest occupancy dominates the features. In the first experiment, 95-99 % of the crystals indexed in C2 (100 % C2c), the rest in P2_1_, limiting the description of structural changes in the CAN pigment to the C2 data (Supplementary tables 7 and 8). In the second experiment, however, equal distributions were obtained for the two crystals forms, and the C2 crystals contained a mixture of the C2o and C2c (60%) conformations (Supplementary tables 7 and 8). However, upon photoexcitation, only the C2c conformer was observed in extrapolated maps (see Supplementary text 5).

#### Photoactivation of the dark-state induces rapid and short-lived conformational changes

TR-SFX data collected on the P2₁ crystals report on the outcome of photoexcitation of the dark OCP^O^ state. Essential features are highly similar in the P2_1_^A^ and P2_1_^B^ protomers although the changes in the pigment and protein differ in detail, as expected from the differences in their ground-state carotenoid tunnel architecture (V53 in P2_1_^B^ is replaced I51 and M47 in P2_1_^A^) (Supplementary figs. S1, S2, S3 and S24). At 0.3 ps, both chains appear to exhibit concerted twisting of the carotenoid polyene that is centered at the C7′–C6′ and C9′–C8′ bonds in P2_1_^A^ (Supplementary figs. S25 and S27-S29), and at C10′=C9′ and C12′=C11′ in P2_1_^B^ (Supplementary figs. S26 and S27-S29). This difference originates from the I51(CD1) methyl group, which is in close proximity to both the C11′ and C9′ methyl group of the carotenoid, preventing their movement in P2_1_^A^ (Supplementary fig. S27). By 10 ps, however, a similar 7-*cis* photointermediate appears to form in both protomers upon twisting around the C6–C7 and C10–C11/C12-C13 (P2 ^A^/P2 ^B^) bonds (Fig. 4 and Supplementary figs. S27), as predicted by excited-state nonadiabatic molecular dynamics simulations^28^. The photo-induced twisting of the carotenoid leads to the sequential rupture of H-bonds to Y203 and W290 at 0.3 ps and 10 ps, respectively, with reformation of at least one H-bond by 100 ps concomitant with relaxation of the carotenoid polyene (Figs. 4 and 5A, and Supplementary fig. S30). Thus, the carotenoid is fully untethered from the CTD at 10 ps, but re-attached at 100 ps (Fig. 4 and 5A). This observation is consistent with both a spectroscopic return to the ‘orange’ state by 100 ps^18,24,25^.

**Figure 4:**
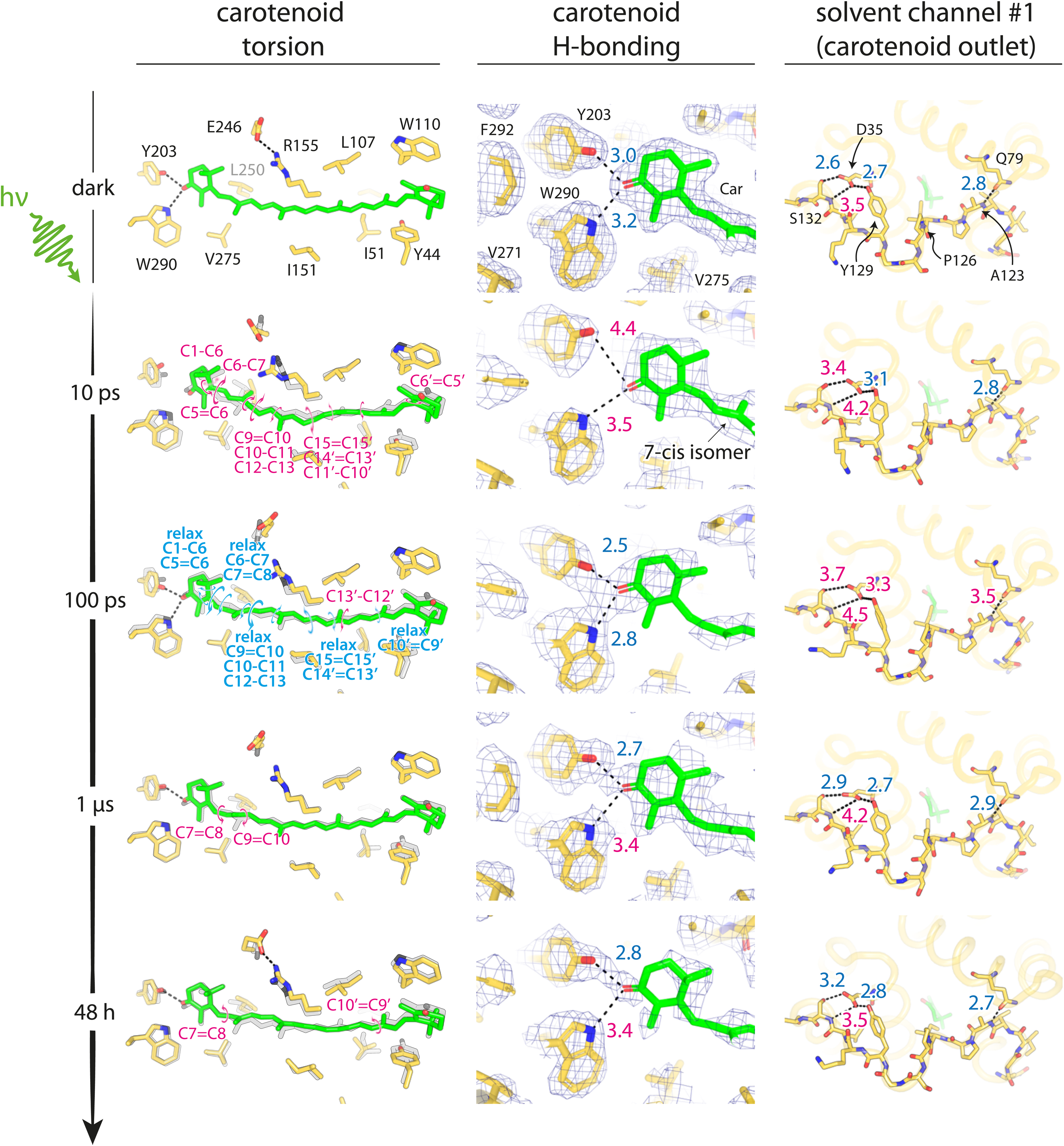
The photo-induced formation of a 7-cis isomer coincides with ultra-fast carotenoid untethering in the dark OCP^O^ state. Results are shown for the P2 ^B^ conformer, most representative of the dark OCP^O^ state. Carotenoid torsion and H-bonding to CTD residues, as well as the αGH loop conformation observed at SC#1, are shown at select time delays across the TR-SFX (10 ps – 1 µs) and serial synchrotron crystallography (SSX) (48h illumination) series. In the left panels, torsions occurring in the carotenoid with respect to the dark state (shown in grey for all time delays) are highlighted by magenta circular arrows whereas relaxation is shown in blue. In the middle panels, electron density is shown at 1 σ for the dark state (conventional 2mFo-DFc map) and the photo-intermediates (extrapolated 2mFextr-DFc maps) captured across our TR-SFX and SSX data series. The right panels focus on H-bonding interactions tethering the αGH loop to the αC and αE helices, as well as on the fold of the αGH loop. Full carotenoid untethering is observed at 10 ps, upon formation of a 7-*cis* isomer, but H-bonds reform upon relaxation of this state at 100 ps and concomitant release of the αGH loop from interactions with αC (D35 mediated H-bonds to Y129 and S132) and αE (Q79 H-bond to A123). Due to crystal packing no refolding occurs. At longer time delays, we observe a single H-bond tethering the carotenoid to the protein, which is known to associate with an orange spectrum. In the steady state structure (P2 ^light-48h^), the αGH loop is released from interaction with αC, as observed in the 100 ps structure.

**Figure 5:**
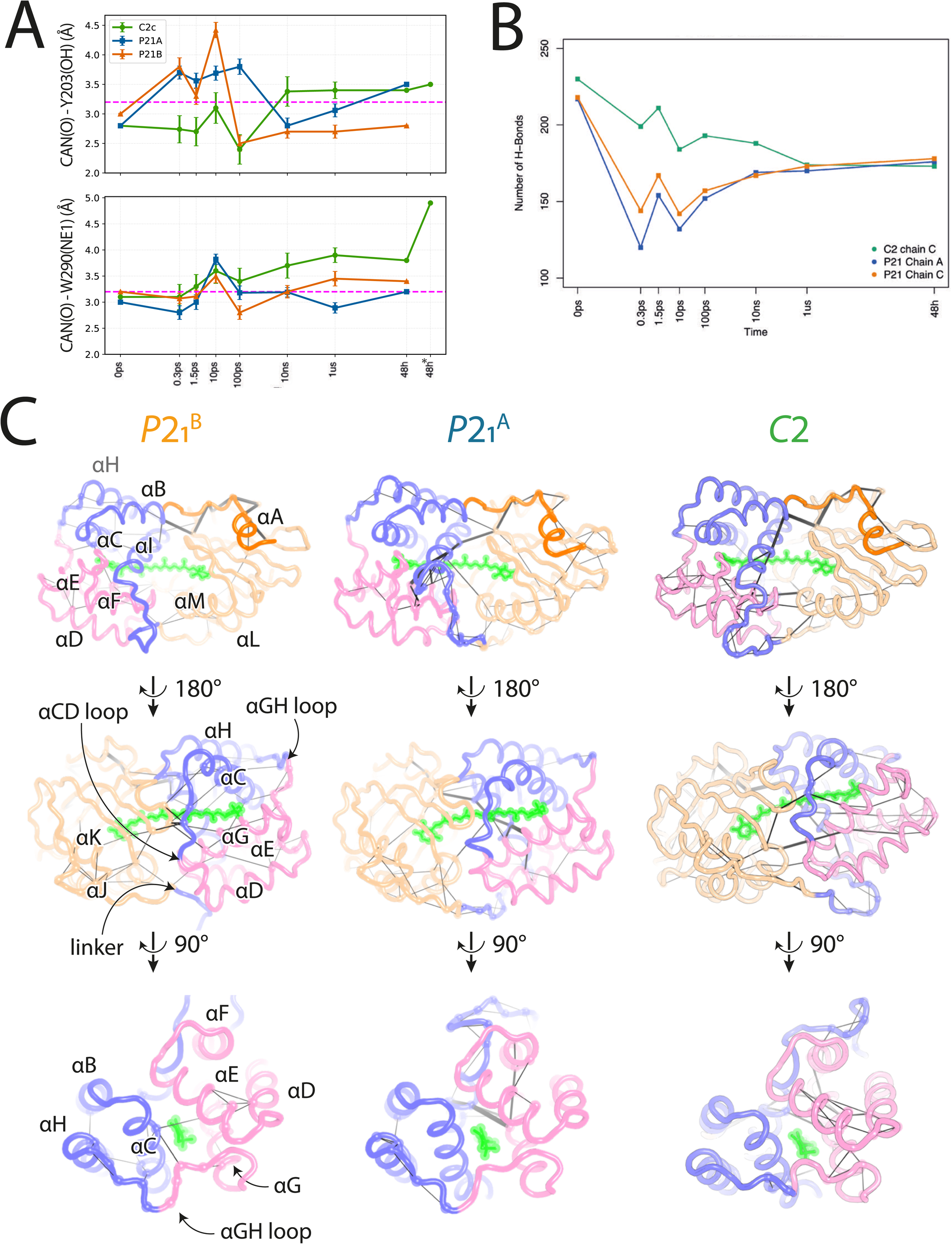
Carotenoid twisting and untethering from the protein sterically propagate to the rest of the scaffold. (A) Canthaxanthin H-bonding to CTD residues Tyr203 and Trp290 in the P2_1_ ^A^, P2_1_ ^B^ and C2c conformers, at all probed time delays across the TR-SFX (0.3 ps – 1µs) and SSX (48h illumination) series. Mind that H-bonding is reported for under the label ‘48h’ for the C2^light-48h^ structure, P2 ^A-light-48h^ and P2 ^B-light-48h^ structures obtained by extrapolations methods, while that for the C2new^light-48h^ structure is reported under the label ‘48h*’. (B) A substantial loss of H-bonding occurs upon photo-excitation of the P2_1_ ^A^, P2_1_ ^B^ and C2c conformers. This loss is ultra-fast in the P2 chains, but slower in the C2c chain. Irrespectively, the three chains feature 75% of their initial (dark-state) H-bond content at the 1 µs time delay, suggesting that photoexcitation primes OCP for conformational changes in the αCD and αGH loops by an overall loosening of the scaffold. (C) A ComPASS analysis, integrating dynamical and interaction-based data into a network framework, reveals how energy propagates through specific residues and structural elements, providing mechanistic insight into long-range allosteric effects. Similar key communication routes are identified in the three chains, propagating the excitation signal from the β-sheet (W290, R291, D306) to the αBCHI to the αDEFG bundle.

Besides rupture of the H-bond to the carotenoid, phototriggered conformational changes occur at W290 (Fig. 4 and Supplementary figs. S25-S27 and S30), which transmit to adjacent R291, resulting in rupture of the H-bond between its guanidinium group and the carbonyl of R9 (Supplementary figs. S31 and S32). This triggers the rupture of the P13^A/B^–A133^B/A^ H-bonds, as early as 0.3 ps and up to 1 µs. Perturbations of both the T17^A/B^–N134 ^B/A^ interactions and the D19(OD2) ^A/B^–R27(NE) ^B/A^ salt bridge also occur on the ∼0.3 ps - 1 µs timescale, supporting transient monomerization of the crystalline OCP^O^ (Supplementary figs. S31 and S33). On the protomer level, photoinduced changes in the carotenoid sterically propagate to tunnel residues (L37, I40, L107, W110, R155, L250, W279, F280), inducing strain within the αBCHI (notably L39, L139, I142, L154) and αDEFG (notably in I71, M83, L86, I94, L113, M117) bundles which further weakens the contacts stabilizing the αGH loop (Supplementary fig. S31 and S34). In P2 ^B^, the loss of the D35(OD2)–S132(OG1), D35(OD1)–Y129(OH) and Q79(OD1)–A123(N) interactions results in complete release of the αGH loop at 100 ps (Fig. 4 and Supplementary fig. S35), while in P2 ^A^, rupture of D35-mediated contacts occurs on the 1.5 to 100 ps timescale (Supplementary fig. S35). Thus, in both P2_1_ chains, two pathways of signal propagation coexist from the photoexcited carotenoid to the αGH loop, priming them for transition to the C2c conformer from 100 ps to 1 µs: one at the protomer level, whereby changes in the carotenoid structure result in loss of the D35 and Q79-mediated intramolecular H-bonds (Fig. 4 and Supplementary figs. S35-S37); and one at the dimer level, whereby transient changes in the carotenoid H-bonding pattern triggers loss of the P13^A/B^–A133^B/A^ H-bonds (Supplementary figs. S31 and S32).

We note that structures determined at early time delays show substantial loss of hydrogen bonds and secondary structure in both chains (Figure 5B). By 10 ns, however, about 75% of these are restored. Thus, a main feature of the photoexcitation of OCP^O^ is a partial refolding of the αGH loop occurring on a 100 ps–1 µs timescale.

#### Photoexcitation of the OCP^1hν^ intermediate results in prolonged lifespan of the untethered carotenoid

The C2c protomer, representing the OCP^1hν^ intermediate, is characterized by a fully shielded carotenoid tunnel and loss of the intermolecular P13^A/B^ –A133^B/A^ H-bonds stabilizing the αGH loop (Figs. 1 and 2). These characteristics affect the structural consequences of photoexcitation (Supplementary figs. S28-S35). For the 0.3 - 1.5 ps time delays, we observe ultra-fast and rapidly-relaxing twists at C6-C7^17^ and C7’-C6’, connecting the β-ionone rings to the polyene, and at C13’-12’ and C10’=C9’, in the vicinity of I51 and Y44 (Supplementary figs. S28, S29 and S38). Additionally, photoinduced conformational changes occur in the β1 ring and its vicinity. Specifically, at 0.3 ps, a combination of twists at C5=C6 and C4-C5 results in a bend at C5, with the attached methyl pointing towards L252 (Supplementary fig. S38). Tunnel residues from which the carotenoid moves away display the largest structural changes, enabling them to maintain contact (e.g., L37, I40, I51, M207, L250, and L252), while those approached (∼0.5 Å) by the carotenoid show no difference (e.g., L37, L107, R155, and V158) (Supplementary fig. S39). As a result, the αBCHI and αDEFG bundles of the NTD shift towards one another and around the carotenoid, allowing steric propagation of the photoexcitation signal from the tunnel-lining (αC, αE, αG, αI) to the protein surface helices (αB, αD, αF, αH) – initially bypassing the αCD and αGH loops (Supplementary fig. S39). A notable marker of this propagation is the change in volume of hydrophobic cavities in each of the two bundles (Supplementary fig. S34); *i.e.* those lined by I71, M83, L86, I94, L113 and M117 in the αDEFG bundle, and by L39, L139, I142 and L154 in the αBCHI bundle (Fig. 1D). It is through these contacts that as early as 0.3 ps, M83(CE) approaches the β2 ionone ring, further blocking the SC#1 outlet (Supplementary fig. S34), while Q79 nears and transiently H-bonds to A123 (Supplementary fig. S35), reproducing a feature of the OCP^O^ state(s) (P2_1_^A^, P2_1_^B^, C2o). These changes revert on the 1.5 to 10 ps timescale, as dihedral flips at C4–C5 and C5=C6 reorient the C5 methyl towards I305, at 1.5 ps, and then back toward L252, at 10 ps (Supplementary figs. S28, S29 and S38). Concurrently, a bicycle-pedal (BP)-like isomerization occurs at C13=C14, C10’=C9’ and C9’–C8’ (∼45–60°) which moves the β1-ionone ring away from W290 (Supplementary figs. S28, S29 and S38). This leads to rupture of the W290 H-bond to the 4-keto oxygen at 10 ps, which remains broken up to 1 µs (Fig. 6 and Supplementary figs. S30 and S38). The BP-like isomerization (Fig. 6 and Supplementary figs. S28, S29 and S38) induces expansion of the hydrophobic cavities within the αBCHI and αDEFG bundles (Supplementary fig. S34), as well as changes in the αGH (Fig. 6 and Supplementary figs. S34, S35, S39 and S40) and αCD loops (Supplementary Fig. 41). Notably, the αGH loop shifts away from the adjacent αC, αD, αE, and αF helices (Supplementary figs. S35 and S39). The β2 ionone ring remains unchanged at this time delay, but the distance from the 4’-keto oxygen (O’) to L37(CD2) increases to 3.7 Å– the largest distance in the C2c TR-SFX series (3.2-3.4 Å in the starting OCP^1hν^ intermediate and at other time delays). By 100 ps, a twisting occurs at C7’-C6’ (Δ∼85°), reminiscent of the changes observed at 0.3 ps, upon relaxation at C13=C14 and redistribution of strain from C9’–C8’ (Fig. 6 and Supplementary figs. S28, S29 and S38). However, torsion is preserved at C10’=C9’. Likewise, no change is seen at the β1 ionone ring, where the H-bond to W290 remains broken and the 4-keto group planar (Fig. 6 and Supplementary figs. S30 and S38). Nevertheless, an “inversion” occurs at the C4’ bend, shifting the O’ of the β-ionone 2 ring toward W110 (−0.4 Å to Trp110 CH2/CZ3) instead of Ile40 as in the starting dark state C2c structure and at other time delays (Fig. 6 and Supplementary figs. S38). These changes are concomitant with a partial re-opening of SC#1, whereby P126 “unplugs” from the tunnel outlet (the ring-to-ring distance between P126 and the β2-ionone ring peaks to 12.4 Å at 100 ps, compared to 6.4 ± 0.1 Å in the starting dark state C2c structure and at other time delays) and D35 H-bonds to Y129 – reminiscent of the SC#1-open OCP^O^ state(s) (P2 ^A^, P2 ^B^, C2o) (Supplementary figs. S35, S38 and S39). The Q79/A123 H-bond does not form, however, nor does the αGH loop fully recoil, as would be required for a complete re-opening of SC#1.

Absence of these features likely underlies the return to the closed C2c state, at 10 ns and 1 µs (Supplementary figs. S35 and S38). Irrespective, rupture of the Y203 H-bond with the carotenoid is observed at 10 ns, resulting in full untethering of the pigment, concomitant with replanarization of the β1-ionone ring and restoration of the ground-state bend at the 4′-keto group (*i.e.,* with O′ oriented toward Ile40) of the β2-ionone ring (Fig. 6 and Supplementary figs. S30, S35 and S38). Along the polyene, the twist around the C7’-C6’ bond relaxes, and the only twist remains at C10’=C9’. Further relaxation of the polyene chain ensues at 1 µs, by concerted twists at C8-C9 and C10-C11 (Fig. 6 and Supplementary figs. S28, S29 and S38), rotating the methyl at C9 by 90° to fit in a hydrophobic groove contributed by L250, V275, T277, M286 and M288. Nevertheless, the carotenoid H-bonds to Y203 and W290 remain broken at 1 µs (Fig. 6 and Supplementary fig S30). Thus, in the C2 conformer representing the OCP^1hν^ intermediate, carotenoid untethering is achieved more slowly, but is prolonged by 5 decades in times compared to the P2_1_ conformer, representing OCP^O^ (Figs. 4, 5A and 6, and Supplementary figs. S30). The slowing of photo-triggered conformational changes is reflected in the gradual decrease of the protein’s H-bond content, which drops to ∼75% over 0.3 ps to 1 µs, contrasting with the observation made on P2_1_ chains (OCP^O^) (Figure 5B). We speculate that the delayed 10-ps bicycle-pedal-isomerization and its relaxation supply the energy that drives both carotenoid untethering, on the 10ps - 1µs timescale, and tunnel reopening on the ∼100-ps timescale (Fig. 6 and Supplementary figs. S35, S38 and S39). The latter timescale coincides with that over which αGH-loop release occurs, in the P2 ^B^ conformer representing OCP^O^ (Fig. 4 and Supplementary figs. S35 and S37).

**Figure 6:**
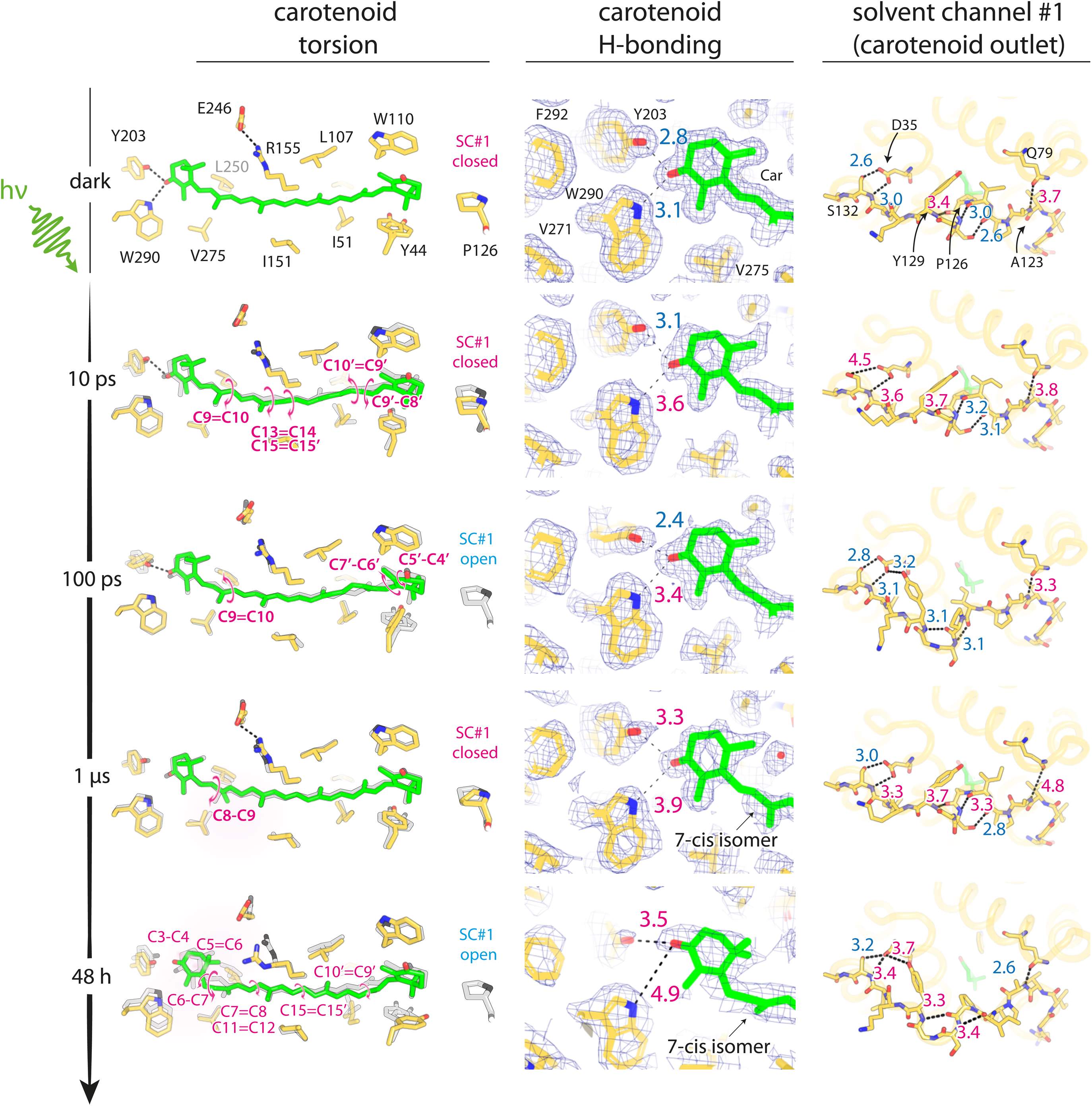
Insulation of the carotenoid in the OCP^1hv^ intermediate delays and extends carotenoid untethering by three and five decades in time, respectively. Carotenoid torsion and H-bonding to CTD residues as well as the αGH loop conformation observed at SC#1 are shown at select time delays across the TR-SFX (10 ps – 1µs) and SSX (48h illumination) series. In the left panels, torsions occurring in the carotenoid with respect to the dark state (shown in grey i for alltime delays) are highlighted by magenta circular arrows. In the middle panels, electron density is shown at 1 σ contour for the dark state (conventional 2mFo-DFc map) and the photo-intermediates (extrapolated 2mFextr-DFc maps) captured across our TR-SFX and SSX data series. The right panels focus on H-bonding interactions tethering the αGH loop to the αC and αE helices, as well as on the fold of the αGH loop. Rupture of the H-bond to W290 is observed at 10 ps, upon bicycle-pedal isomerization at C10’=C9’ and C9’-C8’. Restoration of the open configuration of the αGH loop ensues at 100 ps, upon relaxation of the C9’ isomer and transfer of torsion to C7’-C76’, but the recoiling is transient in that the C2c conformer is again observed at longer times delays. Rupture of the H-bond to Y203 occurs at 10 ns, upon relaxation of the latter bond, and lasts up to at least 1 µs. At this time delay, isomerization occurs at C8-C9, yielding a rotamer that fits its C9 methyl 90° apart, in a groove contributed by the side chains of L250, V275, M286, and M288. In the steady state structure (C2new^light-48h^), the carotenoid is still untethered from the protein but underwent isomerization at C5=C6, C6-C7 and C7=C8 upon relaxation at C8-C9, yielding a 7-cis isomer that differs from that observed at 10 ps in the P2_1_^A^ and P2 ^B^ structures but sees its C9 methyl fit in the same groove occupied in the 1 µs C2c structure. Alongside, the αGH has fully recoiled, with SC#1 present in the open state.

**Figure 7:**
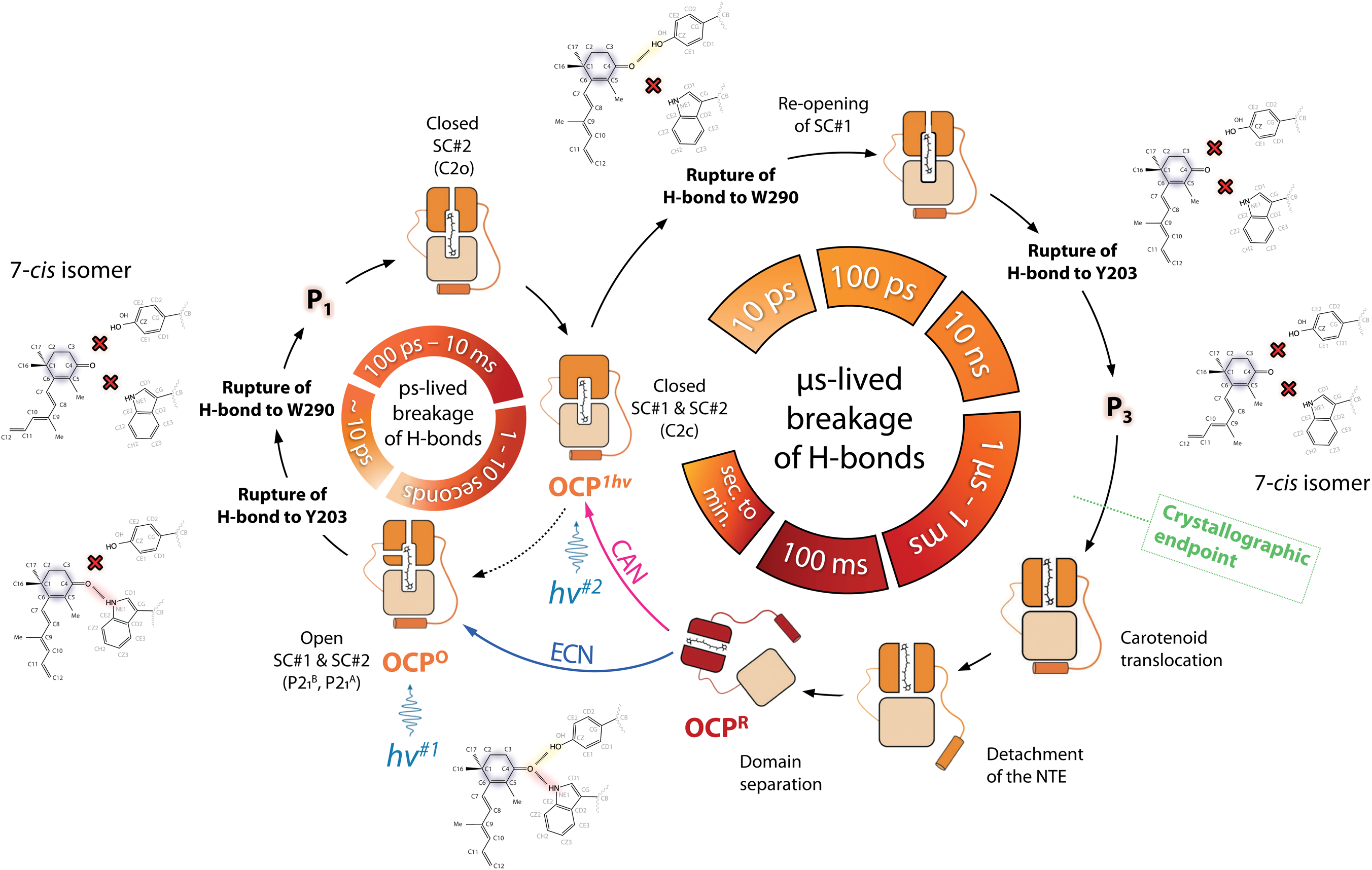
The OCP photocycle from a structural perspective. The SC#1- and SC#2- open dark OCP^O^ state, best represented by P2_1_^B^, shows large conformational change in the ultra-fast regime, but photoexcitation. Rupture of the two H-bonds tethering the carotenoid to the protein scaffold is achieved at 10 ps, upon formation of a 7-*cis* isomer. This state likely corresponds to the spectroscopic P_1_ state. The latter rapidly relaxes, enabling H-bonds of the protein to reset, while steric propagation of the photoexcitation signal fuels the two-step transition to the OCP^1hν^ intermediate, represented by the C2c conformer, upon successive closing of SC#2 (C2o) and SC#1 (C2c). Upon absorption of a photon by the OCP^1hν^ intermediate, conformational changes develop more slowly in the carotenoid, whose untethering from the protein is both delayed and prolonged by three and at least five decades in time, respectively. We speculate that the tighter confinement of the carotenoid in the OCP^1hν^ intermediate entails the succession of multiple-relaxation steps to fully defuse the absorbed energy. Through these, photoexcitation sterically propagates, fueling the re-opening of SC#1, as early as 100 ps, upon recoiling of the αGH loop (C2o). A 7-*cis* isomer of the carotenoid eventually forms, ready to translocate across the tunnel, which differs from the 7-cis isomer observed at 10 ps in the P2_1_^B^ (and P2_1_^A^) conformer(s) and could corresponds to the spectroscopic P_3_ state. Further steps on the pathway to OCP^R^ include complete translocation of the carotenoid across the tunnel, followed by dissociation and re-positioning of the two domains and monomerization. Likewise, it is presumable that monomerization would occur alongside domain dissociation in dimeric OCP (Fig. 3E, F, Supplementary fig. S19), None of these steps are possible in crystalline OCP. In absence of excited phycobilisomes, the OCP^R^ state reverts thermally to the OCP^O^ state. That ECN-OCP follows a strict two-photon absorption photoactivation mechanism implies that the OCP^O^ state is repleted upon recovery. This could occur either via the OCP^1hv^ intermediate, as microreversibility would suggest, or in a single OCP^R^ è OCP^O^ step, as supported by the mono-exponential recovery profile. That CAN-OCP features a population of the OCP^1hv^ intermediate implies that this state is repleted upon recovery, either as an intermediate along a two-step OCP^R^ è OCP^O^ recovery, or through a parallel pathway, as supported by the bi-exponential recovery profile.

#### Photoproduct under steady state illumination

We set out to determine the crystallographic endpoints of the OCP^O^èOCP^1hν^ and OCP^1hν^èOCP^R^ transitions and to this end, performed a serial synchrotron crystallography (SSX) experiment on OCP microcrystals suspended in crystal storage solution and kept either in the dark or illuminated for 48h by weak blue light (Supplementary fig. 20C, B, Supplementary text 7 and Materials and Methods). Illuminated crystals displayed a red spectrum, reminiscent of OCP^R^, while crystals kept in the dark retained a spectrum similar to OCP^O^ (Supplementary fig. S20D). The crystals contained 80% C2 crystals, and 20 % P2_1_ crystals (Supplementary table 9). Illuminated P2_1_ crystals were isomorphous to their dark-state counterparts and while only minor photoinduced conformational changes were observed (most concentrated at the αGH loop or the αA/β-sheet interface), the dimerization interface was disrupted in both chains, with both the P13^A/B^—A133^B/A^ H-bonds and the D19—R27 salt-bridge broken (Fig. 4 and Supplementary figs. S33 and S35-S37). Two C2 populations were present after 48 h illumination: one is isomorphous to the C2 dark crystals (C2c^light-48h^; 75% of crystals), whereas the other shows a 17% increase in unit-cell volume (C2new^light-48h^), essentially matching that of the P2₁ crystals while preserving the crystallographic dimer. In the C2new^light-48h^ crystals, however, the characteristic intermolecular P13—A133 H-bond is formed, again reminiscent of the P21 crystals and in stark contrast with both the dark and illuminated but isomorphous C2 crystals (Fig. 6 and Supplementary figs. S33 and S35).

The structure of the C2c^light-48h^ photo-triggered intermediate was extracted by extrapolation methods, while conventional crystallography was used to determine the structure of the C2new^light-48h^ intermediate (Supplementary table 9). The C2c^light-48h^ and the C2new^light-48h^ structures share the feature of broken H-bonds to the carotenoid (Fig. 5A). The carotenoid and protein conformations however differ markedly in the two structures (Supplementary figs. S28 and S29). In the C2c^light-48h^ structure, the carotenoid conformation closely resembles dark state notwithstanding the presence of the bend at C4’, reminiscent of the TR-SFX C2c structures determined at 10 ns and 1 µs time delays (Supplementary figs. S28 and S29). At the protein level, changes are also minor (Supplementary fig. 39).

By contrast, large conformational changes are observed in the C2new^light-48h^ structure, at both the protein and carotenoid level (Supplementary text 7). Changes in torsion angles are seen at C6–C7 (Δ∼65°), C7=C8 (Δ∼95°), C10–C11 (Δ∼40°), and C11=C12 (Δ∼65°), along with ∼30° bending at C3–C4 and C4–C5 (Fig. 6 and Supplementary figs. S28, S29 and S38). These changes are concomitant with the β1-ring 4-keto oxygen (O) moving away from Y203(OH) and W290(NE1) (3.5 and 4.9 Å distance, respectively) (Fig. 6 and Supplementary figs. S30), in the direction of I305 (3.2 Å distance to CD1), and with the positioning of the C9 methyl into the groove already occupied in the 1-µs C2 structure (formed by L250, V275, T277, M286, and M288). They also bring the C9′-centered terpene unit closer to Y44 (Fig. 6 and Supplementary fig. S38), which coincides with the reopening of SC#1 through a re-coil of the αGH loop (Supplementary figs. S35 and S39) and a re-folding of the αCD loop (Supplementary figs. S39 and S41). Briefly, a ∼30° change in Y44(χ1) positions its aromatic ring against Trp41, forcing the displacement of P126 required for coiling the αGH loop. This fully re-coiled αGH loop conformation is locked-in place by the open-state specific P13/A133 and Q79(OD1)–A123(N) H-bonds, but the D35 mediated H-bonds to Y129 (OH) and S132(OG1) are broken, as expected for a photo-intermediate *en route* to OCP^R^ (Fig. 6 and Supplementary fig. S35). Additionally, the sliding of Tyr44 toward Trp41 liberates space for M47 and I51 to adopt alternative rotamers, coinciding with the αCD loop adopting a P2₁^A^-like conformation (Supplementary fig. 41). Changes in the αCD and αGH loops propagate into the αBCHI and αDEFG bundles, which remodel as their bordering helices shift (Supplementary figs. S34 and S40). Consequently, at the domain level, the NTD–CTD opening angle increases by ∼4°, and most interdomain contacts (4/6), including the R155–E246 salt bridge, are broken. The dimer interface nevertheless remains intact (Supplementary fig S33).

The C2c^light-48h^ structure could bridge the 1-µs C2 and C2new^light-48h^ conformations, requiring two BP-isomerization steps. Alternatively, C2c^light-48h^ could be off-pathway, and the C2new^light-48h^ structure could be reached directly upon relaxation of the 1-µs intermediate. Based on the location of the C9-methyl and adjacency of the twisted bonds, we favor the second hypothesis, where torsional relaxation at C8–C9 (Δ∼120°) initiates the series of twists seen at C5=C6 (Δ∼45°), C6–C7 (Δ∼75°), C7=C8 (Δ∼120°) and C11=C12 (Δ∼50°). Future computational studies could address this issue.

### Conclusion

OCP serves as a biological circuit breaker, protecting the photosynthetic apparatus of cyanobacteria from the damaging effects of strong sun light. To this end, OCP uses a two-photon mechanism^13,11^ (Fig. 1A). The first absorbed photon generates a meta-stable intermediate, OCP^1hν^, whose lifetime matches the intensity threshold required for photoprotection—that is, the time window in which a second blue-green photon must arrive. This lifetime is on the order of seconds; thus, if no second photon is absorbed, OCP^1hν^ reverts thermally^13^ to OCP^O^. Absorption of a second photon by OCP^1hν^ leads to formation of the biologically active, photo-protective OCP^R^. While isomerization or covalent modifications are commonly found in relatively long-lived photo-intermediates, they do not match the prolonged life time of OCP^1hν^. Thus, so far, the basis of OCP^1hν^ stabilization was a mystery. Here we identify the structure of the OCP^1hν^ intermediate (Fig. 2A) and show that the basis of its longevity and meta-stability is the light-induced refolding of a loop in the vicinity of the carotenoid pigment. We used TR-SFX to capture the first light-induced reaction intermediates (0.3 ps–1 µs) along the OCP^O^ è OCP^1hν^ and OCP^1hν^ è OCP^R^ transitions (Figs. 4, 5, 6 and Supplementary figs. S20-S41). Further, we determined the crystallographic “endpoint” of the OCP^1hν^ è OCP^R^ photoreaction, imposed by crystal lattice constraints. The steady state endpoint along the OCP^1hν^ è OCP^R^ transition, characterized after 48-h illumination, could correspond to the P_3_ intermediate, previously suggested to feature changes in helicity upon initiation of carotenoid translocation, on the basis of TR mid-IR spectroscopy^24^ (Supplementary fig. S1). Complementary analysis of mutants designed to test key mechanistic hypotheses (Supplementary figs. S10-S15 and S18-S19) strongly supports that the crystallographic endpoints lie on the pathway to the corresponding functional states, *i.e.*, OCP^1hν^ and OCP^R^, respectively. Thus, despite the fact that our structural insights remain limited to the early steps of the two photon-induced transitions due to lattice constraints, our data support that both transitions involve photoinduced structural changes in the αCD and αGH loops, occurring 100 ps post-absorption of the first or the second photon, respectively, enabling the closure or the opening of the carotenoid tunnel outlet (Supplementary figs. S36, S37, S39). A per-chain comPASS analysis^29^ (Supplementary text 8) of the time-resolved series identifies a common signal-transduction pathway in all three protomers and most marked in the P2_1_^B^ and C2c chains, from the β-sheet through the αBCHI bundle to the αDEGF bundle, such that the largest motions arise in the αCD- and αGH-loops of each chain (Fig. 5C and Supplementary figs. 36, 37 and 39). Our study uncovers a highly-counterintuitive structural mechanism underlying the two-photon activation of OCP, where the protein shields the chromophore binding pocket upon absorption of the first photon, to then re-open upon absorption of the second photon, enabling carotenoid translocation (Fig. 7). Our data reveal that depending on whether OCP^O^ (P2_1_ conformers) or the OCP^1hν^ intermediate (C2c conformer) is excited, the carotenoid is found in a markedly different conformation at 1 µs: it is planar and reconnected to Y203 – and therefore ‘spectroscopically’ orange – when starting from OCP^O^ (Fig. 4 and Supplementary figs. 25-30), whereas it is highly twisted and detached from both Y203 and W290 – i.e. ‘spectroscopically red’, when starting from the OCP^1hν^ intermediate form (Fig. 6 and Supplementary fig. S28-S30 and S38). The two photoreactions differ in the time required for rupture of the two H-bonds tethering the carotenoid to the protein, and in the lifetime of that structure (Fig. 5A). They also differ in the time it takes for the computationally-predicted 7-*cis* CAN isomer^28^ to form (10 ps in the P2_1_ conformers vs. 1 µs in the C2 conformer) and its lifetime (< 100 ps in the P2_1_ conformers vs. up to the steady-state in the C2c conformer) (Figs. 4 and 6, and Supplementary figs. 25-30). We speculate that the five-order-of-magnitude increase in the lifespan of the untethered carotenoid in the OCP^1hν^ è OCP^R^ transition provides the time required for efficient translocation across the NTD (Fig. 7). We propose that the tightly constrained carotenoid geometry in the closed OCP^1hν^ intermediate reduces its conformational entropy, restricting fast relaxation pathways and keeping the carotenoid β1 ionone ring untethered until slower structural rearrangements allow translocation to proceed.

In conclusion, our study provides a structural framework of the OCP reaction mechanism. Sequence analysis and mutagenesis data indicate that our proposed mechanism—centered on αGH-loop transitions—is conserved across all OCP clades^20,30,31^ (Supplementary Table 10), independent of whether the dark state can dimerize (clade 1) or not (clades 2 and X). We show that in clade 1 OCP, formation of the OCP^1hν^ intermediate requires rupture of the key P13–A133 intermolecular tether otherwise blocking the αGH loop, inducing transient weakening of the dimer interface (Fig. 1F and Supplementary Fig. S33)^11^. This requirement is absent in constitutively monomeric OCPs from clades 2 and X, where αGH loop conformational dynamics are solely controlled by H-bonds equivalent to D35—Y129, D35—S132 and Q79—A123. Indeed, the C2c conformational substate characteristic for OCP^1hν^ is observed in the structure of the constitutively monomeric *Gloeobacter* OCP-X (wwPDB code <u>8A0H</u>)^20^. Our TCSF data also show that a significant fraction of OCP^1hν^ is present in CAN-OCP but not in ECN-OCP. This explains the apparent one-photon activation mechanism observed in CAN-OCP^13^.We posit that the apparent one vs. two photon photoactivation response in CAN and ECN, respectively, stems from differential structural dynamics imposed on the αGH loop by the chemical structure of the β2 ionone ring. This is likely of biological relevance since cyanobacterial species express a variety of carotenoids - but for most species the natural OCP pigment is not known. It is thus conceivable that beyond protein sequence and the associated regulation by oligomerization or conformational flexibility in the αCD and αGH loops, different carotenoids modulate the “dynamic range” of OCP-mediated photoprotection through the mechanism described here, thereby extending the range of ecosystems available for cyanobacteria.

## Supporting information

All supplementary tables

All supplementary figures

## Supplementary information includes

- **Supplementary figs. S1-S42**
- **Supplementary tables 1 to 9**
- **Supplementary texts 1 to 8**
- **Materials and Methods.**

## Acknowledgments

We dedicate this paper to the memory of Prof. Israel Silman. This work was supported by the Agence Nationale de la Recherche (grants ANR-17-CE11-0018-01, ANR-20-CE11-0019-02, ANR-22-CE11-0016 to J.-P.C., and ANR-2018CE11-0005-02 to MS, DK, IS and JPC) and the Max Planck Society. RJM is a recipient of a PhD fellowship from GRAL; QG is recipient of a PhD fellowship from the CFR program of the CEA; HEZ is recipient of a PhD fellowship from the IRGA program of the UGA. YK was supported by the ANR under the France 2030 grant reference number « ANR-24-RRII-0002 » operated by the INRIA Quadrant Program. This work was granted access to the HPC resources of IDRIS under the allocation 2025-A0180714660 (granted to Y.K.) made by GENCI. This work used the biophysics characterization, *in crystallo* optical spectroscopy (icOS) and electron microscopy (EM) platforms of the Grenoble Instruct-ERIC Center (IBS and ISBG; UAR 3518 CNRS-CEA-UGA-EMBL) within the Grenoble Partnership for Structural Biology (PSB), with support from FRISBI (ANR-10-INBS-05-02) and GRAL, a project of the University Grenoble Alpes graduate school (Ecoles Universitaires de Recherche) CBH-EUR-GS (ANR-17-EURE-0003). The EM facility is supported by the Auvergne-Rhône-Alpes Region, the Fondation pour la Recherche Médicale (FRM), the Fonds FEDER and the GIS-Infrastructures en Biologie Santé et Agronomie (IBiSA). The authors further acknowledge the provision of in-house experimental time from the CM01 facility at the European Synchrotron Radiation Facility (ESRF) in Grenoble. The cryo-EM data were acquired at the CM02 CRG beamline operated by IBS at the ESRF. The purchase of this microscope was funded by the EquipEx+ France Cryo-EM project (ANR-21-ESRE-0046). The authors thank Guy Schoehn for the establishment of CM02. We are grateful to Sylvain Engilberge, Eric Rives Mathieu and Antoine Royant for dedicated support of our BM7 experiments under proposal numbers A07-1-27 and A07-1-37, and of our iCOS experiments, under proposal numbers IH-MX-354, IH-MX-356, IH-MX-429, IH-MX-431 and IH-MX-462. We acknowledge in this context the French Biology/Health Panel Review Committee for provision of synchrotron radiation beamtime at the ESRF (Grenoble, France) on beamline BM07-FIP2, supported by the French ANR PIA3 (France 2030) EquipEx+ project MAGNIFIX under grant agreement ANR-21-ESRE-0011. We are grateful to Aline Le Roy, Philippe Mas and Caroline Mas for their support at the biophysical platform of the IBS (AUC, FIDA), and thank Marion Albasini, Kritika Sahni Ray, Einar Halldorsson and the Fidabio company (Søborg, Danemark) for providing access to the FIDA NEO instrument. We thank the staff of the ESRF and EMBL Grenoble, and especially Shibom Basu and Daniele de Sanctis, for assistance and support in using beamline(s) ID23-EH2, ID30A-1, ID30A-3 and CM01 under proposal numbers MX2440, MX2498, MX 2586, MX2628, MX2686 and MX2785. We thank Miriam Stricker, Kyprianos Hadjidemetriou, Guillaume Tetreau and the staff of the Coherent X-ray Imaging instrument of the Linear Coherent Light Source for support during preliminary tests of our crystals and Thomas Barends for discussions. We thank Tatiana Domratcheva for stimulating discussions and valuable feedback on our mechanistic hypotheses. We thank Ronald Rios Santacruz for help in testing the injection of crystals embedded in hydroxy-ethyl-cellulose at SACLA. We also thank the SwissFEL staff for supporting beamtimes at the Alvra Prime (experiments p17982 and p18386) and Cristallina MX (commissioning experiments p21231, p21529 and p21592) instruments.

## Authors contribution

JPC designed the project;

JPC, IS, DK and MS supervised parts of the project;

RM and JPC analyzed OCP sequences and designed mutants;

RM, EAA, EH, QG, HEZ and AW expressed and purified WT OCP;

RM, EAA, EH, NZ, QG, HEZ and AW generated, expressed and purified OCP mutants;

RM, QG, HEZ and AW performed photoactivity assays;

RM, QG and HEZ performed TCSF assays;

RM, EAA, EH, QG and HEZ produced crystals for synchrotron experiments;

RM, EAA, QG, HEZ, IS and JPC acquired synchrotron diffraction data;

RM and JPC processed MX data and refined corresponding structures;

SN, MS and GB performed and analyzed ultrafast spectroscopy on OCP crystals;

MB performed spectroscopy analysis;

FD and EB performed molecular dynamics simulations;

YK performed the ComPASS analysis of the time-resolved crystallographic data;

CCh performed metadynamics simulations;

EH and IS established microcrystallization;

RM and BB endeavoured the characterization of OCP photointermediates by NMR RM, EH and IS produced microcrystals for SFX and SSX experiments;

RM, EAA, EDZ, MW, GS, JPC, MK, MLG, SN, AG, LF, MH, MS, GNK, RBD, RLS, IS, DO, KN,

PJMJ, EB, CC, GK, CB, CJM, MVA, GG and FD prepared and performed SFX; MK, MLG, GNK, RBD and RLS performed sample injection;

MVA, JB and PJMJ carried out laser work at the SwissFEL; EAA, GS and JPC prepared SSX data collection;

GNK, GS and JPC performed SSX data collection;

JPC, KN, DO and EDZ performed online SFX data analysis; JPC, DO and NC performed offline SFX data processing; JPC performed online and offline SSX data processing;

JPC refined SFX and SSX structures and performed the related structural analysis; JPC and IS wrote the manuscript with input of RM, MW, GS, CCh and DK;

All authors discussed the results and manuscript.

## Supplementary tables

**Supplementary table 1:** Data processing and refinement statistics for the dark and light room-temperature crystallographic data collected at the ESRF-BM7 beamline on wild-type OCP crystals.

**Supplementary table 2:** Cryo-EM data collection, refinement and validation statistics

**Supplementary table 3:** Data processing and refinement statistics for the dark room-temperature crystallographic data investigating the effect of partial dehydration on wild-type OCP crystals at the ESRF-BM7 beamline.

**Supplementary table 4:** Defining features of the C2o and C2c states.

**Supplementary table 5:** Data processing and refinement statistics for the dark room-temperature crystallographic data collected on crystals of OCP mutants at the ESRF-BM7 beamline.

**Supplementary table 6:** Photoactivity and temperature-controlled scanning fluorimetry profile of CAN and ECN functionalized *Planktothrix aghardii* OCP and mutants thereof.

**Supplementary table 7:** Data processing and refinement statistics for the dark and light serial femtosecond crystallographic data collected at SwissFEL on P2_1_ microcrystals.

**Supplementary table 8:** Data processing and refinement statistics for the dark and light serial femtosecond crystallographic data collected at SwissFEL on C2 microcrystals.

**Supplementary table 9:** Data processing and refinement statistics for the dark and light serial synchrotron crystallographic data collected at ESRF-ID23EH2 on P21 and C2 microcrystals.

**Supplementary table 10:** Residues involved in the OCP^O^ è OCP^1hν^ transition are conserved.

## Supplementary figures

**Supplementary Fig. 1: Secondary structure plots for the various dark-states determined in this study, *i.e.* the cryo-EM structure of the A23C mutant and the wild-type P21 ^A^, P21 ^B^, C2o and C2c crystal structures.**

**Supplementary Fig. 2: The αCD loop adopts distinct conformations in the P2_1_ ^B^, P2_1_ ^A^ and C2 chains, resulting in different interactions with αC, αG, αM, the β5-β6 turn and the linker.** (**A**) Overall conformation and close-up views (CV) of the multiple interactions supported by the αCD loop in the P2_1_ ^B^ chain. **CV1** shows the residues of which the interatomic distances determine opening of SC#2, i.e. W41, Y44, Y111 and E115; no direct H-bonding interaction is observed between Y44 and E115, however a water-bridge exists. **CV2** shows the residues whose interaction stabilizes folding into a α helix of the entire αC, i.e. E46 and K49. **CV3** shows the region where the change in carotenoid steering conditions occurs upon conformational transition in the αCD loop; in the P2_1_ ^B^ chain, V53 plugs the carotenoid tunnel, with no direct interaction with the pigment. **CV4** shows the interaction between the αCD loop and the β5-β6 turn in the P2_1_ ^B^ chain. **CV5** shows that no interaction occurs between the αCD loop and the linker in the P2_1_ ^B^ chain. (**B**) Overall conformation and close-up views (CV) of the multiple interactions supported by the αCD loop in the P2_1_ ^A^ chain. **CV1** shows the residues whose interatomic distances determine opening of SC#2, i.e. W41, Y44, Y111 and E115; no direct H-bonding interaction is seen between Y44 and E115, however a water-bridge exists. **CV2** shows the broken interaction of E46 and K49 in the P2_1_ ^A^ chain and the associated break in the αC helix with residues M47 to T50 folding into a 3^10^ helix stabilized by H-bonding of T50(OG1) to M47(O). **CV3** shows the region where the change in carotenoid steering conditions occurs upon conformational transition in the αCD loop; in the P2_1_ ^A^ chain, I51 plugs the carotenoid tunnel, with direct contribution to the steering of the pigment. **CV4** shows the interface between the αCD loop and the β5β6 turn, upon folding into a 3^10^ helix of the last αC residues, in the P2 ^A^ and C2 chains. The interface features an extra H-bond (T52 contributes two H-bonds instead of a single one contributed by A54 in P2 ^B^), indicating increased stability of the αCD loop as well as stronger connections between the two domains. **CV5** shows that no interaction occurs between the αCD loop and the linker in the P2 ^A^ chain. (**C**) Overall conformation and close-up views (CV) of the multiple interactions supported by the αCD loop in the P2 ^A^ chain. **CV1** shows the residues whose interatomic distances determine opening of SC#2, i.e. W41, Y44, Y111 and E115; H-bonding occurs between Y44 and one of two E115 conformers, with the other H-bonding to Q119 at the N-terminus αGH loop. The distance between the rings of Y44 and W41 is such that SC#2 is closed. H-bonding is seen between Y44 and one of two E115 conformers, double locking SC#2. The other E115 conformer H-bonds to Q119 at the N-terminus αGH loop. **CV2** shows the broken interaction between E46 and K49 in the C2 chain and the associated break in the αC helix with residues M47 to T50 folding into a 3^10^ helix, stabilized by H-bonding of T50(OG1) to M47(O). **CV3** shows the region where the change in carotenoid steering conditions occurs upon conformational transition in the αCD loop; in the C2 chain, I51 plugs the carotenoid tunnel, with direct contribution to the steering of the pigment. Compared to the P2 ^A^ chain, it is a different I51 rotamer that is observed in the C2 chain, which pushes on Y44 thereby forcing the closing of SC#2. CV4 shows the revisited interface between the αCD loop and the β5-β6 turn, upon folding into a 3^10^ helix of the last αC residues, in the P2_1_ ^A^ and C2 chains. The revisited interface features an extra H-bond (T52 contributes two H-bonds instead of a single contributed by A54 in P2_1_ ^B^), indicating increased stability of the αCD loop as well as stronger connections between the two domains. CV5 shows that in the C2 chain, a direct interaction occurs between the αCD loop and the linker, mediated by main chain interactions between A58 and V154.

**Supplementary Fig. 3: The distinct conformations displayed by the αCD loop in the P21 ^B^, P21 ^A^ and C2 chains control access to the carotenoid tunnel via solvent channel #2 (SC#2).** (**A**) Recapitulation of the features of the three chains, and Connolly surface representation showing access of the solvent to the carotenoid tunnel via SC#2, in the P2_1_^B^ and P2_1_^A^ chains, but not in the C2 chain. (**B**) Difference distance matrices between the room-temperature (RT) dark-adapted P2_1_^B^, P2_1_^A^ and C2 structures. Major changes occur at the αCD and αGH loops.

Supplementary Fig. 4: The cryo-electron microscopy (cryoEM) structure of *Planktothrix aghardhii* OCP identifies the P21^B^ conformer as most representative of OCP^O^ in solution. (**A-C**) Wild-type (WT) OCP is unsuited for cryo-EM endeavors because monomers and dimers are in equilibrium, with only the latter being large enough to be amenable to structure determination by the method. We designed a cryo-EM compatible OCP mutant by replacing A23, at the center of the dimerization interface (**A**), by a cysteine (**B**). The A23C mutant featured a covalent disulfide, in X-ray electron density maps, and was also characterized as a constitutive dimer, by analytical centrifugation (**C**). (**D**) Electrostatic potential map for the A23C mutant, at 2.8 Å resolution. The modeled OCP structure is shown as a cartoon, colored by sequence from cold (blue) to hot (red). The αCD and αGH loops are highlighted, respectively colored in cyan and green. (**E**) Overlay of the αCD loop in the cryo-EM map and structure (cyan) with the conformation seen in the P2_1_^B^ (orange), P2_1_ ^A^ (blue), C2o (magenta), and C2c (pink) crystal structures. (**F**) Overlay of the αGH loop in the cryo-EM map and structure (green) with the conformation seen in the P2_1_ ^B^ (orange), P2_1_ ^A^ (blue), C2o (magenta), and C2c (pink) crystal structures. (**G-I**) The A23C mutant is photoactive and functional, though less than the WT protein. (**G**) Photoactivation kinetics. (**H**) Thermal recovery kinetics. (**I**) The A23C mutant is able to quench the core base (CB) of the PBS demonstrating that the accumulated OCP^R^ state is functional.

Supplementary Fig. 5: Supporting data for cryo-EM structure determination of the A23C and A23C-A133P mutant of *Planktothrix aghardhii* OCP. (**A**) A cryo-EM micrograph of the A23C OCP mutant dispensed on carbon grids at 5 mg/mL with a reference scale bar of 50 nm; (**B**) 2D classification experimental images of the A23C OCP mutant; (**C**) Surface views of the processed structure at two different orientations, with colors reflecting the local resolution (4.5 – 2.5 Å); (**D**) a graph plotting the gold standard and map vs model Fourier shell correlation as a function of resolution. Resolution limits as estimated to be 2.8 and 3.0 Å by the two methods (with FSC cutoffs of 0.143 and 0.5), respectively. (D) a cryo-EM micrograph of the A23C-A133P OCP mutant dispensed on carbon grids at 0.3 mg/mL with a reference scale bar of 50 nm; (**F**) 2D classification experimental images of the A23C-A133P OCP; (**G**) surface views of the processed structure at two different orientations, with colors reflecting the local resolution (4.5 – 3.1 Å); (**H**) a graph plotting the gold standard and map vs model Fourier shell correlation as a function of resolution. Resolution limits are estimated to be 3.1 and 3.6 Å by the two methods (with FSC cutoffs of 0.143 and 0.5), respectively.

**Supplementary Fig. 6: OCP crystals are plastic and both the P2_1_ to C2 and C2o to C2c transitions are achievable by controlled dehydration at room-temperature. (A) The space group transition of** from P2_1_ to C2 is favored by decreasing the relative humidity surrounding the crystals from 97 to 90%, notably at pH 7.5. (**B**) Structural similarities between the P21 ^A^ and C2 chains assign P21 ^A^ as an obligate intermediate along the P2 ^B^ to C2 transition, which preludes to closing of SC#2 upon adoption of the C2o state. (**C**) Decrease of the relative humidity does not affect the P2_1_ structures, but in C2 crystals, it favors accumulation of the C2c conformer, where SC#1 closes upon folding of the αGH loop into a 3^10^ helix, in contrast to the C2o conformer where SC#1 is open, mirroring the P2_1_ chains. It can be deduced that structural changes in the αGH loop are packing-limited in the P2_1_ crystals, and only become possible after transition to the C2o conformer. (**D**) Connolly surface representation showing access of the solvent to the carotenoid tunnel via SC#1, in the P2 ^B^, P2 ^A^ and C2o conformers, but not in the C2c conformer.

**Supplementary Fig. 7: Synopsis of the main C2o and C2c features.** The two conformers differ in the fold of the αGH loop, connecting the αBCHI and αDEFG bundles of the NTD and bordering SC#1. In the C2o conformer, the loop is found as an extended coil mainly stabilized by the D35—Y129 and Q79—A123 H-bonds. In the C2c conformer, these two H-bonds break as the loop folds into a 3^10^ helix that plugs P126 into the carotenoid tunnel, fully blocking solvent access to it. Y129 adopts a new rotamer, devoid of H-bonding partner, double locking SC#1. This conformation is stabilized by three H-bonds supporting formation of the 3^10^ helix. Besides H-bonds, adoption of the C2c conformer affects steric interactions with the αC helix, with replacement of the I125 residue sitting atop W41 by P126, in the P2_1_, P2_1_ and C2o conformers. Thus, the local environment of W41 changes significantly along the C2o to C2c transition. This is also true for the carotenoid which is closer to P126 in the C2c conformer than I125 in the C2o conformer.

**Supplementary Fig. 8: Structural changes induced by illumination of macrocrystals**. In the left column, an overview is presented, recapitulating the H-bonding situation of the carotenoid, interactions with closest residues along the tunnel, whether or not the interdomain salt-bridge is formed and if SC#1 is open, using P126 as a marker. In the right column, the focus is on the αGH loop, seen from a 90° rotation perspective compared to the images in the left column. As of 1 min. illumination, the C2c state accumulates in C2 crystals, regardless of the relative humidity. Conformational changes are also seen in the carotenoid structure, but it remains H-bonded to the protein. After 10 min. illumination, no change in the H-bonding of the carotenoid are observed, which furthermore displays less pronounced conformational changes than after 1 min. illumination. Nevertheless, we observe again accumulation of the C2c conformer. The decrease of its occupancy showcases the transient nature of the light—induced C2o to C2c transition.

**Supplementary Fig. 9: Overview of conformational changes triggered by illumination of macrocrystals.** Distance difference matrix plots comparing the various wild-type OCP structures collected from macrocrystals either kept in the dark (dark^MX, C2^ ^#1^ and dark^MX, C2^ ^#2^) or illuminated for 1 or 10 minutes at 537 nm during (light^1min, MX, C2^ ^#1^) or before (light^10min, MX, C2^ ^#2^) X-ray data collection, respectively. Secondary structure elements are shown as cartoons, colored according to sequence and labelled. Red and blue signals indicate elongation and shortening of the distance between compared elements, respectively.

**Supplementary Fig. 10: Temperature controlled scanning fluorimetry (TCSF) allows tracking the OCP^1hv^ intermediate in CAN-functionalized protein.** TCSF series from 15-90°C are presented of CAN-WT, ENC-WT and the constitutively-monomeric CAN-R27L mutant at concentrations of 30 (left), 95 (middle), and 160 µM (right). At these concentrations, 20, 8 and 5 % of the proteins are expected to be present as monomers, given the measured dissociation constant of 8 µM for CAN-WT. The raw fluorescence count at 350 nm is plotted in red, and the associated first derivative used to extract melting temperatures in black. CAN-functionalized proteins feature a concentration-dependent peak, in addition to the (∼54°C) denaturation peak also observed for ECN-functionalized protein. The concentration-dependent peak relates to a change in environment of a Trp residue, occurring in the monomeric (42°C; CAN-WT at low concentrations and CAN-R27L) as well as the dimeric protein (47°C; CAN-WT at high concentrations). The increase in melting temperature indicates that the stability of the structure/structural element containing the Trp reporter is higher in the dimer than in the monomer.

**Supplementary Fig. 11: The concentration dependent peak-shift observed in CAN-OCP (including WT) originates from W41.** (A) TCSF series from 15-90°C are presented of the CAN-W41F mutant at concentrations of 30 (left), 95 (middle), and 160 µM (right). The raw fluorescence count at 350 nm is plotted in red, and the associated first derivative used to extract melting temperatures in black. The amplitude of the concentration-dependent peak is greatly reduced upon replacement of W41 by a phenylalanine, indicating that this residue is that main contributor to the concentration-dependent TCSF signal observed in addition the (∼54°C) denaturation peak. The CAN-W41F mutant photoactivates slower but recovers faster than CAN-WT, resulting in a reduced steady-state accumulation of the OCP^R^ state. This suggests that the Trp acts as a “sliding base” for the αGH loop is important for the photoactivation efficiency and the stability of OCP^R^.

**Supplementary Fig. 12: The closed C2c state is an obligatory intermediate along the photocycle.** The A38C-I125C mutant, covalently locked in the opened state by a disulfide bond tethering the αGH loop to αC, is unable to photoactivate and yield the OCP^R^ state, regardless of whether it is functionalized by CAN (B) or ECN (C). (D-G) The control single mutants (CAN-I125C (D) and CAN-A38C (E)) mutants are photoactive (F), with an increased recovery rate (G). CAN-A38C has a lower photoactivation rate and higher recovery rate than CAN-I125Cys. Resultantly, the steady-state amount of OCP^R^ drops to 50% of that accumulated by CAN-WT.

**Supplementary Fig. 13: The concentration dependent peak-shift observed in CAN-OCP (including WT) is eliminated in the CAN-A38C-I125C mutant, covalently locked in the opened state by a disulfide tethering the αGH loop to αC.** TCSF series from 15-90°C are presented of CAN-A38C-I125C, CAN-A38C and CAN-I125C mutants at concentrations of 30 (left), 95 (middle), and 160 µM (right). The raw fluorescence count at 350 nm is plotted in red, and the associated first derivative used to extract melting temperatures in black. (Top) The concentration-dependent peak is missing in CAN-A38C-I125C, suggesting that in the CAN-WT protein, the peak informs on the melting of the SC#1-closed conformation of the αGH loop, where P126 interacts with W41 (C2c conformer), resulting in the more stable, SC#2-open conformation characterized by the interaction of I125 with W41 (P2 ^A^, P2 ^B^, C2o conformers). The shift of the melting temperature upon dimerization thus suggests that the stability of the C2c conformer increases upon dimerization. (Middle) The concentration dependent peak is present in the A38C mutant, though greatly diminished in amplitude, even more so at low concentrations. The reduced steady-state amount of the persistent OCP^1hν^ intermediate in the A38C mutant could explain its reduced photoactivation efficiency. The acceleration of recovery could originate from unfavorable interactions of the C38 thiol with CAN in the A38C-OCP^R^ state. (Bottom) The peak assigned to the melting of the C2c conformer is present, but has lost the concentration dependency, suggesting that the stability of the C2c conformer is the same in monomers and dimers of this mutant. The lack of stabilization of the C2c conformer upon dimerization suggests that the αGH loop is disconnected from the dimerization interface, *i.e.* the P13/A133P H-bond is broken in the I125C mutant. This was indeed observed in the crystal structure of this mutant.

**Supplementary Fig. 14: Mutants designed to favor accumulation of the C2c conformer preserve photoactivity.** The defining features of the C2c conformer are ruptured intramolecular Q79—A123 and D35—Y129 H-bonds. Adoption of this state is favored by rupture of the intermolecular P13—A133 H-bond in the C2o state. (**A-D**) The Q79L mutant, unable lever αGH loop structural dynamics by H-bonding to A123, is observed in the C2c conformer regardless of the relative humidity and keto-carotenoid pigment (**A, B**). Q79L hardly photoactivates, when functionalized by ECN, but retains ∼half of the WT photoactivation efficiency in the complex with CAN (**C**). The CAN-Q79L protein nevertheless recovers dramatically faster, challenging accumulation of the CAN-Q79L-OCP^R^ state, possibly owing to the desired increased stability of the C2c conformer – but more likely because of its intrinsic instability (**D**). The same could apply to the ECN-Q79L-OCP^R^ state, which would be even less stable than the CAN-Q79L-OCP^R^ state. Indeed, the Q79—A123 H-bond is formed in the CAN-WT-OCP^R^ state. (**E-H**) The D35T mutant, unable to anchor the αGH loop to αC, was also observed to favor the C2c conformer, regardless of the relative humidity and keto-carotenoid pigment (**E, F**). The D35T protein retains ∼half of the WT photoactivation efficiency, which together with an only slightly increased thermal recovery rate, affords accumulation of OCP^R^ to similar levels as CAN-WT. The fact that the D35—Y129 H-bond is replaced by the E34—Y129 H-bond in OCP^R^ explains why the D35T mutation does not affect the stability of OCP^R^. (**H-K**) The CAN-A133P mutant shows a similar propensity as the WT to adopt the C2c structure, i.e., it is hardly present at 97% relative humidity, but accumulates upon dehydration to 90% relative humidity. The mutation is silent in terms of both photoactivation and recovery.

**Supplementary Fig. 15: The peak representing the melting of the C2c conformer increases in amplitude but loses concentration dependency the CAN-Q79L and CAN-D35T mutants.** TCSF series from 15-90°C are presented of the CAN-Q79L, ECN-Q79L, CAN-D35T and ECN-D35T mutants at concentrations of 30 (left), 95 (middle), and 160 µM (right). The raw fluorescence count at 350 nm is plotted in red, and the associated first derivative used to extract melting temperatures in black. (**A**) The CAN-Q79L protein shows no dependency on concentration of the peak associated with the melting of the C2c conformer, indicating that in this protein, the A133P H-bond is unable to form, consistent with the observation that only the C2c conformer crystallized, regardless of the bound co-factor and relative humidity. Notwithstanding, the TCSF profile of the ECN-G79L mutant is identical to that the ECN-WT, indicating that in solution, the protein is mainly present in the open state(s) (P2_1_, P2_1_ and C2o). (**B**) The CAN-D35T protein also shows a complete loss of concentration dependency of the peak associated with the melting of the C2c conformer. This observation is consistent with the CAN-D35T crystal structure which shows an αGH loop devoid of H-bonding interactions at its C-terminus, i.e., where the intermolecular P13—A133 H-bonds are also disrupted. The amplitude of the peak is higher, suggesting an increased fraction of CAN-D35T in the C2c state – consistent with the increase in melting temperature. The TCSF profile of the ECN-D35T mutant is similar to ECN-WT, indicating that in solution, the protein is mainly present in the open conformation(s) (P2 ^A^, P2 ^B^ and C2o). Note the reduced stability of this protein, with a denaturation temperature decreased by nearly 5°C compared to ECN-WT. Like ECN-Q79L, we could trap and observe, in crystals, ECN-D35T in the C2c conformer.

**Supplementary Fig. 16: Molecular dynamics (MD) simulations and well-tempered metadynamics allow estimating the energy barriers between the P21^B^, C2o and C2c conformers. (A)** Enhanced sampling simulation using a path collective variable reaction coordinate built from the distances between I51(CD1) and CAN(C40), V53(CG1) and W279(O), T52 (N) and W279(O), and the RMSDs between P2_1_ ^B^ and C2o; **(B)** molecular dynamics simulation histogram plots showing the distribution of distances between P126(CB)/CAN(C06) and D35(OD2)/Y129(OH) when starting from the C2o (left) and C2c (right) conformers, in WT (top) and Q79L (bottom) OCPs. **(C)** Enhanced sampling simulation using a path collective variable reaction coordinate built from the distances between I51(CD1) and CAN(C40), V53(CG1) and W279(O), T52 (N) and W279(O), and the RMSDs between Q79L-C2o and Q79L-C2c. Presumably, the free energy of the C2c conformer is similar in the CAN-WT and CAN-Q79L proteins, since neither has a H-bond tethering the αGH loop to the αE helix. The barrier is however likely flatter in CAN-WT, in MD simulations the Q79—A123 H-bond easily reforms, favoring transition to the C2o conformer.

**Supplementary Fig. 17: Molecular dynamics (MD) simulations started from the CAN-WT protein in the C2o and C2c conformers point to a higher free energy of the latter.** The defining features of the C2o (blue) and C2c (magenta) conformers (Supplementary table 4) are monitored in terms of interatomic distances in multiple ∼1 µs-long MD trajectories started from the CAN-WT protein in the C2o and C2c conformers. It can be seen that multiple attempts are made by the protein in the C2c conformer to reach the C2o conformer, with a complete transition achieved in one of ten trajectories started from the C2c conformer. Conversely, the C2o conformer is stable on the timescale of our simulations, with no spontaneous attempts to reach the C2c conformer at equilibrium.

**Supplementary Fig. 18: Molecular dynamics (MD) simulations started from the CAN-Q79L protein in the C2o and C2c conformers point to a higher barrier between the two conformers upon suppression of the Q79L—A123 H-bond.** The defining features of the C2o (blue) and C2c (magenta) conformers (Supplementary table 4) are monitored in terms of interatomic distances in multiple ∼1 µs-long MD trajectories started from the CAN-Q79L protein in the C2o and C2c conformers. It can be seen that both the C2o and C2c conformers are stable on the timescale of our simulations, with no spontaneous interstate transition at equilibrium. Thus, the Q79L—A123 H-bond is central to lowering the barrier between the C2c and C2o conformers.

**Supplementary Fig. 19: The P13—A133 intermolecular H-bond is at the origin of the concentration dependance of the peak reflecting the melting of the C2c conformer in the CAN-WT protein.** (**A, B**) The A133P mutation, suppressing the possibility that the αGH loop be anchored by intermolecular H-bonding to P13, has no impact on the photoactivation and recovery rate of the CAN-OCP, indicating that the P13-A133 H-bond is by definition broken, in the C2c conformer, and not rate-limiting to break, upon illumination. Nevertheless, introduction of the A133P mutation in the rigidified, constitutively-dimeric CAN-A23C mutant partly rescues its reduced photoactivation efficiency while fully restoring the thermal recovery properties of the CAN-WT protein. (**C**) The A133P mutation results in a weakened dimer interface, and thus in a higher fraction of monomers for the mutant in analytical ultracentrifugation analyses carried out at 5 µM CAN-WT and CAN-A133P. (**D**) TCSF series from 15-90°C are presented of the CAN-A133P, CAN-A23Cys and CAN-A23C-A133P mutants at concentrations of 30 (left), 95 (middle), and 160 µM (right). The raw fluorescence count at 350 nm is plotted in red, and the associated first derivative used to extract melting temperatures in black. (Top) The CAN-A133P protein shows complete loss of the concentration dependent peak assigned to the melting of the C2c conformer. (Middle) The C2c conformer is not present in the rigidified CAN-A23C protein, whose TCSF profile resembles more ECN-WT than CAN-WT. (Bottom) Introduction of the A133P mutation in the A23C scaffold restores the possibility for the C2c conformation to form at equilibrium.

**Supplementary Fig. 20: Microcrystals and experimental setups.** (**A**) CAN-WT OCP microcrystals used for the TR-SFX experiments at SwissFEL Alvra-prime (left panel) and Cristallina-MX (right panel) beamlines. (**B**) Visualization of the HVE injector used to stream crystals across the XFEL beam at the Cristallina beamline (left) and the HVE stream intercepted by the optical and X-ray lasers (right). (**C**) CAN-WT OCP microcrystals used for the SSX experiment at the ESRF-ID23EH2 beamline. (**D**) Experimental setup of the 48h microcrystal illumination with blue light. A SCHOTT lamp (KL1500) equipped with a double gooseneck light guide was used to illuminate crystals 48h prior to the SSX experiment. **(E)** absorbance spectra of dark OCP microcrystals (black) and those illuminated for 48h with blue light.

**Supplementary Fig. 21: Transient absorption spectroscopy using microcrystals.** (**A**) Peak ΔA signals obtained for 510 nm (ground state bleach), 670 nm (S_1_ and ICT excited states absorption) and 1100 nm (S_2_ excited state absorption) plotted as a function of the peak power density (the pump energies used are 0.2, 0.4, 0.8, 1.6, 3.2 µJ). The 510 nm signal was inverted to obtain a positive value. Green and blue dashed vertical lines indicate equivalents of 32 nJ pulses (used at the TR-SFX experiment at Alvra, low power density (LPD)) and 80 nJ pulses (high power density (HPD). (**B**) Transient absorption spectra covering the carotenoid cation radical band peaking at 950 nm, ≈1ns. ΔA signals at 860 nm and 1090 nm were chosen to estimate the baseline, and this (linear) baseline was then subtracted from the data. Note that the 780-820 nm region is obscured by artifacts due to the residual fundamental beam. (**C**) Amplitude of the 950 nm band plotted as a function of the peak power density (using pump energies of 0.2, 0.4, 0.8, 1.6, 3.2 µJ). The green and blue dashed vertical lines correspond to using 32 nJ (LPD, TR-SFX) and 80 nJ (HPD) pump laser pulses. (**D**) Normalized stationary absorption spectra of the solution and crystalline OCP samples recorded using Jasco V-550 spectrophotometer. Since crystalline samples are affected by a strong scattering, baseline was subtracted. (**E**) Exemplary sample preparation of OCP crystals in LCP in a 100 µm cuvette.

**Supplementary Fig. 22: Overview of transient absorption kinetics (A-D) and spectra (E-I) for all datasets.** Datasets obtained from colloidal microcrystals suspended in PEG (P2_1_) are normalized to-1 at 516 nm, and at 542 nm for data from the microcrystals embedded in viscous LCP (C2, C2c conformer).

**Supplementary Fig. 23: Transient absorption spectra of the solution phase OCP and crystalline OCP**. The experiment on OCP microcrystals in LCP was repeated with a similar setup (70 fs, 530 nm, 0.5 µJ circularly polarized pump, 180 µm FWHM in the sample position). The transient spectra are plotted for 0.3 ps (containing ITC, S_1_ and S* contributions), 2.5 ps (containing S_1_ and S* contributions), 8 ps, 12ps delays (containing mostly S* state). Transient absorption kinetics of the solution phase and crystalline samples compared at (**E**) ground state bleach band and (**F**) excited state absorption band. The kinetics were rescaled in order to overlap them. The data were firstly normalized to-1 close to the bleaching extremum (note that for crystalline samples it was obscured by the excitation scattering) then kinetics were scaled by (**E**) 1.0, 0.8, 1.25 and (**F**) 1.0, 0.98, 0.97 factors. The data show excellent agreement between solution and crystalline phase kinetics.

**Supplementary Fig. 24: Distance difference matrix plots comparing the various dark-state wild-type OCP structures reported and discussed in the manuscript.** Comparison of the dark-state MX structures determined at ESRF-BM7, SFX structures determined at SwissFEL Alvra-prime and Cristallina MX, and SSX structure determined at ESRF-ID23EH2.

**Supplementary Fig. 25: Photoinduced structural changes in the P2_1_^A^ conformer.** Synopsis of the TR-SFX and SSX data, recapitulating the H-bonding situation of the carotenoid, interactions with closest residues along the tunnel, whether or not the interdomain salt-bridge is formed and if SC#1 is open, using P126 as a marker. Torsions and relaxations in the carotenoid are highlighted by magenta and blue circular arrows, respectively, with changes pertaining to the previous time delays. Accordingly, at each time delay, the greyed background structure is that characterized at the previous time delay.

**Supplementary Fig. 26: Photoinduced structural changes in the P2 ^B^ conformer.** Synopsis of the TR-SFX and SSX data, recapitulating the H-bonding situation of the carotenoid, interactions with closest residues along the tunnel, whether or not the interdomain salt-bridge is formed and if SC#1 is open, using P126 as a marker. Torsions and relaxations in the carotenoid are highlighted by magenta and blue circular arrows, respectively, with changes pertaining to the previous time delays. Accordingly, at each time delay, the greyed background structure is that characterized at the previous time delay.

**Supplementary Fig. 27: Photoinduced structural changes in the P21 ^A^ and P21 ^B^ conformers.** Synopsis of the TR-SFX and SSX data, focused on conformational changes occurring in the carotenoid in the two chains, and how this is influenced by interactions with closest residues along the tunnel. Carbon atoms of the P2 ^A^ protein and carotenoid are colored in yellow and green, respectively, while those of the P2 ^B^ protein and carotenoid are colored in pink and blue, respectively.

**Supplementary Fig. 28: Changes in dihedral angles at the β-ionone rings of the CAN pigment in the TR-SFX and SSX data collected on the P2_1_ and C2 crystals.** Data are reported for the P2 ^A^, P2 ^B^ and C2c chains. Vertical separation in the plots point to data collected in different regimes and different occasions, viz. the TR-SFX data collected on the 0.3-100 ps timescale, the TR-SFX data collected on the 10 ns-1µs time scale; SSX collected after 48h illumination. The last point (connected by a dashed line) in the last window corresponds to the presumed next state, *i.e.*, the C2c conformer, in the case of the P2_1_ chains, and the OCP^R^ state, in the case of the C2c chain. Data from the 48h illumination steady-state structures are plotted at apparent time delays of 0.5 ms, for the P21 ^A-light-48h^, P21 ^B-light-48h^ and C2c^light-48h^ photointermediates; and 10 ms, for the C2new^light-48h^ photointermediate.

**Supplementary Fig. 29: Changes in dihedral angles at the polyene of the CAN pigment in the TR-SFX and SSX data collected on the P2_1_ and C2 crystals.** Data are reported for the P2 ^A^, P2 ^B^ and C2c chains. Vertical separation in the plots point to data collected in different regimes and different occasions, viz. the TR-SFX data collected on the 0.3-100 ps timescale, the TR-SFX data collected on the 10 ns-1µs time scale; SSX collected after 48h illumination. The last point (connected by a dashed line) in the last window corresponds to the presumed next state, *i.e.*, the C2c conformer, in the case of the P2_1_ chains, and the OCP^R^ state, in the case of the C2c chain. Data from the 48h illumination steady-state structures are plotted at apparent time delays of 0.5 ms, for the P2 ^A-light-48h^, P2 ^B-light-48h^ and C2c^light-48h^ photointermediates; and 10 ms, for the C2new^light-48h^ photointermediate.

**Supplementary Fig. 30: Carotenoid untethering is rapid and short-lived in the P2_1_ chains, but delayed and prolonged in the C2c chain.** The H-bonding status of the carotenoid is shown for the P21 ^A^, P21 ^B^ and C2c conformers at all time delays across the TR-SFX and SSX data series, alongside its electron density at 1 σ in conventional 2mFo-DFc (dark state structures; C2new^light-48h^) and extrapolated (all other) 2Fextr-DFc maps.

**Supplementary Fig. 31: Steric propagation of the photoexcitation signal in the dark OCP^O^ state**. Results are shown for the P2 ^B^ conformer, most representative of the dark OCP^O^ state. In the left panels, the relative positioning of the αBCHI and αDEFG bundles is shown. In the middle and right panels, the mechanism that allows signal transduction from the carotenoid to the β-sheet (Arg291, via Trp290) to the αA helix (Arg9, Pro13) in one monomer (here P2 ^B^, in pink) to the αGH loop of the facing monomer (Ala133), weakening the dimer interface. These are shown at select time delays across the TR-SFX (0.3 ps – 1µs) and SSX (48h illumination) series, that match those illustrated in the corresponding Figure 4.

**Supplementary Fig. 32: Focus on the mechanism that allows signal transduction from the carotenoid to the β-sheet to the αA helix in one monomer to the αGH loop of the facing monomer.** For the three conformers, we show across the TR-SFX and SSX data series, the changes in interactions occurring at the contact points between the carotenoid and the β-sheet (R291, via W290), the β-sheet and the αA helix (R9, Pro13), and the αGH loop (A133) of the facing monomer (A133) in the dimer.

**Supplementary Fig. 33: Focus on the changes undergone at the dimer interface.** For the three conformers, we show across the TR-SFX and SSX data series, the changes in interactions occurring at the dimer interface.

**Supplementary Fig. 34: Focus on the changes undergone at the interface between the αBCHI and αDEFG bundles.** For the three conformers, we show across the TR-SFX and SSX data series, the conformational changes undergone at the interface between the αBCHI (slate) and αDEFG (pink) bundle.

**Supplementary Fig. 35: Focus on the changes undergone at the αGH loop.** For the three conformers, we show across the TR-SFX and SSX data series, the conformational changes undergone at the αGH loop.

**Supplementary Fig. 36: Overview of conformational changes in the P2 ^A^ conformer across the TR-SFX and SSX data series.** Distance difference matrix plots comparing the photo-triggered (light) P2 ^A^ states captured at different time delays across the TR-SFX and SSX data series, to their corresponding dark-state (light-dark) as well as to one another (light-light)=. Secondary structure elements are shown as cartoons, colored according to sequence and labelled. Red and blue signal indicate increase and decrease of the distance between compared elements, respectively.

**Supplementary Fig. 37: Overview of conformational changes in the P2 ^B^ conformer across the TR-SFX and SSX data series.** Distance difference matrix plots comparing the photo-triggered (light) P2 ^B^ states captured at different time delays across our TR-SFX the SSX data series, to their corresponding dark-state (light-dark) as well as to one another (light-light), shown sequentially in time. Secondary structure elements are shown as cartoons, colored according to sequence and labelled. Red and blue signal indicate increase and decrease of the distance between compared elements, respectively.

**Supplementary Fig. 38: Photoinduced structural changes in the C2c conformer.** Overview of the TR-SFX and SSX data, recapitulating the H-bonding situation of the carotenoid, interactions with closest residues along the tunnel, whether or not the interdomain salt-bridge is formed and if SC#1 is open, using P126 as a marker. Torsions and relaxations in the carotenoid are highlighted by magenta and blue circular arrows, respectively, with changes pertaining to the previous time delays. Accordingly, at each time delay, the greyed background structure is that characterized at the previous time delay.

**Supplementary Fig. 39: Overview of conformational changes in the C2c conformer across the TR-SFX and SSX data series.** Distance difference matrix plots comparing the photo-triggered (light) C2c conformers captured at different time delays across the TR-SFX and SSX data series, to their corresponding dark-state (light-dark) as well as to one another (light-light), shown sequentially with time delays. Secondary structure elements are shown as cartoons, colored according to sequence and labelled. Red and blue signal indicate increase and decrease of the distance between compared elements, respectively.

**Supplementary Fig. 40: Steric propagation of the photoexcitation signal in the dark OCP^1hν^ intermediate.** In the left panels, the relative positioning of the αBCHI and αDEFG bundles is shown. In the middle and right panels, the mechanism that allows signal transduction from the carotenoid to the β-sheet (R291, via W290) to the αA helix (R9, P13) in one monomer (here P2 ^B^, in pink) to the αGH loop of the facing monomer (A133), weakening the dimer interface. These are shown at select time delays across the TR-SFX (0.3 ps – 1µs) and SSX (48h illumination) series, that match those illustrated in the corresponding Figure 6.

**Supplementary Fig. 41: Focus on the changes undergone at the αCD loop, at the αΗ/αI turn and at the αH/αN interface.** For the three conformers, we show the changes in interactions occurring at the αΗ/αI turn and at the αH/αN interface, across the TR-SFX and SSX data series.

**Supplementary Fig. 42:** (**A**) **Characterization of parameters of the transient absorption experiments.** Raman signal extracted from the OCP signal by subtraction of 645 nm kinetic. (**B**) The resulting Raman kinetics was fitted with a Gaussian function, resulting in a 149 fs (FWHM) instrument response function (IRF). (**C**) The pump spatial profile was determined by a horizontal razor edge scan of the pump beam in the sample position, resulting in 246 µm (FWHM). (**D, E**) The energy density and power density profiles of the 532 nm pump pulse were calculated using the experimentally derived (see **B, C**) pump beam parameters The peak values are 1.17 mJ/cm^2^ (**D**) and 10.44 GW/cm^2^ (**E**) for 0.8 µJ pump laser energy.

