## Supplementary material for "Structural basis of the two-photon photoactivation mechanism of orange carotenoid protein": All supplementary tables

**Supplementary Table 1**: Room temperature C2 and $P2_1$ macrocrystal structures. Data processing and refinement statistics for OCP MX experiments.

| | **P21 dark** | **C2 dark 1** | **C2 dark 2** | **C2 1 min illumination** | **C2 10 min illumination** |
|---|---|---|---|---|---|
| **PDB ID** | **30MP** | **30MO** | **30LU** | **30LV** | **30LW** |
| **Data collection and processing statistics** | | | | | |
| Beamline | BM07 - FIP2 | BM07 - FIP2 | BM07 - FIP2 | BM07 - FIP2 | BM07 - FIP2 |
| Wavelength (Å) | 0.9795 | 0.9795 | 0.9795 | 0.9795 | 0.9795 |
| Space group | *P 1 2 1* | *C 2* | *C 2* | *C 2* | *C 2* |
| Unit cell parameters | | | | | |
| a, b, c (Å) | 63.7 69.8 77.7 | 82.7. 69.3. 62.7 | 82.8. 69.1. 62.8 | 83.8. 69.8. 64.0 | 83.0. 68.8. 62.9 |
| $\alpha, \beta, \gamma$ (°) | 90 101.67 90 | 90.0. 117.7. 90.0 | 90.0. 117.7. 90.0 | 90.0. 117.9 .90.0 | 90.0. 117.9. 90.0 |
| Diffraction limit (Å) | 46.54 - 2.14 (2.22 - 2.14) | 50.34 - 1.78 (1.81 - 1.78) | 55.58 - 1.64 (1.67 - 1.64) | 50.81 - 2.20 (2.27 - 2.20) | 55.64 - 2.00 (2.05 - 2.00) |
| $R_{meas}$ (%) | 16.5 (130.1) | 12.1 (117.7) | 13.8 (159.6) | 23.1 (161.4) | 8.0 (78.7) |
| $R_{pim}$ (%) | 8.7 (68.5) | 6.5 (62.9) | 7.4 (84.9) | 12.3 (86.9) | 6.0 (123.7) |
| $CC_{1/2}$ (%) | 99.5 (49.1) | 99.5 (39.5) | 99.5 (28.4) | 97.5 (33.1) | 99.8 (67.7) |
| $\langle I/\sigma(I) \rangle$ | 6.6 (1.3) | 6.4 (0.9) | 6.4 (0.9) | 4.9 (1.5) | 11.70 (2.09) |
| No. of reflections | 125518 (12002) | 96287 (5047) | 125237 (6066) | 53136 (4726) | 70927 (5198) |
| No. of unique reflections | 35981 (3448) | 28229 (1492) | 37189 (1802) | 15528 (1379) | 20784 (1529) |
| Multiplicity | 3.5 (3.5) | 3.4 (3.4) | 3.4 (3.4) | 3.4 (3.4) | 3.41 (3.39) |
| Completeness (%) | 97.5 (96.0) | 93.6 (86.8) | 96.9 (94.7) | 93.5 (94.2) | 97.5 (98.5) |
| $R_{iso}$ (%)[‡] | N.A. | N.A. | N.A. | 21.7 (47.0) | 11.19 (24.48) |
| **Refinement statistics** | | | | | |
| **Refinement strategy**[#] | Classic | Classic | Classic | Extrapolated | Extrapolated |
| Diffraction limit (Å) | 44.09 – 2.14 (2.2 – 2.14) | 36.60 – 1.77 (1.81 – 1.78) | 55.58 – 1.64 (1.66 – 1.64) | 32.47 – 2.04 (2.17 – 2.04) | 19.52 – 2.0 (2.11 – 2.0) |
| No. of reflections | 35912 (2675) | 28220 (1377) | 37185 (1479) | 16525 (2050) | 20028 (2837) |
| R-work (%) | 19.81 | 16.55 | 17.40 | 33.98 | 33.26 |
| R-free (%)[§] | 23.90 | 21.00 | 20.27 | 37.29 | 36.51 |

| | | | | | |
|---|---|---|---|---|---|
| Number of atoms | 5403 | 2874 | 3098 | 2647 | 2692 |
| macromolecules | 4967 | 2615 | 2732 | 2499 | 2478 |
| ligands | 84 | 42 | 42 | 42 | 84 |
| solvent | 352 | 217 | 324 | 106 | 130 |
| Protein residues | 620 | 309 | 309 | 309 | 309 |
| RMS | | | | | |
| bonds (Å) | 0.009 | 0.011 | 0.006 | 0.003 | 0.007 |
| angles (°) | 0.44 | 0.91 | 0.79 | 0.46 | 0.50 |
| Ramachandran (%) | | | | | |
| favored | 98.53 | 99.02 | 98.69 | 97.05 | 96.07 |
| allowed | 1.47 | 0.98 | 0.98 | 2.62 | 3.28 |
| outliers | 0.00 | 0.00 | 0.33 | 0.33 | 0.66 |
| Rotamer outliers (%) | 0.75 | 2.47 | 2.03 | 2.61 | 2.25 |
| Clashscore | 2.60 | 4.54 | 2.89 | 4.37 | 3.77 |
| Average B-factor (Å$^2$) | 41.15 | 35.17 | 30.70 | 23.33 | 38.45 |
| macromolecules | 40.94 | 34.34 | 28.93 | 23.27 | 38.74 |
| ligands | 28.07 | 22.30 | 18.85 | 26.33 | 26.65 |
| solvent | 47.29 | 47.73 | 47.14 | 23.59 | 40.47 |

*Values between parentheses refer to the highest resolution shell; [§]Rfree is calculated using 5 % of random reflections excluded from refinement; ‡ Riso is defined as $\sum|F_{obs}^{dark} - F_{obs}^{light}| \Big/ \sum|F_{obs}^{dark} + F_{obs}^{light}|/2$

[#]Refinement has been carried out in extrapolated structure factor amplitudes (“extrapolated”) or the original structure factor amplitudes/intensities (classic); N.A.: Not applicable

$P2_1$ dark is the structure also used for $P2_1$ at 97% RH coupled which is coupled with ‘$P2_1$ 90% RH’ in Table 3.

**Supplementary Table 2:** Cryo-EM data collection, refinement and validation statistics

| | #1 Mutant A23C<br>(EMDB-56580)<br>(PDB 28KP) | #2 Mutant A23C – A133P<br>(EMDB-56539)<br>(PDB 28JA) |
|---|---|---|
| **Data collection and processing** | | |
| Magnification | 215 000 | 215 000 |
| Voltage (kV) | 300kV | 300kV |
| Electron exposure (e–/Å$^2$) | 60 | 60 |
| Defocus range (μm) | -1 to -2.2 | -1 to -2.2 |
| Pixel size (Å) | 0.57 | 0.57 |
| Symmetry imposed | C2 | C2 |
| Final particle images (no.) | 230 709 | 100 860 |
| Map resolution (Å) | 2.8 | 3.1 |
| FSC threshold | 0.143 | 0.143 |
| **Refinement** | | |
| Model resolution (Å) | 3.0 | 3.6 |
| FSC threshold | 0.5 | 0.5 |
| Model composition | | |
| Non-hydrogen atoms | 5052 | 4900 |
| Protein residues | 636 | 618 |
| Ligands | 2 | 2 |
| R.m.s. deviations | | |

| | | |
|---|---|---|
| Bond lengths (Å) | 1.3 | 1.3 |
| Bond angles (°) | 5.3 | 5.4 |
| Validation | | |
| MolProbity score | 2.2 | 1.9 |
| Clashscore | 9.9 | 9.2 |
| Poor rotamers (%) | 3.0 | 2.3 |
| Ramachandran plot | | |
| Favored (%) | 95.6 | 97.1 |
| Allowed (%) | 3.8 | 2.6 |
| Disallowed (%) | 0.3 | 0.3 |

Supplementary Table 3: Room temperature structures from macrocrystals at 90 and 97% relative humidity. Data processing and refinement statistics for OCP MX experiments.

| | **C2 $WT_{CAN}$ 97% RH** | **C2 $WT_{CAN}$ 90% RH** | **P21 $WT_{CAN}$ 90% RH** | **C2 $WT_{ECN}$ 97% RH** | **C2 $WT_{ECN}$ 90% RH** |
|---|---|---|---|---|---|
| **PDB ID** | **30LR** | **30LS** | **30LQ** | **30LO** | **30LP** |
| **Data collection and processing statistics** | | | | | |
| Beamline | BM07 - FIP2 | BM07 - FIP2 | BM07 - FIP2 | BM07 - FIP2 | BM07 - FIP2 |
| Wavelength (Å) | 0.9795 | 0.9795 | 0.9795 | 0.9795 | 0.9795 |
| Space group | *C 2* | *C 2* | *P 1 2 1* | *C 2* | *C 2* |
| Unit cell parameters | | | | | |
| a, b, c (Å) | 82.8 69.4 62.7 | 82.5 68.7 68.7 | 63.3 69.2 77.0 | 83.5 69.4 63.1 | 83.3 68.7 63.2 |
| $\alpha, \beta, \gamma$ (°) | 90 117.76 90 | 90 117.68 90 | 90 101.05 90 | 90 117.96 90 | 90 118.04 90 |
| Diffraction limit (Å) | 45.21 - 1.78 (1.84 - 1.78) | 55.60 - 1.89 (1.98 - 1.89) | 46.17 - 2.70 (2.83 - 2.70) | 45.38 – 2.31 (2.39 – 2.31) | 50.21 – 2.81 (2.96 – 2.81) |
| $R_{meas}$ (%) | 15.8 (115.1) | 15.4 (124.2) | 32.2 (208.9) | 20.7 (124.2) | 22.4 (117.4) |
| $R_{pim}$ (%) | 8.3 (61.0) | 8.2 (66.0) | 17.4 (110.4) | 11.0 (65.4) | 11.9 (61.3) |
| $CC_{1/2}$ (%) | 99.0 (40.6) | 99.4 (41.5) | 98.1 (44.5) | 98.7 (43.4) | 98.2 (49.3) |
| $\langle I/\sigma(I) \rangle$ | 4.0 (0.7) | 7.7 (1.3) | 4.6 (0.9) | 5.4 (1.5) | 5.6 (1.4) |
| No. of reflections | 101497 (9855) | 83954 (5084) | 58361 (8203) | 47216 (4754) | |
| No. of unique reflections | 29291 (2860) | 24271 (1470) | 17575 (2347) | 13794 (1353) | 7605 (1082) |
| Multiplicity | 3.5 (3.4) | 3.5 (3.5) | 3.3 (3.5) | 3.4 (3.5) | 3.4 (3.6) |
| Completeness (%) | 97.3 (96.9) | 97.6 (92.9) | 97.4 (98.4) | 98.3 (99.0) | 98.1 (96.7) |
| $R_{iso}$ (%)$^{‡}$ | N.A. | 17.66 | N.A. | N.A. | 16.16 (26.28) |
| **Refinement statistics** | | | | | |
| **Refinement strategy**$^{\#}$ | Classic | Extrapolated (occ 27.9%) | Classic | Classic | Extrapolated (occ 30%) |
| Diffraction limit (Å) | 40.44 – 1.78 (1.84 – 1.78) | 29.43 – 1.89 (1.98 – 1.89) | 43.62 – 2.70 (2.87 – 2.70) | 28.58 – 2.31 (2.49 – 2.31) | 40.78 – 2.81 (3.22 – 2.81) |
| No. of reflections | 29176 (2803) | 23498 (2622) | 17486 (2925) | 13760 (2739) | 7467 (2457) |
| R-work (%) | 16.43 | 26.85 | 21.86 | 18.53 | 29.74 |

| | | | | | |
|---|---|---|---|---|---|
| R-free (%)[§] | 20.77 | 31.40 | 26.88 | 23.59 | 35.11 |
| Number of atoms | 2873 | 2730 | 5105 | 2708 | 2453 |
| macromolecules | 2612 | 2520 | 4907 | 2441 | 2364 |
| ligands | 42 | 42 | 84 | 41 | 41 |
| solvent | 219 | 168 | 114 | 226 | 48 |
| Protein residues | 310 | 310 | 619 | 309 | 307 |
| RMS | | | | | |
| bonds (Å) | 0.012 | 0.010 | 0.012 | 0.002 | 0.002 |
| angles (°) | 0.88 | 0.64 | 0.46 | 0.40 | 0.58 |
| Ramachandran (%) | | | | | |
| favored | 99.02 | 99.35 | 96.73 | 98.69 | 97.03 |
| allowed | 0.98 | 0.65 | 3.11 | 1.31 | 2.64 |
| outliers | 0.00 | 0.00 | 0.16 | 0.00 | 0.33 |
| Rotamer outliers (%) | 1.42 | 1.48 | 0.95 | 0.38 | 0.40 |
| Clashscore | 3.79 | 2.75 | 4.76 | 3.22 | 7.11 |
| Average B-factor ($Å^2$) | 35.77 | 44.06 | 52.19 | 42.32 | 44.96 |
| macromolecules | 34.98 | 43.93 | 52.48 | 41.63 | 45.45 |
| ligands | 22.20 | 35.05 | 40.98 | 30.95 | 29.37 |
| solvent | 47.88 | 48.25 | 47.86 | 51.74 | 33.85 |

*Values between parentheses refer to the highest resolution shell; [§]Rfree is calculated using 5 % of random reflections excluded from refinement; ‡ Riso is defined as $\sum |F_{obs}^{dark} - F_{obs}^{light}| \Big/ \sum |F_{obs}^{dark} + F_{obs}^{light}|/2$

[#]Refinement has been carried out in extrapolated structure factor amplitudes ("extrapolated") or the original structure factor amplitudes/intensities (classic); N.A.: Not applicable

**Supplementary Table 4**: Defining features of the C2o and C2c states.

| Type of interaction | Involved residues | Distance in C2o (Å) | Distance in C2c (Å) |
|---|---|---|---|
| H-bonds between the αGH loop and neighboring structural elements (αE, αC) | Q79(OD1)—A123(N) | 2.7 | **4.0** |
| | D35(OD1)—Y129(OH) | 3.2 | **9.7** |
| | D35(OD2)—S132(OG1) | 2.6 | 3.0 |
| | D35(OD1)—S132(N) | 3.1 | 3.0 |
| Van der Waals interactions between the αGH loop and neighboring structural elements (αE, αC) | W41(CD1)—I125(CB) | 3.9 | **8.6** |
| | W41(CB)—I125(CG1) | 3.6 | **9.5** |
| | W41(CD1)—P126(CB) | **8.6** | 4.5 |
| | W41(CB)—P126(CB) | **9.7** | 3.6 |
| | T80(CG2)—Y129(OH) | **9.6** | 3.3 |
| H-bonds stabilizing the $3_{10}$ helix | I125(O)—G128(N) | **6.3** | 3.3 |
| | P124(O)—S127(N) | **6.3** | 3.2 |
| | P124(O)—S127(OG1) | **8.4** | 2.6 |
| Closest van der Waals interactions between the αGH loop and canthaxanthin. | I125(CG1)—CAN(C3′) | 5.2 | **8.4** |
| | P126(CG)—CAN(C3′) | **11.0** | 4.2 |

**Supplementary Table 5**: Room temperature structures of OCP mutants in complex with CAN or ECN.

| | **A23C$_{CAN}$ at 100K** | **A38C-I125C$_{CAN}$ 97% RH** | **A38C$_{CAN}$ at 100K** | **I125C$_{CAN}$ 97% RH** | **A38C-I125C-T80W$_{CAN}$ 97% RH** | **Q79L$_{CAN}$ 97% RH** | **Q79L$_{ECN}$ 97% RH** |
|---|---|---|---|---|---|---|---|
| **PDB ID** | **30LX** | **30LZ** | **30LY** | **30MA** | **30MD** | **30MB** | **30MC** |
| **Data collection and processing statistics** | | | | | | | |
| Beamline | ID30A-1 | BM07 - FIP2 | BM07 - FIP2 | BM07 - FIP2 | BM07 – FIP2 | BM07 - FIP2 | BM07 - FIP2 |
| Wavelength (Å) | 0.9654 | 0.9795 | 0.9795 | 0.9795 | 0.9795 | 0.98 | 0.9795 |
| Space group | *P 1 2 1* | C 1 2 1 | *P 1 2 1* | C 1 2 1 | C 1 2 1 | C 1 2 1 | C 1 2 1 |
| Unit cell parameters | | | | | | | |
| a, b, c (Å) | 62.49 69.02 76.91 | 83.8 68.0 62.8 | 62.6 63.7 74.5 | 82.7 69.2 62.9 | 83.45 68.22 62.36 | 83.1 70.6 62.8 | 82.2 68.8 62.7 |
| $\alpha, \beta, \gamma$ (°) | 90 100.67 90 | 90 118.65 90 | 90 104.51 90 | 90 117.58 90 | 90 118.46 90 | 90 117.43 90 | 90 117.47 90 |
| Diffraction limit (Å) | 43.85 – 1.62 (1.68 – 1.62) | 55.12 - 2.50 (2.60 - 2.50) | 72.16 – 1.99 (1.95 – 1.92) | 55.79 - 1.87 (1.90 - 1.87) | 44.88 – 1.89 (1.94 – 1.89) | 45.56 - 1.99 (2.06 - 1.99) | 50.09 - 2.00 (2.05 - 2.00) |
| $R_{meas}$ (%) | 4.9 (122.6) | 15.9 (110.8) | 20.7 (135.4) | 19.8 (177.5) | 6.5 (148.2) | 14.5 (177.6) | 17.4 (130.7) |
| $R_{pim}$ (%) | 3.0 (73.6) | 8.5 (58.3) | 10.9 (71.0) | 10.4 (93.0) | 3.5 (78.7) | 8.8 (107.5) | 9.3 (69.1) |
| $CC_{1/2}$ (%) | 99.8 (60.2) | 99.3 (50.6) | 99.1 (35.1) | 99.0 (33.2) | 99.9 (49.2) | 99.1 (34.1) | 99.3 (44.7) |
| $\langle I/\sigma(I) \rangle$ | 10.8 (0.9) | 7.3 (1.6) | 4.6 (1.1) | 5.5 (0.9) | 10.6 (1.0) | 5.3 (0.7) | 4.8 (0.9) |
| No. of reflections | 179686 (17734) | 35049 (4279) | 138501 (7010) | 87138 (4394) | 83598 (5402) | 49698 (4908) | 71005 (5268) |
| No. of unique reflections | 78714 (7741) | 10476 (1208) | 399301 (1991) | 25229 (1248) | 24360 (1562) | 20740 (2040) | 20742 (1522) |
| Multiplicity | 2.3 (2.3) | 3.3 (3.5) | 3.5 (3.5) | 3.5 (3.5) | 3.4 (3.5) | 2.4 (2.4) | 3.4 (3.5) |
| Completeness (%) | 96.5 (96.9) | 97.0 (99.3) | 97.3 (97.6) | 96.9 (97.1) | 99.3 (97.6) | 94.0 (95.3) | 98.5 (98.9) |
| $R_{iso}$ (%)$^{‡}$ | N.A. | N.A. | N.A. | N.A. | N.A. | N.A. | N.A. |
| **Refinement statistics** | | | | | | | |
| **Refinement strategy**[#] | Classic | Classic | Classic | Classic | Classic | Classic | Classic |
| Diffraction limit (Å) | 41.87 – 1.62 (1.64 -1.62) | 27.56 – 2.50 (2.63 – 2.50) | 72.16 – 1.95 (2.01 – 1.96) | 36.63 – 1.87 (1.95 – 1.87) | 44.88 – 1.89 (1.97 – 1.89) | 40.55 – 1.99 (2.1 – 1.99) | 40.17 – 2.0 (2.05 – 2.00) |
| No. of reflections | 78513 (2946) | 10463 (1533) | 39847 (2794) | 25166 (2724) | 24337 (2666) | 20697 (3006) | 20732 (1461) |
| R-work (%) | 19.94 | 20.45 | 21.97 | 16.28 | 20.53 | 17.10 | 16.86 |

| | | | | | | | |
|---|---|---|---|---|---|---|---|
| R-free (%)[§] | 24.67 | 24.72 | 25.97 | 20.29 | 24.83 | 22.22 | 21.69 |
| Number of atoms | 5496 | 2585 | 5347 | 2754 | 2602 | 2659 | 2758 |
| macromolecules | 4894 | 2423 | 4861 | 2476 | 2385 | 2444 | 2496 |
| ligands | 96 | 42 | 84 | 42 | 42 | 42 | 41 |
| solvent | 506 | 120 | 402 | 236 | 175 | 173 | 221 |
| Protein residues | 619 | 309 | 612 | 309 | 305 | 307 | 309 |
| RMS | | | | | | | |
| bonds (Å) | 0.011 | 0.005 | 0.010 | 0.013 | 0.013 | 0.012 | 0.002 |
| angles (°) | 0.99 | 0.44 | 0.48 | 0.92 | 0.93 | 0.89 | 0.48 |
| Ramachandran (%) | | | | | | | |
| favored | 98.85 | 98.69 | 99.00 | 99.34 | 98.67 | 98.68 | 99.34 |
| allowed | 1.15 | 1.31 | 0.83 | 0.66 | 1.33 | 1.32 | 0.33 |
| outliers | 0.00 | 0.00 | 0.17 | 0.00 | 0.00 | 0.00 | 0.33 |
| Rotamer outliers (%) | 1.33 | 2.30 | 0.57 | 0.75 | 1.96 | 0.00 | 2.23 |
| Clashscore | 4.25 | 6.15 | 4.18 | 3.82 | 3.34 | 2.03 | 3.93 |
| Average B-factor ($Å^2$) | 36.91 | 51.64 | 30.67 | 27.05 | 49.12 | 39.53 | 30.59 |
| macromolecules | 36.34 | 51.68 | 30.40 | 26.04 | 48.74 | 39.11 | 29.83 |
| ligands | 27.92 | 39.70 | 21.83 | 15.55 | 37.45 | 26.51 | 19.07 |
| solvent | 44.10 | 55.01 | 35.82 | 39.77 | 57.05 | 48.53 | 41.32 |

| | **D35T$_{CAN}$ 97% RH** | **D35T$_{CAN}$ 90% RH** | **A133P$_{CAN}$ 97% RH** | **A133P$_{CAN}$ 90% RH** |
|---|---|---|---|---|
| **PDB ID** | **30ME** | **30MF** | **30MG** | **30MH** |
| **Data collection and processing statistics** | | | | |
| Beamline | BM07 - FIP2 | BM07 - FIP2 | BM07 - FIP2 | BM07 - FIP2 |
| Wavelength (Å) | 0.9795 | 0.9795 | 0.9795 | 0.9795 |

| | | | | |
|---|---|---|---|---|
| Space group | C 1 2 1 | C 1 2 1 | C 1 2 1 | C 1 2 1 |
| Unit cell parameters | | | | |
| a, b, c (Å) | 82.4 69.0 62.8 | 82.4 68.8 62.8 | 82.7 68.8 62.6 | 82.7 67.8 62.7 |
| $\alpha, \beta, \gamma$ (°) | 90 117.61 90 | 90 117.69 90 | 90 117.59 90 | 90 117.76 90 |
| Diffraction limit (Å) | 50.15 - 2.20 (2.27 - 2.20) | 55.61 - 1.94 (1.99 - 1.94) | 55.51 - 2.00 (2.05 - 2.00) | 55.52 – 2.24 (2.32 – 2.24) |
| $R_{meas}$ (%) | 10.7 (63.4) | 14.1 (113.4) | 16.7 (100.2) | 30.5 (203.7) |
| $R_{pim}$ (%) | 6.0 (37.0) | 7.5 (62.9) | 8.9 (53.4) | 16.3 (108.3) |
| $CC_{1/2}$ (%) | 99.4 (80.2) | 99.6 (38.1) | 99.2 (51.6) | 98.0 (23.9) |
| $\langle I/\sigma(I) \rangle$ | 5.3 (2.2) | 8.9 (1.4) | 6.9 (1.5) | 4.8 (0.8) |
| No. of reflections | 40255 (3197) | 77138 (4614) | 72135 (5250) | 49726 (4544) |
| No. of unique reflections | 14021 (1184) | 22598 (1487) | 20843 (1519) | 14557 (1308) |
| Multiplicity | 2.9 (2.7) | 3.4 (3.1) | 3.5 (3.5) | 3.4 (3.5) |
| Completeness (%) | 88.4 (85.1) | 98.4 (96.0) | 98.9 (99.2) | 98.5 (97.5) |
| $R_{iso}$ (%)$^{‡}$ | N.A. | N.A. | N.A. | 21.92 (39.98) |
| **Refinement statistics** | | | | |
| **Refinement strategy**$^{\#}$ | Classic | Classic | Classic | Extrapolated (occ 45%) |
| Diffraction limit (Å) | 40.25 – 2.2 (2.28 – 2.2) | 55.61 – 1.94 (2.03 – 1.94) | 55.51 – 2.0 (2.05 – 2.00) | 34.4 – 2.24 (2.41 – 2.24- |
| No. of reflections | 13927 (1340) | 22593 (2775) | 20833 (1462) | 14445 (2665) |
| R-work (%) | 17.21 | 16.27 | 15.14 | 27.97 |
| R-free (%)$^{§}$ | 22.17 | 21.22 | 20.89 | 32.26 |
| Number of atoms | 2728 | 2732 | 2872 | 2593 |
| macromolecules | 2500 | 2492 | 2600 | 2406 |
| ligands | 42 | 42 | 42 | 42 |
| solvent | 186 | 198 | 230 | 145 |
| Protein residues | 309 | 309 | 310 | 309 |
| RMS | | | | |

| | | | | |
|---|---|---|---|---|
| bonds (Å) | 0.010 | 0.014 | 0.010 | 0.015 |
| angles (°) | 0.52 | 0.97 | 0.82 | 0.55 |
| Ramachandran (%) | | | | |
| favored | 99.01 | 99.34 | 98.69 | 96.39 |
| allowed | 0.66 | 0.33 | 0.98 | 3.61 |
| outliers | 0.33 | 0.33 | 0.33 | 0.00 |
| Rotamer outliers (%) | 1.85 | 1.12 | 3.19 | 0.78 |
| Clashscore | 3.96 | 2.98 | 3.23 | 1.24 |
| Average B-factor ($Å^2$) | 33.63 | 31.07 | 29.08 | 24.60 |
| macromolecules | 33.25 | 30.30 | 28.25 | 24.84 |
| ligands | 22.87 | 19.83 | 17.61 | 18.19 |
| solvent | 41.11 | 43.09 | 40.64 | 22.52 |

*Values between parentheses refer to the highest resolution shell; [§]Rfree is calculated using 5 % of random reflections excluded from refinement; ‡ Riso is defined as $\sum |F_{obs}^{dark} - F_{obs}^{light}| / \sum |F_{obs}^{dark} + F_{obs}^{light}|/2$

[#]Refinement has been carried out in extrapolated structure factor amplitudes (“extrapolated”) or the original structure factor amplitudes/intensities (classic); N.A.: Not applicable

**Supplementary Table 6**: Photophysical and biophysical properties of the OCP mutants in complex with CAN or ECN.

| | Photoactivation rate (change in initial $OD^{550nm}$ /sec) | Recovery rate (change in initial $OD^{550nm}$ /sec) | Fold-change in photoactivation rate compared to WT | Fold-change in recovery rate compared to WT | Steady-state accumulation of the $OCP^{R}$ state (% change compared to WT) | Concentration - dependent peak-shift in temperature-controlled scanning-fluorimetry (TCSF) | $1^{st}$ TCSF peak (°C; informing on the recoiling of the αGH-loop in the $OCP^{O}$ monomer) | $2^{nd}$ TCSF peak (°C; informing on the recoiling of the αGH-loop in the $OCP^{O}$ dimer) | $3^{rd}$ TCSF peak (°C; informing on the denaturation of OCP) |
|---|---|---|---|---|---|---|---|---|---|
| **$WT_{CAN}$** | 19.3 | -0.3 | 1 | 1 | 100 | Yes | 42.12±0.20 | 47.13±0.06 | 53.64±0.21 |
| $A133P_{CAN}$ | 19.1 | -0.2 | 0.99 | 0.67 | 93.32±5.88 | No | 40.37±1.59 | *na* | 53.90±0.21 |
| $A23C_{CAN}$ | 6.9 | -0.6 | 0.36 | 2 | 81.33±2.84 | No | *na* | *na* | 58.59±0.24 |
| A23C-$A133P_{CAN}$ | 7.9 | -0.2 | 0.41 | 0.67 | 96.98±1.51 | No | 36.85±0.33 | *na* | 57.10±0.60 |
| $A38C_{CAN}$ | 8.3 | -4.5 | 0.43 | 15 | 51.11±0.97 | Yes | 44.60 | 46.86±0.39 | 53.31±1.97 |
| A38C-$I125C_{CAN}$ | *na* | *na* | *na* | *na* | 2.00±0.71 | Yes | *na* | *na* | 53.58±0.54 |
| $D35T_{CAN}$ | 12.2 | -0.4 | 0.63 | 1.33 | 92.47±2.01 | Yes | 38.10 | 45.59±0.55 | 50.93±0.25 |
| $I125C_{CAN}$ | 14.1 | -1.3 | 0.74 | 4.33 | 88.78±4.85 | Yes | 34.37±0.37 | 44.40 | 51.13±0.68 |
| $Q79L_{CAN}$ | 9.3 | -8.4 | 0.48 | 28 | 35.95±2.84 | No | 38.61±0.38 | *na* | 52.37±0.45 |
| $W41F_{CAN}$ | 9.6 | -0.9 | 0.50 | 3 | 78.47±1.93 | No | *na* | *na* | 52.9±0.44 |
| A38C-I125C-T80W-CAN | 2.8 | 11.7 | 0.14 | 39 | 15.21±1.12 | No | *na* | *Na* | 54.9±0.20 |

| | Photoactivation rate (change in initial $OD^{550nm}$ /sec) | Recovery rate (change in initial $OD^{550nm}$ /sec) | Fold-change in photoactivation rate compared to WT | Fold-change in recovery rate compared to WT | Steady-state accumulation of the $OCP^{R}$ state (% change compared to WT) | Concentration - dependent peak-shift in temperature-controlled scanning-fluorimetry (TCSF) | $1^{st}$ TCSF peak (°C; informing on the recoiling of the αGH-loop in the $OCP^{O}$ monomer) | $2^{nd}$ TCSF peak (°C; informing on the recoiling of the αGH-loop in the $OCP^{O}$ dimer) | $3^{rd}$ TCSF peak (°C; informing on the denaturation of OCP) |
|---|---|---|---|---|---|---|---|---|---|
| **$WT_{ECN}$** | 4.2 | -0.2 | 1 | 1 | 100 | No | *na* | *na* | 54.60±0.57 |
| A38C-$I125C_{ECN}$ | *na* | *na* | *na* | *na* | 0 | No | *na* | *na* | 53.99±0.26 |
| $Q79L_{ECN}$ | 0.7 | *na* | 0.23 | *na* | 1.44±0.53 | No | *na* | *na* | 51.27±0.80 |
| $D35T_{ECN}$ | 1.6 | -0.5 | 0.38 | 2.5 | 82.51±1.01 | No | *na* | *na* | 51.25±0.15 |

**Supplementary Table 7**. Data processing and refinement statistics for OCP TR-SFX experiments on the $P2_1$ crystals .

| | **OCP - P21 dark** | **OCP - P21 300fs - darkinter** | **OCP - P21 300fs** | **OCP - P21 1500fs - darkinter** | **OCP - P21 1500fs** | **OCP - P21 10ps - darkinter** | **OCP - P21 10ps** | **OCP - P21 100ps -darkinter** | **OCP - P21 100ps** |
|---|---|---|---|---|---|---|---|---|---|
| **PDB ID** | **30MQ** | | **30MR** | | **30MS** | | **30MT** | | **30PD** |
| **Data collection and processing statistics** | | | | | | | | | |
| Beamline | SwissFEL Alvra-Prime | SwissFEL Alvra-Prime | SwissFEL Alvra-Prime | SwissFEL Alvra-Prime | SwissFEL Alvra-Prime | SwissFEL Alvra-Prime | SwissFEL Alvra-Prime | SwissFEL Alvra-Prime | SwissFEL Alvra-Prime |
| Wavelength (Å) | 1.05 | 1.05 | 1.05 | 1.05 | 1.05 | 1.05 | 1.05 | 1.05 | 1.05 |
| Space group | $P\ 2_1$ | $P\ 2_1$ | $P\ 2_1$ | $P\ 2_1$ | $P\ 2_1$ | $P\ 2_1$ | $P\ 2_1$ | $P\ 2_1$ | $P\ 2_1$ |
| Unit cell parameters | | | | | | | | | |
| a, b, c (Å) | 63.5 ± 0.3<br>69.3 ± 0.4<br>77.0 ± 0.5 | 63.5 ± 0.5<br>69.7 ± 0.4<br>77.2 ± 0.8 | 63.4 ± 0.5<br>69.7 ± 0.4<br>77.2 ± 0.8 | 63.4 ± 0.3<br>69.6 ± 0.3<br>77.1 ± 0.4 | 63.4 ± 0.3<br>69.6 ± 0.3<br>77.1 ± 0.4 | 63.5 ± 0.3<br>69.6 ± 0.3<br>77.2 ± 0.4 | 63.5 ± 0.3<br>69.6 ± 0.3<br>77.2 ± 0.4 | 63.5 ± 0.4<br>69.6 ± 0.4<br>77.2 ± 0.5 | 63.5 ± 0.3<br>70.0 ± 0.4<br>77.2 ± 0.5 |
| $\alpha, \beta, \gamma$ (°) | 90 ± 0.4<br>101.8 ± 0.4<br>90 ± 0.4 | 90.1 ± 0.6<br>102.0 ± 0.8<br>90.1 ± 0.5 | 90.1 ± 0.6<br>102.0 ± 0.9<br>90.1 ± 0.5 | 90.0 ± 0.4.<br>101.9 ± 0.4<br>90.0 ± 0.4 | 90.0 ± 0.4.<br>101.9 ± 0.4<br>90.0 ± 0.4 | 90.0 ± 0.4<br>101.9 ± 0.4<br>90.1 ± 0.4 | 90.0 ± 0.4<br>101.9 ± 0.4<br>90.0 ± 0.4 | 90.0 ± 0.4<br>101.9 ± 0.4<br>90.0 ± 0.4 | 90.0 ± 0.4<br>101.9 ± 0.4<br>90.0 ± 0.4 |
| Diffraction limit (Å) | 8.00 - 2.20<br>(2.25 - 2.20) | 8.00 - 2.80<br>(2.86 - 2.80) | 8.00 - 2.80<br>(2.86 - 2.80) | 8.00 - 2.40<br>(2.45 - 2.40) | 8.00 - 2.40<br>(2.45 - 2.40) | 8.00 - 2.60<br>(2.66 - 2.60) | 8.00 - 2.60<br>(2.66 - 2.60) | 8.00 - 2.70<br>(2.76 - 2.70) | 8.00 - 2.70<br>(2.76 - 2.70) |
| $R_{split}$ (%) | 12.6 (116.5) | 74.5 (184.9) | 77.3 (181.7) | 31.5 (193.4) | 24.1 (1.22) | 37.8 (180.5) | 38.8 (202.7) | 39.5 (159.8) | 39.2 (162.0) |
| CC* (%) | 99.9 (75.8) | 80.4 (37.5) | 90.0 (58.1) | 98.9 (51.7) | 98.7 (90.5) | 97.6 (67.5) | 97.3 (59.4) | 97.0 (62.0) | 97.1 (60.4) |
| $\langle I/\sigma(I) \rangle$ | 6.0 (1.0) | 1.3 (0.6) | 1.3 (0.57) | 2.5 (0.6) | 2.5 (0.7) | 2.2 (0.6) | 2.2 (0.6) | 2.2 (0.8) | 2.2 (0.7) |
| No. of reflections | 20889536 (950718) | 329142 (16396) | 314099 (15632) | 1739784 (74670) | 1766003 (75265) | 790805 (35921) | 772368 (34790) | 600954 (29073) | 614571 (29318) |
| No. of unique reflections | 32705 (2167) | 15579 (1019) | 15577 (1019) | 25083 (1649) | 25083 (1649) | 19619 (1308) | 19619 (1308) | 17461 (1168) | 17461 (1168) |
| Multiplicity | 638.7 (438.7) | 21.1 (16.1) | 20.2 (15.3) | 69.4 (45.3) | 70.4 (45.6) | 40.3 (27.5) | 39.4 (26.6) | 34.4 (24.9) | 35.2 (25.1) |
| Completeness (%) | 100.0 (100.0) | 100.0 (100.0) | 100.0 (100.0) | 100.0 (100.0) | 100.0 (100.0) | 100.0 (100.0) | 100.0 (100.0) | 100.0 (100.0) | 100.0 (100.0) |
| $R_{iso}$ (%)‡ | N.A. | N.A. | 35.0 (43.4) | N.A. | 15.7 (31.1) | N.A. | 16.8 (32.4) | N.A. | 18.2 (35.4) |
| **Refinement statistics** | | | | | | | | | |
| **Refinement strategy#** | Classic | | Extrapolated (occ 40 %) | | Extrapolated (occ 20 %) | | Extrapolated (occ 20 %) | | Extrapolated (occ 20 %) |
| Diffraction limit (Å) | 10.00 – 2.20<br>(2.26 – 2.20) | | 7.99 – 2.80<br>(2.97 – 2.80) | | 7.99 – 2.40<br>(2.49 – 2.40) | | 7.99 – 2.60<br>(2.73 – 2.60) | | 7.99 – 2.70<br>(2.86 – 2.70) |

| | | | | | | | | | |
|---|---|---|---|---|---|---|---|---|---|
| No. of reflections | 33021 (2729) | | 15510 (2551) | | 25003 (2722) | | 19581 (2766) | | 17442 (2891) |
| R-work (%) | 15.83 | | 37.52 | | 36.10 | | 36.26 | | 37.60 |
| R-free (%)[§] | 20.23 | | 41.05 | | 41.01 | | 40.27 | | 40.74 |
| Number of atoms | 5431 | | 5061 | | 5135 | | 5122 | | 5187 |
| macromolecules | 5024 | | 4851 | | 4862 | | 4871 | | 4893 |
| ligands | 84 | | 84 | | 84 | | 84 | | 84 |
| solvent | 323 | | 126 | | 189 | | 167 | | 210 |
| Protein residues | 615 | | 619 | | 619 | | 619 | | 619 |
| RMS | | | | | | | | | |
| bonds (Å) | 0.009 | | 0.003 | | 0.003 | | 0.005 | | 0.005 |
| angles (°) | 0.92 | | 0.51 | | 0.43 | | 0.61 | | 0.66 |
| Ramachandran (%) | | | | | | | | | |
| favored | 98.35 | | 95.42 | | 98.04 | | 91.00 | | 96.56 |
| allowed | 1.65 | | 4.42 | | 1.96 | | 9.00 | | 3.27 |
| outliers | 0.00 | | 0.16 | | 0.00 | | 0.00 | | 0.16 |
| Rotamer outliers (%) | 2.77 | | 2.69 | | 0.00 | | 0 .96 | | 0.76 |
| Clashscore | 5.13 | | 6.95 | | 4.18 | | 13.76 | | 13.48 |
| Average B-factor (Å$^2$) | 45.70 | | 37.57 | | 41.57 | | 39.65 | | 42.57 |
| macromolecules | 45.46 | | 37.67 | | 42.39 | | 40.43 | | 43.54 |
| ligands | 33.09 | | 55.04 | | 33.34 | | 32.81 | | 33.96 |
| solvent | 52.68 | | 21.96 | | 24.03 | | 20.50 | | 23.26 |

| | **OCP - P21 dark** | **OCP - P21 10ns - darkinter** | **OCP - P21 10ns** | **OCP - P21 1us - darkinter** | **OCP - P21 1us** |
|---|---|---|---|---|---|
| **PDB ID** | **30MU** | **30OO** | **30MV** | **30MW** | **30MX** |
| **Data collection and processing statistics** | | | | | |
| Beamline | SwissFEL Cristallina-MX | SwissFEL Cristallina-MX | SwissFEL Cristallina-MX | SwissFEL Cristallina-MX | SwissFEL Cristallina-MX |

| | | | | | |
|---|---|---|---|---|---|
| Wavelength (Å) | 1.1 | 1.1 | 1.1 | 1.1 | 1.1 |
| Space group | $P\ 2_1$ | $P\ 2_1$ | $P\ 2_1$ | $P\ 2_1$ | $P\ 2_1$ |
| Unit cell parameters | | | | | |
| a, b, c (Å) | 63.9 ± 1.4<br>70.2 ± 0.3<br>77.5 ± 1.2 | 64.3 ± 3.2<br>70.2 ± 0.3<br>77.2 ± 2.5 | 64.2 ± 2.8<br>70.2 ± 0.3<br>77.3 ± 2.2 | 63.8 ± 0.3<br>70.2 ± 0.4<br>77.6 ± 0.4 | 63.8 ± 0.3.<br>70.2 ± 0.4.<br>77.6 ± 0.4 |
| $\alpha, \beta, \gamma$ (◦) | 90.0 ± 0.3<br>102.1 ± 1.2<br>90.0 ± 0.4 | 90.0 ± 0.1<br>102.4 ± 2.6<br>90.0 ± 0.1 | 90.0 ± 0.1<br>102.3 ± 2.3<br>90.0 ± 0.1 | 90.0 ± 0.5<br>102.0 ± 0.4<br>90.1 ± 0.5 | 90.0 ± 0.5<br>102.0 ± 0.4<br>90.1 ± 0.5 |
| Diffraction limit (Å) | 40.00 - 1.70<br>(1.74 - 1.70) | 40.00 - 1.70<br>(1.74 - 1.70) | 40.00 - 1.70<br>(1.74 - 1.70) | 40.00 - 1.80<br>(1.84 - 1.80) | 40.00 - 1.80<br>(1.84 - 1.80) |
| $R_{split}$ (%) | 3.1 (163.7) | 8.9 (160.4) | 9.09 (171.97) | 8.22 (298.41) | 8.44 (375.41) |
| CC* (%) | 100.0 (73.3) | 99.8 (68.8) | 99.8 (67.8) | 99.9 (62.4) | 99.9 (54.4) |
| $<I/\sigma(I)>$ | 12.0 (0.7) | 6.29 (0.71) | 6.29 (0.66) | 6.37 (0.39) | 6.09 (0.31) |
| No. of reflections | 176776669<br>(7195347) | 36257817<br>(1428207) | 35749924<br>(1410475) | 40916197<br>(1750876) | 40349872<br>(1728274) |
| No. of unique reflections | 73152 (4825) | 73898 (4867) | 73898 (4867) | 62321 (4125) | 62321 (4125) |
| Multiplicity | 2416.6 (1491.2) | 490.1 (293.4) | 483.8 (289.8) | 656.54 (424.5) | 647.5 (419.0) |
| Completeness (%) | 100.0 (100.0) | 100.0 (100.0) | 100.0 (100.0) | 100.0 (100.0) | 100.0 (100.0) |
| $R_{iso}$ (%)‡ | N.A. | N.A. | 7.2 (25.1) | N.A. | 7.8 (33.6) |
| **Refinement statistics** | | | | | |
| **Refinement strategy#** | Classic | Classic | Extrapolated<br>(occ 7 %) | Classic | Extrapolated<br>(occ 7 %) |
| Diffraction limit (Å) | 37.99 – 1.70<br>(1.72 – 1.70) | 43.97 – 1.70<br>(1.72 – 1.70) | 14.93 – 1.81<br>(1.87 – 1.81) | 28.53 – 1.80<br>(1.83 – 1.80) | 14.93 – 1.80<br>(1.85 – 1.80) |
| No. of reflections | 73613 (2823) | 73749 (2844) | 44931 (739) | 61094 (2444) | 53613 (2207) |
| R-work (%) | 17.28 | 17.54 | 30.50 | 17.93 | 39.45 |
| R-free (%)§ | 19.34 | 21.07 | 34.15 | 22.18 | 42.86 |
| Number of atoms | 5540 | 5647 | 5534 | 5599 | 5263 |
| macromolecules | 5052 | 5101 | 5078 | 5112 | 4935 |
| ligands | 84 | 84 | 84 | 84 | 84 |
| solvent | 404 | 462 | 372 | 403 | 244 |
| Protein residues | 619 | 619 | 619 | 619 | 615 |

| | | | | | |
|---|---|---|---|---|---|
| RMS | | | | | |
| bonds (Å) | 0.007 | 0.011 | 0.004 | 0.011 | 0.003 |
| angles (°) | 0.80 | 0.88 | 0.48 | 0.89 | 0.54 |
| Ramachandran (%) | | | | | |
| favored | 99.18 | 99.18 | 97.05 | 99.18 | 93.74 |
| allowed | 0.82 | 0.82 | 2.95 | 0.82 | 5.77 |
| outliers | 0.00 | 0.00 | 0.00 | 0.00 | 0.49 |
| Rotamer outliers (%) | 2.20 | 3.09 | 4.40 | 2.54 | 1.32 |
| Clashscore | 1.37 | 2.14 | 2.83 | 1.74 | 5.72 |
| Average B-factor ($Å^2$) | 43.01 | 39.26 | 31.03 | 45.06 | 31.85 |
| macromolecules | 42.21 | 38.34 | 31.08 | 44.29 | 32.06 |
| ligands | 29.39 | 25.43 | 27.44 | 30.35 | 26.88 |
| solvent | 55.74 | 51.97 | 31.11 | 57.99 | 29.27 |

*Values between parentheses refer to the highest resolution shell; §Rfree is calculated using 5 % of random reflections excluded from refinement; ‡ Riso is defined as $\sum |F_{obs}^{dark} - F_{obs}^{light}| \Big/ \sum |F_{obs}^{dark} + F_{obs}^{light}|/2$

#Refinement has been carried out in extrapolated structure factor amplitudes (“extrapolated”) or the original structure factor amplitudes/intensities (classic); N.A.: Not applicable.

**Supplementary Table 8**. Data processing and refinement statistics for OCP TR-SFX experiments on the C2 crystals.

| | **OCP – C2 dark** | **OCP - C2 300fs - darkinter** | **OCP - C2 300fs** | **OCP - C2 1500fs - darkinter** | **OCP - C2 1500fs** | **OCP - C2 10ps - darkinter** | **OCP - C2 10ps** | **OCP - C2 100ps - darkinter** | **OCP - C2 100ps** |
|---|---|---|---|---|---|---|---|---|---|
| **PDB ID** | **30MY** | | **30MZ** | | **30NA** | | **30NB** | | **30NC** |
| **Data collection and processing statistics** | | | | | | | | | |
| Beamline | SwissFEL Alvra-Prime | SwissFEL Alvra-Prime | SwissFEL Alvra-Prime | SwissFEL Alvra-Prime | SwissFEL Alvra-Prime | SwissFEL Alvra-Prime | SwissFEL Alvra-Prime | SwissFEL Alvra-Prime | SwissFEL Alvra-Prime |
| Wavelength (Å) | 1.05 | 1.05 | 1.05 | 1.05 | 1.05 | 1.05 | 1.05 | 1.05 | 1.05 |
| Space group | *C 2* | *C 2* | *C 2* | *C 2* | *C 2* | *C 2* | *C 2* | *C 2* | *C 2* |
| Unit cell parameters | | | | | | | | | |
| a, b, c (Å) | 82.5 ± 0.3<br>68.8 ± 0.5<br>62.7 ± 0.2 | 82.8 ± 0.3<br>69.2 ± 0.3<br>62.9 ± 0.2 | 82.8 ± 0.3<br>69.2 ± 0.3<br>62.9 ± 0.2 | 825 ± 0.3,<br>68.9 ± 0.4<br>62.8 ± 0.2 | 82.5 ± 0.3<br>68.9 ± 0.4<br>62.8 ± 0.2 | 82.5 ± 0.3<br>68.8 ± 0.5<br>62.7 ± 0.2 | 82.5 ± 0.3<br>68.8 ± 0.5<br>62.7 ± 0.2 | 82.4 ± 0.3<br>68.7 ± 0.4<br>62.7 ± 0.2 | 82.4 ± 0.3<br>68.7 ± 0.4<br>62.7 ± 0.2 |
| $\alpha, \beta, \gamma$ (°) | 90.1 ± 0.3<br>117.6 ± 0.2<br>89.8 ± 0.3 | 90.1 ± 0.3<br>117.5 ± 0.2<br>89.8 ± 0.2 | 90.1 ± 0.3<br>117.5 ± 0.<br>89.8 ± 0.2 | 90.1 ± 0.3<br>117.6 ± 0.2<br>89.8 ± 0.2 | 90.1 ± 0.3<br>117.6 ± 0.2<br>89.8 ± 0.2 | 90.1 ± 0.3<br>117.6 ± 0.2<br>89.8 ± 0.3 | 90.1 ± 0.3<br>117.6 ± 0.2<br>89.8 ± 0.3 | 90.1 ± 0.3<br>117.6 ± 0.2<br>89.8 ± 0.3 | 90.1 ± 0.3<br>117.6 ± 0.2<br>89.8 ± 0.3 |
| Diffraction limit (Å) | 50.00 - 1.60 (1.64 - 1.60) | 8.00 - 1.67 (1.71 - 1.67) | 8.00 - 1.67 (1.71 - 1.67) | 8.00 - 1.67 (1.71 - 1.67) | 8.00 - 1.67 (1.71 - 1.67) | 8.00 - 1.67 (1.71 - 1.67) | 8.00 - 1.67 (1.71 - 1.67)* | 8.00 - 1.67 (1.71 - 1.67) | 8.00 - 1.67 (1.71 - 1.67) |
| $R_{split}$ (%) | 3.0 (57.9) | 8.5 (128.4) | 8.7 (132.1) | 7.0 (102.1) | 6.9 (95.1) | 9.6 (144.1) | 9.6 (158.8) | 8.1 (140.9) | 8.2 (129.4) |
| CC* (%) | 99.9 (91.3) | 99.9 (73.0) | 99.9 (69.5) | 99.9 (79.6) | 99.9 (79.6) | 99.9 (65.9) | 99.6 (25.4) | 99.9 (68.4) | 99.9 (68.3) |
| $\langle I/\sigma(I) \rangle$ | 21.6 (2.0) | 7.4 (0.9) | 7.2 (0.9) | 9.1 (1.2) | 9.0 (1.2) | 6.6 (0.8) | 6.5 (0.7) | 7.7 (0.9) | 7.7 (0.9) |
| No. of reflections | 321102200 (15208894) | 31117368 (1487068) | 29515513 (1411620) | 47912674 (2292312) | 47057813 (2253353) | 26941230 (1291549) | 26086939 (1247812) | 36896773 (1769111) | 36180851 (1731259) |
| No. of unique reflections | 41250 (2740) | 35975 (2371) | 35975 (2371) | 35975 (2371) | 35975 (2371) | 35975 (2371) | 35975 (2371) | 35975 (2371) | 35975 (2371) |
| Multiplicity | 7784.3 (5550.7) | 865.0 (627.1) | 820.4 (595.4) | 1331.8 (966.8) | 1308.1 (950.4) | 748.9 (544.7) | 725.1 (526.3) | 1025.6 (746.1) | 1005.7 (730.2) |
| Completeness (%) | 100.0 (100.0) | 100.0 (100.0) | 100.0 (100.0) | 100.0 (100.0) | 100.0 (100.0) | 100.0 (100.0) | 100.0 (100.0) | 100.0 (100.0) | 100.0 (100.0) |
| $R_{iso}$ (%)$^{‡}$ | N.A. | N.A. | 7.5 (29.2) | N.A. | 6.3 (25.9) | N.A. | 8.1 (31.6) | N.A. | 7.4 (29.4) |
| **Refinement statistics** | | | | | | | | | |
| **Refinement strategy$^{\#}$** | Classic | | Extrapolated (occ 7 %) | | Extrapolated (occ 7 %) | | Extrapolated (occ 7 %) | | Extrapolated (occ 7 %) |
| Diffraction limit (Å) | 9.98 – 1.60 (1.64 – 1.60) | | 7.99 – 1.67 (1.71 – 1.67) | | 7.999 – 1.67 (1.71 – 1.67) | | 7.99 – 1.67 (1.71 – 1.67) | | 7.99 – 1.67 (1.71 – 1.67) |

| | | | | | | | | | |
|---|---|---|---|---|---|---|---|---|---|
| No. of reflections | 41084 (2727) | | 35740 (2540) | | 35959 (2756) | | 35889 (2696) | | 35948 (2754) |
| R-work (%) | 15.08 | | 34.98 | | 34.04 | | 35.93 | | 33.91 |
| R-free (%)[§] | 17.38 | | 39.25 | | 41.69 | | 42.91 | | 40.45 |
| Number of atoms | 3030 | | 3041 | | 2882 | | 2842 | | 2849 |
| macromolecules | 2711 | | 2547 | | 2563 | | 2547 | | 2561 |
| ligands | 42 | | 42 | | 42 | | 42 | | 42 |
| solvent | 277 | | 452 | | 277 | | 253 | | 246 |
| Protein residues | 309 | | 309 | | 309 | | 309 | | 309 |
| RMS | | | | | | | | | |
| bonds (Å) | 0.006 | | 0.005 | | 0.004 | | 0.004 | | 0.013 |
| angles (°) | 0.77 | | 0.60 | | 0.59 | | 0.48 | | 1.25 |
| Ramachandran (%) | | | | | | | | | |
| favored | 99.34 | | 96.72 | | 97.38 | | 95.41 | | 94.75 |
| allowed | 0.33 | | 2.95 | | 2.62 | | 4.59 | | 4.26 |
| outliers | 0.33 | | 0.33 | | 0.00 | | 0.00 | | 0.98 |
| Rotamer outliers (%) | 3.06 | | 6.93 | | 3.97 | | 6.18 | | 11.59 |
| Clashscore | 2.73 | | 9.11 | | 5.99 | | 6.59 | | 10.26 |
| Average B-factor ($Å^2$) | 28.64 | | 27.63 | | 28.52 | | 27.66 | | 25.93 |
| macromolecules | 27.22 | | 27.66 | | 28.50 | | 28.11 | | 26.43 |
| ligands | 18.83 | | 17.58 | | 15.93 | | 18.20 | | 16.47 |
| solvent | 44.02 | | 28.39 | | 30.69 | | 24.68 | | 22.36 |

| | OCP - C2 dark | OCP - C2 10ns - darkinter | OCP - C2 10ns | OCP - C2 1us - darkinter | OCP - C2 1us |
|---|---|---|---|---|---|
| **PDB ID** | **30ND** | **30NF** | **30NE** | **30NG** | **30NH** |
| **Data collection and processing statistics** | | | | | |
| Beamline | Cristallina | Cristallina | Cristallina | Cristallina | Cristallina |
| Wavelength (Å) | 1.1 | 1.1 | 1.1 | 1.1 | 1.1 |
| Space group | C 1 2 1 | C 1 2 1 | C 1 2 1 | C 1 2 1 | C 1 2 1 |
| Unit cell parameters | | | | | |
| a, b, c (Å) | 83.1 ± 0.2<br>69.4 ± 0.3<br>63.2 ± 0.2 | 83.1 ± 0.3<br>69.4 ± 0.3<br>63.2 ± 0.2 | 83.1 ± 0.2<br>69.4 ± 0.3<br>63.2 ± 0.0 | 83.1 ± 0.3<br>69.4 ± 0.3<br>63.2 ± 0.2 | 83.1 ± 0.3<br>69.4 ± 0.3<br>63.2 ± 0.2 |
| $\alpha, \beta, \gamma$ (°) | 90.1 ± 0.3<br>117.5 ± 0.2<br>89.9 ± 0.2 | 90.0 ± 0.4<br>117.5 ± 0.3<br>89.9 ± 0.2 | 90.0 ± 0.1<br>117.5 ± 0.1<br>89.9 ± 0.1 | 90.0 ± 0.4<br>117.5 ± 0.3<br>89.8 ± 0.2 | 90.0 ± 0.4<br>117.5 ± 0.3<br>89.8 ± 0.2 |
| Diffraction limit (Å) | 40 - 1.52<br>(1.55 - 1.50) | 40 – 1.57<br>(1.61 – 1.55) | 40 - 1.57<br>(1.61 - 1.55) | 40 – 1.57<br>(1.61 – 1.57) | 40 - 1.57<br>(1.61 - 1.55) |
| $R_{split}$ (%) | 3.7 (49.4) | 7.1 (59.3) | 5.7 (54.2) | 7.9 (108.9) | 8.15 (132.95) |
| CC* (%) | 99.8 (77.2) | 99.8 (79.6) | 99.9 (85.4) | 99.8 (77.1) | 99.8 (71.3) |
| $\langle I/\sigma(I) \rangle$ | 16.0 (2.4) | 9.36 (2.26) | 11.1 (2.3) | 7.2 (1.0) | 7.0 (0.9) |
| No. of reflections | 152745942 (6798532) | 36525661 (1691607) | 58000433 (2689957) | 28117870 (1304103) | 27840378 (1290152) |
| No. of unique reflections | 50626 (3364) | 46211 (3077) | 46211 (3077) | 46211 (3077) | 46211 (3077) |
| Multiplicity | 3017.1 (2021.0) | 790.4 (549.8) | 1255.1 (874.2) | 608 (423.8) | 602.5 (419.3) |
| Completeness (%) | 100.0 (100.0) | 100.0 (100.0) | 100.0 (100.0) | 100.0 (100.0) | 100.0 (100.0) |
| $R_{iso}$ (%)[‡] | N.A. | N.A. | 4.7 (18.3) | N.A. | 6.3 (32.5) |
| **Refinement statistics** | | | | | |
| **Refinement strategy[#]** | Classic | Classic | Extrapolated<br>(3.5% occ) | Classic | Extrapolated<br>(occ 3.5 %) |
| Diffraction limit (Å) | 28.65 – 1.50<br>(1.53 – 1.50) | 29.94 – 1.55<br>(1.60 – 1.55) | 14.97 – 1.67<br>(1.72 – 1.67) | 28.65 – 1.55<br>(1.58 – 1.55) | 14.97 – 1.60<br>(1.64 – 1.60) |
| No. of reflections | 50883 (2788) | 46159 (2858) | 34215 (2207) | 46135 (2854) | 37629 (2138) |
| R-work (%) | 15.58 | 15.50 | 39.76 | 15.85 | 41.12 |

| | | | | | |
|---|---|---|---|---|---|
| R-free (%)[§] | 17.78 | 17.64 | 43.67 | 18.65 | 45.08 |
| Number of atoms | 3214 | 3215 | 2852 | 3214 | 2799 |
| macromolecules | 2883 | 2883 | 2579 | 2883 | 2561 |
| ligands | 42 | 42 | 42 | 42 | 42 |
| solvent | 289 | 290 | 231 | 289 | 196 |
| Protein residues | 309 | 309 | 309 | 309 | 309 |
| RMS | | | | | |
| bonds (Å) | 0.009 | 0.009 | 0.003 | 0.009 | 0.003 |
| angles (°) | 0.86 | 0.84 | 0.41 | 0.88 | 0.42 |
| Ramachandran (%) | | | | | |
| favored | 99.02 | 98.36 | 95.41 | 99.02 | 95.74 |
| allowed | 0.98 | 1.64 | 4.26 | 0.98 | 4.26 |
| outliers | 0.00 | 0.00 | 0.33 | 0.00 | 0.00 |
| Rotamer outliers (%) | 1.93 | 2.89 | 1.81 | 2.89 | 2.90 |
| Clashscore | 2.92 | 2.40 | 5.74 | 3.26 | 5.62 |
| Average B-factor ($Å^2$) | 30.59 | 29.82 | 22.16 | 33.29 | 19.81 |
| macromolecules | 29.08 | 28.24 | 22.75 | 33.75 | 20.20 |
| ligands | 20.25 | 18.69 | 17.32 | 22.61 | 15.69 |
| solvent | 47.14 | 47.21 | 16.46 | 50.24 | 15.48 |

*Values between parentheses refer to the highest resolution shell; [§]Rfree is calculated using 5 % of random reflections excluded from refinement; ‡ Riso is defined as $\sum |F_{obs}^{dark} - F_{obs}^{light}| \Big/ \sum |F_{obs}^{dark} + F_{obs}^{light}|/2$

[#]Refinement has been carried out in extrapolated structure factor amplitudes (“extrapolated”) or the original structure factor amplitudes/intensities (classic); N.A.: Not applicable.

**Supplementary Table 9**. Data processing and refinement statistics for OCP SSX experiments.

| | **OCP – P21**<br>**dark** | **OCP – P21**<br>**light – 48 h** | **OCP - C2**<br>**dark** | **OCP - C2**<br>**light – 48h I** | **OCP - C2**<br>**light – 48h II** |
|---|---|---|---|---|---|
| **PDB ID** | **30NI** | **30NJ** | **30NK** | **30NL** | **30NM** |
| **Data collection and processing statistics** | | | | | |
| Beamline | ESRF ID23-EH2 | ESRF ID23-EH2 | ESRF ID23-EH2 | ESRF ID23-EH2 | ESRF ID23-EH2 |
| Wavelength (Å) | 0.8731 | 0.8731 | 0.8731 | 0.8731 | 0.8731 |
| Space group | $P\ 2_1$ | $P\ 2_1$ | *C 2* | *C 2* | *C 2* |
| Unit cell parameters | | | | | |
| a, b, c (Å) | 63.0 ± 0.4<br>69.1 ± 0.6<br>77.2 ± 0.6 | 63.1 ± 0.3<br>69.1 ± 0.6<br>77.3 ± 0.56 | 81.9 ± 0.4<br>68.2 ± 0.5<br>62.3 ± 0.3 | 82.0 ± 0.5<br>68.4 ± 0.5<br>62.5 ± 0.3 | 89.2 ± 0.9<br>69.0 ± 0.6<br>63.1 ± 0.3 |
| $\alpha, \beta, \gamma$ (°) | 90.0 ± 0.4<br>101.5 ± 0.4<br>90.0 ± 0.4 | 90.0 ± 0.4<br>101.5 ± 0.3<br>90.0 ± 0.4 | 90.0 ± 0.4<br>117.5 ± 0.3<br>90.0 ± 0.4 | 90.1 ± 0.4<br>117.6 ± 0.3<br>90.0 ± 0.4 | 90.0 ± 0.5<br>122.0 ± 0.5<br>90.0 ± 0.5 |
| Diffraction limit (Å) | 50.00 - 2.23<br>(2.28 - 2.23) | 50.00 - 2.43 (2.49 - 2.43) | 50.00 - 1.80<br>(1.82 - 1.80) | 50.00 - 1.90<br>(1.97 - 1.92) | 50.00 - 2.20<br>(2.45 - 2.40) |
| $R_{split}$ (%) | 34.73 (160.55) | 23.28 (149.21) | 13.0 (80.95) | 18.7 (96.0) | 26.0 (229.6) |
| CC* (%) | 97.1 (64.84) | 98.94 (70.56) | 99.53 (81.05) | 99.0 (77.8) | 98.6 (53.0) |
| $\langle I/\sigma(I) \rangle$ | 2.04 (0.66) | 2.87 (0.74) | 5.20 (1.36) | 3.69 (1.10) | 2.42 (0.45) |
| No. of reflections | 1516017 (68455) | 1980876 (91212) | 4798384 (215716) | 1682005 (71012) | 866357 (40580) |
| No. of unique reflections | 34215 (2257) | 26462 (1774) | 28329 (1906) | 24218 (1574) | 16647 (1125) |
| Multiplicity | 44.31 (30.33) | 74.85 (51.42) | 169.4 (113.2) | 69.5 (45.1) | 52.0 (36.1) |
| Completeness (%) | 99.98 (100.0) | 100 (100) | 100.0 (100.0) | 100 (100) | 100 (100) |
| $R_{iso}$ (%)$^{\ddagger}$ | N.A. | 19.88 (28.18) | N.A. | 9.0 (15.0) | N.A |
| **Refinement statistics** | | | | | |
| **Refinement strategy**$^{\#}$ | Classic | Extrapolated (occ 27.9%) | Classic | Extrapolated (occ 20%) | Classic |
| Diffraction limit (Å) | 46.53 – 2.20<br>(2.26 – 2.20) | 35.96 – 2.40<br>(2.50 – 2.40) | 49.73 – 1.80<br>(1.86 – 1.80) | 19.41 – 2.40<br>(2.64 – 2.40) | 46.10 – 2.40<br>(2.59 – 2.40) |
| No. of reflections | 33997 (2750) | 26385 (2939) | 28194 (2786) | 11976 (2978) | 12777 (2512) |
| R-work (%) | 22.89 | 37.51 | 16.18 | 29.22 | 31.98 |

| | | | | | |
|---|---|---|---|---|---|
| R-free (%)[§] | 26.75 | 41.31 | 19.00 | 33.97 | 35.32 |
| Number of atoms | 5206 | 5047 | 2947 | 2681 | 2658 |
| macromolecules | 4896 | 4866 | 2677 | 2498 | 2417 |
| ligands | 84 | 84 | 42 | 42 | 42 |
| solvent | 226 | 97 | 228 | 141 | 199 |
| Protein residues | 621 | 621 | 310 | 308 | 302 |
| RMS | | | | | |
| bonds (Å) | 0.004 | 0.004 | 0.008 | 0.005 | 0.005 |
| angles (°) | 0.47 | 0.58 | 0.66 | 0.55 | 0.56 |
| Ramachandran (%) | | | | | |
| favored | 98.86 | 95.27 | 99.67 | 97.37 | 91.28 |
| allowed | 1.14 | 4.24 | 0.33 | 2.63 | 6.71 |
| outliers | 0.00 | 0.49 | 0.00 | 0.00 | 2.01 |
| Rotamer outliers (%) | 2.09 | 2.29 | 2.41 | 3.35 | 9.69 |
| Clashscore | 2.64 | 7.65 | 3.13 | 12.09 | 13.96 |
| Average B-factor ($Å^2$) | 47.91 | 49.50 | 32.01 | 39.21 | 57.80 |
| macromolecules | 48.07 | 49.96 | 30.90 | 39.63 | 58.26 |
| ligands | 33.69 | 38.87 | 22.18 | 40.65 | 52.39 |
| solvent | 49.94 | 35.63 | 46.77 | 31.28 | 53.27 |

*Values between parentheses refer to the highest resolution shell; [§]Rfree is calculated using 5 % of random reflections excluded from refinement; ‡ Riso is defined as $\sum |F_{obs}^{dark} - F_{obs}^{light}| / \sum |F_{obs}^{dark} + F_{obs}^{light}|/2$

[#]Refinement has been carried out in extrapolated structure factor amplitudes (“extrapolated”) or the original structure factor amplitudes/intensities (classic); N.A.: Not applicable.
