## Supplementary material for "Structural basis of the two-photon photoactivation mechanism of orange carotenoid protein": Materials and Methods

for

by

Rory Munro, Elena A. Andreeva[#], Elisabeth Hartmann[#], Quentin Goor[◊], Hosni El Zein[◊], Stanislaw Nizinski, Adjélé Wilson, Elke De Zitter, Gregory Effantin, Nicolas Coquelle, Ninon Zala, Martin V. Appleby, Shira Bar-Zvi, Camila Bacellar, Emma Beale, Emmanuelle Bignon, Bernhard Brutscher, Martin Byrdin, Claudio Cirelli, Florian Dworkowski, Lutz Foucar, Guillaume Gotthard, Alexander Gorel, Marie Luise Grünbein, Mario Hilpert, Philip J.M. Johnson, Marco Kloos, Gregor Knopp, Karol Nass, Gabriela Nass Kovacs, Dmitry Ozerov, Christopher J. Milne, Gotard Burdzinski, Christophe Chipot, Yasaman Karami, François Dehez, Martin Weik, R. Bruce Doak, Robert L. Shoeman, Giorgio Schiro, Michel Sliwa, Diana Kirilovsky, Ilme Schlichting[*] and Jacques-Philippe Colletier[*]

[#] these authors contributed equally.
[◊] these authors contributed equally.
[*] correspondance to

**Authors affiliations:**

[1]**Institut de Biologie Structurale, Grenoble, F-38000 France**:

Rory Munro, Elena A. Andreeva, Quentin Goor, Hosni El Zein, Elke De Zitter, Nicolas Coquelle, Gregory Effantin, Ninon Zala, Bernhard Brutscher, Martin Byrdin, Martin Weik, Giorgio Schiro, Jacques-Philippe Colletier

[2]**Max Planck Institute for Medical Research, D-69120 Heidelberg, Germany**:

Elisabeth Hartmann, Elena A. Andreeva, Marco Kloos, Marie Luise Grünbein, Stanislaw Nizinski, Alexander Gorel, Lutz Foucar, Mario Hilpert, Gabriela Nass Kovacs, R. Bruce Doak, Robert L. Shoeman, Ilme Schlichting

[3]**Institute for Integrative Biology of the Cell, Gif-sur-Yvette, France**:

Adjélé Wilson, Shira Bar-Zvi, Diana Kirilovsky

[4] **SwissFEL, SLS, Paul Scherrer Institute, Villigen, Switzerland**:

Dmitry Ozerov, Karol Nass, Philip J.M. Johnson, Emma Beale, Claudio Cirelli, Gregor Knopp, Camila Bacellar, Christopher J. Milne, Martin V. Appleby, Guillaume Gotthard, Florian Dworkowski

[5] **Université de Lorraine, CNRS, LPCT, F-54000 Nancy, France**:

Christophe Chipot, Emmanuelle Bignon, François Dehez

[6] **European XFEL, Hamburg, Germany**:

Marco Kloos, Christopher J. Milne

[7] **Quantum Electronics Laboratory, Faculty of Physics and Astronomy, Adam Mickiewicz University, Uniwersytetu Poznańskiego 2, Poznan 61-614, Poland**

Stanislaw Nizinski, Gotard Burdzinski

[8] **Theoretical and Computational Biophysics Group, Beckman Institute, and Department of Physics, University of Illinois at Urbana-Champaign, Urbana, Illinois 61801, USA;** [9] **Department of Biochemistry and Molecular Biology, The University of Chicago, Chicago, Illinois 60637, USA:**

Christophe Chipot

[9] **Université de Lorraine, CNRS, Inria, LORIA, F-54000 Nancy, France**:

Yasaman Karami

[10] **Laboratoire d'Optique et Biosciences, CNRS, Inserm, Ecole polytechnique, Institut Polytechnique de Paris, 91120 Palaiseau, France**:

Michel Sliwa

***OCP mutagenesis and protein production***

The *Planktothrix aghardhii* OCP-containing pCDFDuet-1 plasmid vector was constructed in the lab of Dr. Diana Kirilovsky at the I2BC institute (Gif-sur-Yvette, France) using *EcoR1* and *HindIII* restriction sites, and likewise for the pAC-BETA and pBAD plasmids (expressing genes to synthesize β-carotene and ketolate the carotenoid, respectively)[1]. Plasmid amplification, 3-plasmid transformation into *E. coli* cells, holo-protein expression and purification were performed as described in[2]. Protein expression and purification were performed similarly as described previously[1], with exception of a very large batch that was produced for time-resolved serial femtosecond crystallography experiment at the *Alvra-prime* beamline of the SwissFEL (*vide infra*). Scaling up protein expression resulted in a large fraction of apo protein, which copurified with holoprotein by polyhistidine affinity chromatography. Most of the non-functional protein was removed by successively dialyzing the protein against 5 M NaCL, 40 mM Tris pH 7.4 followed by 0.15 M NaCl, 40 mM Tris pH 7.4 buffers, with three rounds of centrifugation at 13500 rpm after each dialysis step to eliminate the precipitated apo protein (white pellet) as well as misfolded holoprotein (violet pellet). Typically, this procedure resulted in an absorption ratio of $A_{500\ nm}/A_{280\ nm}$ ~ 1.5. The protein was further purified by crystallization. In all cases, the protein concentration was determined using an extinction coefficient at 500 nm of 63,000 $M^{-1}$ $cm^{-1}$ as determined in [3]. Protein samples were stored in 40 mM Tris pH 7.4, 150 mM NaCl, at either -80°C or 4°C.

To produce mutant variants, oligonucleotide primers were designed for the desired mutation, following which the 5 – 3' and 3 – 5' sequences (~ 30 – 40 bases each) were ordered lyophilized from Eurofins Genomics (Ebersberg, Germany). Point mutations were introduced using the QuikChange™ XL Site-Directed Mutagenesis Kit (Agilent Technologies, Santa Clara, CA, USA). After amplification, plasmids were purified using the mini-prep Plasmid DNA Purification kit (Macherey-Nagel, Düren, Germany), and then sequenced using the Eurofins Genomics LightRun service (Ebersberg, Germany).

***Steady-state absorption spectroscopy***

Steady-state dark and light spectra were recorded using a Specord S600 spectrophotometer (Analytik Jena, Jena, Germany). For all proteins (WT-OCP and mutants), spectra were recorded for the dark-adapted (before illumination) and light-adapted (after 5 minute of illumination) states, in a wavelength range of 250 to 800 nm with an integration time of 68 ms. Spectra were acquired from samples adjusted to an OD of 0.3 at 500 nm, using a quartz cuvette with 10 mm path length and a sample temperature of 8 °C. To record the light adapted spectra, samples were illuminated from above (30 mm distance to the sample) using a bright white lamp delivering 7,000 μM photons $s^{-1}$ $m^{-2}$ (KL 1500 electronic, Schott AG, Mainz, Germany).

To measure the photoactivation kinetics, spectra were recorded in a wavelength range of 500 to 550 nm with a 30 ms integration time. This narrow spectral window allowed spectra to be collected with short time delays between them. A dark spectrum was taken prior to illumination, then samples (WT-OCP and mutants) were illuminated under the conditions detailed above.

To follow the kinetics from the dark-state up to the eventual plateau of the final $OCP^R$ steady-state, spectra were collected in the following way: 25 spectra separated by 800 ms, followed by 140 spectra separated by 2 s. To measure the recovery kinetics, the lamp was turned off and spectra were recorded in the following order: 10 spectra separated by 4 s, 10 spectra separated by 8 s, 10 spectra separated by 20 s, and 22 spectra separated by 40 s, covering 1,200 s total. To evaluate photoactivation and recovery rates, absorbance at 550 nm was plotted as a function of time and initial slopes (3-4 first points) were determined and expressed as a fraction of $OCP^R$ formed per second.

***Isolation of PBS, OCP-PBS Complexes and fluorescence measurements***

*Planktothrix agardhii* OCP efficiently quenches the *Synechocystis PCC6803* phycobilisome[1]. Here, we used the basal part or core base (CB) of the phycobilisome, composed of only the allophycocyanin cylinders and linker proteins and featuring the binding epitope for OCP. The purification of CB phycobilisomes (CB-PBS) from *Synechocystis* PCC 6803 mutants was performed as previously described[4]. CB-PBS consists of an allophycocyanin core from which six rods extend, each containing one phycocyanin hexamer[5]. The OCP/CB-PBS complexes were prepared in 0.8 M K-phosphate buffer pH 7 by illuminating for 5 min at 23°C isolated *Synechocystis* PCC8603 CB-PBS in the presence of OCP (OCP:CB-PBS ratio = 20) with 5000 μM photons $m^{-2}$ $s^{-1}$ of white light. The PBS fluorescence emission spectra were measured at room temperature from 600 to 800 nm in either a CARY Eclipse fluorescence spectrophotometer fluorometer (Varian) using a 1 cm path length cuvette, or a Biotek Synergy-H4 96-well plate-reader using a volume of 50 μl per well. The sample was excited by 590 nm light.

***Temperature-controlled scanning-fluorimetry***

Protein samples were centrifuged at 39,000 g for 10 minutes at 4°C, before the supernatant was transferred to a new tube and diluted to concentrations of 30, 95, and 160 μM (45 μl total each). For each concentration, three Prometheus capillaries (NanoTemper Technologies, Munich, Germany) were filled with 15 μl of sample. The capillaries were loaded into a Prometheus Nanotemper instrument (NanoTemper Technologies, Munich, Germany) and a temperature gradient of 1°C/min from 15 – 90°C was set. The sample was excited at 280 nm with an excitation power of 30%, with the detection of fluorescence at 350 nm. We refer to these data as TCSF, because we here only monitor the fluorescence signal at 350 nm. By contrast, in differential scanning fluorimetry (nanoDSF) the ratio of the 350 and 330 nm fluorescence signals is determined which was in the present case fully uninformative. However, the un-exploitability of the nanoDSF signal provided hints that something specific to OCP was going on, which we further explored.

***Analytical ultracentrifugation***

Sedimentation velocity experiments were carried out in an XL-I analytical ultracentrifuge (Beckman Coulter, Brea, CA, USA) at 25 °C and 42,000 rpm, using either an AnTi-50 or AnTi-60 rotor (Beckman Coulter, Brea, CA, USA). OCP samples were adjusted to the required concentrations (5 μM for the constitutively dimeric A23C mutant, 160 μM for the WT protein and the A133P mutant) and loaded into 1.5-mm path-length Ti double-sector centerpieces equipped with sapphire windows (Nanolytics GmbH, Potsdam, Germany). The protein sample and reference buffers contained of 40 mM Tris pH 7.4 and 150 mM NaCl. Radial scans were recorded at 280 nm and 500 nm, as well as using interference optics (DMA 5000 densitometer and AMVn viscometer, Anton Paar, Graz, Austria). Data were processed using standard procedures implemented in Sedfit v17.0 (www.analyticalultracentrifugation.com). Buffer parameters, including the sedimentation coefficient corrected to 25 °C in water ($s_{(25,w)}$), were calculated with Sednterp (sednterp.unh.edu), assuming a density (ρ) of 1.005 $g \cdot mL^{-1}$ and a viscosity (η) of 1.01 mPa·s. Sedimentation profiles were analyzed using both the continuous c(s) distribution of sedimentation coefficients[6] and a non-interacting species model, yielding experimental values for s and concentration, as well as estimates of the molar mass M. Predicted values for the partial specific volume and refractive index increment ($\partial n/\partial c$)— 0.739 and 0.188 $mL \cdot g^{-1}$, respectively — were used to determine whether the proteins sedimented as monomers, dimers, or both. Figures were generated using Gnuplot.

***Estimation of the dimer dissociation constant of WT-CAN and A133P-CAN using Flow-Induced Dispersion Analysis (FIDA)***

All experiments were conducted on a FIDA Neo instrument (FIDABio) using light-emitting diode induced fluorescence with an excitation wavelength of 640 nm. We used coated capillaries (inner diameter 75 μm) and 40mM HEPES pH 7.5, 150mM NaCl buffer. All measurements were performed in triplicate with a hydrodynamic radius ($R_h$) estimated using the FIDA software on the basis of the coordinates of the CAN-functionalized *Planktothrix aghardii* OCP in the C2 space group (PDB code 7QCZ). The Protein Labelling Kit ALC 640 Fida 1 was used for cross-linking OCP and the fluorescent label (Alexa Fluor 647 (AF 647 NHS-ester; excitation and emission maxima: 655/680nm); the incubation time was 30 minutes at room temperature with a concentration of 2 mg/mL of OCP. Free Dye was eliminated with a 180 μL PD Spintrap G-25 column with an exclusion limit of 5000 Da (Cytiva, Malborough, MA, USA). Experiments were performed with a 1:1 ratio of dye to protein, 50 nM of labelled OCP (*indicator*), and increasing concentrations of unlabeled OCP (*analyte*; 0 – 350 μM) with a laminar flow created by a 400 mbar pressure difference at 25°C. The measurement was performed using the standard coated-capillary protocol, *i.e.*, the capillary was rinsed with NaOH at 3500 mbar, followed by deionized water at 3500 mbar, then filled with the analyte solution at the same pressure. The premixed indicator was injected at 50 mbar, after which mobilization and data acquisition were carried out using the analyte solution at 400 mbar. Data were processed with the FIDA analysis Software

(V3.1.1.0) with an adjustment of the viscosity via the reference residence time ($tR_{REF}$) for calculating the apparent $R_h$. The dissociation constants ($K_D$) were estimated by fitting the titration curves to the 1:1 binding stoichiometry model available of the FIDA Analysis software. The fit was very good for the data collected on the wild-type protein, but of lesser quality for the A133P mutant. Nevertheless, the data point to a 5-10 fold decrease in affinity, upon elimination of the P13/A133 H-bonds at the dimerization interface.

#### *OCP macro-crystallization by vapor diffusion*

Macrocrystals of OCP (~100-200 μm) were grown using the hanging drop geometry, as described in [1]. Crystals were grown in 24-well Linbro plates using 22 mm diameter non-siliconized cover slips (Epredia, Portsmouth, HA, USA). The precipitant solution consisted of 18% (+/- 2) polyethylene glycol (PEG) 4000 and 0.1 M sodium acetate pH 5. The protein sample was spun at 16,000 g for 10 minutes (4°C) and the supernatant was separated from the pellet to remove aggregates and then diluted to the desired concentration (200 – 300 μM). The well reservoir was filled to 1 mL with the precipitant solution, then 1 μL of protein solution and 1 μL of precipitant were sequentially pipetted onto the cover slip (no mixing) before the well was immediately sealed with grease. Crystallization setups were pipetted (and inspected) under red light and the plates were wrapped in aluminum foil for storage at 20°C. Crystals grew overnight. WT $OCP_{ECN}$ and mutant constructs crystallized using similar conditions as WT $OCP_{CAN}$, though tuning of the PEG 4000 concentration was often required to optimize crystal size.

#### *Single crystal data acquisition and processing*

Diffraction data from single crystals kept at ambient temperature were collected at the BM07 (FIP2) beamline of the European Synchrotron Radiation Facility (ESRF). Crystals of 150 – 200 μm in length were fished with a nylon cryoloop (Hampton Research, Aliso Viejo, CA, USA) and transferred to a soaking solution of 18% PEG 4000, 0.1 M Bis-Tris Propane pH 7.5, where they equilibrated for one minute before being fished out with a cryoloop and mounted on the goniometer under a stream of humid air at room temperature (90 or 97% relative humidity) using an HC-Lab Humidity Controller (Arinax, Moirans, France) or liquid nitrogen (100 K). Exposing the crystals to 100 K required the soaking solution to contain an additional 10% glycerol to mitigate ice formation. Diffraction data were collected by exposing crystals to a monochromatic X-ray beam with a wavelength of 1 Å (12.65 keV) and a size of 250 x 250 μm. Crystals were rotated at an oscillation range of 0.2° with an exposure time of 100 ms per frame, for a total of 600 – 900 diffraction images. As structures were collected over many beamtimes, the ring current was variable (ranging from 74 – 200 mA), requiring attenuation of the beam transmission to between 5.9 and 35%. All datasets were collected with doses below 0.3 MGy, following recommendations from de la Mora et al. 2020[7].

The illumination of crystals was performed using the *in crystallo* UV-vis absorption microspectrophotometer[8] available at the BM07 (FIP2) beamline, where a 532 nm LED emitting at a nominal power density of 22 mW (Thorlabs, Newton, NJ, USA) was connected to the lower 4× demagnifying objective with a 110 μm (ID) optical fiber (0.3 mW at the sample position). The crystals were either illuminated for 10 minutes, with the ~1 min data collection with continuous goniometer rotation started after 9 min of illumination, or were illuminated concomitant with data collection, resulting in 1 minute of total illumination.

Data was processed both manually and automatically. For manual data processing, the *XDS* package[9] was used. For the highest quality datasets, however, automatic data processing was used (on the *ISPyB* platform) which also relied on *XDS* and was performed using *XDSAPP* or *grenades_parallelproc*[10]. The resulting mtz files were phased with *Phaser-MR*[11] using WT OCP (PDB 7QD2 and 7QCZ for the $P2_1$ and C2 space groups, respectively) as a search model. The structures were refined using *phenix.refine* in the *Phenix* suite[12]. Real-space refinement was performed using *Coot*[13]. For all structures, iterative cycles of real space and reciprocal space refinements were performed until the model fit the 2mFo-DFc map while maintaining a reasonable geometry, no major discrepancies were present in the Fo-Fc map, and an R-free/R-work gap of 3-5%. Structure factor extrapolation was performed using *Xtrapol8*[14], where k-weighted extrapolation was performed on isomorphous datasets with a scaling factor of 0.5 to produce Fourier difference maps and kfextr extrapolated maps. A range of occupancies were sampled to determine the best occupancy estimate of the triggered-state (using the *difference-map* method), and Xtrapol8 was used to pre-refine structures in reciprocal space before they were iteratively refined using *Coot* and phenix.refine. Of important note is, however, that the occupancy estimates were only considered trustworthy when allowing to see full density around the carotenoid at a cutoff of 1 σ in the extrapolated 2mFext-DFc maps. As this was often not the case, we opted, in each dataset, for the lowest occupancy value allowing to meet that condition. The differences between dark and triggered structures are thus likely underestimated; however, we felt that it is better to not trust conformational changes in the absence of electron density. Unless stated otherwise, all graphical illustrations of the crystallographic structures were produced using PyMOL (Schrödinger, New York, USA).

***OCP microcrystallisation***

OCP crystallizes readily. However, in contrast to macro-crystallization the presence of apoprotein matters for microcrystallization as protein aggregates can result in “fishnet” or “cotton wool” like structures (see Fig. S4 in [15]) that can trap microcrystals, complicating both crystal delivery and diffraction data analysis[15]. Since crystallization is a very efficient purification method, we integrated it into the purification protocol, which allowed to omit the hydrophobic column in the purification protocol[2] (it has no negative effects on microcrystallization), in particular following the high salt concentration dialysis step mentioned for the large-scale protein purification. In the following we describe batch crystallization of OCP, in conjunction with microseeding [15], with the first crystallization step either used for protein purification and/or preparation of a seed stock solution. The volumes given are for a small-scale preparation, they can be scaled up to 100s of ml of protein. To produce seeds, 4 mL of purified protein was crystallized at an initial concentration of 4 mg/ml using the batch method with a precipitant concentration of 26% PEG 4000, 0.1 M sodium acetate pH 5. Crystals grew overnight, and were centrifuged the next day at 4,000 g for 3 minutes; the supernatant was discarded and the crystals were suspended in mother liquor (~ 100 µL of concentrated crystals in 1 mL). The microcrystal solution was then transferred to a 2 mL tube pre-filled with zirconium beads (3.0 mm diameter) (Benchmark Scientific, Sayreville, NJ, USA) and the tube was shaken in a Bead Bug microtube homogenizer (Benchmark Scientific, Sayreville, NJ, USA) for 45 s at 4000 rpm. After shaking, the tube was immediately transferred to an iced water bath for 45 s and the cycle was repeated 10 times. The crushed crystals were then transferred to a pre-filled tube containing zirconium beads of mixed sizes (0.5 and 1.0 mm diameter) and the cycle of shaking for 45 s followed by cooling for 45 s was repeated until an homogenous sample of crushed crystals under 1 µm in size was obtained. The seeds were transferred to a 1.5 ml Eppendorf tube and spun at 2,000 g for 3 minutes to sediment the crystals. The supernatant was adjusted such that ~ 100 µL of crushed crystals was matched by 400 µL of mother liquor. The seeds were then transferred back to the pre-filled bead tube and stored at -80°C. Before use for seeding crystallization setups, the thawed seeds underwent one shaking cycle in the Bead Bug.

A separate batch of freshly purified sample was then diluted to 4 mg/mL and an equal volume of 26% PEG 4000, 0.1 M sodium acetate pH 5 was prepared. Following this, 1% (v/v) of suspended seeds were added to the precipitant solution (0.5% final volume after addition of the soluble protein). The protein solution was quickly mixed with the precipitant and seeds, and the crystals were grown overnight at 20°C, wrapped in aluminum foil. The dimensions of the microcrystals were determined using a Hirox HR-5000 microscope (Hirox, Hackensack, NJ, USA). Upon crystal sedimentation, the supernatant was discarded and ~ 1 mL of fresh mother liquor was added to the sediment for the storage of the microcrystals at 4°C.

The carotenoid composition of microcrystalline $OCP_{CAN}$ used for the time-resolved serial femtosecond crystallography experiment at the Alvra-prime beamline of the SwissFEL (*vide infra*) was determined by HPLC analysis (CaroteNature.com) and shown to consist of echinone and canthaxanthine in a roughly 50:50 mixture. This observation is in line with a number of crystal structures of $OCP_{CAN}$ deposited at the PDB, showing clear difference density peaks at the β2 keto oxygen of CAN in electron density maps downloaded from the Electron Density Server.

Since the O' oxygen of canthaxanthin was always clearly visible in the electron density maps (conventional and extrapolated, alike) calculated from data collected on these crystals, the carotenoid was modelled as canthaxanthin.

#### *Cryo-electron microscopy of the A23C and A23C-A133P mutants*

*Sample preparation and image acquisition:* 3.0 µL of the constitutively dimeric A23C and A23C-A133P mutants (5 and 0.3 mg/mL, respectively) were applied to 1.2/1.3 C-Flat holey carbon grids (Protochips Inc, Morrisville, NC, USA) and plunge-frozen in liquid ethane using a Vitrobot Mark IV (Thermo Fisher Scientific, Waltham, MA, USA) with a 6 to 8 seconds blot time and a blot force of 0. These manipulations were carried out in darkness, to avoid undesired photoactivation of OCP. The sample grids were then measured at the CRG beamline CM02, operated by IBS/ISBG (Grenoble, France) and located at the ESRF (Grenoble, France). A Titan Krios G4 (Thermo Fisher Scientific, Waltham, MA, USA), operated at 300 kV, and equipped with an energy filter of 10 eV slit width (Selectris X, Thermo Fisher Scientific, Waltham, MA, USA) was used. For both mutants, movies were recorded automatically using EPU (Thermo Fisher Scientific, Waltham, MA, USA) with a Falcon 4i detector (Thermo Fisher Scientific, Waltham, MA, USA). For each movie, the total exposure time was 3.16 s, corresponding to 54 frames at ~0.9 $e^{-}/Å^{2}$ (*i.e.,* a total dose of ~60 $e^{-}/Å^{2}$ per movie). The magnification was 215,000×, corresponding to a nominal pixel size of 0.57 Å at the camera level. The defocus range for the images was between −1.0 and −2.2 µm. For the A23C mutant, a total of 8,168 movies were collected while for the A23C-A133P mutant, a total of 17 584 and 12 096 movies were collected untilted and with a 40° tilt respectively.

*Image analysis of the A23C mutant :* The movies were imported into CryoSPARC[16] for processing. Patch motion correction and patch CTF estimation were performed. Blob picking followed by 2D classification was conducted to identify 2D classes representing distinct particle views, which were then used for template-based particle picking. Particles were extracted with a box size of 128 pixels (1.8 Å/pixel sampling). A 20 Å resolution rendering of the crystal structure of an OCP dimer was used as a 3D reference for a heterogeneous refinement with four classes. The best 3D class was selected for further processing. An *ab initio* reconstruction was performed followed by a heterogeneous refinement (both requesting three 3D classes). Particles from the best 3D class were again selected for another round of *ab initio* reconstruction and heterogeneous refinement (again with three 3D classes). This process led to the isolation of 230,709 particles, which were re-extracted with a box size of 286 pixels (1.0 Å/pixel sampling). A sequence of processing steps—initial non-uniform refinement, local and global CTF refinement, and a second non-uniform refinement—yielded a 3D map at an overall resolution of 2.8 Å (FSC = 0.143).

*Image analysis of the A23C-A133P mutant :* For the untilted dataset, image processing was performed with RELION 5.0[17]. The movies were first drift-corrected with MOTIONCOR2[18], then CTF estimation was carried out with CTFFIND4[19] within RELION. Micrographs with a defocus of less than -3 µm and a CTF resolution worse than 5 Å were discarded. Particles were first picked with a Laplacian-of-Gaussian approach and classified in 2D in 2 successive rounds. Only the best looking 2D classes corresponding to monomers or dimers of OCP A23C – A133P were kept to train and pick new particles with TOPAZ[20]. 2 512 197 particles were extracted and 2D classified in 2 successive rounds. The best 2D classes corresponding to monomers and dimers were kept and a 3D classification (in 10 classes, C2 symmetry applied, circular mask diameter of 120 Å) was performed using a 20 Å resolution rendering of the crystal structure of the dimer formed by OCP A23C – A133P as an initial 3D model. The particles belonging to the best dimer 3D class (101 230 particles) were kept. Many attempts in getting a 3D map were made but were unsuccessful because of a problem of preferred orientation. The particles from the untilted data identified with RELION were further processed in CRYOSPARC[16] in combination with tilted data (see below).

For the tilted dataset, the movies were first drift-corrected with MOTIONCOR2[18] in RELION. The micrographs were then imported in CRYOSPARC for further processing. Patch CTF was performed and micrographs with an average defocus between -1 and -3 µm and a CTF resolution fit better than 10 Å were kept. The best particles identified in the untilted data (see above) were then imported in CRYOSPARC and a 2D classification was performed. The 2D classes representing unique views were selected to perform template picking on the tilted micrographs. Following two rounds of 2D classifications, the best 2D classes from the tilted data were selected for second template picking hoping to improve the quality of the picking. After two successive 2D classifications, a non-uniform refinement (with 77 873 tilted particles) gave a first 3D map at 4.2 Å resolution using a 20 Å resolution rendering of the crystal structure of an OCP dimer as an initial 3D model.

The untilted (101 230 particles) and tilted (77 873 particles) data were then pooled together in CRYOSPARC, for another non uniform refinement that gave a 3D map at 3.3 Å resolution. Hetero-refinement (3 classes, C2 symmetry) further identified one main class with 100 860 particles. The final 3D map of the OCP A23C–A133P dimer was obtained at an average resolution of 3.1 Å (FSC = 0.143) following a workflow of non-uniform refinement, global CTF refinement, another non-uniform refinement, local CTF refinement, and a final non-uniform refinement. This last map was subsequently sharpened with DeepEMhancer for manual model fitting[21].

***Refinement of the cryo-electron microscopy structures of the A23C and A23C-A133P mutants***

For both mutants, refinement started with the real-space fitting of the $P2_1^A$, $P2_1^B$, and C2 chains in the raw cryo-EM maps using Chimera[22]. Upon finding that these are best fit by the $P2_1^B$ model, we used iterative cycles of *phenix.real_space_refine*[12] and *Coot*[13] to refine the structures. Figures were produced using Chimera X[23]. Statistics of the final structures are shown in Supplementary table 2.

***Comparison of visible–near infrared (NIR) femtosecond time-resolved absorption spectroscopy on solution and crystalline OCP***

We performed ultrafast spectroscopy on both crystal forms and compared the results with those obtained from solution samples. Both crystal forms were investigated, with the $P2_1$ crystal form suspended in PEG and the C2 form (C2c conformer) embedded in LCP.

*Setup description:* Femtosecond Vis-NIR transient absorption spectra were collected using a *Helios* system (*Ultrafast Systems*). It consists of an 80 MHz titanium sapphire oscillator (*Mai Tai* (*Spectra Physics*)) generating ≈70 fs pulses followed by a high-energy titanium sapphire regenerative amplifier (*Spitfire Ace* (*Spectra Physics*)) generating ≈100 fs pulses with a 1 kHz repetition rate. The resulting 800 nm beam was split into two beams to generate: (1) the pump (532 nm) using the optical parametric amplifier *Topas Prime* coupled with a *Nir-UVVis* frequency mixer and (2) the probe, a white light continuum in the vis-NIR range generated by focusing the fundamental beam into a sapphire (430–780 nm), or YAG (820–1390 nm) crystal. The residual 800 nm photons were filtered out. The pump beam was depolarized to avoid anisotropy effects. The sample-containing cuvette was translated using a XY translational stage generating a pseudo Lissajous pattern. The sample and experimental setup was kept at 22°C. OCP microcrystals (~ 3 µm) in PEG (17% PEG, 0.1 M natrium acetate pH 5.0) were measured in cuvette with 2 mm optical path length, while microcrystals (~ 3 µm) embedded in LCP (see section on serial femtosecond crystallography at Alvara (below) for details) were dispensed in a 100 µm cuvette. OCP solutions in 2 mm cells were stirred during measurements. For each delay, a transient absorption spectrum was obtained by averaging 1000 spectra with and without excitation, respectively. The measurement of the entire set of pump-probe time delay points was repeated four times, then the data were inspected and averaged. The transient absorption data were corrected for the chirp of the white light continuum taking into account the given thickness of sapphire, water and BK7 glass that the probe pulse had to transverse before reaching the sample. Data were projected onto a 5 nm spaced grid to get smoother kinetic traces. For each sample and energy, the experiment was performed twice, once with probing at NIR (820–1390 nm) and once with probing at visible wavelengths (430–780 nm). Both datasets were merged later.

*Pump pulse characterization:* The pump pulse duration was determined using the stimulated Raman signal clearly visible in the data obtained from OCP microcrystals in PEG using a 3.2 µJ pulse, peaking at 645 nm. This signal stems from the instantaneous pump-probe interaction with the solvent and represents cross-correlation signal between pump and probe. It is followed by the excited state absorption (ESA) signal originating from the OCP. The 660 nm kinetics contains almost no stimulated

Raman signal, but a very similar the ESA contribution as for the 645 nm kinetics (since the stimulated Raman signal is spectrally narrow and ESA is spectrally broad). We separated the stimulated Raman signal by subtracting the kinetics at 660 nm (multiplied by factor of 0.78) from the kinetics at 645 nm (Supplementary fig. S42A). The resultant stimulated Raman signal free of ESA contributions (blue curve, fig. S42A) was fitted with a Gaussian curve (Supplementary fig. S42B), yielding a FWHM of 149 fs, representing the instrument response function (IRF). The pump and probe pulse duration was therefore 105 fs (149 fs/ $\sqrt{2}$), assuming that both pump and probe have Gaussian temporal profiles and equal durations, and other factors contributing to the IRF are negligible. The pump spatial profile at the sample position was determined using a horizontal razor edge scan. The resultant curve was fitted by the integrated Gaussian curve (Supplementary fig. S42C), yielding 246 μm FWHM of the Gaussian curve. The energy and power density profiles were calculated using these pump parameters (Supplementary fig. S42D, E). We performed power dependence measurements using a pump laser energy range of 0.2–3.2 μJ. Time-resolved spectra to determine the photodynamics of the various samples in the linear excitation range were obtained using 0.8 μJ laser energy, corresponding to 1.17 mJ/cm$^2$ peak energy density and 10.44 GW/cm$^2$ peak power density.

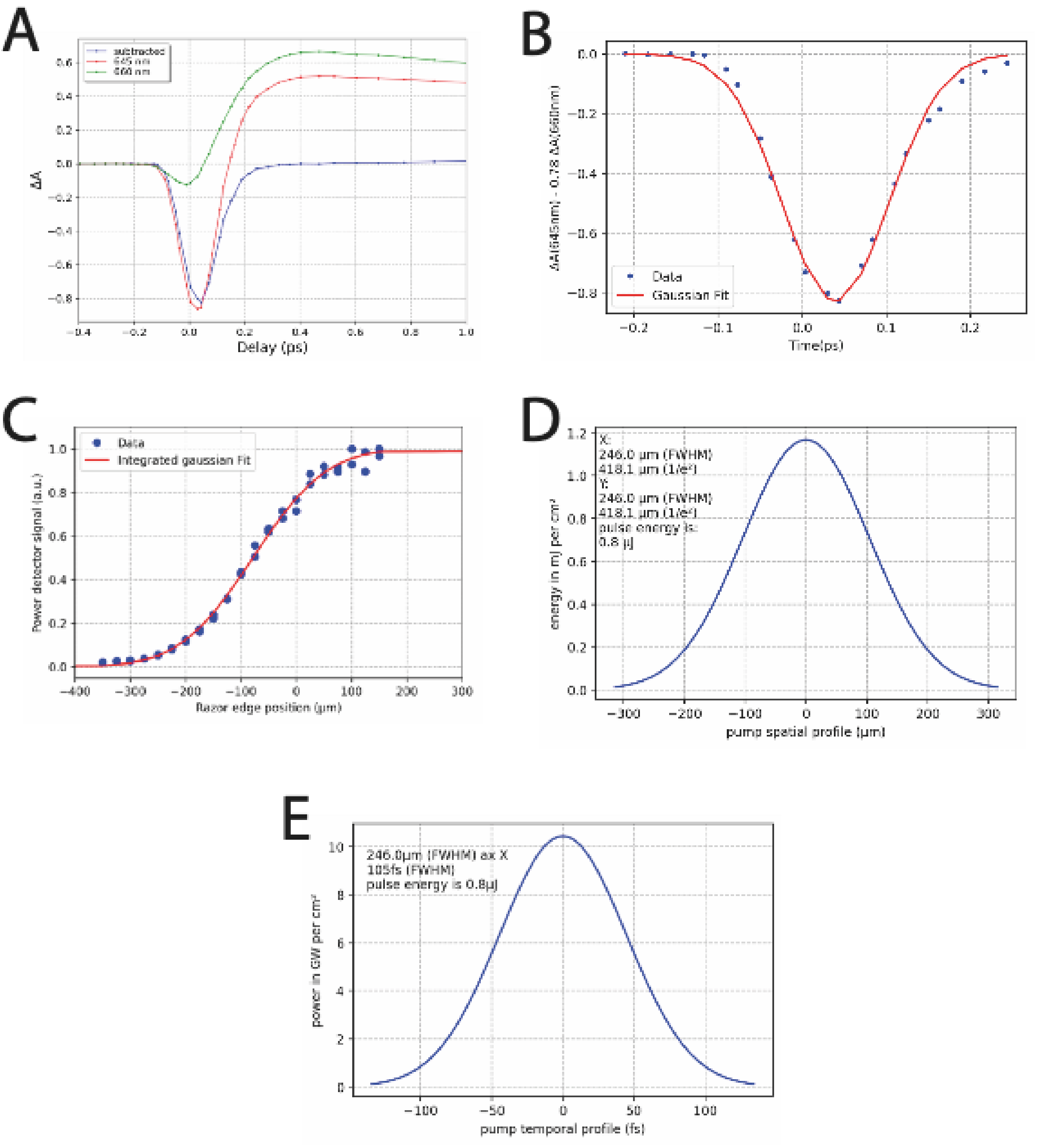


**Supplementary fig. 42**: A) Raman signal extracted from the OCP signal by subtraction of 645 nm kinetic. B) The resulting Raman kinetics was fitted with a Gaussian function, resulting in a 149 fs (FWHM) instrument response function (IRF). C) The pump spatial profile was determined by a horizontal razor edge scan of the pump beam in the sample position, resulting in 246 µm (FWHM). D, E) The energy density and power density profiles of the 532 nm pump pulse were calculated using the experimentally derived (see B,C) pump beam parameters The peak values are 1.17 mJ/cm$^2$ (D) and 10.44 GW/cm$^2$ (E) for 0.8 µJ pump laser energy.

***Serial femtosecond crystallography at the Alvra prime instrument at SwissFEL***

OCP microcrystals with a size of ~ 3 µm were embedded in a lipidic cubic phase (LCP) matrix for high viscosity extrusion (HVE) delivery.  LCP was prepared by mixing 60 µl of monoolein (42 °C) and 35-40 µl of a solution consisting of 76 mM Bis Tris Propane pH 7.5, 18 % (w/v) pluronic F-108 (Sigma) and 35 % (w/v) PEG 3350 using coupled Hamilton® Gastight® syringes until the mixture turned transparent. Before embedding in the LCP viscous matrix, the OCP microcrystal suspensions were pelleted by gentle centrifugation. 12 µl of a rather dry crystalline pellet were mixed homogeneously with 95-100 µl of the transparent LCP matrix using coupled Hamilton® Gastight® syringes. For SFX data collection the contents of three of such syringes was mixed homogeneously, pooled and filled into a custom-built, large volume HVE injector (Doak *et al* in preparation). For HVE delivery, the microcrystal containing LCP was extruded from capillaries with 75 µm inner diameter at a flow rate of ~4.2 µl/min. The extruded jet was stabilized in the Alvra prime sample chamber (filled with 100 mbar He atmosphere) using a He gas sheath At this flow rate, the spacing between consecutive X-rays shots (delivered at 100 Hz) was ~90 µm and 180 µm between consecutive optical laser shots (delivered at 50 Hz see below), avoiding light contamination. The actual spacing on the jet between two consecutive X-ray pulses was monitored in real time during SFX experiments, based on the analysis of the light transmission change imprinted by X-rays in the viscous material. Runs during which the jet speed varied significantly between the shots or became too slow, were discarded from the analysis. SFX data were collected in October 2020 using attenuated XFEL pulses (40 fs in length, nominal photon energy 11840 eV (FWHM 20 eV), a photon flux of ~1.7 × $10^{11}$ photons/pulse, pulse energy  ~330 µJ) focused to 2.7 µm (h) × 2.2 µm (v) (FWHM). Data were recorded using a JUNGFRAU 16 M pixel detector operating in 4M mode. The outer panels were excluded to reduce the amount of data.

The SFX experiments were carried out in a time-resolved mode using an optical pump – X-ray probe scheme. As detailed above, transient absorption spectroscopy power titrations carried out on dilute OCP solutions as well as on OCP microcrystals (either suspended in mother liquor ($P2_1$ crystal form, colloidal sample) or embedded in LCP (C2 crystals, C2c conformation) show that the $S_2$ yield (1100 nm band) deviates significantly from linearity above 20 GW/$cm^2$, yielding a long-lived, off-pathway, radical-state[24]. This finding guided the choice of the pump laser parameters of the TR-SFX experiment.

Briefly, the Alvra experimental laser system (~20 mJ, central wavelength 800 nm, 35 fs FWHM pulses at 100Hz) pumped an OPA to deliver 535 +/- 10 nm pulses which were made circularly polarised by an achromatic quarter wave plate. This beam was focused at the interaction region to a Gaussian spot of size 43 x 43 $µm^2$ FWHM (73 µm $1/e^2$), at an angle of ~3 degrees with respect to the X-rays, and with an estimated pulse duration of ~85 fs FWHM. The pump laser fluence was 1.5 mJ/$cm^2$, (pulse energy 32 nJ), corresponding to nominally 0.4 absorbed photons on average per chromophore (power density: 18 GW/$cm^2$). The synchronization between laser and XFEL pulses was monitored and corrected by an online diagnostic of their arrival time (timing tool) and by regularly determining and correcting the nominal time t=0.

Diffraction data with and without pump-laser excitation were collected in an interleaved fashion with the XFEL operating at 100 Hz and the pump laser at 50 Hz. Online monitoring of the hit-rate (*i.e.* the average percentage of images containing crystal diffraction) was carried out using the beamline analysis tools. A series of TR-SFX experiments were carried out at pump-probe delays of 0.3, 1.5, 10 and 100 ps, corresponding to the peak occupancy of the ICT, $S_1$, S* and $P_1$ states.

***Serial femtosecond crystallography at the Cristallina-MX instrument at SwissFEL***

SFX data were collected in December 2023 as part of the commissioning of the Cristallina-MX instrument, including the setup of HVE injection at Cristallina-MX and pump–probe experiments using an HVE injector. OCP microcrystals with an average size of 5 x 3 x 3 µm were embedded in a lipidic cubic phase matrix as described above for the experiments performed at Alvra. The HVE injection was performed and monitored as described above for the Alvra experiment, however this time the sample was extruded in air at atmospheric pressure and a rotating catcher[25] was used to stabilize the extruded jet. X-ray pulses with a nominal photon energy of 11.2 keV (FWHM 20 eV) and a duration of 25 fs (FWHM) were used, with a pulse energy of ~160 µJ at the sample position, accounting for losses and attenuation, corresponding to $\sim 9.17 \times 10^{10}$ photons/pulse. The beam was focused to 3.6 µm (h) × 2.8 µm (v) (FWHM). Data were recorded using a JUNGFRAU 8 M pixel detector with sixteen sensor modules. The pump laser (EKSPLA NT230, 535 ± 10 nm, 3.7 ns pulse length (FWHM)) was fiber coupled (70 m, 100 µm core, 0.12 NA (Ceram Optec)) to the sample position such that the optical beam was nearly collinear (4° deviation) to the X-rays and focused to a spot size of 32 x 35 $\mu m^2$ ($1/e^2$) at sample position. Diffraction data with and without pump-laser excitation were collected in an interleaved manner with the XFEL operating at 100 Hz and the pump laser at 50 Hz. Online monitoring of the hit-rate (*i.e.,* the average percentage of images containing crystal diffraction) was carried out using the beamline analysis tools. A series of TR-SFX experiments were carried out at pump-probe delays of 10 ns and 1 µs, with a pump-laser fluence of 17 $mJ/cm^2$, corresponding to nominally 5 absorbed photons on average per chromophore (power density: 0.005 $GW/cm^2$). Online monitoring of diffraction data, such as determination of hit-rate and estimation of the fraction of multiple hits, was carried out using the data processing pipeline using the *CrystFEL*[26] engine.

***Processing of the Serial femtosecond crystallography (SFX) data collected at the Alvra Prime instrument at SwissFEL***

SFX data were collected either in complete dark (*dark-only*) or according to a pump-probe scheme with dark (no laser; *dark-interleaved*) and light (laser excitation; *light*) images interleaved. A total of ~1.3 M indexed patterns were collected. After automatic hit-finding by a dedicated beamline tool, the data were split into *light* and *dark* images, which were then separately indexed, integrated and merged using *CrystFEL* 0.8.0[26]. A dataset assembled from the first 20,000 *dark-only* patterns indexed in the C2 space group served to refine the detector geometry. We then switched to the automatic indexing pipeline available at Alvra, ~95-99% of the data collected during this beamtime indexed in the C2 space group. Data were reprocessed offline (*i.e.,* after the experiment) using crystFEL 0.10.2, with the crystal to detector distance refined on a per-run basis. Best parameters for the indexing were grid-searched, which maximized the number of patterns indexed in $P2_1$ and C2. Comparable data (same space group and illumination conditions) were merged using the MonteCarlo approach, where the intensity estimate for each unique reflection is obtained by averaging of redundant measurements. Neither polarization correction nor rescaling by a second pass was applied to the data, after it was found that these approaches resulted in degraded figures of merit. Final data statistics are shown in Supplementary Table 6.

As noted above, data were acquired either in complete dark (*dark-only*) or using a pump–probe sequence in which dark (no-laser; *dark-interleaved*) and light (laser-excited; *light*) images were alternated. In all cases, the jet velocity was ~9 mm/s, providing a spacing of approximately 90 µm between successive shots to ensure that no light contamination occurred. This assumption was verified by comparing the *dark-interleaved* dataset assembled from all *dark-interleaved* indexed patterns with the *dark-only* dataset assembled from *dark-only* patterns collected without interleaving light patterns. Specifically, we checked for structural differences by means of *dark-interleaved* vs. *dark-only* Fourier difference maps and extrapolated structures, using the program Xtrapol8. Even with the highly redundant C2 data, we found structural differences to be negligible, amounting to 5% occupancy for a state with a virtually identical structure (rmsd=0.275 Å) to that solved with the merged data, for both the protein and carotenoid. Accordingly, nearly identical extrapolated maps and structures were obtained when comparing the *light* data to either sets of *dark* data or their combination thereof. Therefore, we decided to opt for high redundancy, and all *dark* data were merged together (*all-darks*). Note that given the scarcity of $P2_1$ crystals in the sample used during this first experiment, the decision to merge dark-only and dark-interleaved (all-dark) data was derived from examination of the C2 dark data, and then applied to the $P2_1$ data as well. All refinements in the reciprocal-space were carried out using *phenix.refine*, from the Phenix suite of crystallographic software[12]. Likewise, all real-space refinements were performed using *Coot*[13]. As for the other structures reported herein, geometry restraints used for refinement of the canthaxanthin molecule were downloaded from the Grade2 web server (https://grade.globalphasing.org/cgi-bin/grade2_server.cgi). To ensure that the geometry of canthaxanthin did not deviate from ideal values, notably in terms of bond lengths, it was re-fit using *Coot* in the 2mFextr-DFc electron density map with very strong weight on the Lennard-Jones parameters (0.1, with an overall X-ray/geometry weight of 5), after each refinement step, including the last one. The resulting structures are admittedly the most conservative that can be fit in the observed electron density, and hardly make room for the observation of features such as bond elongation. Nevertheless, we found this stringency to be the better option to avoid unjustified claims rooted in an overfitting of the data.

As for the *dark-interleaved* vs. *dark-only* datasets, structural changes between the *dark* datasets and the *light* datasets collected at various time delays and energies were assessed by means of Fourier difference maps and extrapolated structures, using the program Xtrapol8. We used q-weighting of structural differences to produce the Fourier difference maps, and k-weighting to generate extrapolated structure factors. This was justified by the higher completeness of the k-weighted extrapolated data, which translated to higher quality extrapolated maps as well as better statistical indicators for the refined extrapolated structures. Likewise, the Fourier difference maps and extrapolated maps obtained by comparing the *light* datasets to the *all-dark* dataset were of higher quality than those obtained using the *dark-only* or *dark-interleaved* datasets. This was most visible for the 0.3 ps data, where the peaks heights on the carotenoid are highest, and thereafter applied to the other datasets of the data series. Thus, our analysis was based on the *light* vs. *all-darks* Fourier difference maps and extrapolated structure factors, respectively. It needs be acknowledged, however, that due to the very low optical laser fluence, the Fourier difference maps did not feature strong peaks. Nevertheless, conformational changes were clearly visible in the extrapolated maps.

Occupancy determination is central in scalar structure factor amplitude extrapolations. In first approximation, we used the estimate from the *difference-map* method, found to be most robust in our tests[14]. With the $P2_1$ data, occupancy estimates were high and varied from ~30-40% at 0.3 ps to 15-30% at 1.5, 10 and 100 ps. Occupancy values derived from the *Pearson-CC* method were of the same magnitude or higher. We suspected these estimates to be highly biased by the comparatively small number of indexed lattices that was used to generate the *light* (and *dark*) datasets. Indeed, the highest occupancy is found for the dataset composed of the lowest number of frames (0.3 ps). Using the *distance-refinement* method as well as our new SVD method (see section SSX data analysis), we found the occupancies to be likely lower (**Materials and Method Table 1**), confirming this hypothesis. Nevertheless, we used, in the end, the occupancy estimate given by the *difference-map* method for all datasets as extrapolated electron density maps degraded at lower occupancies – notably in terms of definition of the carotenoid in the electron density. The refined extrapolated structures are thus presumably very conservative; however, we decided not trust conformational changes in case where electron density did not contour the whole carotenoid at 1 RMSD in the extrapolated 2MFextr-DFc maps.

**Materials and Method Table 1**

| **Occupancy estimates of the phototriggered states in the $P2_1$ data collected at the Alvra prime instrument** | | | | | | | |
|---|---|---|---|---|---|---|---|
| **Extrapolated dataset** | Difference-map | Pearson CC | Distance refinement | SVD, $2^{nd}$ lSV | SVD, $3^{rd}$ lSV | Average estimate excluding the Pearson CC | **Applied occupancy based on the carotenoid electron density as the sole criterion** |
| **Light$^{0.3ps}$** | 40 | 40 | 32.6 | 20 | 15 | 26.9 | **40** |
| **Light$^{1.5ps}$** | 20 | 30 | 25.3 | 15 | 8 | 17.1 | **20** |
| **Light$^{10ps}$** | 20 | 35 | 20.2 | 15 | 15 | 17.6 | **20** |
| **Light$^{100ps}$** | 20 | 35 | 20.8 | 10 | 6 | 14.2 | **20** |

The C2 data were of much higher quality, presumably due to significantly higher multiplicity of observations. Indeed, the *all-dark* reference dataset used for structure-factor extrapolation comprised 1.32 million patterns, while the *light* datasets ranged from ~85,000 patterns (0.3 ps, 45 GW/cm²) to ~165,000 patterns (100 ps). Regardless of the probed time delay, occupancy estimates obtained with the *difference-map* method were consistently similar (6–7%), prompting us once again to examine alternative approaches for determining occupancies (**Materials and Method Table 2**). The *distance-refinement* and SVD methods yielded occupancy estimates similar to those found with the difference map approach, viz ~6-9 and ~9%, respectively, while the *PearsonCC* method again yielded larger estimates, viz.10-15%. Based on the confirmation that occupancy in the C2 data series does not vary as a function of the time delay, we refined the intermediate state structures using the occupancies determined by the difference-map method, viz. 7%.

**Materials and Method Table 2**

| **Occupancy estimates of the phototriggered states in the C2 data collected at the Alvra prime instrument (%)** | | | | | | |
|---|---|---|---|---|---|---|
| **Extrapolated dataset** | **Difference-map** | Pearson CC | Distance refinement | SVD, $2^{nd}$ lSV | SVD, $3^{rd}$ lSV | Average estimate including (or excluding) the Pearson CC |
| **Light$^{0.3ps, 18GW/cm2}$** | **7** | 15 | 6.7 | 10 | 9 | 9.5 (8.2) |
| **Light$^{1.5ps, 18GW/cm2}$** | **7** | 10 | 6.4 | 9 | 9 | 8.3 (7.9) |
| **Light$^{10ps, 18GW/cm2}$** | **7** | 15 | 5.8 | 10 | 9 | 9.4 (8.0) |
| **Light$^{100ps, 18GW/cm2}$** | **7** | 10 | 6 | 9 | 9 | 8.2 (7.8) |

***Processing of the Serial femtosecond crystallography (SFX) data collected at the Cristallina-MX instrument at SwissFEL***

Data were recorded on a Jungfrau 8M detector (Dectris, Villigen, Switzerland) either in complete dark (*dark-only*) or according to a pump-probe scheme with dark (no laser; *dark-interleaved*) and light (laser excitation) images interleaved, amounting to a total of 915,951 indexed patterns. After automatic hit-finding by a dedicated beamline analysis tool (see above), the data were split into *light* and *dark* images, which were then separately indexed, integrated and merged using crystFEL v 0.10.2. A dataset assembled from 20,000 indexed *dark-only* patterns in the C2 space group served to refine the detector geometry. We then switched to the automatic indexing pipeline provided by the facility (using a CrystFEL[26] engine) to index diffraction patterns in both the $P2_1$ and C2 space groups, respectively. During this experiment, roughly 50 % of crystals belonged to the $P2_1$ space group, with the remainder corresponding to C2 crystals where the C2c and the C2o conformers were populated at ~50%. The crystal to detector distance was refined on a per injector-filling basis and data were reprocessed offline (*i.e.,* after the experiment). Comparable data (same space group and illumination conditions) were merged using the MonteCarlo approach, where the intensity estimate for each unique reflection is obtained by averaging of redundant measurements. Neither polarization correction nor rescaling by a second pass was applied to the data, after it was found that these approaches resulted in degraded figures of merit. Of note, the jet was found to be extremely stable during this experiment, possibly due to a new version of the catcher[25] that was used during this beamtime. Nevertheless, data collection runs during which jet issues occurred were systematically discarded. Final data statistics are shown in Supplementary Table 7.

Pump-probe data were collected with *dark* (no laser) and *light* (laser excitation) images interleaved. Again, we had adjusted the injection conditions such that the jet advanced by ~90 µm/180 µm between consecutive X-ray/pump laser shots to prevent that the dark data were contaminated by stray illumination. It was found using structure factor extrapolation methods that the *dark-interleaved* and *dark-only* data these were not strictly equivalent, most likely due to differing injection conditions, preventing the merging of the *dark-only* and *dark-interleaved* data. Thus, for both crystal forms and time-delays, only the *dark-interleaved* data collected alongside the *light* data were used as references for intermediate-state structure extrapolation. Fortunately, this drawback was compensated by the high fraction of $P2_1$ crystals (50% of all indexed patterns) which together afforded all our *dark-interleaved* and *light* datasets to be assembled from 100-k indexed patterns or more. Using an average of all occupancy estimates the triggered-state occupancy was found to amount to ~3-4 % for the C2 data (**Materials and Methods Table 3**) and 6-8 % for the $P2_1$ data **(Materials and Methods Table 4)**. In the end, we used a 3.5% occupancy for the C2 data, and a 7% occupancy for the $P2_1$ data ().

The C2 crystals were found to feature the C2o and C2c states at ~50% occupancy in both the *dark-interleaved* and the *dark-only* data. Nevertheless, in the C2 *light* vs. *dark* extrapolated maps, electron density is observed only for the C2c conformation. This observation implies either that the C2o→C2c transition occurs within 10 ns or that any light-induced structural changes in the C2o form have decayed by 10 ns.

**Materials and Methods Table 3**

| Occupancy estimates of the phototriggered states in the C2 data collected at the Cristallina-MX instrument | | | | | | | |
|---|---|---|---|---|---|---|---|
| **Extrapolated dataset** | **Difference-map** | Pearson CC | Distance refinement | SVD, 2$^{nd}$ lSV | SVD, 3$^{rd}$ lSV | Average estimate including (or excluding) the Pearson CC | Occupancies used |
| **Light$^{10ns}$** | **2** | 3.5 | 3.0 | 4 | 2.5 | 3.0 (2.9) | 3.0 |
| **Light$^{1µs}$** | **3.5** | 7 | 3.0 | 4 | 3 | 4.1 (3.3) | 3.5 |

### Materials and Methods Table 4

| Occupancy estimates of the phototriggered states in the $P2_1$ data collected at the Cristallina-MX instrument | | | | | | | |
|---|---|---|---|---|---|---|---|
| **Extrapolated dataset** | **Difference-map** | Pearson CC | Distance refinement | SVD, 2$^{nd}$ lSV | SVD, 3$^{rd}$ lSV | Average estimate including (or excluding) the Pearson CC | Occupancies used |
| **Light$^{10ns}$** | **7** | 10 | n.a. | 4 | 3 | 6.0 (4.7) | 7 |
| **Light$^{1µs}$** | **10** | 15 | 3.0 | 4 | 3 | 8.0 (5.0) | 7 |

#### *Refinement of the extrapolated SFX structures*

All refinements in reciprocal-space were carried using *phenix.refine* from the Phenix suite of crystallographic software[12]. Likewise, all real-space refinements were performed using *Coot*[13]. The geometry restraints used for refinement of the canthaxanthin molecule were downloaded from the Grade2 web server (https://grade.globalphasing.org/cgi-bin/grade2_server.cgi). To ensure that the geometry of canthaxanthin did not deviate from ideal values, notably in terms of bond lengths, it was re-fit in real space with very strong weight on the Lennard-Jones parameter (0.1, with an overall X-ray/geometry weight of 5). Thus, our structures are strongly constraint with conservative estimates for structural changes that could be fit in the observed electron density. Nevertheless, we found this stringency to be the better option to avoid unjustified claims rooted in an overfitting of the data.

#### *Hydrogen bond analysis of TR-SFX structures*

The hydrogens were added using Reduce and the environment of  histidines were checked using MolProbity[27]. Hydrogen bonds (H-bonds) were detected using the HBPLUS algorithm[28]. H-bonds occur between donor (D) and acceptor (A) atoms that satisfy the following geometric criteria: (i) maximum distances of 3.20 Å for D-A and 2.70 Å for H-A, (ii) minimum value of 90° for D-H-A, H-A-AA and D-A-AA angles, where AA is the acceptor antecedent. We then counted the total number of H-bonds for each structure. The analysis was performed by comparing: i) the total number of H-bonds in each structure, ii) total number of H-bonds that were removed, iii) total number of H-bonds that were added, and iv) total number of H-bonds that remained the same. Such H-bond inventories were made for the dark state structure as well as the structures determined from the time-resolved data series and then compared.

***Pathway analysis using ComPASS***

The ComPASS tool was used to perform communication network analysis[29]. Starting from an ensemble of conformations of a given system, ComPASS extracts several properties: generalized correlations, non-covalent interactions, communication propensity (variance of inter-residue distances), and inter-residue distances. It then applies principal component analysis (PCA) on the generalized correlations, non-covalent interactions, and communication propensity to construct an adjacency matrix. A graph is then built, in which nodes represent amino acids, and edges are defined by applying a distance cutoff to the average minimum distance matrix. The adjacency matrix is then used to assign weights to the graph. Finally, ComPASS identifies the following features from the graph: *i)* the shortest pathways between all pairs of residues, *ii)* hotspots, determined using standard centrality measures, *iii)* cliques, defined as fully connected groups of residues, and *iv)* communities, corresponding to highly interconnected groups of residues that may share a common functional role.

For each set of structures (C2c, $P2_1^A$, $P2_1^B$), we used Reduce to add hydrogens and checked the environment of the histidines using MolProbity. Then we modelled the gaps (2-3 missing residues at the N-terminus, and 10-16 missing residues within the linker) using MODELLER[30,31] and applied ComPASS to extract the network of communication pathways. Finally, we report the set of top 10% most significant shortest pathways for each ensemble. The ligand was excluded from the analysis, as the method supports only natural amino acids and nucleic acids as input.

***Prolonged blue light illumination of OCP microcrystals and mounting on SOS chips***

Half a milliliter of gravity-sedimented OCP microcrystals (~7 µm) was deposited in a single-isolated well of 6 well Greiner plate (well diameter: 35 mm). An equivalent volume of 17% PEG 4000, 0.1 M NaCl, 0.1 M sodium acetate pH 5.8 was added to the crystals and the crystal/reagent suspension was agitated at room temperature to prevent crystals sedimentation. An ice pack was scotch-taped under the plate to prevent crystal heating under illumination. The ice pack was replaced every 6-8 hours. The plate and the ice pack were together placed on a slowly rotating orbital shaker, and the well containing the crystals was illuminated for 48h using a gooseneck light guide of 4.5 mm diameter (P/N: 154 202; Schott, Germany) to deliver the ~450 nm light produced by a SCHOTT KL1500 halogen lamp (P/N: 150 700; Schott, Germany) equipped with a blue filter (maximum transmission at ~450–500 nm, with cut-on and cut-off wavelengths of 420 and 520 nm, respectively). The distance from the light guide to the Greiner plate was adjusted (~10 cm) so that the well containing the crystals remained constantly and uniformly illuminated while shaking. Before data collection, ~50 µl of the crystal suspension were transferred to an Eppendorf tube and centrifuged for 10 seconds using a benchtop centrifuge. 2 to 5 µL of the crystal pellet were pipetted and deposited per SOS chip[32,33], consisting of two 2.5 µm thick Mylar sheets. After placing a Mylar sheet on the lower part of the chip holder the OCP microcrystals were deposited and spread using a pipette tip, before being covered by the upper Mylar membrane; then the SOS chip was closed. It was attached via a magnetic base to the beamline goniometer. The same treatment was applied to crystals kept in the dark, but similarly transferred into a Greiner plate placed over an icepack and agitated for 48 hours. Before mounting, the absorption spectra of dark and illuminated crystals were recorded using a NanoDrop™ One (Ozyme, Saint-Cyr-l'École, France) spectrophotometer. Background subtraction was applied to extract the absorption peaks.

***Serial synchrotron crystallography (SSX) at the microfocus ID23EH2 beamline***

SSX data were collected at the ID23-EH2 beamline of the European Synchrotron Radiation Facility (ESRF) using a fly-scan approach, where the SOS chips is raster-scanned across the X-ray beam by vertical and horizontal translations of the MD3 High Precision X-ray Microdiffractometer (Arinax, Moirans, France), with the shutter continuously open during vertical translations. A Pilatus 2M detector (Dectris, Villigen, Switzerland) was used to collect the diffraction data. Raster-scanning steps and exposure times were 6 µm and 6 ms at a transmission of 100%, respectively, data collection per chip took 90 minutes. Three and two chips were collected for the dark and light datasets, respectively. All chips were prepared fresh, *i.e.* just before data collection. For collection of the light datasets, the crystals in the SOS chip remained illuminated with a 440 nm LED during data acquisition. The photon energy was 14200 eV, corresponding to 0.873 Å wavelength. The average diffraction weighted dose was 26 kGy per crystal.

***Processing of the SSX data***

Data processing was performed using NanoPeakCell[34] for hit finding and CrystFEL v. 0.8.2[26] for the indexing, integration and merging of the diffraction data. For the latter step, we generally used the MonteCarlo approach, where the intensity estimate for each unique reflection is obtained by simple-averaging of multiple measurements, and in these case, neither polarization correction nor rescaling by a second pass was applied. Only the C2newlight-48h dataset (see below) was merged using the scaling approach in partialator (model: *unity*; iterations: *5*) after it was found that better statistics could be obtained than with the MonteCarlo approach (this was not the case for other datasets). From three SOS chips[32,33] filled with crystals continuously kept in the dark, we collected 141902 patterns, containing 103914 hits of which 9134 and 33366 were indexed in $P2_1$ and C2 space groups, respectively. These datasets are thereafter referred to as $P2_1^{dark48h}$ and $C2^{dark48h}$, respectively. We also collected data from two continuously light-exposed chips filled with crystals that had been kept under ~450 nm illumination for 48h. 114949 patterns were collected, containing 68929 hits of which 38464 were indexed. Specifically, 13953 and 14220 patterns indexed in the $P2_1$ and C2 space group, using as search unit cell parameters those of the $P2_1^{dark48h}$ and $C2^{dark48h}$ datasets, respectively. These datasets, which are isomorphous to their dark counterparts, are thereafter referred to as $P2_1^{light48h}$ and $C2^{light48h}$, respectively. A new population of C2 crystals was also found, characterized by a 16% increase in unit cell volume. The dataset merged from these crystals 10291 crystals is thereafter referred to as the $C2new^{light48h}$ dataset.

***Refinement of the SSX structures***

Reciprocal-space refinement of the SSX structures was carried out using *phenix.refine* from the Phenix suite of crystallographic software[12]. Likewise, all real-space refinements were performed using *Coot*[13]. The geometry restraints used for refinement of the canthaxanthin pigment were downloaded from the Grade2 web server (https://grade.globalphasing.org/cgi-bin/grade2_server.cgi). To ensure that the geometry of the canthaxanthin did not derive from ideal values, notably in terms of bond lengths, it was refit in real space with very strong weight on the Lennard-Jones parameter (0.1, with an overall X-ray/geometry weight of 5) to prevent overfitting of the data.

Dark state structures in the $P2_1$ and C2 space groups ($P2_1^{dark48h}$ and $C2^{dark48h}$) were solved by rigid-body refinement using as the search model the deposited dark-state $P2_1$ (wwPDB id: 7QD2) and C2 (7QCZ) structures obtained via conventional cryo-crystallography at 100 K, respectively[1]. After real-space fitting in the electron density of the residues displaying different conformations at RT and 100 K, the structures were refined in reciprocal space. For residues present in alternate conformations, occupancies were as well refined. Iterative cycles of refinement in reciprocal space and real space were carried out to obtain the models of which the statistics are shown in Supplementary table 5.

The $P2_1^{light48h}$ and $C2c^{light48h}$ datasets were isomorphous to the $P2_1^{dark}$ and $C2c^{dark}$ datasets, respectively. Structures solved from these datasets did not readily show differences with respect to the dark-state structures. Therefore, extrapolations methods were used to obtain insights into the putative conformational changes occurring upon photoexcitation. We used Xtrapol8[14] to calculate extrapolated structure factors amplitudes (ESFA) at various occupancies of the phototriggered-state, with application of (i) the k-weighting approach to down-weight poorly measured reflections ($Fext=\alpha*m^{dark}*k*(Fobs^{light}-Fobs^{dark}) + Fobs^{dark}$, with $\alpha=1/occupancy$) and (ii) the truncate method to "rescue" negative reflections that would otherwise not be usable by reciprocal-space refinement programs. Use of k-weighting was preferred over the q-weighting scheme because of its lower stringency that resulted in more complete datasets (less rejection of reflections), which in turn afforded better refinement statistics and extrapolated electron density maps (2mFext-DFc) of higher quality. Nevertheless, we stuck to the more stringent q-weighting approach for the calculation of Fourier difference maps ($m^{dark}*q*(Fobs^{light}-Fobs^{dark})* \exp(i\varphi^{dark})$) as no obvious improvement was seen in the quality of these maps upon switching between the k- and q-weighting schemes. Notwithstanding, the Fourier difference maps did not feature strong peaks. The occupancy-estimate for the triggered state in the $P2_1$ crystals significantly differed depending on whether the difference-map method (20 and 44 % for the C2 and $P2_1$ datasets) or the PearsonCC-method (15 and 28% for the C2 and P21 datasets) was used. Recall that the former method assigns as the correct occupancy that at which the height of peaks is maximized in the initial mFext-DFc difference maps, while the latter picks that at which the location and height of peaks in the initial extrapolated difference map $((m^{dark}Fext\text{-}DFc)*\exp(i\varphi^{dark}))$ compare best with the Fourier difference map. Distance-refinement[14] did not help, due to its stringency in the selection of the tracked interatomic distances (n.d. for the C2 and $P2_1$ datasets). Thus, we sought to obtain an additional occupancy-estimate using an orthogonal approach. Given the richness in information of extrapolated electron density maps (2mFext-DFc), compared to both Fourier difference map and initial extrapolated difference maps, we developed a method based on (2mFext-DFc) maps. This new occupancy-determination approach is based on singular value decomposition (SVD[35]) of the refined 2mFext-DFc extrapolated maps, i.e. those obtained after reciprocal space refinement of the dark-state structure against the various sets of extrapolated structure factor amplitudes (ESFA). When applying SVD, the left-singular vectors (lSV) represent the separated states and the right-singular vectors their evolution across the input datasets, while the singular values themselves (S) point to the overall weight of each state in the data series. Specific to our SVD-based occupancy-determination approach, *i.e.* applied to a dataset composed of extrapolated electron density maps calculated for various presumed occupancies of the triggered state, the first lSV is an average of all the input maps while the second lSV yields a map similar to that of the dark-state and thus features its minimal rSV value at an occupancy value closed to that of the triggered state. By contrast, the third lSV informs on the triggered state, and thus features its maximal rSV at the occupancy where the latter is best defined in terms of electron density. In practice, for a given pair of reference and triggered datasets, the second and third lSV thus offer upper (conservative estimate) and lower (permissive estimate) bounds for the range of trustworthy occupancy values. For both the C2 and $P2_1$ datasets, we used the permissive estimates (20 and 44 % for the C2 and $P2_1$ datasets) rather than the conservative estimates (30 and 52 % for the C2 and $P2_1$ datasets) on the basis that these were closer to the estimates derived by the difference-map method and Pearson-CC methods. Following occupancy determinations, refinement of the extrapolated $P2_1^{light\text{-}48h}$ and $C2c^{light\text{-}48h}$ structures was straightforward, with iterative cycles of real-space fitting and reciprocal-space refinement. A summary of the occupancy determination efforts is shown in **Material and Methods Table 5**. Refinement statistics are shown in Supplementary table 5.

**Material and Methods Table 5**

| Occupancy estimates of the phototriggered states in the SSX data collected at the ID23EH2 beamline (%) | | | | | | |
|---|---|---|---|---|---|---|
| Extrapolated dataset | Difference-map | Pearson CC | Distance refinement | SVD, $2^{nd}$ lSV | SVD, $3^{nd}$ lSV | Average estimate including (or excluding) the Pearson CC |
| $C2^{light48h}$ | 15 | 15 | n.a. | 30 | 25 | 23 (21.25) |
| $P21^{light48h}$ | 28 | 28 | n.a | 52 | 44 | 47 (42) |

The C2new$^{light-48h}$ structure was solved by molecular replacement with Phaser[11]. For this approach to be successful (as judged from the quality of maps and $R_{free}/R_{work}$ values after the first refinement round), we had to use as search models the coordinates of the isolated NTD and CTD, respectively. Indeed, rigid-body refinement was unable to place the two domains of the proteins, even when these were labelled as separated chains in the input model and rigid-body options sets to fit each chain individually. In retrospect, this failure is not surprising, given the large change in conformation undergone by the protein upon reopening of SC#1 in presence of the untethered carotenoid. After obtaining the first phased electron density map, multiple cycles of rebuilding in the electron density and reciprocal-space refinement were carried out. The resulting model could not be improved further, and while the $R_{free}$ and $R_{work}$ value remain high, they likely relate to the comparatively large distribution of unit cell parameters that characterized the C2new crystals as and the correspondingly high Wilson-B of the merged data. Refinement statistics are shown in Supplementary table 5.

***Evaluation of the precision of atomic coordinates in our time-resolved SFX structures.***

All coordinate errors reported in the manuscript (Figure 5) were estimated using the bootstrap method. Briefly, the bootstrap resampling approach can be used in serial crystallography to estimate the precision of atomic coordinates when individual observations are varied[36]. In this context, individual observations correspond to indexed diffraction patterns, which serve as a pool from which resampled datasets are constructed. In the bootstrap method, a given indexed pattern can be drawn multiple times (random sampling with replacement), and each resampled dataset contains the same number of indexed patterns as the original dataset. All SFX datasets (dark and light) were resampled 100 times to compute, for each time delay and space group, 100 resampled sets of extrapolated structure factors computed with the appropriate α value. Subsequently, 100 refinements were performed with *phenix.refine* for each intermediate state using the corresponding refined model as the starting model. All atomic coordinates were perturbed by 0.3 Å using the *sites.shake* option in *phenix.refine*. All coordinate errors reported in the manuscript (Figure 5) were estimated using the bootstrap method. For the distances between two atoms, errors were computed using the RMSD of the 100 distances obtained from the 100 refined models.

***Molecular dynamics***

All molecular dynamics simulations were performed using NAMD 3.0[37] at 300 K and 1 atm. The pressure and temperature were maintained using a Langevin barostat[38] and Langevin dynamics, respectively. Hydrogen mass repartitioning[39] and the r-RESPA multiple time-step integration algorithm[40] were employed to integrate the equations of motion with time steps of 4 fs for short-range and 8 fs for long-range interactions. Covalent bonds involving hydrogen atoms were constrained using the SHAKE/RATTLE[41] and SETTLE[42] algorithms. Long-range electrostatic interactions were treated using the particle-mesh Ewald method[43]. A 9-Å cutoff was applied to truncate both van der Waals and short-range Coulombic interactions. After 2,000 minimization steps, all systems were equilibrated for 5 ns with harmonic restraints on all protein and canthaxanthin heavy atoms, followed by an additional 20 ns with restraints kept only on the protein backbone atoms. The systems were further thermalized for 100 ns with soft harmonic restraints applied to the backbone atoms of the αGH loop (residues 120–129). For production purposes, ten and five independent 1-μs trajectories were generated for WT OPC in the monomeric C2o and C2c and the dimeric C2o, C2c, and $P2_1^A$/$P2_1^B$ conformations, respectively.

#### ***Well-tempered metadynamics simulations***

*Construction of the systems*: The initial systems were built using the C2o, C2c, $P2_1^A$ and $P2_1^B$ X-ray structures in complex with canthaxanthin. Missing residues (2-3 missing residues at the N-terminus, and 10-16 missing residues within the linker) were added using Modeller[31]. The Q79L mutants were generated for C2o and C2c by manually mutating the glutamine residue to leucine. The proteins were then embedded in ca. 90 × 90 × 90 Å$^3$ water boxes, an Na+ and Cl- ions were added at a concentration of 0.15 M to neutralize the total charge of the system using the *tleap* module of the AMBER 2024 suite[44]. Water molecules were modeled using the TIP3P model[45]. The ff14SB AMBER force field[46] was used to describe the proteins and ions, and canthaxanthin was parameterized using the modified GAFF[47] parameters derived by Bondanza et al.[48]. Multiple-walker well-tempered metadynamics extended adaptive biasing force (MW-WTM-eABF) simulations were then run to examine the $P2_1^B$ to C2o and C2o to C2c transitions, respectively.

*Enhanced-sampling simulations :* The free-energy change underlying the transition between the $P2_1^B$ and the C2o conformations was determined with the well-tempered metadynamics extended adaptive biasing force (WTM-eABF) algorithm[49,50], using, as a reaction coordinate model, a path collective variable[51], *s*, along a surrogate pathway connecting the end states of the transition, built from five collective variables, namely (i) the distances between I51(CD1) and CTX(C40), between V53(HA) and W279(O), and between T52 (H) and W279(O), and the RMSDs with respect to $P2_1^B$ and to C2o. The free-energy change underlying the transition between the C2o and the C2c conformations of the Q79L mutant was determined with the well-tempered metadynamics extended adaptive biasing force (WTM-eABF) algorithm[49,50], using, as a reaction coordinate model, a path collective variable[51], *s*, along a surrogate pathway connecting the end states of the transition, built from three collective variables, namely the distances between P126(CB) and CTX(CO6), between D35(OD1/2) and Y129(OH), and between I125(CG2) and Y129(OH). In each case, the distances were chosen on the basis of the amplitude of their difference in the compared structures. In a nutshell, the WTM-eABF algorithm relies upon the integration of the average force acting on $s$, obtained from unconstrained molecular dynamics simulations. In the course of the simulation, a biasing force is estimated such that, once applied to the system, it yields a Hamiltonian devoid of an average force exerted along $s$. As a result, all values of $s$ are sampled with an equal probability, which, in turn, greatly improves the accuracy of the calculated free energies. The transition pathway spanned the [0; 1] interval, and was discretized in bins 0.02 wide, where samples of the local force acting along $s$ were accrued. To mitigate the risk of deleterious nonequilibrium effects, no time-dependent bias was applied until a threshold of 10,000 samples was reached[52]. In consideration of the complexity of the transition, likely to be related to slowly relaxing degrees of freedom coupled to $s$, likely to hamper convergence of the free-energy calculation, a multiple-walker strategy with four walkers was employed to improve ergodic sampling[53]. The statistical error associated to the free-energy change was estimated from the variance of the free-energy differences measured by each walker.
