## Supplementary texts for "Structural basis of the two-photon photoactivation mechanism of orange carotenoid protein"

for

by

Rory Munro, Elena A. Andreeva[#], Elisabeth Hartmann[#], Quentin Goor[◊], Hosni El Zein[◊], Stanislaw Nizinski, Adjélé Wilson, Elke De Zitter, Gregory Effantin, Nicolas Coquelle, Ninon Zala, Martin V. Appleby, Shira Bar-Zvi, Camila Bacellar, Emma Beale, Emmanuelle Bignon, Bernhard Brutscher, Martin Byrdin, Claudio Cirelli, Florian Dworkowski, Lutz Foucar, Guillaume Gotthard, Alexander Gorel, Marie Luise Grünbein, Mario Hilpert, Philip J.M. Johnson, Marco Kloos, Gregor Knopp, Karol Nass, Gabriela Nass Kovacs, Dmitry Ozerov, Christopher J. Milne, Gotard Burdzinski, Christophe Chipot, Yasaman Karami, François Dehez, Martin Weik, R. Bruce Doak, Robert L. Shoeman, Giorgio Schiro, Michel Sliwa, Diana Kirilovsky, Ilme Schlichting[*] and Jacques-Philippe Colletier[*]

[#] these authors contributed equally.
[◊] these authors contributed equally.
[*] correspondance to

**Authors affiliations:**

[1]**Institut de Biologie Structurale, Grenoble, F-38000 France**:
Rory Munro, Elena A. Andreeva, Quentin Goor, Hosni El Zein, Elke De Zitter, Nicolas Coquelle, Gregory Effantin, Ninon Zala, Bernhard Brutscher, Martin Byrdin, Martin Weik, Giorgio Schiro, Jacques-Philippe Colletier

[2]**Max Planck Institute for Medical Research, D-69120 Heidelberg, Germany**:
Elisabeth Hartmann, Elena A. Andreeva, Marco Kloos, Marie Luise Grünbein, Stanislaw Nizinski, Alexander Gorel, Lutz Foucar, Mario Hilpert, Gabriela Nass Kovacs, R. Bruce Doak, Robert L. Shoeman, Ilme Schlichting

[3]**Institute for Integrative Biology of the Cell, Gif-sur-Yvette, France**:
Adjélé Wilson, Shira Bar-Zvi, Diana Kirilovsky

[4] **SwissFEL, SLS, Paul Scherrer Institute, Villigen, Switzerland**:
Dmitry Ozerov, Karol Nass, Philip J.M. Johnson, Emma Beale, Claudio Cirelli, Gregor Knopp, Camila Bacellar, Christopher J. Milne, Martin V. Appleby, Guillaume Gotthard, Florian Dworkowski

[5] **Université de Lorraine, CNRS, LPCT, F-54000 Nancy, France**:
Christophe Chipot, Emmanuelle Bignon, François Dehez

[6] **European XFEL, Hamburg, Germany**:
Marco Kloos, Christopher J. Milne

[7] **Quantum Electronics Laboratory, Faculty of Physics and Astronomy, Adam Mickiewicz University, Uniwersytetu Poznańskiego 2, Poznan 61-614, Poland**
Stanislaw Nizinski, Gotard Burdzinski

[8] **Theoretical and Computational Biophysics Group, Beckman Institute, and Department of Physics, University of Illinois at Urbana-Champaign, Urbana, Illinois 61801, USA;** [9] **Department of Biochemistry and Molecular Biology, The University of Chicago, Chicago, Illinois 60637, USA:**
Christophe Chipot

[9] **Université de Lorraine, CNRS, Inria, LORIA, F-54000 Nancy, France**:
Yasaman Karami

[10] **Laboratoire d'Optique et Biosciences, CNRS, Inserm, Ecole polytechnique, Institut Polytechnique de Paris, 91120 Palaiseau, France**:
Michel Sliwa

**Supplementary text 1: Dark state structures of OCP**

Recently, six cryo-crystallographic (100 K) structures of dark-adapted *Planktothrix aghardii* OCP were determined[1] using two different monoclinic crystal forms (C2 and $P2_1$) showing three distinct conformational sub-states regardless of the functionalizing carotenoid (echinenone or canthaxanthin). In the $P2_1$ crystals, a biological dimer is present in the asymmetric unit, consisting of conformationally distinct chains that are hereafter referred to as $P2_1^A$ and $P2_1^B$. In the C2 crystals, a single monomer is present in the asymmetric unit, from which a perfectly-symmetric biological dimer can be derived by crystallographic operations, referred to as C2 in the following. This conformational diversity originates at a specific crystal packing interface (coined interface X in [1]), consisting of the αCD and αGH loops in the $P2_1$ crystals (Supplementary fig. S1). In the C2 crystals, also αD, αE and linker residues participate, resulting in a 50% increase in buried surface area at this interface as well as in a very compact OCP conformation. While the CTD is overall well conserved structurally in the $P2_1^A$, $P2_1^B$ and C2 chains, this is not the case for the NTD, which comprises two four-helix bundles (αBCHI and αDEFG)[3] that face one another across the carotenoid tunnel and are connected by the αCD and αGH loops (Fig. 1). We noted previously[1] that the $P2_1^A$, $P2_1^B$ and C2 conformations diverge mostly in the αC (conserved segment is A33-L45) and αD (conserved segment is A63-Q73) helices and the interconnecting αCD (K49-Q62) loop (Supplementary fig. S2). Because structural equilibria can shift during crystal cryo-cooling – potentially obscuring the relationship between αCD loop conformational dynamics and function – we here describe structures determined from diffraction data collected at room-temperature (RT) in the dark as well as under various illumination conditions, using macrocrystals kept at various relative humidities or microcrystals in a variety of conditions. We mainly focused on cantaxanthin (CAN)-functionalized OCP, but structures were also determined of echinenone (ECN)-functionalized protein, when needed to test our structural and functional hypotheses.

From RT structures determined in the two space groups at 97% relative humidity (Supplementary tables 1 and 3), we confirmed our previous hypothesis[1,2] that structural dynamics in the αCD loop impacts interaction of the carotenoid within the tunnel, access of bulk-solvent to it (Fig. 2A), inter-helix and inter-bundle distances in the NTD (similar in $P2_1^B$ and C2, but different in $P2_1^A$), as well as the coupling of distinct structural elements (αC to αG, via the Y44/E115 H-bond ; αCD loop to β5-β6 turn, via the T52/P278 and T52/W279 H-bonds ; αCD loop to linker, via the A58/V178 H-bond ; αH to αI, via the L145/Q150 H-bond; and αH to αN, via the Q149/A309 H-bond) (Supplementary fig. S2). These differences are described in the following for the three conformers.

Briefly, in the $P2_1^B$ structure (Supplementary fig. S3), the αCD loop is stretched as an extended coil between the two straight αC (A33-K49) and αD (F63-Q73) helices, with stabilization of the loop structure by a single H-bond to the β5-β6 turn (A54(N)/W279(O)) as well as hydrophobic interactions (L56 fits into a groove formed by P278 and W279, while A55, A54 and V53 arrange around Y111)

(Supplementary fig. S2). The straight helical conformation of αC, reminiscent of that seen in the *Synechocystis* PCC6803 (Syn)-OCP structure (3MG1)[4], is in many structures stabilized by an intra-helix H-bond between E46(OD2) and K49(NZ) (though often broken in pH 7.5 structures), while that of αD seems to only result from the absence of constraining interactions with other structural elements (Supplementary fig. S3). The $P2_1^B$ conformation of the αCD loop "plugs" V53 into the carotenoid tunnel, placing its CG1 atom ~4.4 Å from the carotenoid C13' methyl group, while the I51 and M47 side chains together fit into a groove surrounded by the side chains of S147, I151, L154, Q150, F280 and V284 (Supplementary fig. S2). This conformation of the protein, where T52 is exposed to the solvent, prevents direct contact between the M47 and I51 side chains and the carotenoid (closest interatomic distances: 6.4 and 8.5 Å). The position of M47(SD) impacts nearby Q150, at the αHI turn, which displays two conformers, both of which H-bond to I142(O). At the C-terminus of the αCD loop, N60 and Q62 participate in forming a groove at the interface with αG (T102, I105) and β4 (E248) within which linker residues E176 and P177 fit. Specifically, the E176 carboxylate oxygens fit between Q248(NE2) and N104(ND2), the latter forming a conserved interdomain H-bond with W279. There is, however, in the $P2_1^B$ chain, no direct coupling between the αCD loop and the linker region (Supplementary fig. S2).

These observations contrast sharply with those in the $P2_1^A$ and C2 chains which share three distinguishing features: (i) a kink at E46, preventing the intra-helix H-bond to K49, resulting in the refolding of M47-T50 as a $3_{10}$ helix (Supplementary figs. S1 and S3). This conformational change positions the I51 side chain in the carotenoid tunnel, packed between the aromatic rings of Y44, Y111, W279 and F280, replacing V53 in $P2_1^B$. (ii) A similar coiling of the T52-A55 segment, stabilized by two inter-domain H-bonds from T52(OG1) and (N) to P278(O) (2.9 Å in the C2 and $P2_1^A$ chains) and W279(O) (2.9 and 2.8 Å in the C2 and $P2_1^A$ chains), respectively; and (iii) an identical H-bonding pattern for Q150 at the αHI turn, *i.e.*, with its NE2 atom H-bonded to both I142(O) and L145(O) (Supplementary figs. S2 and S41).

Despite these similarities, the $P2_1^A$ and C2 chains also show significant differences, all of which originate from the structurally-divergent L56-Q62 segment (Supplementary fig. S2). Notably in the latter, A58(O) draws near the linker residue V178(N) (3.3 and 7.8 Å in the C2 and $P2_1^A$ chains, respectively), inducing rearrangements at the αCD/β5β6 hydrophobic interface. These changes ultimately bring αD ~1.3 Å closer to both αG—with a water bridge connecting E65 and Q112—and αC, and are accompanied by a rotation of the αDEFG bundle toward αC. Despite the fact that in both chains A55 faces W279 (3.5 Å in the C2 and $P2_1^A$ chains) and L56 fits into a hydrophobic furrow created by V53(CG1), M61(CG), G108(CA), Y111(CD1) and W279(CB) (Supplementary figs. S2, S3 and S41), the "pull" exerted by the A58(O)/V178(N) H-bond results in a "looser" fit in the C2 chain (Supplementary fig. S2), allowing the aromatic ring of Y111 to nudge between the L56 and I51 side chains. This induces a ~180° change in I51($\chi$1), which in turn displaces Y44 towards W41, effectively closing SC#2 (Fig. 2A, B, and Supplementary fig. S2). This makes room for M47 to reorient between the Y44 and F280 rings such that its side chain also interacts with the carotenoid (the CE atom is at 5.5 Å from the carotenoid C12'). Indeed, the CD1 atom of I51 remains in the C2 chain at a similar distance to the carotenoid C11' (4.1 Å), despite the rotamer change associated with the above-described structural reorganisation. It is of note that two conformers are observed for Y44 in the C2 chain, both of which close off SC#2 (Fig. 2A, B, and Supplementary fig. S2). One of the two conformers is locked in place by an H-bond to one of two E115 conformers, in effect coupling the two four-helix bundles through a "tether" between the αC and αG helices. The alternate E115 conformer H-bonds to Q119 – the terminal residue of αG. Water-bridges complement the two modes of interaction, with each of the E115 conformers connecting via a water molecule to the non-H-bonded conformers of Q119 and Y44. It must be stressed that in the $P2_1$ chains as well as in the *Synechocystis* PCC6803 (Syn)-OCP structure (3MG1)[4], E115 and both Y44 ($P2_1^A$ ; $P2_1^B$, *Syn*-OCP) and Q119 ($P2_1^B$, *Syn*-OCP) interact via a water molecule. All in all, the above-described changes at αC and the αCD loop decrease the distances from αC to αD, αE and αG on average by ~1.5 Å in the C2 chain, compared to either of the $P2_1$ chains. No such changes are observed at the second point of linkage between the two bundles, *i.e.* the αGH loop, notwithstanding the stronger attachment of the αGH loop onto αC, through a (second) D35/S132 H-bond (OD1 to N, in addition to OD2 to OG1).

*Space group and structural transitions*

We found that the $P2_1$ crystals can transition to C2 upon change in relative humidity (97 % to 90 %), precipitant concentration, high phosphate concentration (not shown) as well as embedding in viscous matrices such as lipidic cubic phase (LCP) or hydroxy-ethyl-cellulose (HEC, not shown). This transition is favored by an increase of pH from 5.0 (crystallization) to ≥ 5.8 (we typically used pH 7.5, to match conditions in which photoactivity and recovery are measured) as well as (*i*) a sudden (but limited) increase in the PEG concentration (up to 100 % enrichment); or, (ii) a sudden decrease in relative humidity (RH; 50-100 % enrichment). The latter was found to be achievable through ~10-seconds of air-drying of $P2_1$ crystals in the mounting loop, or by directly mounting $P2_1$ crystals in a humidifier stream at RH ≤ 97%. In the case of micro-crystals, the space group transition occurred upon transfer into a LCP (see TR-SFX experiments) or HEC (15 %) matrix (not shown). In general, the $P2_1$ to C2 transition results in an improvement of the diffraction quality, all the more upon embedment in LCP (Supplementary table 1, 3, 5 and 9 vs. Supplementary tables 7 and 8). By contrast, HEC embedded microcrystals show reduced stability, with first a progressive change in the C2 unit cell parameters and then loss of diffraction power, on the ~1.5 to 2-hour timescale.

The above-outlined structural similarities between the $P2_1^A$ and C2 chains strongly suggest that the $P2_1^A$ state is an obligatory intermediate along the $P2_1$ to C2 transition (Fig. 2A, B, and Supplementary fig. S6A, B), *i.e.* the $P2_1^B$ chains must first transition to the $P2_1^A$ conformation, restructuring the tunnel wall, before progression to the C2 arrangement can occur, whereby the two chains become related by crystallographic symmetry. The C2 dimerization interface features the strictly-conserved D19$^{A/B}$(OE1)-R27$^{B/A}$(NH2) salt-bridge and two H-bonds between T17(N) /(O) and N134(ND1)/(OD1), respectively, but lacks the intermolecular P13$^{A/B}$(O)/A133$^{B/A}$(N) H-bond, also strictly conserved across the OCP1 clade (Fig. 1E). Absence of this H-bond releases the αGH loop (Fig. 2C). Importantly, the optical spectrum of C2 crystals at RH ≥ 97% or preserved in their mother liquor at RT resembles that of OCP$^{O}$ in solution and of the P2$_1$ crystals, with two vibrionic peaks at ~495 and 505 nm, respectively (Supplementary fig. S21D). Thus, the C2 conformation at RH ≥ 97% can be considered as one of the several OCP$^{O}$ states[5].

*Conformational transitions in the C2 space group*

The humidity-driven conformational changes at interface X[1] not only permit conversion of P2$_1$ crystals to C2 but can also drive a coil-to-3$^{10}$ helix refolding of the highly-conserved 122-VAPIPSGY-129 segment of the αGH loop (G120-A133) (Supplementary figs. S1, S6C, S6D, and S7). This structural change closes the carotenoid tunnel at solvent channel #1 (SC#1)[1], resulting in complete insulation of the carotenoid from bulk solvent (Fig. 2A, B). Briefly, at RH ≥ 97%, the carotenoid is accessible to the bulk *via* the open SC#1 in both the P2$_1^{A,B}$ and C2 structures. The coiled, 'SC#1-open' conformation of the αGH loop is stabilized by the strictly conserved H-bonds to αE (at the N-terminus) and αC (at the C-terminus), viz. Q79(OE1)—A123(N), D35(OD2)—Y129(OH) and D35(OD1)—S132(OG) H-bonds, respectively. Upon decreasing the RH to ≤ 90%, or when the crystals are dehydrated by one of the means described above, an alternate conformer accumulates in the C2 crystals, where the αGH loop folds into a 3$_{10}$ helix, effectively closing SC#1 (Supplementary figs. S6 and S7). We refer to this conformer, which cannot be accessed in the P2$_1$ crystals and features both SC#2 and SC#1 in the closed conformation as C2c (Supplementary fig. S6C) – in contrast to C2o where only SC#2 is closed (Fig. 2A, B). The defining features of the C2c conformer are rupture of the strictly conserved Q79(OE1)—A123(N) and D35(OD1)—Y129(OH) H-bonds, upon folding of residues 122-VAPIPSGY-129 into a 3$_{10}$ helix. This conformational change switches I125 and P126 as the residue laying atop the W41 side chain, in effect "plugging" P126 into the carotenoid tunnel (CG is 4.2 Å from C3') (Supplementary fig. S7). It is accompanied by a 180° change in Y129($\chi$1) which places its phenol ring in the exit path of SC#1. The table below, which is also provided as Supplementary table 2, summarizes the differences between the C2c and C2o structures.

**Synopsis of the defining features of the C2o and C2c structures:**

| Type of interaction | Involved residues | Distance in C2o (Å) | Distance in C2c (Å) |
|---|---|---|---|

| | | | |
|---|---|---|---|
| H-bonds between the αGH loop and neighboring structural elements (αE, αC) | Q79(OD1)—A123(N) | 2.7 | **4.0** |
| | D35(OD1)—Y129(OH) | 3.2 | **9.7** |
| | D35(OD2)—S132(OG1) | 2.6 | 3.0 |
| | D35(OD1)—S132(N) | 3.1 | 3.0 |
| Van der Waals interactions between the αGH loop and neighboring structural elements (αE, αC) | W41(CD1)—I125(CB) | 3.9 | **8.6** |
| | W41(CB)—I125(CG1) | 3.6 | **9.5** |
| | W41(CD1)—P126(CB) | **8.6** | 4.5 |
| | W41(CB)—P126(CB) | **9.7** | 3.6 |
| | T80(CG2)—Y129(OH) | **9.6** | 3.3 |
| H-bonds stabilizing the $3_{10}$ helix | I125(O)—G128(N) | **6.3** | 3.3 |
| | P124(O)—S127(N) | **6.3** | 3.2 |
| | P124(O)—S127(OG1) | **8.4** | 2.6 |
| Closest van der Waals interactions between the αGH loop and canthaxanthin. | I125(CG1)—CAN(C3′) | 5.2 | **8.4** |
| | P126(CG)—CAN(C3′) | **11.0** | 4.2 |

Besides these differences, the C2c structure shares all features of the C2o structure, including (i) the tunnel wall rearrangements bringing I51(CD1) and M47(CE) in van der Waals contact with canthaxanthin (or echinenone); (ii) the closure of SC#2 upon decrease of the W41 and Y44 distance and (optional) formation of the Y44/E115 H-bond, linking αC and αG; and (iii), the broken $P13^{A/B}$(O)/$A133^{B/A}$(N) intermolecular H-bond at the dimerization interface (Figs. 1E and 2C). Importantly, we never observed a closed state in $P2_1$ crystals, irrespective of the conditions in which they were probed (LCP embedding, solid supports, mother liquor at various RH) (Supplementary figs. S6C and S24). Thus, the three conformational changes characteristic of the $P2_1$ to C2o transition – (i) kink at αC; (ii) refolding of the αCD loop which tightens carotenoid stabilization, closes SC#2, and establishes new inter-domain contacts; (iii) rupture of the P13/A133 H-bond which increases αGH loop flexibility) are obligatory steps for the coil-to-$3_{10}$-helix αGH loop conformational switch required for forming the C2c state (Fig. 2B). We note that the C2c conformational substate was first observed in the structure of *Gloeobacter* OCP-X (PDB code 8A0H), where it was proposed to cause the reduced activity of the protein[6]. Importantly, the C2c conformation is incompatible with the crystal lattice contacts of Q130 observed in both crystal forms ($P3_221$, containing a monomer, a dimer is obtained via crystallographic symmetry operations and $P3_2$, containing an asymmetric dimer, both similar to the C2/$P2_1$ situation) obtained of OCP from *Synechocystis sp. PCC 6803* (for example, wwPDB entries 5TUX and 3MG1, respectively).

In macrocrystals, the C2c state can be accumulated to 100% by either directly mounting C2 crystals at 70 % RH or by gradually decreasing the RH to 60%. Both approaches, however, generally degrade diffraction quality. More plastic and prone to afford the C2o to C2c conformational transition without impact on diffraction quality are microcrystals (typically 3-8 µm), in which C2c can be accumulated to 100%. Accordingly, our first structural characterisation of the C2c state was achieved by serial femtosecond crystallography at the Coherent X-ray Imaging instrument of the Linear Coherent Light Source (CXI-LCLS), where ~7 µm microcrystals were injected into the X-ray beam by high-viscosity extrusion (HVE), after embedding the crystals into an LCP matrix prepared at pH 5.8 – the minimum pH value to obtain transparent LCP. All crystals were found to belong to the C2 space group, and the 1.4 Å resolution structure solved from ~60,000 indexed patterns showed 100% occupancy of the C2c conformer. In follow-up experiments, we found that the C2c conformer can also be elicited in microcrystals embedded in HEC or mounted in MISP[7] or SOS[8,9] chips.

*Equilibrium structural dynamics of dark-adapted CAN-functionalized OCP*

CryoEM analysis identifies the $P2_1^B$ conformer, characterized by open SC#1 and SC#2 and an extended and straight αC helix, as that most representative of $OCP^O$ is solution and in the dark (Supplementary figs. S4 and S5). X-ray structures additionally suggests that $P2_1^A$ and C2o are obligate intermediates along the $P2_1^B$ → C2c transition ($P2_1^B$ → $P2_1^A$ → C2o → C2c). It remains unclear, however, whether absorption of the first blue-green photon—delivering roughly 55–60 kcal.mol$^{-1}$— drives the $P2_1^B$ → ($P2_1^A$ →) C2o step, the C2o→C2c step, or both. We examined the energetics of these transitions by applying computational methods to CAN-functionalized OCP. Free-energy calculations for the $P2_1^B$→C2o transition yield a barrier of 5.5 kcal·mol$^{-1}$ and a stabilization of $P2_1^B$ relative to C2o by approximately 3.5 kcal.mol$^{-1}$, consistent with µs-timescale molecular dynamics (MD) simulations, which at equilibrium show no spontaneous conversion of $P2_1^B$ to C2o (Supplementary fig. S16A and Materials and Methods).

Attempts to map the free-energy landscape of the C2o→C2c transition proved more challenging. The C2o conformation was found to be highly stable on the µs-timescale. Contrastingly, the C2c conformation displayed transient C2o-like features such as reformation of the D35(OD1)–Y129(N) or the Q79(OD1)–A123(N) H-bond, as early as within the first ns of the C2c simulations (Supplementary figs. S16B, S17, and Materials and Methods). The latter H-bond was generally found to then only rarely break. A complete C2c to C2o transition was observed in only one out of the ten 1µs-long simulations simulation started from the C2c state (Supplementary figs. S16B, S17, and Materials and Methods), suggesting a high-barrier between the two conformers. Free energy calculations pointed to a similar stabilization of the C2o and C2c conformers, in agreement of both being present in CAN-OCP solutions at RT, but determining the barrier between them was beyond the reach of the methods applied for the wild-type protein. Resorting to calculations on the Q79L mutant (see Main text and

Supplementary texts 3, 4), experimentally shown to recover $OCP^{O}$ faster from $OCP^{R}$ than WT, resulted in a more stable C2c conformer, unable to transition to the C2o state within the duration of our MD simulations (multiple trajectories, each spanning ~1 μs), though a free-energy barrier of a 1.8 kcal·mol$^{-1}$ could be estimated for the C2o→C2c transition (Supplementary figs. S16B, S16C and S18). These findings suggest that rupture of the Q79(OD1)–A123(N) H-bond is the main obstacle to the C2o→C2c transition, and that the latter features a higher barrier than the $P2_1^{B}$ → C2o transition in the WT protein. In conclusion, molecular dynamics and metadynamics simulations suggest that in the OCP-CAN conformational energy landscape, the $P2_1^{B}$, $P2_1^{A}$, C2o (SC#1-open) and C2c (SC#1-closed) conformations are distinct persistent conformers present at RT. While the crystallographic WT-C2c conformation is stable in crystals, we see in simulations a propensity of the Q79—A123 H-bond to reform (Supplementary S17), which would dramatically elevate the barrier between the C2o and C2c state and explain presence of the latter, at equilibrium, in protein solution (Supplementary figs. S17 and S18). Absence of the Q79—A123 H-bond, involved in the stabilization of $OCP^{R}$, could explain the experimentally observed faster recovery of the Q79L mutant (Supplementary text 3 and Supplementary fig. 14A, D). The increased stability of the C2c state in this mutant could play a role in its faster recovery. Indeed, as reformation of $OCP^{O}$ and $OCP^{1hv}$ associates with a similar 'dark' state spectrum, it is not directly possible to assess whether the $OCP^{R}$ repopulates the former via the latter (as expected from micro reversibility) or if a direct path exists from $OCP^{R}$ to $OCP^{O}$. Fits of our kinetics data, which go beyond the scope of the current paper but will be reported elsewhere, are suggestive of the latter hypothesis.

**Supplementary text 2: The C2o to C2c transition can be controlled by illumination**

The relevance of the C2c conformation was, at a first glance, questionable. Indeed, it is known, by inference from the isolated NTD structure[10] and from cryoEM structures of photoactivated OCP$^{R}$ in complex with the phycobilisome[11], that for OCP to photoactivate to OCP$^{R}$, the carotenoid must undergo at 12 Å translocation across the carotenoid tunnel (Fig. 2A), emerging at the exit of SC#1[11]– which is closed in the C2c conformer (Fig. 2A, B). Thus, the C2c conformer could be a candidate for an inhibited state involved in some sort of regulation – as suggested earlier by Maksimov and coworkers[6], a crystalline artifact or a functionally – though counterintuitively – relevant conformational sub-state. Our first test of a possible functional relevance of the C2c conformation was to illuminate C2o crystals for various periods of time to determine if it can be induced by light (Supplementary table 1). We then used structure-factor extrapolation to assess the structural changes induced by illumination. Again, unless stated otherwise, CAN-functionalized OCP was used in the experiments described below.

Starting from C2 macrocrystals (~150 µm) at 97 % RH featuring the C2o conformer at 100% occupancy, we found that the C2c conformation accumulates to ~50% upon 1 min illumination at 537 nm, as revealed by structure factor extrapolation[12,13] between datasets collected on the same crystal, before (dark) and during/after illumination (data collection time was ~1 min) (Fig. 3A and Supplementary figs. S8 and S9). Briefly, in the 1-min illuminated structure, the D35(OD1)—Y129(OH) and D35(OD1)/S132(N) H-bonds are broken, the $3_{10}$ helix has formed and thus both P126 and Y129 block the outlet of SC#1 (Fig. 3A and Supplementary fig. S8). Notably, D35 undergoes a conformational shift, positioning its side chain within the cavity created by the displacement of Y129 ($\Delta\chi1$~180°), allowing D35(OD2) to now H-bond with both S132(OG1) and (N). The Q79(OE1)/A123(N) H-bond, however, is seen to rupture. This observation is suggestive of the main chain oxygen of A123 being able to serve as a pivot point for conformational changes occurring in the αGH loop (Supplementary fig. S8). Overall, changes at the αC/αGH and αE/αGH interfaces result in the αGH loop being pushed away from the NTD core (and therefore, from the CTD as well) (Supplementary fig. S9). Resultingly, two conformations are observed for W41 and close-by L45 and M117. Photo-triggered changes also include loss of the $P2_1^A$ and C2-specific interdomain Q149(ND2)—A309(O) H-bond which, in the dark state structure, anchors the base of αN to the tail of αH (Supplementary figs. S2 and S8). Loss of the latter interaction concurs with a repositioning of the D306-R319 segment (end of β7 and αN), with a simple H-bond swap at D306/R291 (the OD1 to NH2 H-bond is replaced by an H-bond to OD2), but complete rupture of the A309(N)—L16(O), L314(O)—R256(NH1), and N316(ND2)—N14(O) H-bonds. Absence of these interactions results in loss of the R319(NE)—E260(OE2), P261(O)—Q268(NE2), and R291(NH1)—R9(O) H-bonds, partly clarifying how light-induced changes "travel" from the carotenoid in the CTD to W290 to the β4-β5-β6-β7 portion of the β-sheet to the αA and αN helices. Illustratively, we see a large conformational change in L293, at the interface between αA (F3, A8, I11), β4 (A262), β5 (Y266) and β7 (F301, F302). No changes are seen at the β1 ionone ring, which remains H-bonded to Y203 and W290 after 1-min illumination (Supplementary fig. S8). Nevertheless, there are changes in the carotenoid structure. At the β2 ionone ring, we observe a ~25° "bending" of the carbonyl bringing the carotenoid O' closer to W110 (0.5 Å) together with a change in pucker state at C3'. Large (Δ ≥ 40-55°) alternating twists in the polyene chain are observed at C7'-C6', C10'=C9', C14'=C13', C15-C14, C13=C14 and C12-13 (Supplementary fig. S8), as well as more minor (≤ 30 °) twists at C9'-C8', C11'-C10', C15'-C14', C11=C12 and C9=C10. This combination of twists is accompanied with positional changes of tunnel-lining residues (Y44, I51, M83, L86, L107, Y111, I151, R155 and F280) as well as of the structural elements that harbour them (αC, αE, αG, αI and the β5-β6 turn). Notably, we see a transduction path for structural changes trailing from the carotenoid to the β5-β6 turn (T277-F280), to the αCD loop (loss of the T52(N)—W279(O) and T52(OG1)—P278(O) interdomain H-bonds), and to the C-terminus of αC (loss of the T50(OG1)—M47(O) H-bond stabilizing the $3^{10}$ helix characteristic of C2o, C2c and $P2_1^A$). The repositioning of the linker is accompanied by rupture of H-bonds to αK (V180(O)—N210(ND2) and V180(N)—A209(N) are broken) and β4 (Q248(NE2)—V178((O) and Q248(OE1)—V180((N) are broken) and by the formation a new Q248(ND2)—R244(O) H-bond. This coincides with stabilization of the R244 side chain between the E176 and E213 carboxylates (alternate conformers), while that of R241 draws away from E176 to interact with either D237 or E245 (alternate conformers). A new salt-bridge forms between E217 and K236 side chains, in addition to that linking E217(O) and K299(NZ), in the C2o dark-state structure. Thus, the face of the β-sheet opposite to αN and αA, also propagates the light-induced changes in the carotenoid to the surface of the protein, with an apparent effect on the electrostatics.

We examined whether photo-triggered conformational changes accumulate in the C2 crystal form. For this, we illuminated a rotating crystal (~150 µm) for 10 minutes while rotating it in the optical beam and collected diffraction data during the final minute. A pre-illumination dark dataset was recorded on the same crystal prior to the extended illumination enabling structure-factor extrapolation to probe the structural changes induced by the additional light exposure (and/or reaction time). (Supplementary table 1). The largest structural changes occur in the αGH loop. Indeed, relative to the dark-state starting point—which contained the C2o and C2c conformers in a 6:4 ratio—the 10-minute illumination structure again shows light-driven accumulation of the C2c conformer, with an apparent ~30% increase of its occupancy (Supplementary table 1). In extrapolated electron density maps, the side chain of Q79 is well defined and aligns perfectly with that in the C2c conformer accumulated upon dehydration, *i.e.* with the OE1 oxygen at 3.6 Å from A123(N), and NE2 at H-bonding distance from I72(O) and M74(O) (2.6 and 3.2 Å) (Fig. 3A and Supplementary figs. S8 and S9). At the N-terminus of the αC helix, we observe D35 in two alternate conformations, corresponding to either the dark-state conformation (as seen in C2c and C2o), or the conformation observed after 1 min illumination (see above). The αH conformation is, however, more dark-like than in the 1-min illumination structure. Notably, the C2-specific Q149—A309 H-bond bridging αH to αN reforms, allowing E146, N287, and (consequently) P311 to shift back toward their dark-state positions. Likewise, we observe the reformation of the A309(N)—L16(O), L314(O)—R256(NH1), and N316(ND2)—N14(O) H-bonds upon the repositioning of αN. At the nearby salt bridge between R291 and D306, the observed configuration is, alike for D35, in-between the dark and the 1-minute illumination structures, *i.e.*, a single H-bond is observed from D306(OD2) to R291(NH2) and (NE), however the OD1 atom is now closer to NH2 and the R291(NH1)/R9(O) H-bond is restored. The Q268(NE2)/P261(O) H-bond is also reformed, with the terminal R319 side chain now H-bonding to Q268(OE1) via its NH1 atom – as opposed to being H-bonded to D260 in the dark-state, and without H-bonding partner, in the 1-min illumination structure. The large conformational change at L293, observed after 1-min illumination, has also reverted. Nevertheless, substantial conformational changes occur in the αCD loop, whose C-terminal segment refolds such that it now matches the conformation observed in the P21$^{A}$ protomer—a configuration not previously seen in the C2 crystals (Supplementary fig. S9). The C2-specific Y44/E115 H-bond is accordingly lost. However, the T52(OG1)–P278(O) and T52(N)–W279(O) H-bonds that tether the αCD loop to the β5β6 turn in the P21$^{A}$ and C2/C2c dark conformers are restored, as is the T50(OG1)–M47(O) H-bond, stabilizing the kink at the C-terminal end of αC. Nevertheless, the Q248(OE1)—V180(N) H-bond remains broken, connecting the linker to β4. Thus, at the protein level, the 10-minute illumination structure appears in between the C2/C2c dark-state structures and the 1-minute illumination structure. Likewise, at the carotenoid level, most of the conformational changes listed for the 1-min illumination structure have reverted after 10-min illumination. Notably, the β2 ionone ring dark-state configuration is restored, alongside planarity at most polyene bonds. An exception is the C14'=C13' bond, which undergoes dihedral "inversion" upon prolonged illumination (Δ[10-minute – 1-minute] ~90°). Minor accompanying twists are seen at C8-C9, C15=C15' and C15'-C14'. Together, these changes "push" the C14'-C8' segment of the carotenoid toward the tunnel floor, paved by the side chains of Y44, M47, I51, I151, in the NTD, and V275, T277, F280, M286 and M288 in the CTD.

**Supplementary Text 3: The C2c conformer is an obligatory intermediate in OCP photoactivation**

Our structure determined after 1 min illumination shows that the conformational transition between the C2o and C2c conformers can be induced by light. Given the recent demonstration that OCP photoactivation requires the absorption of two consecutive photons[14,15], we hypothesized the following mechanism: (i) upon absorption of the first photon by dark-state $OCP^{O}$, with SC#1 open, the protein transitions to the closed $OCP^{1hv}$ intermediate; (ii) upon absorption of a second photon by this intermediate, with a potentially higher likelihood of carotenoid untethering, the protein reverts to the open state, enabling carotenoid translocation through the NTD tunnel and formation of $OCP^{R}$. We used mutagenesis to test this hypothesis. The effect of amino acids substitutions on the structure, photoactivation and recovery rates were probed with the tools of X-ray crystallography (RT data collection, unless stated otherwise) and steady-state spectroscopy (extracting both the photoactivation and recovery rates). Note, that compared to $WT_{ECN}$, $WT_{CAN}$ shows photoactivation and recovery rates that are 74 and 33 % faster[1], respectively (Fig. 3C).

We questioned the relevance of the C2c conformation by engineering a constitutively-open A38C-I125C mutant, unable to reach the C2c conformation due to formation of a disulfide bridge, thereby coVently trapping the open form (Supplementary fig. S12). The structure, solved in the C2 space group, confirmed the design – notably the fact that the SC#1 channel is open, regardless of the RH at which data are collected (Supplementary fig. S12A and Supplementary tables 4 and 5). The disulphide-bridged A38C-I125C OCP mutant was inactive, regardless of whether it was functionalized with CAN or ECN (Supplementary fig. S12B, C and Supplementary table 6), while the control A38C and I125C mutants were both active, though with reduced photoactivity (60 and 30% drop in initial rate of photoactivation with respect to $WT_{CAN}$, respectively) and faster recovery (620 and 470 % increase in initial rate of recovery with respect to $WT_{CAN}$, respectively) (Supplementary fig. S12D, G and Supplementary table 6). Importantly, the A38C-I125C mutant is unable to quench the PBS (here assessed using the core base (CB) of the PBS), regardless of the functionalizing carotenoid (Fig. 3D, E). This establishes that transition to the C2c conformation is an obligatory step in the OCP photoactivation mechanism.

We investigated the importance of (temporarily) shielding the carotenoid from the solvent by engineering the A38C-I125C-T80W mutant, also unable to reach the C2c conformer but whose SC#1 channel can close with the aromatic side chain acting as a surrogate door (Supplementary figs. S12 and S13). Photoactivity was partially restored in this protein, demonstrating the importance of carotenoid insulation from the bulk and suggesting that it is this shielding why why the C2c state needs to form (Supplementary figs. S12 and S13).

In *Planktothrix agardhii* OCP, the defining features of the C2c conformation are ruptures of the intramolecular Q79/A123 and D35/Y129 H-bonds. We investigated the importance of the Q79/A123 H-bond for stabilizing the open or closed conformation by characterizing the Q79L mutant where H-bonding between αE and the αGH loop is suppressed (Supplementary fig. S14A, B and Supplementary tables 5 and 6). Photoactivity was halved in the $Q79L_{CAN}$ mutant (-52% and +1900 % change in photoactivation and recovery rate) and quartered in the ECN counterpart (-77% change in photoactivation rate but recovery too fast to be measured) (Supplementary fig. S14A, B and Supplementary tables 5 and 6). The structures of the CAN- and ECN-functionalized Q79L mutant were solved in the C2 space group and showed 100% occupancy of the C2c structure regardless of the RH during data collection (Supplementary fig. S14A, B and Supplementary table 5). Although the structure of OCP-X (PDB 8a0h) shows a closed C2c-like conformation in the presence of the analogous Q78/A122 H-bond, our computational analysis (see Supplementary Text 1) strongly supports that in case of *Planktothrix agardhii* OCP, the H-bond must be formed for the stability of the C2o state to exceed that of the C2c state, and that preventing formation of this H-bond skews the dark-state equilibrium toward the C2c state – at least for $Q79L_{CAN}$. This hypothesis is supported by MD simulations (Supplementary text 1). The reduced $OCP^R$ accumulation for the Q79L mutant is consistent with the fact that this H-bond also forms in the $OCP^R$ state, which is particularly noticeable for ECN-functionalized $Q79L_{ECN}$, showing an exceptionally fast recovery rate with a clear bi-exponential behavior. Nevertheless, both $Q79L_{CAN}$ and $Q79L_{ECN}$ remain able to quench the PBS (here assessed using the core base (CB) of the PBS) (Fig. 3D, E). This result indicates that upon binding of this mutant to the PBS, the $OCP^R$ state stabilizes despite absence of an H-bond to A123 to contribute to its stabilization. Together with data obtained on the A38C-I125C mutant, these results establish that access to the C2c conformation is required for the protein to be functional: while it is possible for the protein to access $OCP^R$ from the C2c conformer, it cannot do so directly from the open conformations ($P2_1^B$, $P2_1^A$, C2o).

We similarly interrogated the role of the D35/Y129 and D35/S132 H-bonds by producing the D35T mutant, expected to lack both H-bonds due to the shorter side chain (Supplementary fig. S14E, H and Supplementary tables 5 and 6). The polar threonine side chain was selected to maintain the local hydrophilicity. The structures of the $D35T_{CAN}$ and $D35T_{ECN}$ variants were solved in the C2 space group. They confirmed lack of interaction with Y129 and/or S132 and showed high occupancy of the C2c conformer, with fractional occupancies of 0.8 and 1.0 at 97% RH, respectively (Supplementary fig. S14E, F and Supplementary table 5). Regardless of the functionalizing carotenoid, the mutant was photoactive (Supplementary fig. S14G, H and Supplementary table 6), with $D35T_{CAN}$ and $D35T_{ECN}$ showing 40 and 70 % reduced photoactivity, and an increase of 33 and 300 % for thermal recovery, respectively (Supplementary table 6). These results indicate that like the Q79/A123 H-bond, the D35-mediated H-bonds are dispensable to navigate between the C2o and C2c states. Unlike the Q79/A123 H-bond, however, they have a limited impact on the $OCP^R$ to $OCP^O$ recovery, consistent with the replacement of the D35—Y129 H-bond, in the $OCP^O$ state, by the E34—Y129 H-bond, in the $OCP^R$ state. Again, the increased effect of the D35T mutation in the ECN variant highlights the impact of the carotenoid on the equilibrium between the open and closed αGH loop conformations (Supplementary table 6).

The C2o and C2c conformers share the feature of broken intermolecular P13/A133 H-bonds, suggesting that rupture of the latter could be a pre-requisite to accessing the C2c state. We investigated the impact of this H-bond by engineering the A133P mutant, unable to engage in a H-bond with its tertiary main chain nitrogen (Supplementary fig. S14I, J and Supplementary table 5). Relative to the WT, the mutation was fully silent in terms of photoactivation and recovery (Supplementary fig. S14K, L and Supplementary table 6). As expected, only C2 crystals (wherein the P13/A133 H-bond is constitutively broken) were obtained, which showed a similar propensity as the WT to accumulate the C2c conformer (Supplementary figure S14I, J). This was further confirmed by solving the cryoEM structure of the A23C-A133P mutant, which again best matched the $P2_1^B$ like conformation characterized by open SC#1 and SC#2 and an extended and straight αC helix. The mutation nevertheless fully restored the recovery rate of the WT when introduced in the A23C scaffold, demonstrating the importance of this H-bond in navigating back and forth between the $P2_1^B$/$P2_1^A$/C2o and C2c states.

Altogether, these data strongly support that that back-and-forth transitions between the C2o and C2c conformers are required for $OCP^O$ to form $OCP^R$, upon photoactivation, and reform thereof, upon thermal recovery. Accordingly, the $A133P_{CAN}$, $D35T_{CAN}$ and $Q79L_{CAN}$ proteins were able to quench the phycobilisome, whereas the $A38C\text{-}I125C_{CAN}$ mutant was not (Fig. 3D, E). In this context, it must be highlighted that the central residues for this transition, including P13, D35, W41, Q79, A123, I125, P126, S132 and A133, are strictly conserved in OCPs from OCP1 clade[16] (Supplementary table 10).

**Supplementary Text 4: Temperature-controlled scanning-fluorimetry on CAN- and ECN-functionalized WT and mutant OCP**

To obtain further information on the energetics of the structural dynamics of OCP, we performed temperature-controlled scanning-fluorimetry (TCSF) of WT and mutant variants, complexed with CAN or ECN, complementing the structural and functional insights described in Supplementary Text 3. TCSF measurements were recorded at 350 nm (excitation at 280 nm), reporting on the overall level of solvation of tryptophan side chains, to probe structural distributions and stability *in vitro*[17]. We found, by characterizing the WT proteins, that the thermophoresis profile of $OCP_{CAN}$ features a concentration-dependent peak, which is unrelated to denaturation of the protein (no increase in scattering), shifts from 42 to 47°C upon increase of the concentration from 30 to 160 μM (corresponding to dimer fractions of 70% and 95%, respectively[2]), but is absent from the thermophoresis profile of $OCP_{ECN}$, where only the major denaturation peak (associated with scattering) is observed at ~54°C (Fig. 3B and Supplementary fig. S10). By assaying the constitutively monomeric $R27L_{CAN}$ mutant, known to be more photoactive and recover faster than WT[18], it was established that observation of a concentration dependent temperature-induced peak-shift depends on the presence of dimeric protein and thus informs on changes in a structural element that is affected by dimerization (Supplementary fig. S10) .

None of the structural elements contributing to the dimer interface (by decreasing contribution to the buried surface area: αB, αA, αH/αGH, the αEF turn and the β2-β3 turn) feature a tryptophan residue. However, the αGH loop, contributing to the interface via the intermolecular P13(O)/A133(N) H-bond, can adopt two conformations that are characterized by distinct van der Waals interactions of W41 – namely, with I125 and P126 in the C2o and C2c structures, respectively (Supplementary fig. S7). Additionally, the ring-to-ring W41—Y44 distance differs in the open ($P2_1^A$, $P2_1^B$, C2o) and closed (C2c) forms of the proteins, which could modulate the efficiency of a potential energy transfer from Y44 to W41. Indeed, the C2c state associates with the conformation of Y44 that is H-bonded to E115 and resultantly closer to W41 (Fig. 2A and Supplementary fig. S2).

We tested the hypothesis that the peak informs on αGH loop structural dynamics by producing the W41F mutant (Supplementary fig. S11A). The $W41F_{CAN}$ protein displays a 2-fold decrease and 3-fold increase of the initial photoactivation and recovery rates compared to $WT_{CAN}$, respectively (Supplementary fig. S11B, C and Supplementary table 6). It features a dramatic decrease of the concentration-dependant peak (Supplementary fig. S11A), supporting that in the CAN-WT, this peak relates to changes occurring in the local environment of W41 upon elevation of temperature to around 40° C or 47 °C, depending on oligomerization state (monomer vs. dimer).

Next, we used TCSF to examine the series of mutants that had been designed to probe the role of the different αGH loop configurations, and are thus characterized by varied propensity to take up the C2o and C2c conformations (see Supplementary text 3). The structural and functional characteristics of the mutants are described in Supplementary text 3 and Supplementary table 5.

Since the intermolecular P13(O)/A133(N) H-bond is an essential part of the dimer interface, we first investigated the A133P$_{CAN}$ mutant (displaying photoactivation and recovery rates similar to WT$_{CAN}$). The effect of the A133P mutation on dimer affinity was probed by analytical ultra-centrifugation (Supplementary fig. S19C), which showed that the P13/A133 H-bond stabilizes the dimer to a lesser extent than the R27/D19 salt bridge. An attempt was made to obtain a dissociation constant using the same method employed for the WT. Data for the mutant were of lesser quality than those recorded on the WT protein, though indicative of a 5-to-10-fold reduced affinity, in line with analytical ultracentrifugation results. Nevertheless, the thermophoresis profiles of A133P$_{CAN}$ and R27L$_{CAN}$ are rather similar, lacking the concentration-dependent peak around 47 °C (Supplementary figs. S10 and S19C). This strongly supports the interpretation that the P13(O)/A133(N) H-bond –and thus the αGH loop structure– contributes to the conformation associated with the concentration-dependent temperature peak-shift seen in the TCSF data of CAN-functionalized proteins. To further our analysis of the role played by this H-bond, we introduced it in the constitutively dimeric A23C mutant. In the thermophoresis profile of the A23C$_{CAN}$ protein, showing a high propensity to form the P13(O)/A133(N) H-bond, the concentration-dependent 42-47°C peak is strongly reduced (Supplementary fig. S19C). Addition of the A133P substitution to the A23C scaffold allowed to restore the 42°C peak; however, similar to the observation for the A133P single mutant, no concentration dependency was observed (Supplementary fig. S10). Thus, the 40-47° C peak observed in the WT$_{CAN}$ TCSF data reflects the C2c to C2o structural changes in the αGH loop that alter the environment of W41, and its concentration-dependent temperature shift arises from formation of the P13(O)/A133(N) H-bond upon dimer formation (Supplementary figs. S10, S11 and S19), further stabilizing the C2o conformation.

We next examined the constitutively-open A38C-I125C mutant (Supplementary fig. S13). We found that regardless of whether CAN or ECN is used as the functionalizing pigment, the constitutively open A38C-I125C mutant features only the denaturation peak. This result further supports the hypothesis that the peak observed at 42-47°C reflects the αGH-loop structural transition leading to formation of the closed state. We also examined the control single A38C and I125C mutants. Unexpectedly, the peak is present at 34-44°C in the I125C$_{CAN}$ mutant (Supplementary fig. S13), indicating reduced stability (Δ~8 °C) of the closed state conformation upon insertion of a cysteine at position 125 – a result in line with the changed photoactivation and recovery rates of the mutant (decreased by 27% and increased by 5-fold, respectively). Nevertheless, the protein accumulates to a similar level in the OCP$^R$ state. Conversely, the A38C$_{CAN}$ mutant features a concentration dependent peak at 44-47°C, indicating increased stability of the closed state conformation. This is supported by a 12-fold increased recovery rate of this mutant, which together with a 60% drop in the photoactivation rate results in a markedly reduced accumulation of the $OCP^R$ state (Supplementary table 5). These observations illustrate that an unbalance in free energy between the conformationally open and closed states negatively impacts the $OCP^R$ yield, as also supported by computational analysis of the Q79L mutant (see section *Equilibrium structural dynamics* in Supplementary text 1).

The different carotenoid complexes of the Q79L mutant also show interesting differences in the TCSF data (Supplementary fig. S15A). The $Q79L_{CAN}$ variant, displaying a 51% decreased photoactivation rate alongside a 20-fold increased recovery rate (Supplementary fig. S14C, D), features the 42°C peak (though observed at 39°C) at all concentrations (Supplementary fig. S15A), *i.e.* the concentration-dependent peak-shift is fully abolished. This result is indicative of the protein residing in the closed-state *in vitro*, with no possibility for the P13–A133 to form, even in the dimer. Interestingly, the photoactivation and accumulation of the $OCP^R$ state, as well as recovery of the dark-state(s) ($OCP^O$/C2c), are comparable in the $Q79L_{CAN}$ and $A38C_{CAN}$ mutant.

Last, we examined the D35T mutants in their CAN- and ECN functionalized states. A loss of dependency on concentration and increase in amplitude of the 42°C peak is also observed in the $D35T_{CAN}$ mutant (Supplementary fig. S15B), whereas the $D35T_{ECN}$ mutant shows the same profile as $WT_{ECN}$. These results confirm the hypothesis that αGH loop dynamics are, at equilibrium, only present in the CAN-functionalized protein, *i.e.,* the C2c conformer is not accessible thermally in the ECN-functionalized scaffold – neither in the WT nor the mutant variants. Indeed, the thermophoresis profiles of the ECN-functionalized mutants are all devoid of the peak at 42-47°C (Supplementary figs. S10 and S15 and Supplementary table 6).

In conclusion, by combining the TCSF results with the structural and functional information on WT and mutant OCP (Main text, Supplementary Text 3 and Supplementary table 6), we infer that the concentration-dependent peak represents the fraction of the protein that resides in the C2c state; because this state is metastable, it depopulates with raising temperature, for the benefit of the C2o population. The presence and absence of the peak in the CAN vs ECN-functionalized proteins (Fig. 3B) suggests that *in vitro*, $OCP_{ECN}$ occurs in a homogeneous open dark-state ($P2_1^B$, $P2_1^A$) whereas $OCP_{CAN}$ is also present in the closed C2c conformer. Since it has been shown previously that in $OCP_{CAN}$, the one-photon intermediate state is thermally accessible, we assign the TCSF peak at 42-47°C – and thus the C2c conformer – to $OCP^{1hv}$. Evidently, the difference between the two carotenoid complexes stems from the presence of the β2 carbonyl oxygen of CAN and its interactions within the tunnel; these favor the transition to the $OCP^{1hv}$ intermediate at equilibrium, in the dark. Nevertheless, SC#1 closing and opening can as well be triggered in $C2_{ECN}$ crystals, by means of both illumination and a decrease in the humidity. Thus, the proposed mechanism, whereby a first absorbed photon induces carotenoid insulation, enabling a second photon to yield an untethered state with a lifetime compatible with that required to reopen SC#1, would apply to both ECN- and CAN-functionalized OCP. In the latter, however, a fraction of OCP molecules is already present in the $OCP^{1hv}$ intermediate state (C2c) and therefore evades the requirement to sequentially absorb two photons to access the $OCP^R$ state.

**Supplementary text 5: Pump-probe TR-SFX illuminates OCP photocycle**

Taken together our structural and functional data very strongly support that the $P2_1^{A,B}$ and C2c structures correspond to dark-adapted $OCP^O$ and the 1 photon-intermediate $OCP^{1hv}$, respectively. Thus, two light-driven reactions are involved; that enabling transition from the dark-state to the $OCP^{1hv}$ intermediate; and that yielding $OCP^R$ from the 1-photon intermediate. By applying time-resolved serial femtosecond crystallography (TR-SFX) to the $P2_1$ and C2 crystal forms, we shed light on both pathways. Experiments were performed using a ~5 μm X-ray beam at the SwissFEL, and focused either on the ultrafast events using 85 fs optical pulses to trigger the reaction (Δt=0.3-100 ps; Alvra instrument), or on characterizing longer-lived intermediates, using ns pump pulses (Δt=10 ns, 1 μs; Cristallina-MX instrument) (Supplementary tables 7 and 8). OCP microcrystals (average size ~3 μm ± 0.5 μm) were grown by the micro-batch method[19], embedded in LCP (see Methods) and shortly afterwards presented to the X-ray beam by means of a High-Viscosity-Extruder (HVE) operating at RT[20] (Supplementary fig. S20, A, B). With view of maximizing light penetration into the optically-dense crystalline sample (and as well because transient absorption spectroscopy had shown simpler photodynamics at this wavelength compared to that upon excitation at 490 nm[21]), we used 535 nm laser pulses as the actinic source ($\varepsilon^{535\ nm}$ = 47,250 $M^{-1}.cm^{-1}$) and probed structural changes using ~5 μm-focused, 40 fs-duration X-ray pulses. Based on transient absorption spectroscopy power titrations carried out on OCP solutions[21] as well as on OCP microcrystals ($P2_1$ and C2, see Supplementary Text 6), it is known that the $S_2$ yield (1100 nm band)[21] deviates significantly from linearity at power densities above 18 GW/$cm^2$, yielding a long-lived, off-pathway, radical-state[21] (Supplementary text 6 and Supplementary figs. S21-S23). Accordingly, our first set of TR-SFX data was collected at a power density of 18 GW/$cm^2$ (fs laser) – translating to 0.4 and 0.75 nominally absorbed photons per carotenoid on average and on the first crystal layer, respectively – and at time-delays of 0.3, 1.5, 10 and 100 ps, corresponding to the peak occupancy of the ICT, S1, S* and P1 states. A second set of TR-SFX data probed time delays of 10 ns and 1 μs following excitation by a 3.7 ns pulse at 17 mJ/$cm^2$ energy density, resulting in a very low power density of 5 MW/$cm^2$. In this safe excitation regime, five nominally absorbed photons were delivered per carotenoid with hope for re-excitation during the pulse resulting in increased photoproduct occupancy.

Due to the experimental nature of the work, different ratios of $P2_1$ and C2 crystals were probed during the two experiments. The $P2_1$ crystals amounted to 2-10% of all probed crystals in the TR-SFX data collected at short time delays, with the remainder corresponding to C2 crystals featuring the C2c conformer at an occupancy of 1. Data were collected in an interleaved pump-probe fashion, resulting in a large amount of dark-interleaved data (800,000 and 80,000 patterns indexed in C2 and $P2_1$, respectively). We tested the feasibility of using these in a highly-redundant dark reference dataset (which also allowed to check for light-contamination, that is, the accidental illumination crystals that contribute to the laser-off data set), by collecting a large dark-only dataset (200 and 20 k patterns indexed in C2 and $P2_1$, respectively), which allowed investigation of potential structural differences by means of dark-interleaved vs. dark-only Fourier difference maps and extrapolated structures. Even with the highly redundant C2 data, we found structural differences to be negligible, amounting to 5% occupancy for a state with structures virtually identical (rmsd=0.275 Å) to that solved with the merged data, for both the protein and carotenoid. Accordingly, nearly identical extrapolated maps and structures were obtained when comparing the light data to either sets of dark data or their combination thereof. Hence, we opted for redundancy, and all dark data were merged together, resulting in higher quality extrapolated maps as well as better statistical indicators for the refined extrapolated structures (Supplementary tables 7 and 8). Note that given the scarcity of $P2_1$ crystals in the sample used during this first experiment, the decision to merge dark-only and dark-interleaved (all-dark) data was drawn from examination of the C2 dark data, and thereafter applied to the $P2_1$ data as well. As judged from the comparison to the C2c *all-dark* dataset, the occupancy of triggered states in the C2 light datasets yield an occupancy of 8% at 0.3 ps, and 7% at all other short time delays (1.5, 10, 100 ps). These occupancies are in stark contrast to those determined for the $P2_1$ light datasets, where they vary from 40% at 0.3 ps to 20% at the other time delays and energies. Most likely, The $P2_1$ occupancy values are artificially-high due to the significantly lower quality of the $P2_1$ short-time delay data. Indeed, the apparent occupancy of intermediate states is highly skewed by the quality (strongly coupled to redundancy, when it comes to serial data) of the compared reference and triggered data in structure factor extrapolations endeavours. It is crucial to understand that regardless of the method used to determine intermediate state occupancies, these only marginally inform on the actual occupancy, merely on the best multiplication factor to apply to difference data to observe the intermediate at the highest signal to noise ratio in extrapolated electron density maps. Illustratively, structure factor extrapolations between datasets collected on crystals of wild-type and mutant proteins do not yield an occupancy of 1. Occupancies determined by extrapolation methods should thus only be compared across datasets of the same quality – *i.e.*, in the present case, assembled from a similar number of crystals of similar symmetry – and never be taken as absolute. They can be compared across the C2c data collected at short time delays, because of the similarly high redundancy of the triggered (light) data and usage of the same reference (dark) dataset, but they cannot be compared to those determined from the $P2_1$ data, which are of much lesser quality due to the ~10 times reduced number of crystals used in the datasets.

Merging of the dark-only and dark-interleaved data was not possible for the data collected at longer time-delays (Supplementary tables 7 and 8). Indeed, for both crystal forms and both time-delays, only the dark-interleaved data collected intermittently with the light data could be used as reference for intermediate-state structure extrapolation. Fortunately, this drawback was compensated by a larger fraction of $P2_1$ crystals (50% of all indexed patterns) in the batch of sample used for this experiment; all our dark-interleaved and light datasets exceeded 100,000 indexed patterns. In both the dark-interleaved and dark-only data, C2 crystals were found to feature the C2o and C2c conformers at ~50% occupancy. Nevertheless, in the corresponding light vs. dark extrapolated maps, electron density is observed only for the C2c conformation. Occupancies of the triggered states were found to amount to 3.5-4% in the C2 crystals, and to 7-8% in the P21 crystals (Materials and Methods). Since the P21 and C2 data acquired at long time delays have been assembled from a similar number of crystals, one could be inclined to here trust that the occupancy of intermediate states is higher in the $P2_1$ than in the C2c crystals, and from this infer that the photoactivation yield is higher for $OCP^O$ than for the $OCP^{1h\nu}$ intermediate. Indeed, the spectral properties of these two dark states are identical and their concentration is similar in the two crystal forms. However, the symmetry of the C2 crystals is higher, which likely explain both the higher resolution of the data and the ability to utilize higher multiplication factors (1/occupancy) of the difference data to observe electron density of the intermediate state at the highest signal to noise ratio.

Overall, our data show that the structural changes triggered by photoexcitation of the carotenoid are distributed across the whole OCP dimer – including the dimerization interface – but with different outcomes depending the starting dark ground-state. In absence of strong peaks in the Fourier difference maps – possibly also reflecting a mixture of excited states characterized by a different electronic structures, conformations and lifetimes – extrapolated structures formed the main source of insights into photo-triggered conformational changes. Under the working hypothesis that at any time delay, the intermediate at peak occupancy is that dominating in the extrapolated density maps, we refined structures at each time delay and energy and examined the change in overall conformation of the protein and the carotenoid (Supplementary tables 7 and 8). In the three chains, the most noticeable outcome of photoexcitation is large conformational changes in the αCD and αGH loops, triggered as early as 0.3 ps and up to 1 µs (Supplementary figs. S32-S37 and S39). Thus, at short as well as at longer time delays, and under pulsed as well as continuous illumination (conventional MX and SSX data), the two NTD loops are the main "relays" for transfer of excitation energy from the carotenoid to the protein scaffold (Figs. 4 and 6, and Supplementary figs. S8, S9, S32-S37 and S39). Noticeable in the two $P2_1$ chains is a sudden change in carotenoid structure upon photoexcitation, incurring loss in the protein's H-bonding interactions as early as 0.3 ps (Supplementary fig. 30). By 100 ps - 10 ns, however, both the carotenoid structure and the H-bonding network of the protein have virtually recovered, despite the latter still exhibiting minor deviations from the dark state (Figs. 4 and 5A, and Supplementary figs. S25-S29). By contrast, in the C2 chain changes in the carotenoid structure are less dramatic at short time delays but span the whole time series with a slow but continuous loss of protein's H-bonding interactions up to the 10 ns timescale (Figure 5A and 6, and Supplementary figs. S28-S29 and S38). However, the triggered cascade of conformational changes lasts for more than 1 µs as evidenced by the $C2new^{light\text{-}48h}$ structure (Fig. 6 and Supplementary figs. 30-35 and 38-41) but as well the $P2_1^{B\text{-}light\text{-}48h}$, $P2_1^{A\text{-}light\text{-}48h}$ and $C2^{light\text{-}48h}$ structures (*vide infra*).

**Supplementary Text 6: Ultrafast optical spectroscopy on $P2_1$ and C2 OCP microcrystals**

A recent visible–near infrared (NIR) femtosecond time-resolved absorption spectroscopy studied the ultrafast photodynamics of OCP from *Synechocystis* PCC 6803 as a function of the excitation pulse power and wavelength[21] and found that a carotenoid radical cation can form at relatively low excitation power[21]. Since it was also shown that His-tagging at the N- or C-termini affects the excited-state photophysical properties[21] we decided to characterize the photodynamic response of the two *Planktothrix aghardii* crystal forms ($P2_1$, C2) by NIR femtosecond transient absorption spectroscopy to establish (i) conditions for single photon excitation, avoiding non-linear effects reflected in the radical formation and (ii) whether the crystal lattice or crystallization conditions affect the photodynamics. To this end we performed (i) a power titration and compared (ii) the ultrafast dynamics of solution phase OCP and crystalline OCP.

*Power titration results*

We performed power dependence measurements in the energy range of 0.2–3.2 μJ. When plotting the temporal and spectral evolution of the ground state bleach (GSB, 510 nm), $S_1$ and ICT excited state absorption (ESA, 670 nm) and of the $S_2$ excited state absorption (1100nm) signals as a function of the pump peak power density, one can clearly see deviations of linearity (Supplementary fig. 21A). The power density used for microcrystal photoexcitation during the SFX experiment performed at the Alvra instrument at SwissFEL was 18 GW/cm$^2$ (low power density (LPD), green dashed vertical line in Supplementary fig. 21A). The LPD pump energy selection was governed by the need to maximize the excitation energy (to increase the occupancy of light-induced states), while keeping the pump energy density close to the linear excitation regime and the carotenoid cation radical formation below the detection threshold. We have chosen an energy that is near the end of the linear regime to compensate for optical losses by Fresnel reflections at the air/jet interface. For comparison, we also investigated the dynamic response when exciting with much higher power density (HPD), indicated by a red dashed line in (Supplementary fig. 21A. The HPD pump power excitation results in saturated signals, moreover significant contribution of multiphoton excitation events can be expected resulting in formation of a certain number of carotenoid cation radical species. Usage of HPD pump power density in the power titration experiment resulted in fast sample degradation and presence of detectable carotenoid cation radical signal at 950 nm (Supplementary fig. 21B, C). In general, the signals saturate slightly faster for the colloidal crystal sample (PEG, $P2_1$) compared to one of the crystals embedded in viscous LCP (C2, C2c conformer), despite the fact that the sample stationary absorbance peak overlaps better with 532 nm excitation for the LCP sample. Supplementary fig. 21D,E shows an overview of the evolution of the spectra and kinetics.

*Comparison of the ultrafast dynamics observed in crystalline and solution phase OCP*

To investigate a potential influence of the crystal lattice or crystallization conditions on the ultrafast dynamics of photoexcited OCP probed by TR-SFX we used the same experimental setup as for the power titration and a pump laser peak energy and power density of ~1.2 mJ/cm$^2$ and ~10 GW/cm$^2$, respectively. We performed time-resolved ultrafast pump probe UV-vis spectroscopy studies on solution phase $OCP_{CAN}$ as well as microcrystalline OCP suspended in PEG solution ($P2_1$ crystals) and embedded in LCP (C2 crystals, C2c conformer). The solution phase sample was excited with a 540 nm pump; it was contained in a 2 mm cuvette and stirred during the measurement. The crystalline samples were translated during the measurement. Two different LCP samples were investigated at different times, using different setups and slightly different excitations (data reported here utilized peak energy and power densities of ~1.2 mJ/cm$^2$ and ~10 GW/cm$^2$, while the data corresponding to the green line in Supplementary fig. 21A utilized a peak energy and power densities of ~1.4 mJ/cm$^2$ and ~18 GW/cm$^2$), resulting in similar results that confirmed reproducibility.

Supplementary fig. 22 shows stationary spectra of the crystalline samples compared to solution phase OCP. Since crystalline samples show substantial scattering, the scattering-induced baseline was subtracted for the sake of easy comparison. The stationary $S_0 \rightarrow S_2$ absorption band in the PEG sample ($P2_1$ crystals) is redshifted with respect to the one in solution; in the LCP sample (C2 crystals, C2c conformer) the redshift is even larger. The spectral shifts are also manifested in the transient absorption spectra. In particular, the excited state bands of the crystalline samples are significantly redshifted and have different shapes compared to the solution sample. Nevertheless, the crystalline and solution phase samples show comparable dynamics – all bands decay with comparable speed as one can see in Supplementary fig. 23, where the relative magnitude between bands stays similar for different delays. A better way to demonstrate this is by superimposing the kinetics, taking into account the redshift of the bands. Supplementary fig. 21E, F show that a very good overlap can be obtained this way. Therefore, we conclude that the changes in the protein environment are tuning the bands but not altering significantly the excited state relaxation dynamics. Since the relaxation time of the excited population is limited by the $S_0$-$S_1$ gap, it is likely that this gap is not altered, otherwise the kinetics would have changed. Unfortunately, the $S_0 \rightarrow S_1$ band is not visible in carotenoid spectra, since it is a forbidden transition. In conclusion, we did not detect significant differences in the ultrafast photodynamics in solution phase and crystalline ($P2_1$, C2) OCP samples.

**Supplementary text 7: Light-induced untethering of the carotenoid in the C2c state**

Data collected on C2 macrocrystals featuring the C2o state at 100% occupancy show that upon illumination, the SC#1 closes (1 min illumination) (Fig. 3A and Supplementary text 2). However, carotenoid untethering was not observed, *i.e.* the H-bond to Y203 and W290 were intact in the 1 and 10 min. illumination structures (Fig. 3A and Supplementary fig. S8). To investigate the structure after prolonged illumination under steady state conditions we illuminated microcrystals, featuring less unit cells and therefore being both more light-penetrable and potentially more plastic. OCP microcrystals (~7 µm) were transferred to 2.5 cm Petri dishes filled with mother liquor at pH 5.8 and subjected to 48h-illumination with mild blue light, or kept in the dark (Supplementary fig. S20B, C). The 'dark' crystal slurry contained $P2_1$ and C2 microcrystals at a 1:5 ratio (as judged from the indexing of 10,000 vs. 39,000 patterns), with the latter featuring the C2c conformation at 100 % occupancy (Supplementary table 9). Illuminated $P2_1$ crystals were all isomorphous with their dark-state counterparts but in the case of C2 crystals, two populations were present after illumination, only one of which was isomorphous to the dark C2c state while the other was characterized by a 16.6% increase in the unit cell volume. Extrapolation methods were used to extract the steady-state accumulated light-induced contribution from isomorphous dark and light datasets, which afforded to model the $P2_1^{light\text{-}48h}$ and $C2c^{light\text{-}48h}$ structures (Materials and Methods). The non-isomorphous C2 crystals were used to merge a $C2new^{light\text{-}48h}$ dataset, from which a structure was solved using standard crystallographic approaches (see below). In the main text and figures, we mostly plot results for the two 'extreme' steady-state structures that were characterized, *i.e.* the $P2_1^{light\text{-}48h}$ and $C2new^{light\text{-}48h}$ structures, though carotenoid dihedral angle distribution (Supplementary figs. S28 and S29) and H-bonding distances (Fig. 5A) are shown for the $C2c^{light\text{-}48h}$ structure as well. Below, we further develop our findings and rationalize this choice.

In the $P2_1$ crystals, an expansion of the protein is observed upon illumination Supplementary figs S36 and S37), with changes in the overall configuration. Briefly, in the two chains, we observe partial untethering of αA from the β-sheet (R9/R291 H-bond broken in $P2_1^A$ and $P2_1^B$; and T15/K279 H-bonds broken in $P2_1^A$) (Supplementary figs. S31 and S32) as well as from the facing monomer in the dimer (broken $P13^{A/B}/A133^{B/A}$ H-bonds) (Supplementary fig. S33). Complete disruption is seen at the dimerization interface (Supplementary fig. S33) upon release of the αGH loop by loss of the $P13^{A/B}$/A133B/A intermolecular H-bonds. In the $P2_1^B$ chain, we additionally observe rupture of the D35-mediated H-bond to Y129 and S132, at the αC/αGH interface (Supplementary fig. S35). The Q79/A123 H-bond is nevertheless maintained at the αE/αGH interface, in line with the observation of the open conformation of the αGH loop in the two chains (Supplementary fig. S35). Likewise, the αCD loop conformation is overall unchanged (Supplementary figs. S36, S37 and S41). Thus, prolonged illumination of $P2_1$ crystals leads to the accumulation of structural changes in crystalline OCP, notably at the αGH loop, but do not trigger a conformational transition to the closed-conformation –

presumably as a result of crystal lattice constraints (Supplementary fig. S6C). At the carotenoid level, slight differences are seen in the illuminated $P2_1^A$ and $P2_1^B$ carotenoid conformation (Supplementary figs. S25-S29), reflecting differences in the dark-state tunnel architecture (Supplementary figs. S2, S3 and S24). However, the changes are overall minor, with carotenoid dihedral angles varying by less than 30°, in the two chains (Supplementary figs. S25-S29), while H-bonding is preserved to either Y203 ($P2_1^B$) or W290 ($P2_1^A$), mirroring observations in the P21 structure determined at the 1 μs time delay (Figs. 4 and 5A and Supplementary fig. S30).

In stark contrast, the carotenoid is fully untethered from the CTD in the 48h-illumination extrapolated C2c structure ($C2c^{light-48h}$; O/Y203(OH) and O/W290(NE1) distances are 3.4 and 3.8 Å, respectively). The carotenoid conformation is virtually dark-like notwithstanding a bending at C4' , reminiscent of the observation made in the C2 structure determined at the 1 μs time delay (Fig. 6 and Supplementary figs. S28-S29). The latter change results in the O' atom nearing I40(CG1) (-0.7 Å, compared to the C2c dark state). At the protein level, conformational changes are overall minor (Supplementary fig. S39). Notably, no changes are observed in the αGH loop configuration, *i.e.* the protein remains in the C2c conformation and the Q79/A123 H-bond broken, while interactions are preserved at the interfaces with αC (N32/D35 and D35/S132 H-bonds) and αE (Q79/I71 and Q79/M74 H-bonds). The αCD loop also retains the dark-state configuration, although the Y44/E115 H-bond double-locking SC#2 in the C2c dark-structure is now water-mediated. The dimerization interface does not change (T17/N134, T17/D19 and D19/R27 H-bonds are all maintained), and interdomain interactions are for the most part preserved (the T52/P278, T52/W279, A58/V178, N104/W279, Q149/A209 and R155/E246 H-bonds are all formed). Thus, the 48h-illumination extrapolated C2c-structure differs from the dark C2c conformer mainly by the loss of H-bonding between the carotenoid and the protein, and the change in puckering state of the C2 atom (Fig. 5A).

Extrapolation methods could not be utilized for data collected on the non-isomorphous illuminated C2 crystals, characterized by a 16% increase in unit cell volume (Supplementary table 9). Instead, a new model was phased and iteratively built and refined based on the raw electron density map. We refer to this dataset and structure as $C2new^{light-48h}$. In the final $C2new^{light-48h}$ structure, the carotenoid is observed untethered from both Y203(OH) (3.5 Å) and W290(NE1) (4.9 Å), echoing the observations made on the isomorphous $C2c^{light-48h}$ structure. Nevertheless, the carotenoid structure differs largely in the two structures. Notably, large twists are seen at C6-C7, C7=C8 and C11=C12 (Δ~75°, 120° and 65°, respectively; Fig. 6 and Supplementary fig. S28-S29 and S38), yielding another 7-*cis* isomer of canthaxanthin, distinct from both the 7-*cis* isomer observed in the $P21^A$ and $P21^B$ structures determined at the 10 ps time delay differ and the 7-*cis* isomer observed in the C2c structure determined at the 1 μs time delay. This dramatic change in the polyene structure results in the C9 methyl fitting into a grove lined by L250, V275, T277, M286, and M288 – that is, the same place that it occupied in the C2c structure determined at the 1 µs time delay as opposed to being located above the M286 and M288 side chains as in the dark C2c and C2c$^{light-48h}$ structure. W290 and Y203 both undergo substantial displacement upon loss of their H-bond to the carotenoid, but it is notable that the Y203(OH) accompanies the β1 carbonyl oxygen (O) as it slides towards I305 (distance to CD1 is 3.2 Å) The carotenoid is thus in very different configurations in the C2c$^{light-48h}$ and C2new$^{light-48h}$ structures (Supplementary figs. S28-S29), though they share the common feature of the rupture of H-bonds to Y203 and W290 (Fig. 5A). The part of the carotenoid accommodated in the NTD only shows twists at C15'-C14' and C7'-C6', and a slightly increased twist at C10'=C9' (Fig6 and Supplementary figs. S28-S29 and S38). Nevertheless, the C2new$^{light-48h}$ structure shows substantial rearrangements within the NTD. Notably, αG has moved inward toward αF and αE, producing a local compaction, whereas αD is shifted outward relative to αC and the remainder of the αDEFG bundle (Supplementary fig. S39). The reason for such large conformational changes in αC, αD, αE and the αCD loop is the main feature of the C2new$^{light-48h}$ structure, that is, the reopening of the carotenoid tunnel at SC#1 upon complete restoration the open state conformation of the αGH loop and re-folding of the αCD loop (Fig. 6 and Supplementary figs. S34, S5, S39 and S41). The re-open αGH loop conformation, which features virtually no electron density for I125, is stabilized by the Q79/A123 H-bond, but the D35 mediated H-bonds to Y129 and S132 are broken (Supplementary fig. 35). Strikingly, however, we observe the re-setting of the P13/A133 H-bond, contrasting with all C2 structures determined thus far (Supplementary fig. S33). Accompanying changes include substantial displacements of the αG, αC, and αI helices, which predominantly contribute to the carotenoid tunnel wall (Supplementary fig. S39). Notably, in αC, large rotamer changes are observed in W41, whose aromatic ring serves as a "sliding base" for the αGH loop, and L45, which maintains W41 in position; in Y44, which due to positional changes in the carotenoid atoms undergoes a ~30° change in χ1 that results both in a change of the I51 rotamer and H-bonding of Y44(OH) to E115 (different rotamer than in the C2c or C2o structures) and Y111; in M 47, whose side chain has moved away from the carotenoid upon changes in Y44 and I51, resulting in the loss of the M47(O)/T50(OG1) H-bond (Supplementary fig. 41). The loss of this H-bond is concomitant with the unwinding the C-terminus of αC and a pull of the αCD loop, which only remains anchored to the CTD via the T52/W279 H-bond – *i.e.*, both the T52/P278 H-bond (β5-β6 turn) and the A58/V178 (linker) H-bonds are broken. Of note is that the αD helix is also largely displaced, moving towards the position formerly occupied by the linker (notably N60 and F63) and resultingly sliding over αG and αF (Supplementary fig. S39). Comparatively large positional and/or rotameric changes occur in other tunnel-lining residues, viz. L37, L107, L154, R155, V158, in the NTD, and A224, L225, F243 L250, V275, W279, M288 and I305, in the CTD. In this context it must be noted that L107 shows very poor electron density and a 0.3 Å increase in distance between its CD2 and the carotenoid C12'.

The C2new$^{light-48h}$ structure is also characterized by an increased opening angle between the two domains (~ +4°). Accordingly, of six interdomain tethers observed in the dark C2o and C2c structures, only the N104(OD1)—W279(NE1) and T52(N)/W279(O) H-bonds are preserved in the C2new$^{light-48h}$ structure – *i.e.,* the T52(OG1)/P278(O), Q149(ND2)/A309(O) and A58/V178 (linker) H-bonds are broken, and likewise the R155—E246 salt-bridge. Interestingly, we see N156(OD1) acting as a compensatory H-bonding partner for R155 (3.2 Å distance to NH2), with the resulting N156 conformation itself stabilized by a water bridge to Q230. Nevertheless, we observe conservation of all interactions at the dimerization interface. At the surface of the β-sheet, αN remains overall well anchored to the β-sheet and αA, with conservation of the N14(O)/N316(ND2), L16(N)/E313(OE1), L16(O)/A309(N) and A18(N)/L307(O) H-bonds. Only the H-bond from N316 to K279 breaks, which partly liberates αN from the β-sheet and further reduces the coupling with αA upon rupture of the K279(NZ) H-bridge to T15(OG1). At the αA/β6/β7 interface, we concomitantly see rupture of the R9(O)/R291(NH1) and R291(NH2)/D306(OD1) H-bonds, as R291 "slides" over the D306 carboxylate (Supplementary fig. S32).

**Supplementary Text 8: ComPASS analysis of correlated light-triggered structural changes**

The networks of communication pathways in the three chains were extracted based on dynamical correlations and interaction propensities using ComPASS[22] (Materials and Methods). These per-chain analyses of the covariance of conformational changes point to a similar transduction pathway of the excitation energy in the three chains, “trailing” from the β-sheet to the dimerization interface to the αBCHI and then αDEGF bundles of the NTD (Fig. 5C). Accordingly, in the three chains, the main outcome of photoexcitation are large conformational changes in the αCD and αGH loops, triggered as early as 0.3 ps and persisting up to 1 µs (Figs. 4, 5, 6 and Supplementary figs. 25-41). Thus, as already noted, at short, as well as at longer time delays, the two loops connecting the four-helix bundles of the NTD are the main ‘recipients’ of the energy transferred from the carotenoid to the protein scaffold. The efficiency of photo-excitation signal propagation to the αGH loop is higher in the $P2_1^B$ and C2 chains than in the $P2_1^A$ chain, presumed to represent an intermediate between the two formers (Fig. 5C). The ComPASS analysis was crucial to systematically identify the key communication routes that underly the structural response to photoexcitation. By integrating dynamical and interaction-based data into a network framework, thereby going beyond one to one structural comparison between successive time delays, the ComPASS analysis revealed correlated changes across specific residues and structural elements, providing mechanistic insight into long-range allosteric effects (Fig. 5C, Materials and Methods).

Irrespective of the time delay, the effect of the starting ground state ($P2_1^A$ vs $P2_1^B$ vs (C2o/C2c vs) C2c) (Supplementary figs. S1-S3 and S24) was evident on both the carotenoid (Supplementary figs. S25-S28 and S38) and the protein structures (Fig. 5B and Supplementary figs. S31-S37 and S39-S40). As a proxy to judge of the appearance and accumulation of a photoproductive state – *i.e.*, a state *en route* to $OCP^R$, we used the criterion of carotenoid H-bonding to the protein, given the known requirement that the two H-bonds break for OCP to display a red spectrum[23–25] (Fig. 5A). Indeed, the changes at the carotenoid level (Supplementary figs. S25-S27 and S38) associate with different patterns of H-bond rupture to the protein (Figs 4, 5A, 6 and Supplementary fig. S30). In the two $P2_1$ chains, the H-bonds to Y203 break at 0.3 ps and persists as such until 10 ps, in the $P2_1^B$ chain, and 100 ps, in $P2_1^A$. The H-bond to W290 breaks as well at 10 ps but reforms by 100 ps. At longer time delays, the carotenoid is seen to attach by at least one H-bond to either Y203 or W290 (Fig. 4 and Supplementary figs. S25-S26 and S30). Thus, in the two $P2_1$ chains, complete untethering of the carotenoid, proposed to correspond to the formation of the $P_1$ photoproduct, lasts for about 10 ps, which is evidently too short for the carotenoid to exit the tunnel. We see in both chains that as of 1 µs, only one H-bond is set between the carotenoid and the protein, suggesting that the energy barrier to fully untether the carotenoid is significantly and durably lowered upon absorption of a first photon. In the C2c chain, rupture of the H-bond to W290 occurs at 10 ps (*i.e.,* as in the $P2_1$ chains) and remains broken until 1 µs. This is followed by the complementary rupture of the H-bond to Y203, observed at

10 ns, which remains broken at 1 μs. Thus, the carotenoid remains fully untethered for at least 1 μs in the C2 chain, which is five decades of time longer than observed in the $P2_1$ chains, and would be compatible with progression of the carotenoid along the tunnel – at least to the extent that is compatible with the crystal lattice constraints as seen in the $C2new^{light\text{-}48h}$ structure.
