## Supplementary material for "Structural basis of the two-photon photoactivation mechanism of orange carotenoid protein": All supplementary figures

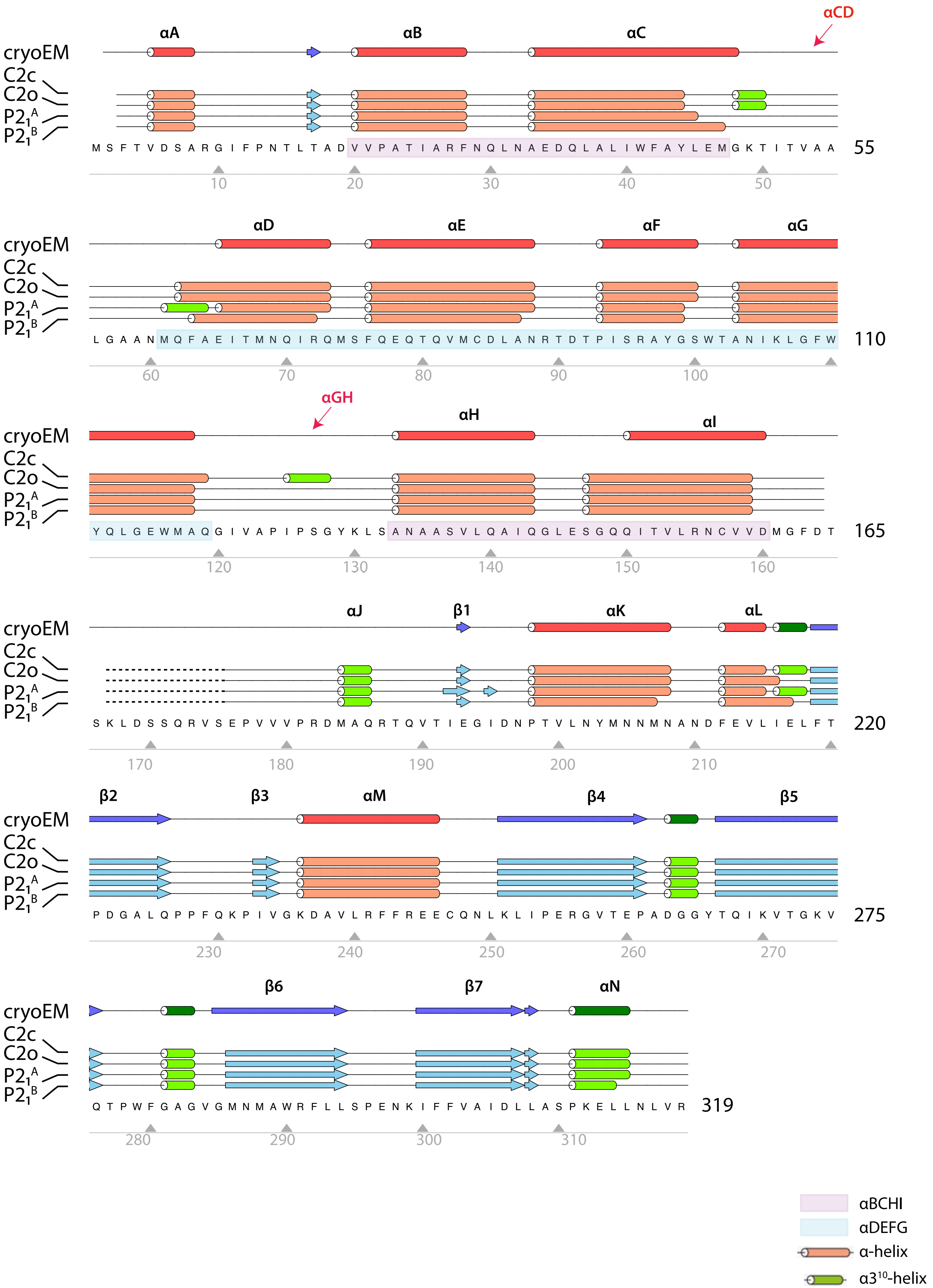


Supplementary Fig. S1

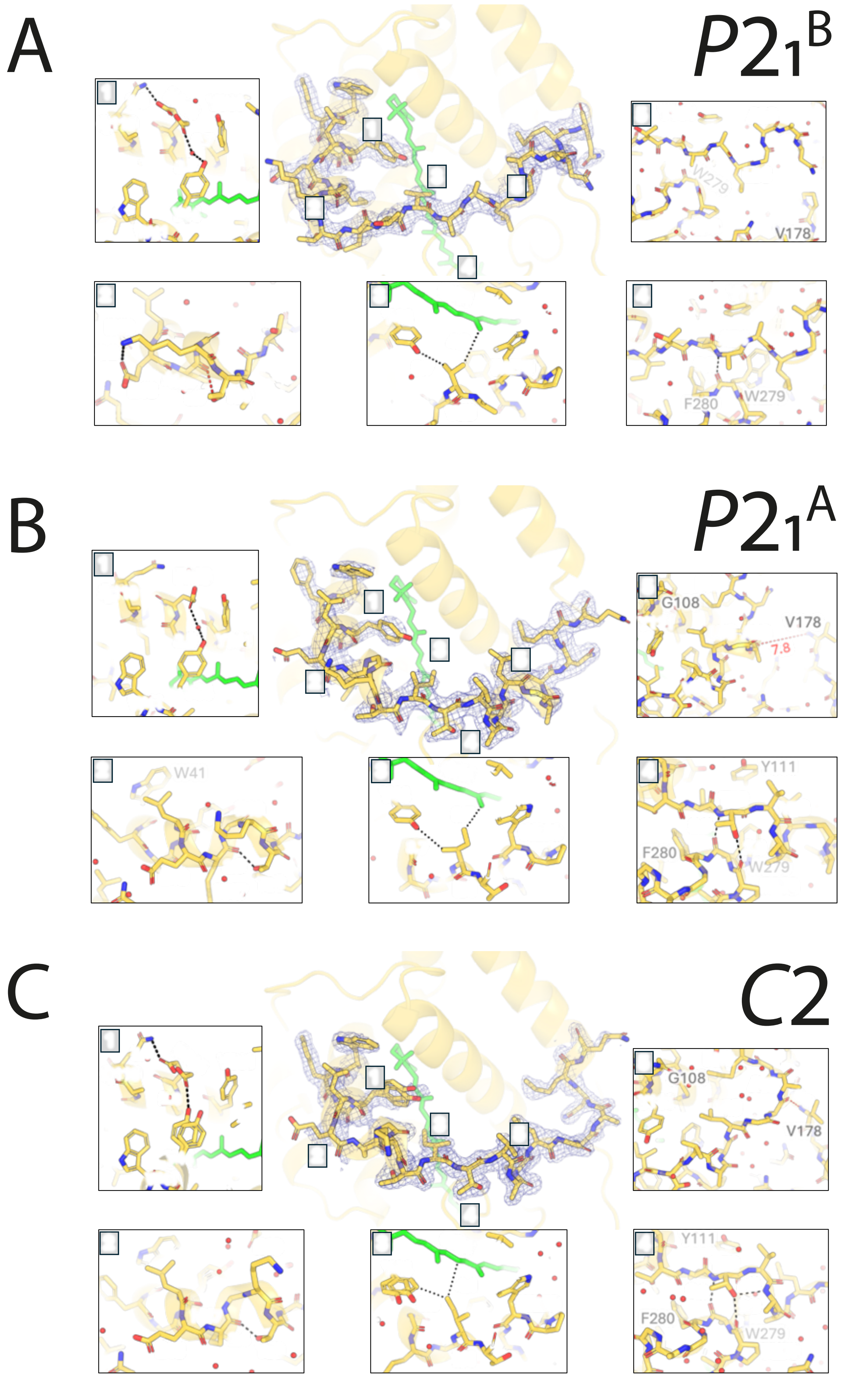


Supplementary Fig. S2

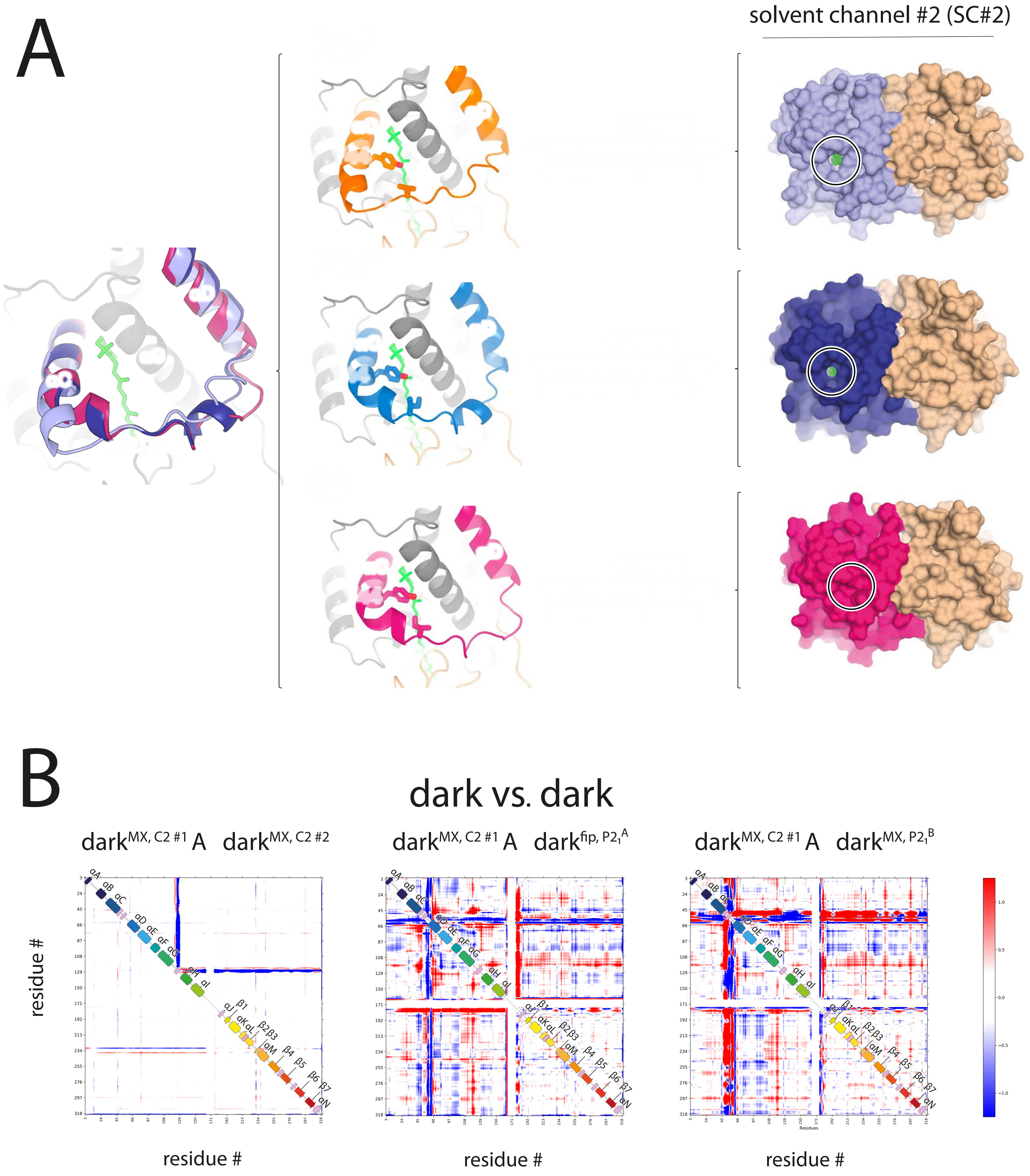


Supplementary Fig. S3

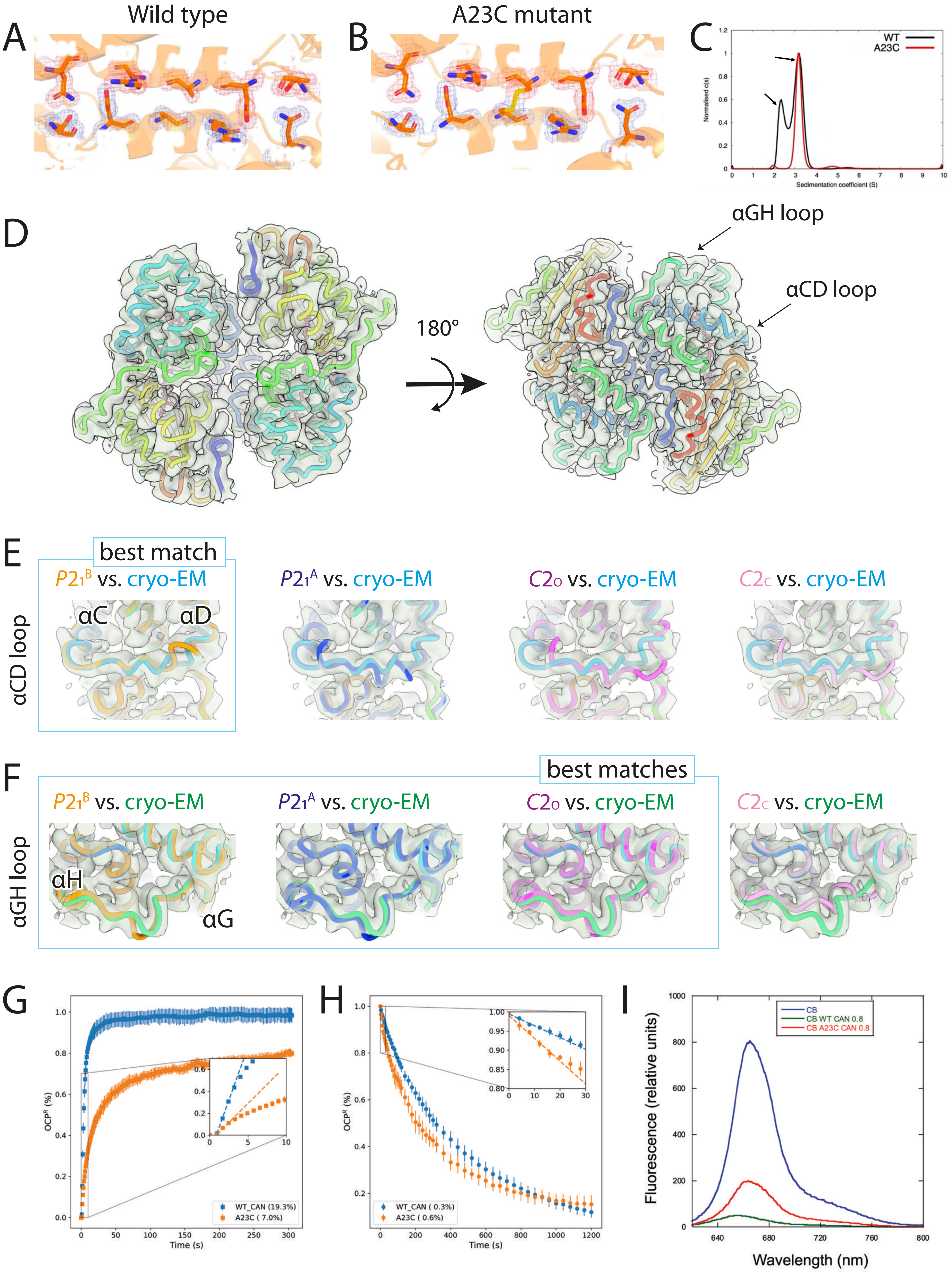


Supplementary Fig. S4

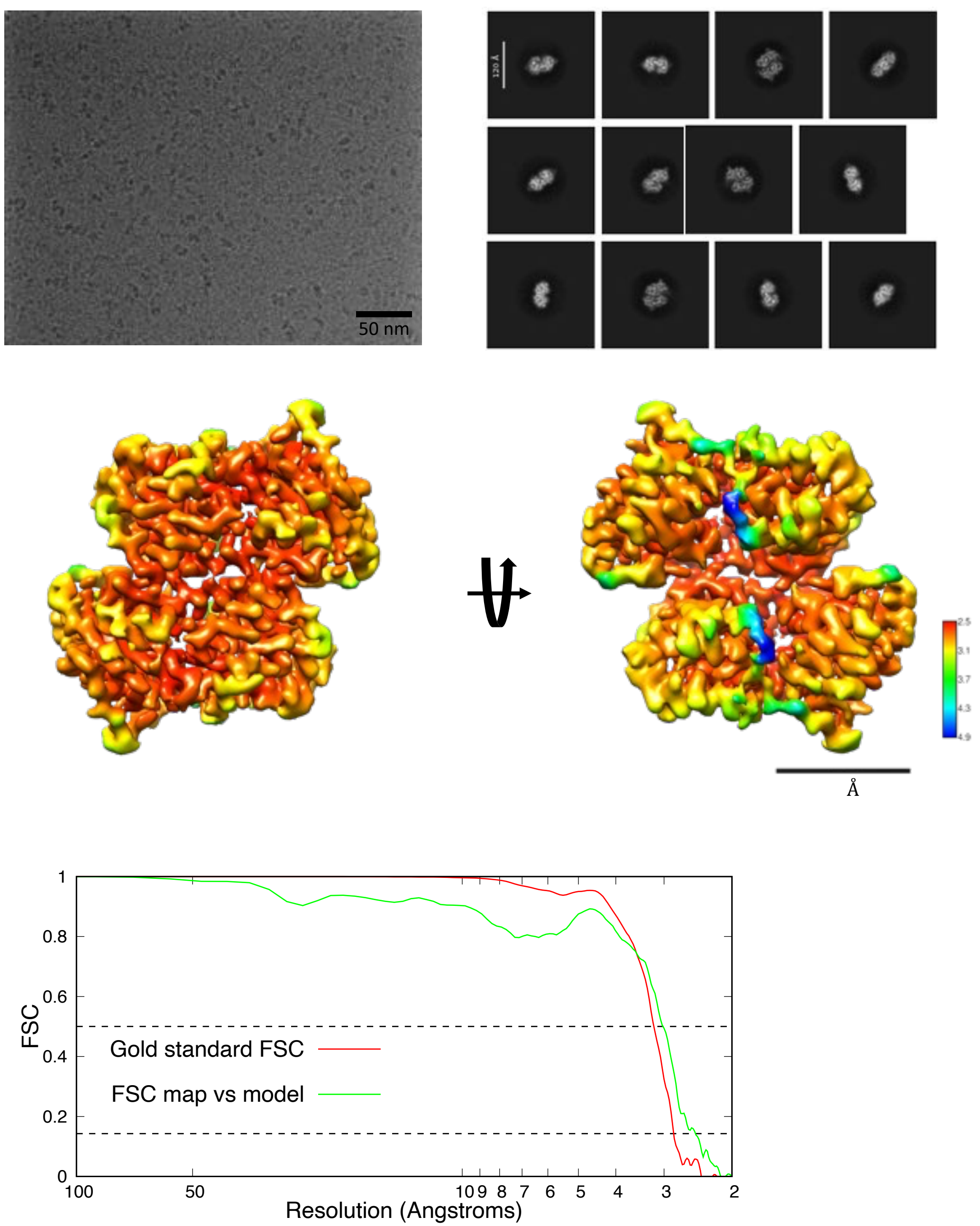
50 nm
2.5
3.1
3.7
4.3
4.9
Å
1
0.8
0.6
0.4
0.2
0
FSC
Gold standard FSC
FSC map vs model
100
50
10 9
8
7
6
5
4
3
2
Resolution (Angstroms)

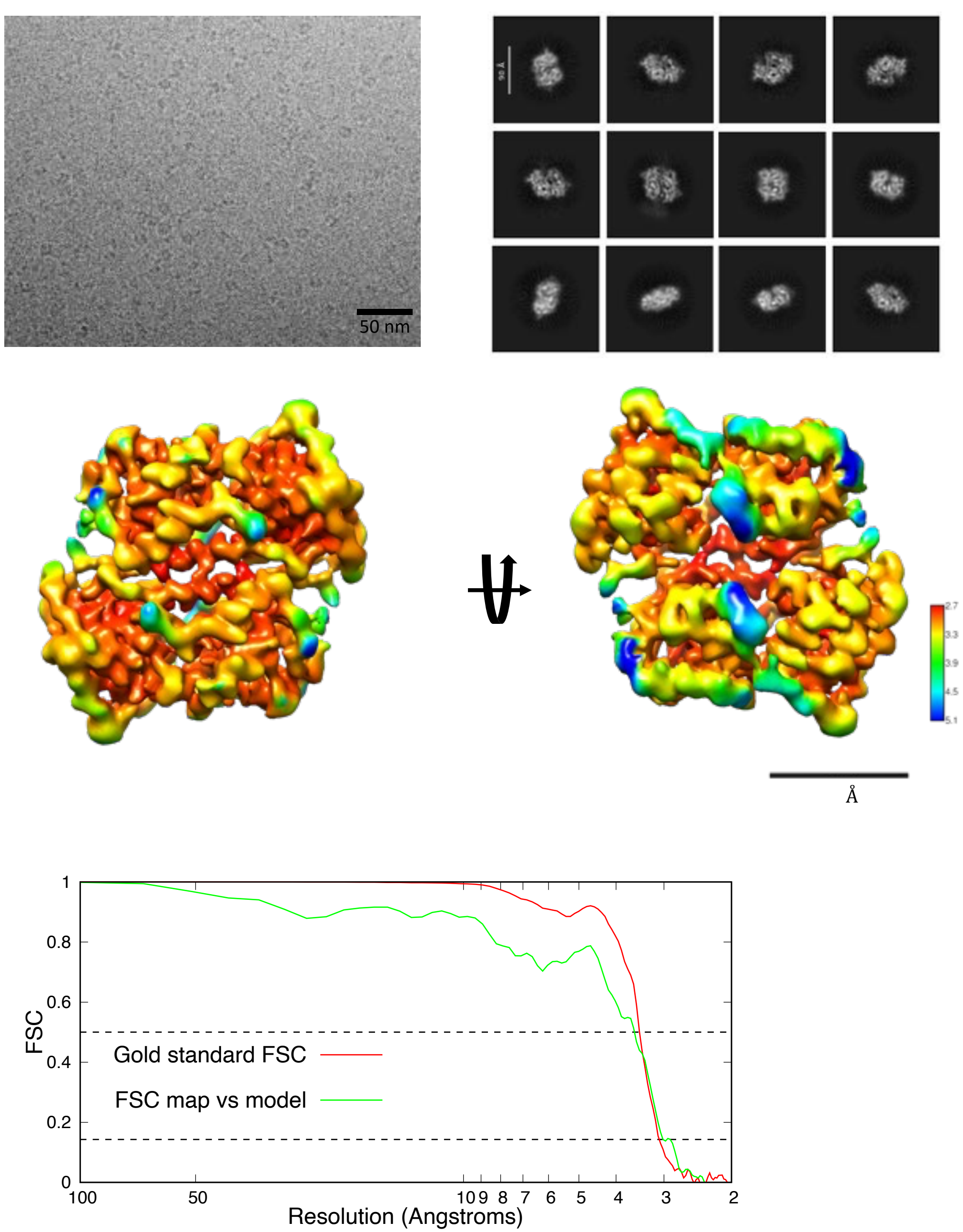
50 nm
2.7
3.3
3.9
4.5
5.1
Å
1
0.8
0.6
0.4
0.2
0
FSC
Gold standard FSC
FSC map vs model
100
50
10 9
8
7
6
5
4
3
2
Resolution (Angstroms)

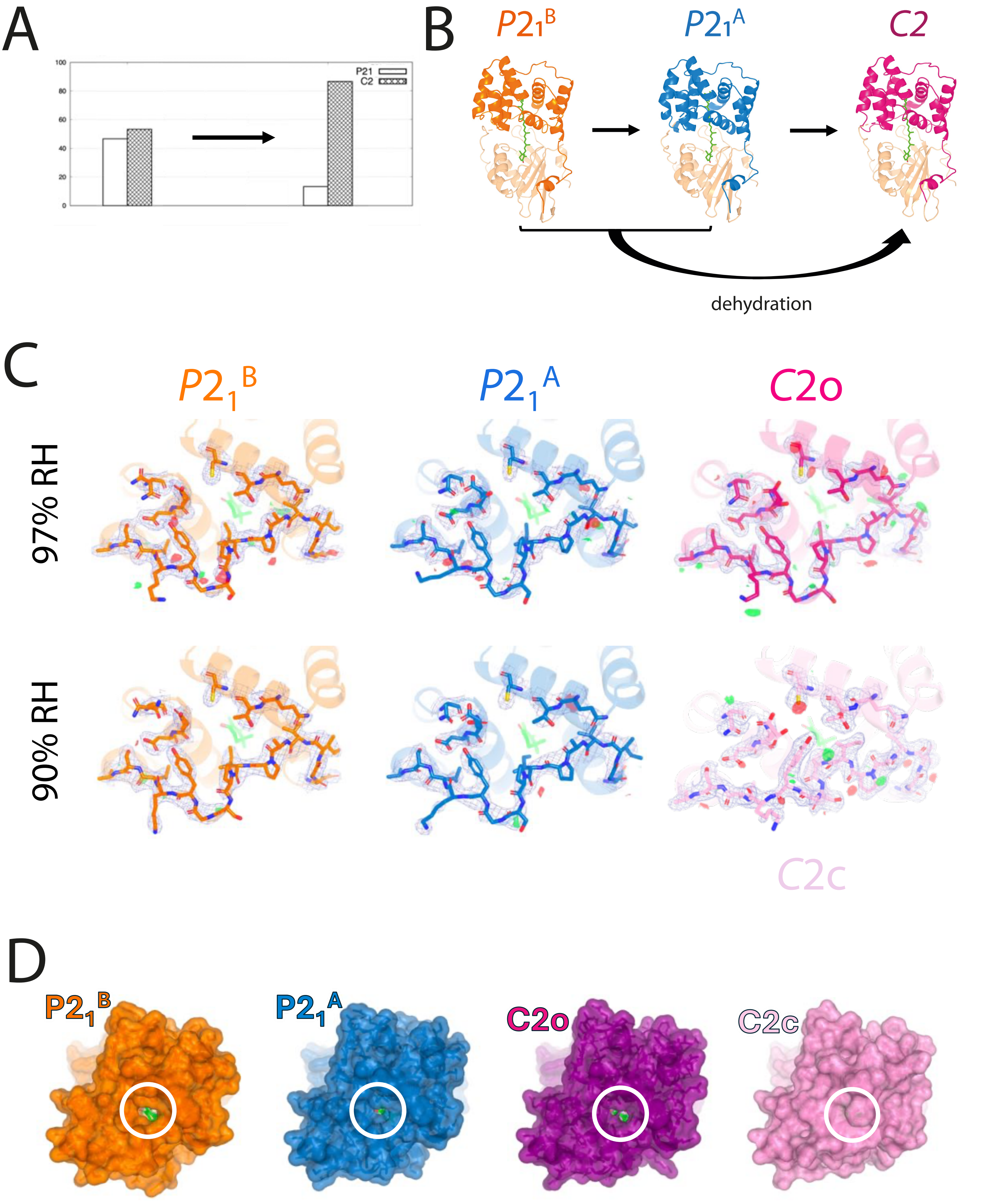


Supplementary Fig. S6

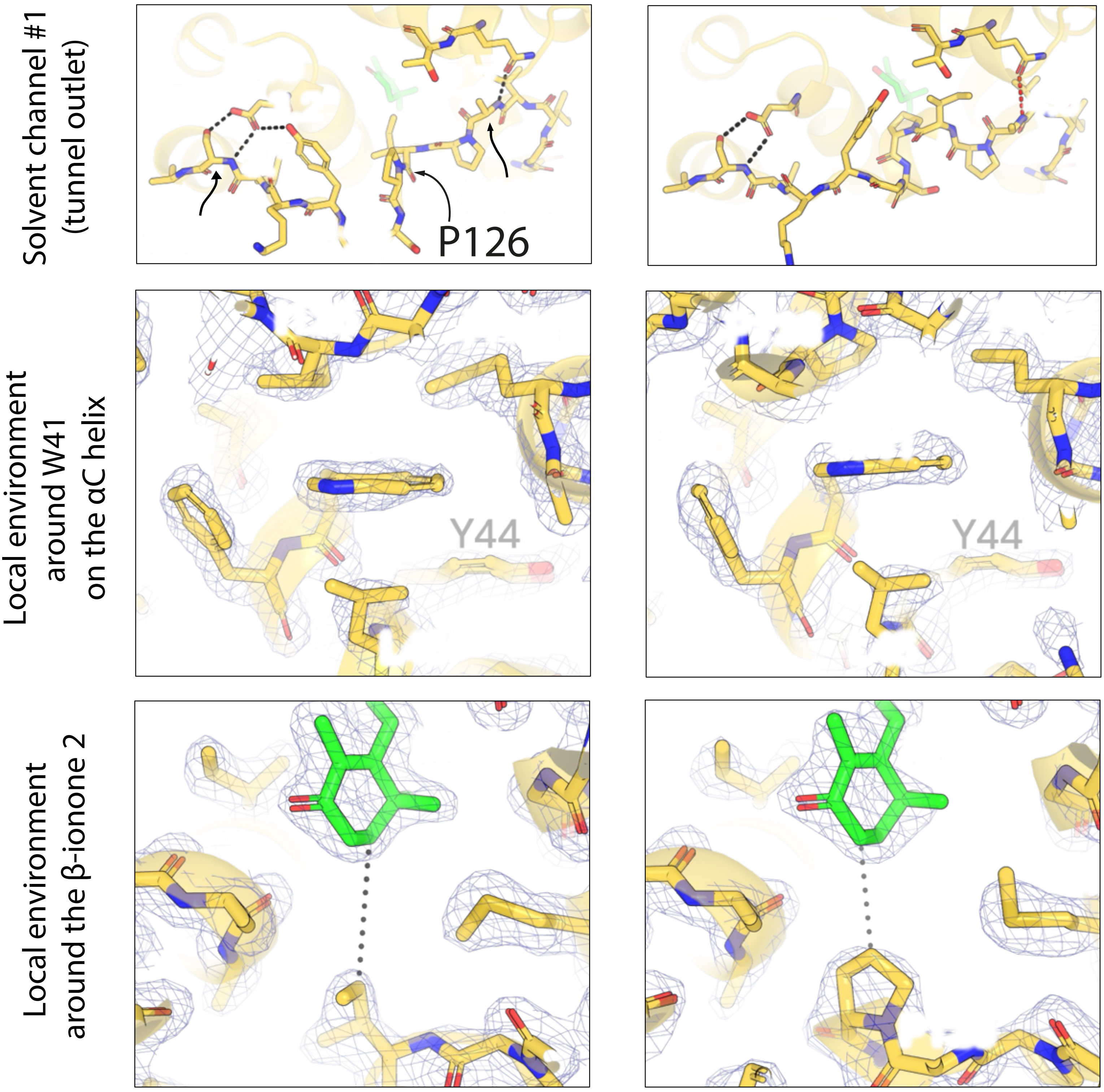

Solvent channel #1 (tunnel outlet)
P126
Local environment around W41 on the αC helix
Y44
Y44
Local environment around the β-ionone 2

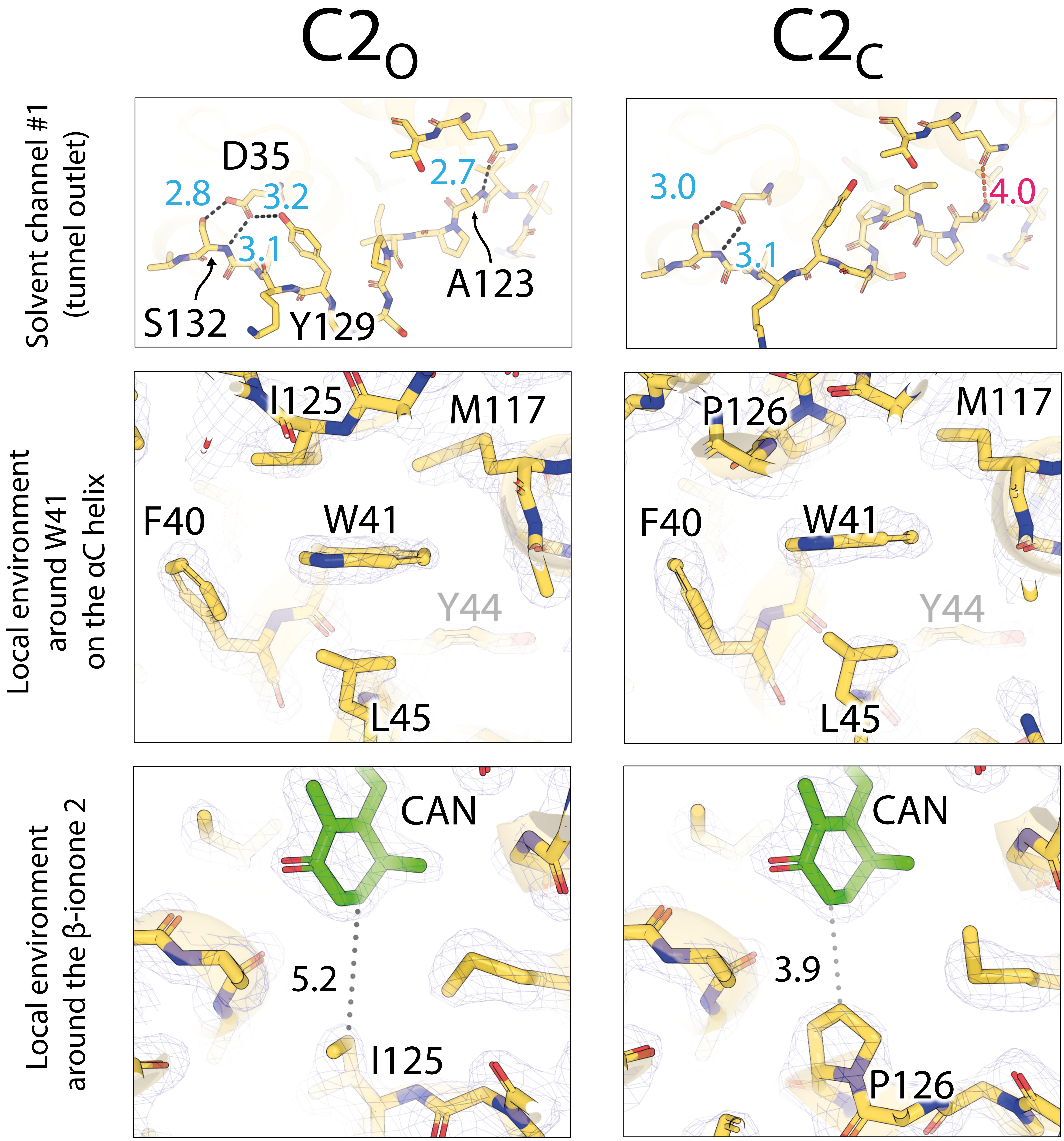


Supplementary Fig. S7

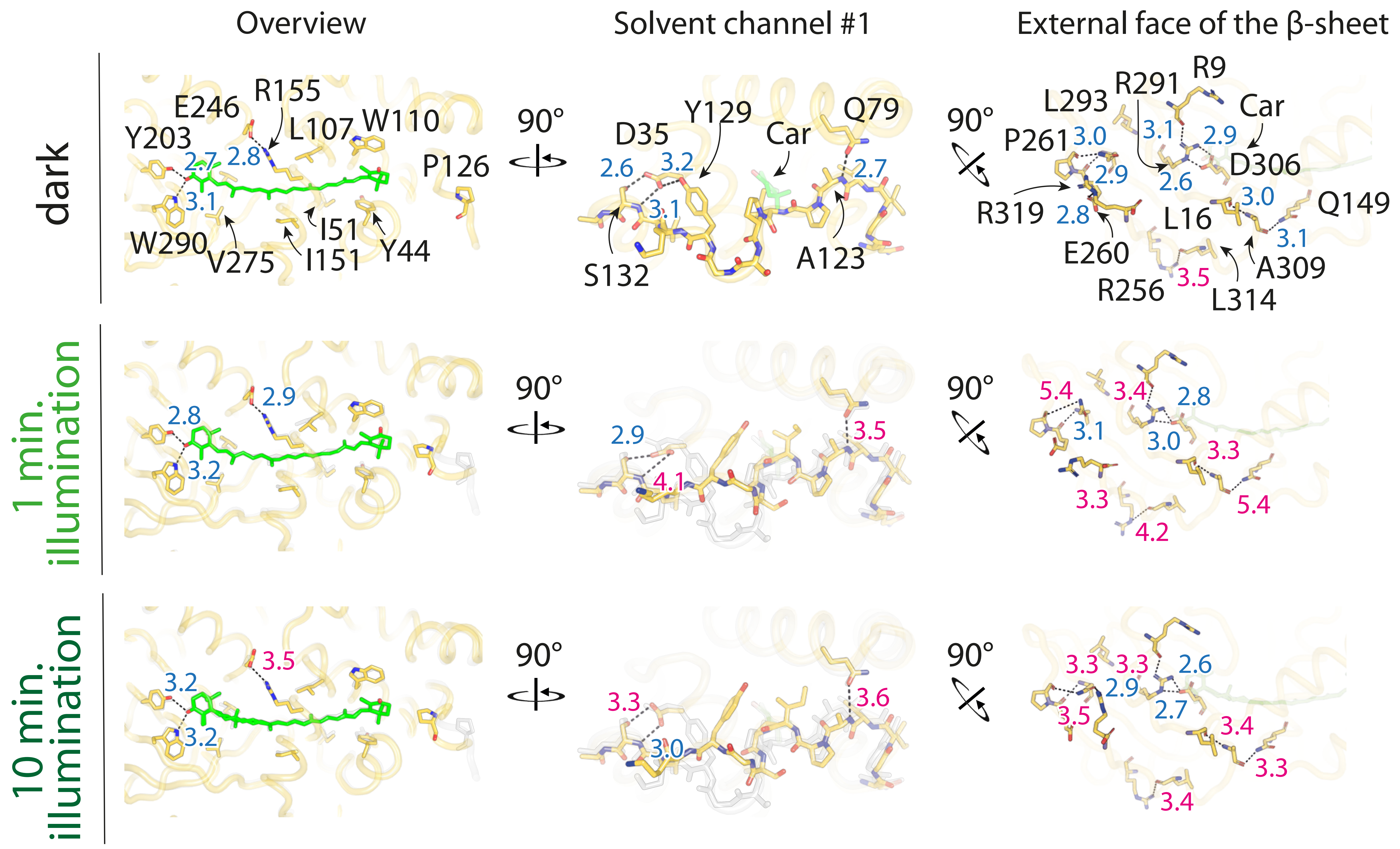


Supplementary Fig. S8

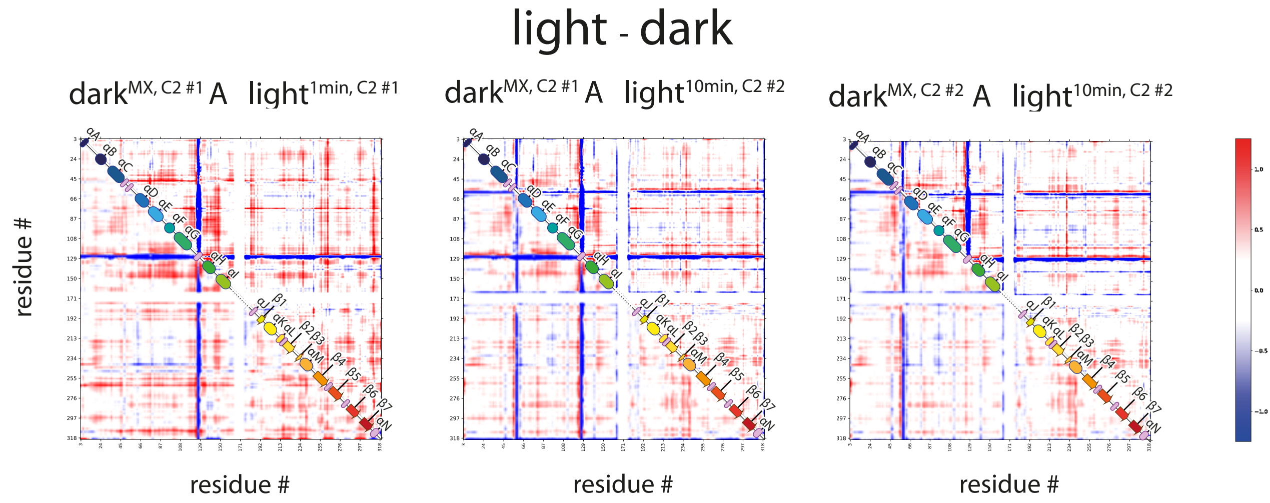


Supplementary Fig. S9

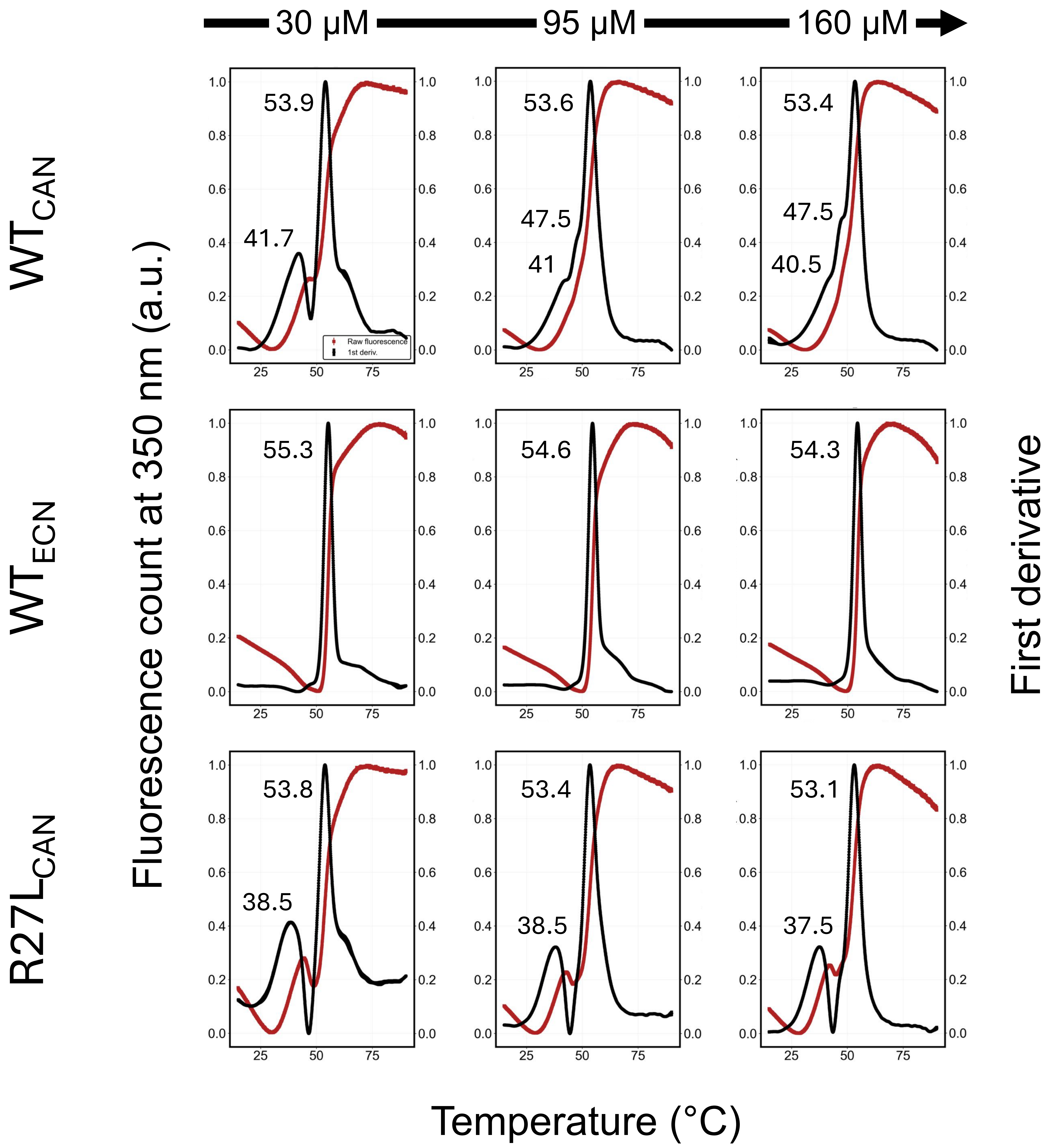


Supplementary Fig. 10

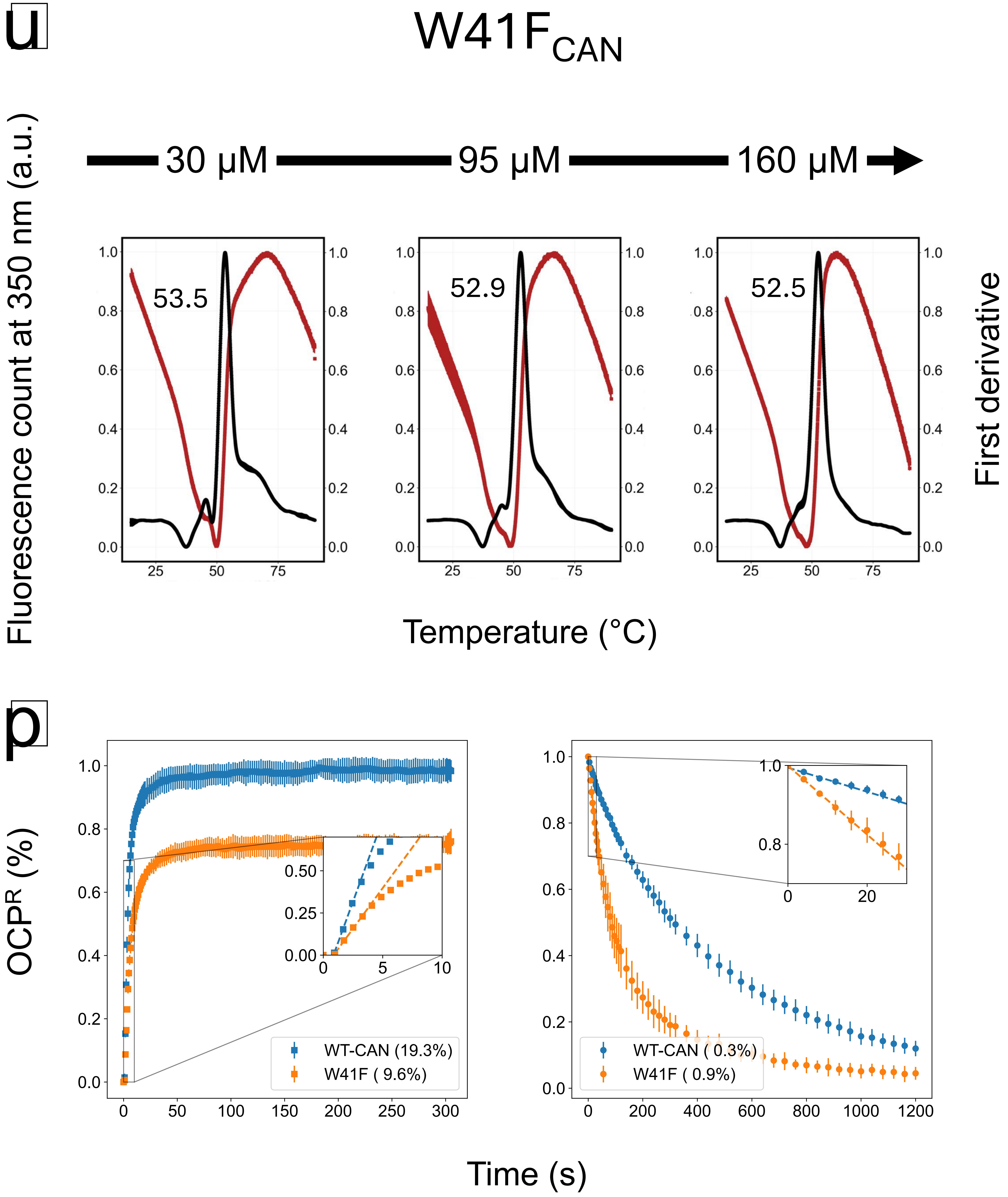


Supplementary Fig. 11

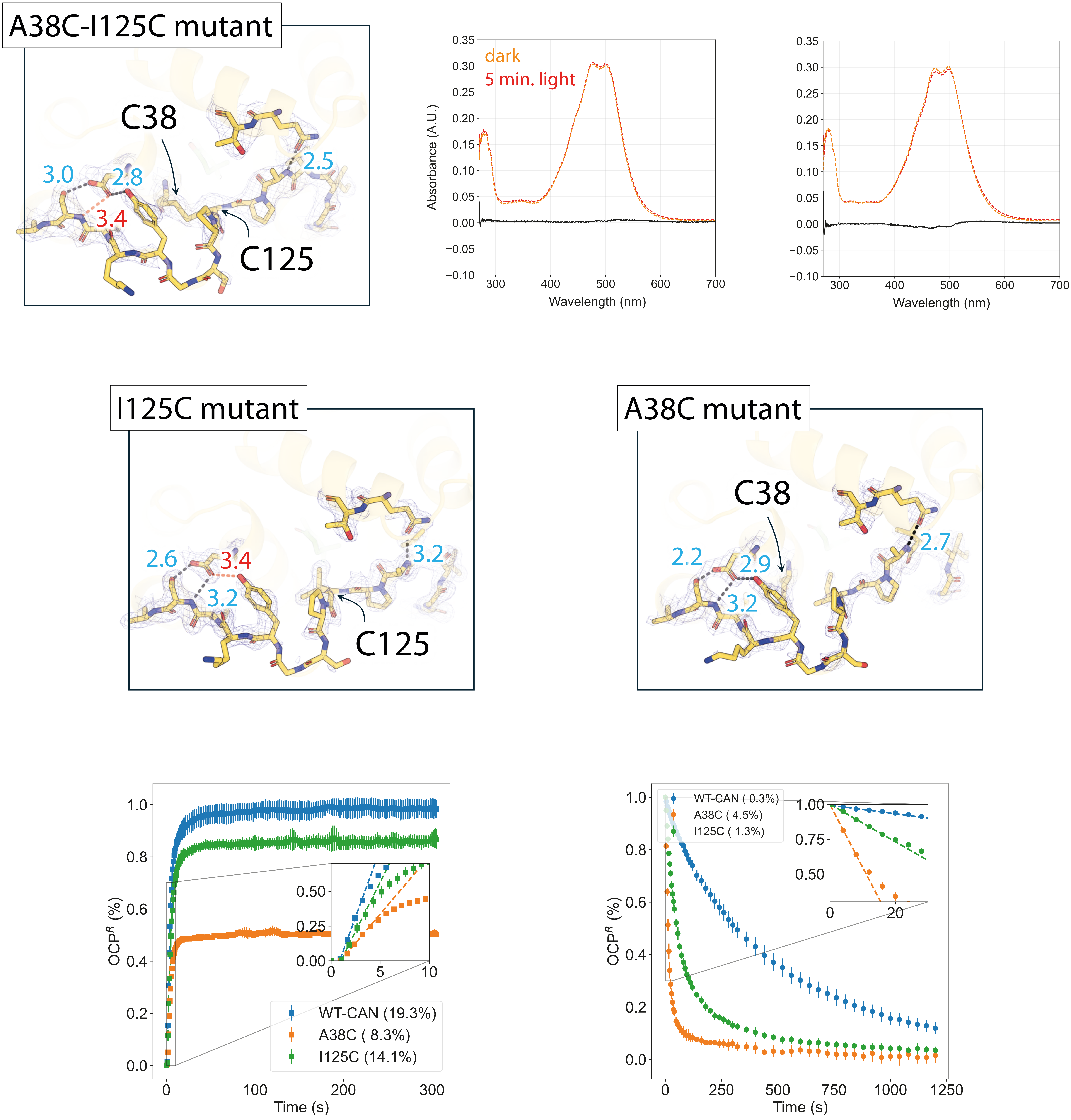


Supplementary Fig. S12

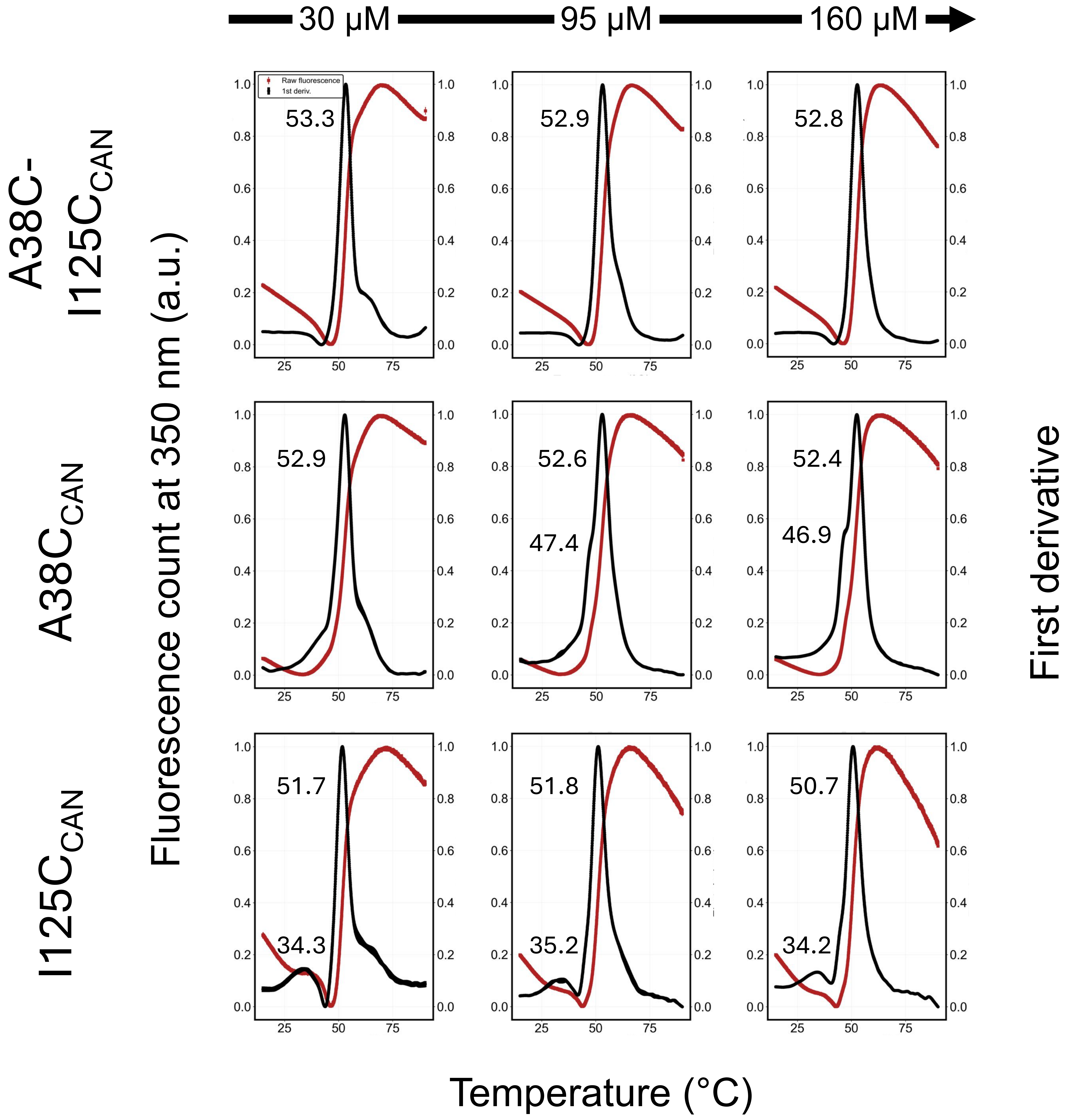


Supplementary Fig. 1

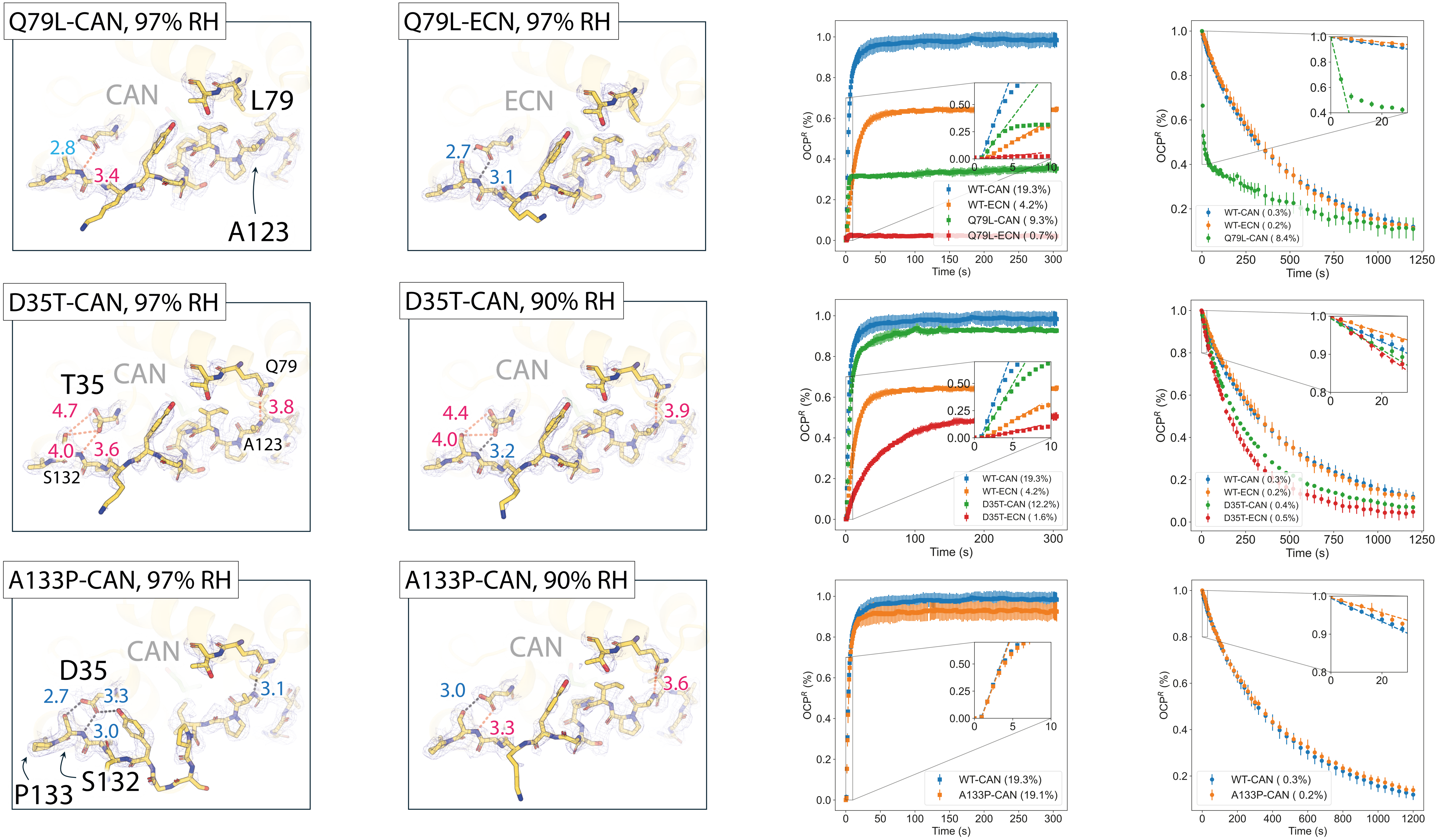


Supplementary Fig. S14

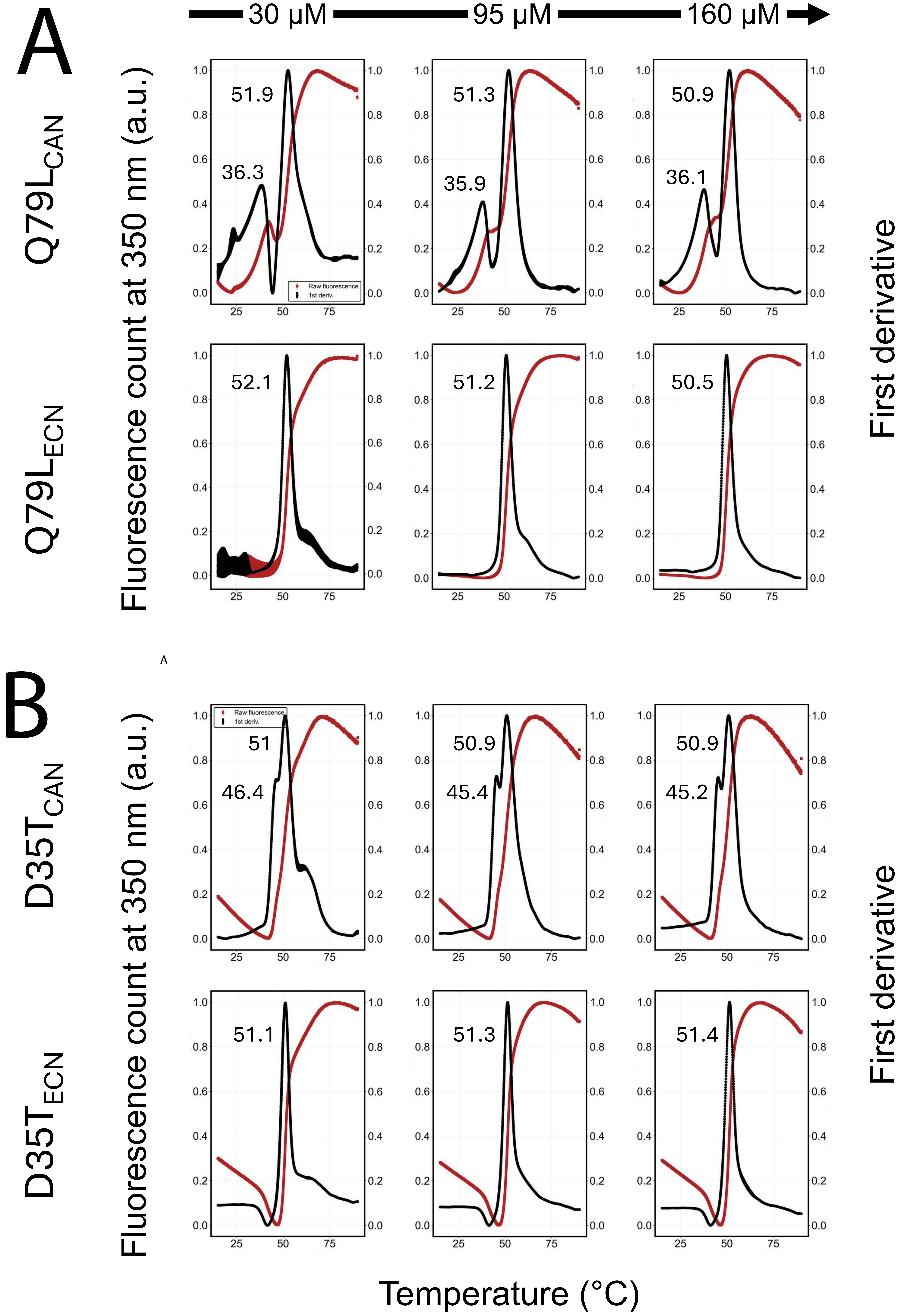

A
30 µM
95 µM
160 µM
Q79L$_{CAN}$
Q79L$_{ECN}$
Fluorescence count at 350 nm (a.u.)
First derivative
51.9
36.3
51.3
35.9
50.9
36.1
52.1
51.2
50.5
Raw fluorescence
1st deriv.
B
D35T$_{CAN}$
D35T$_{ECN}$
51
46.4
50.9
45.4
50.9
45.2
51.1
51.3
51.4
Temperature (°C)

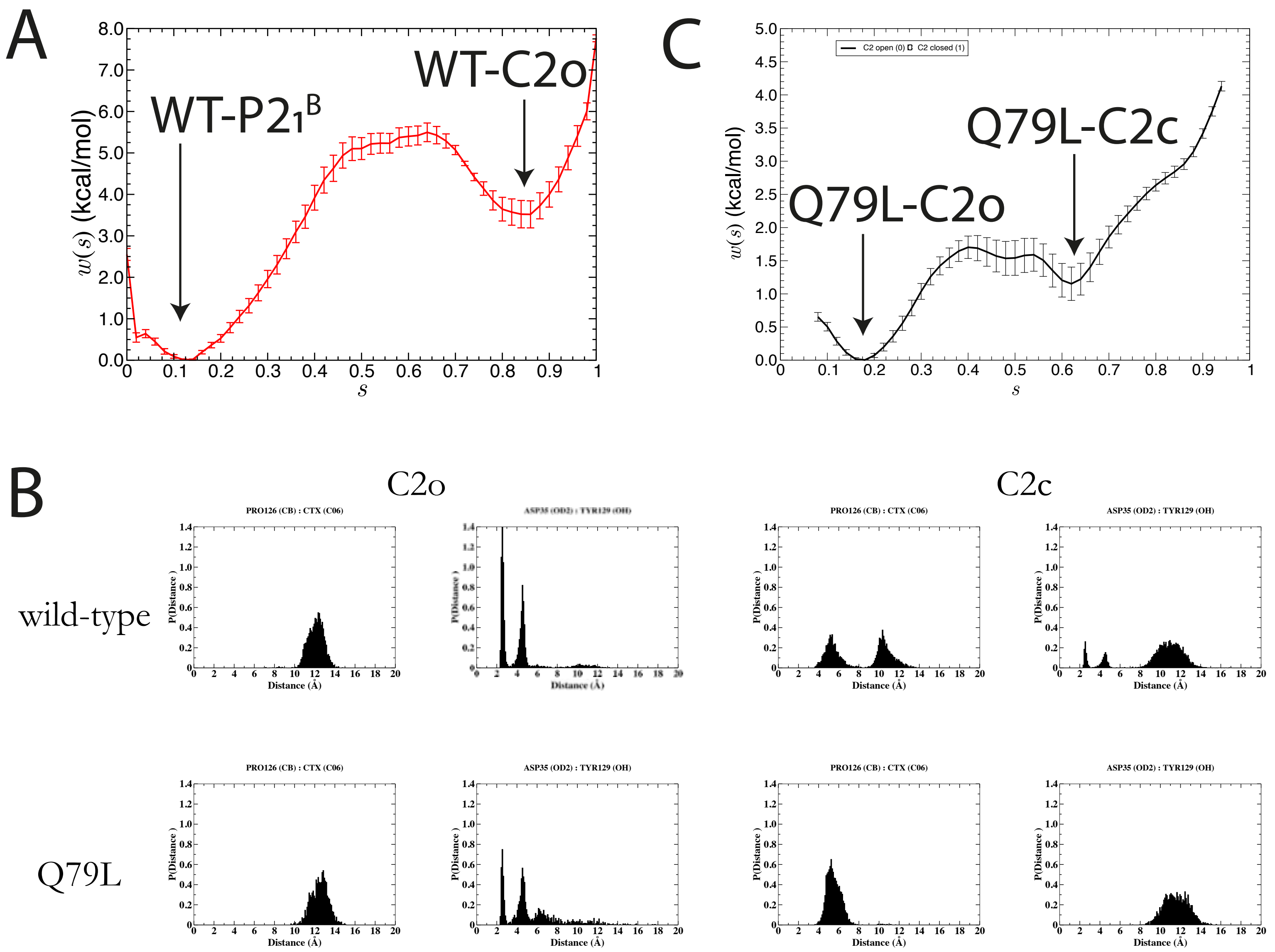


Supplementary Fig. S16

MD simulations starting from
OCP-**WT** in the **C2c monomeric state**

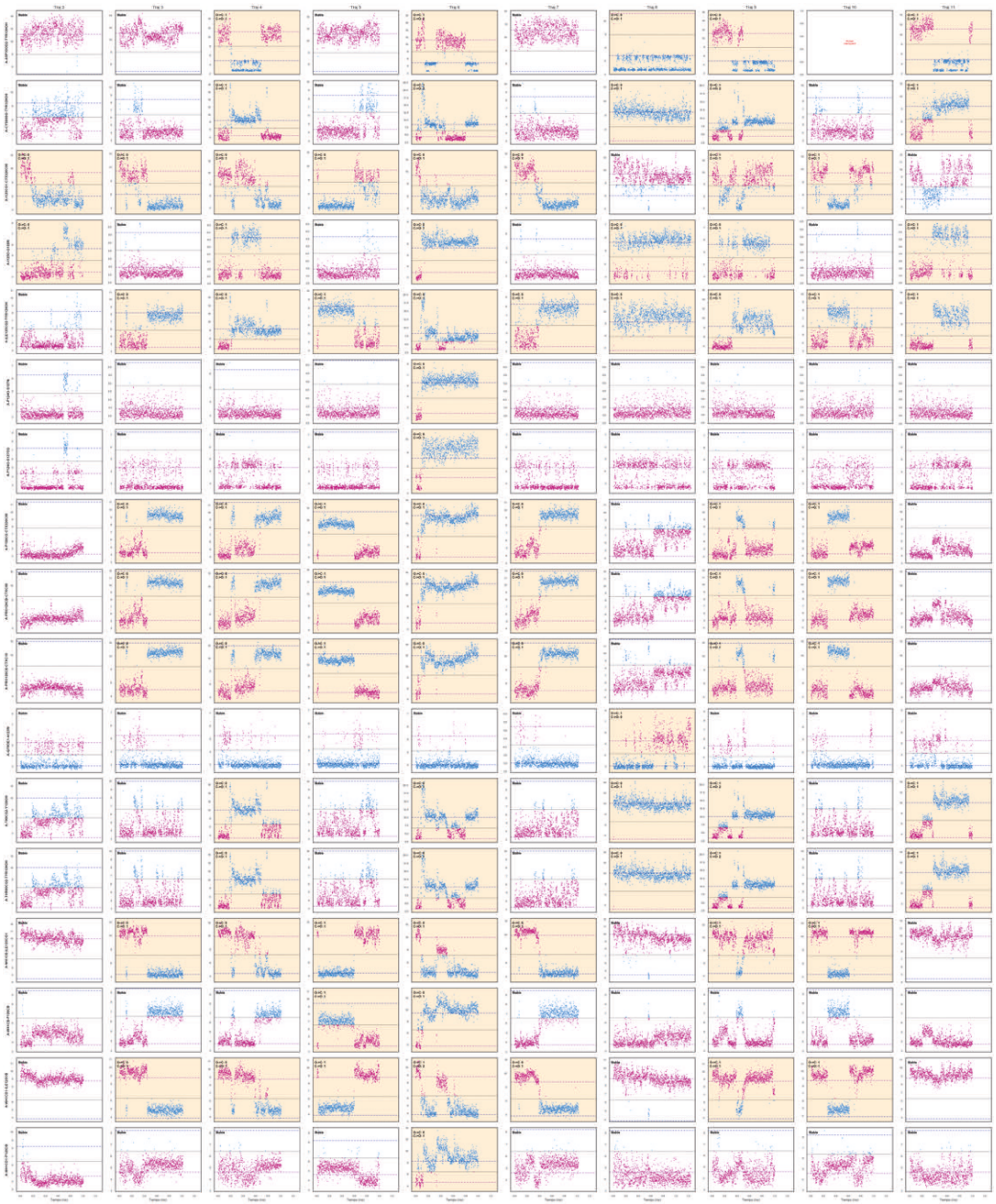

MD simulations starting from
OCP-**WT** in the **C2o monomeric state**

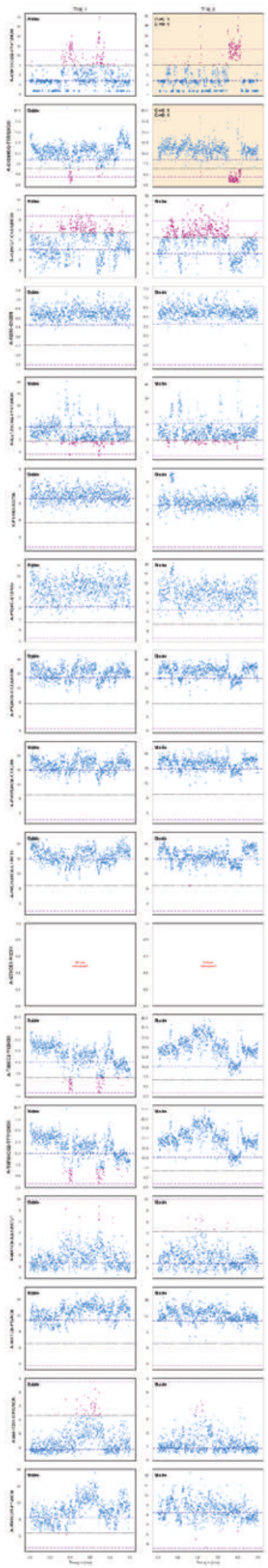

Supplementary Fig. S17

MD simulations starting from
OCP-**Q79L** in the **C2c monomeric state**

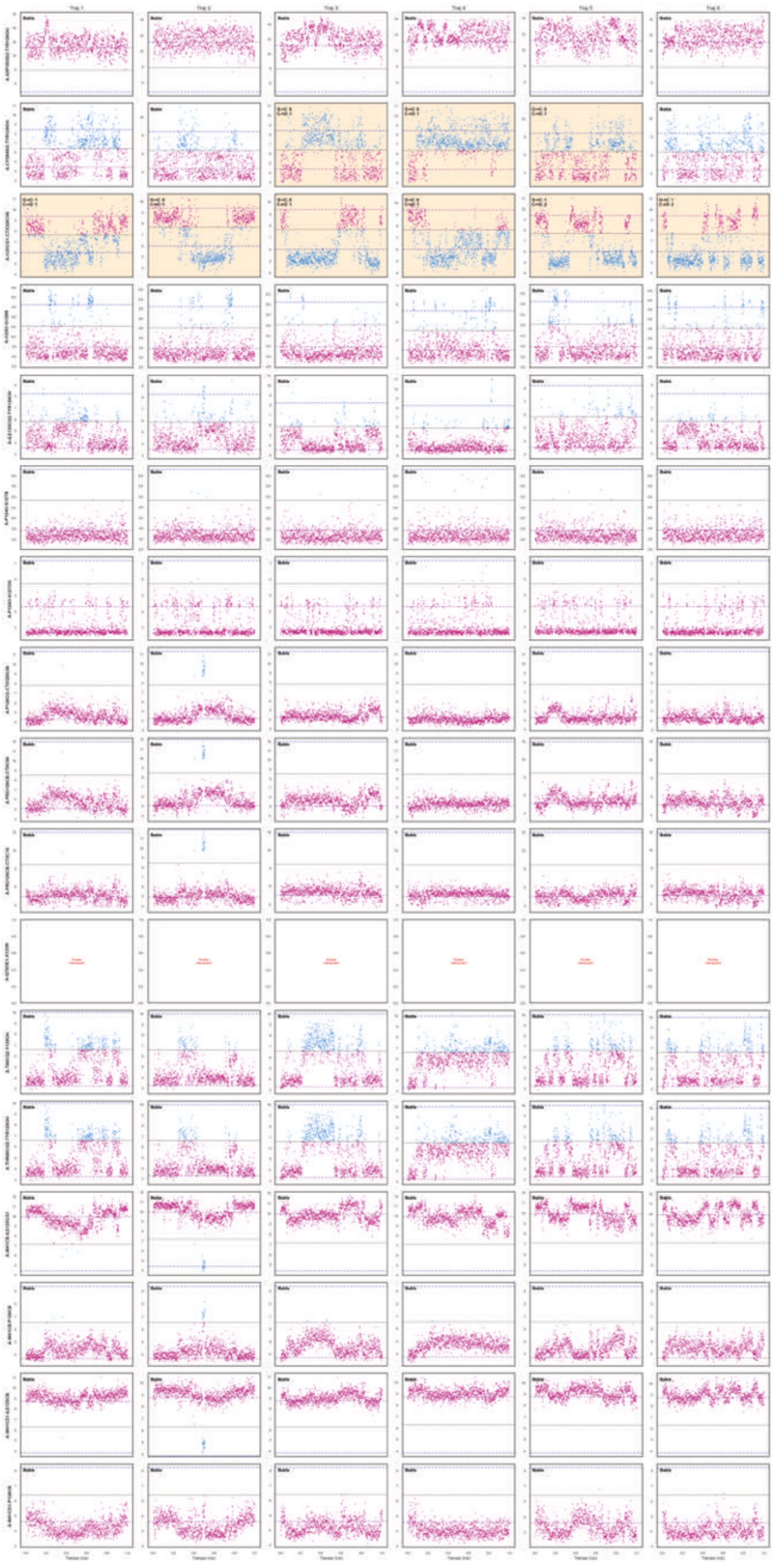

MD simulations starting from
OCP-**Q79L** in the **C2o monomeric state**

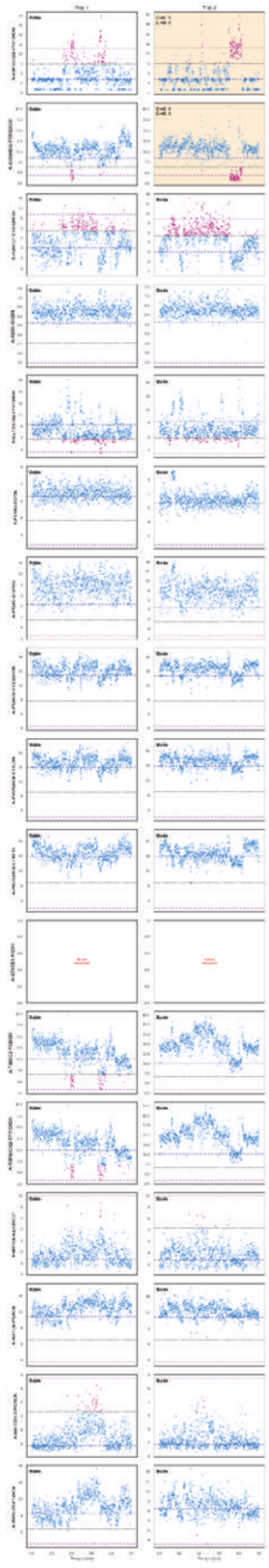

Supplementary Fig. S18

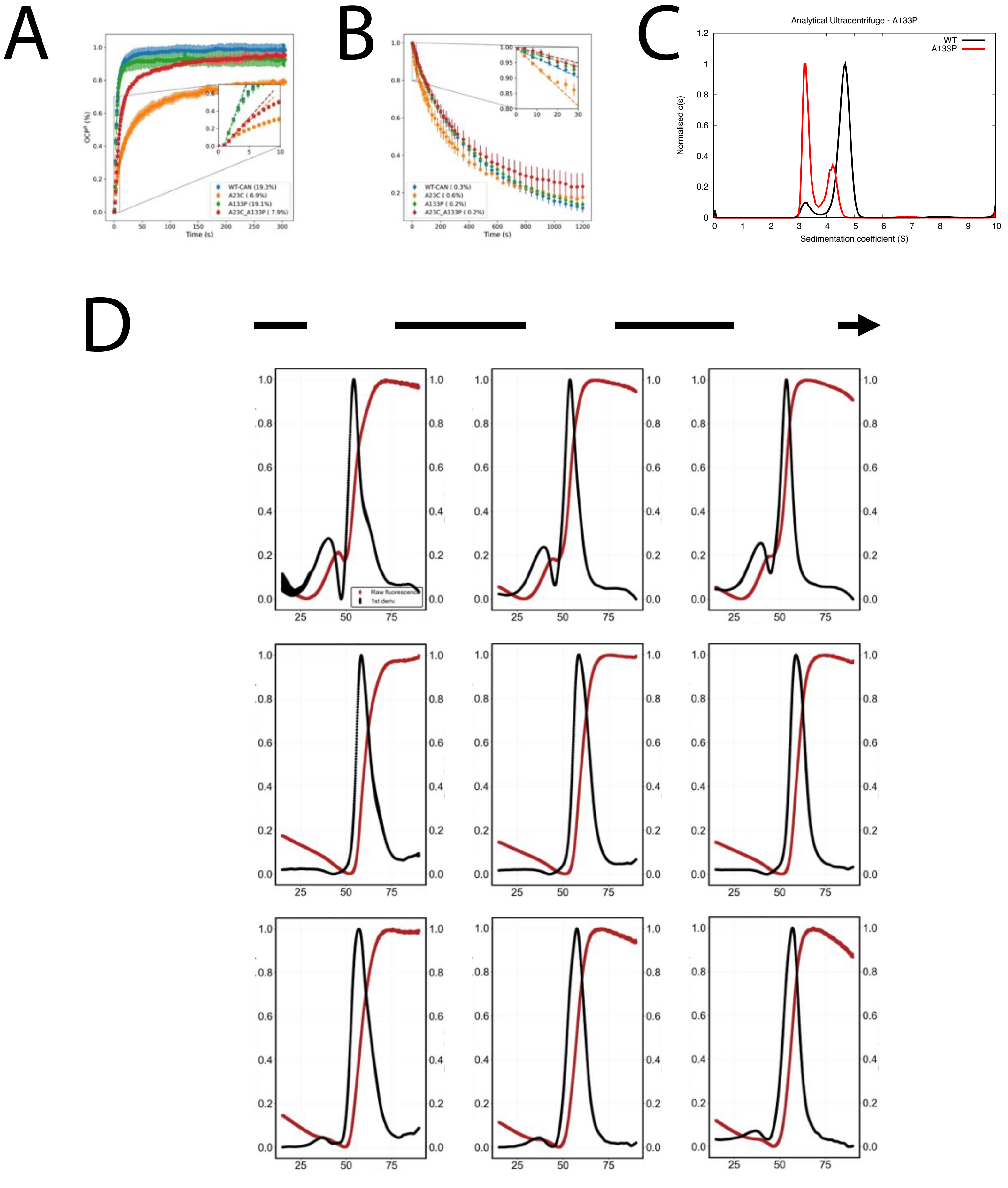


Supplementary Fig. 19

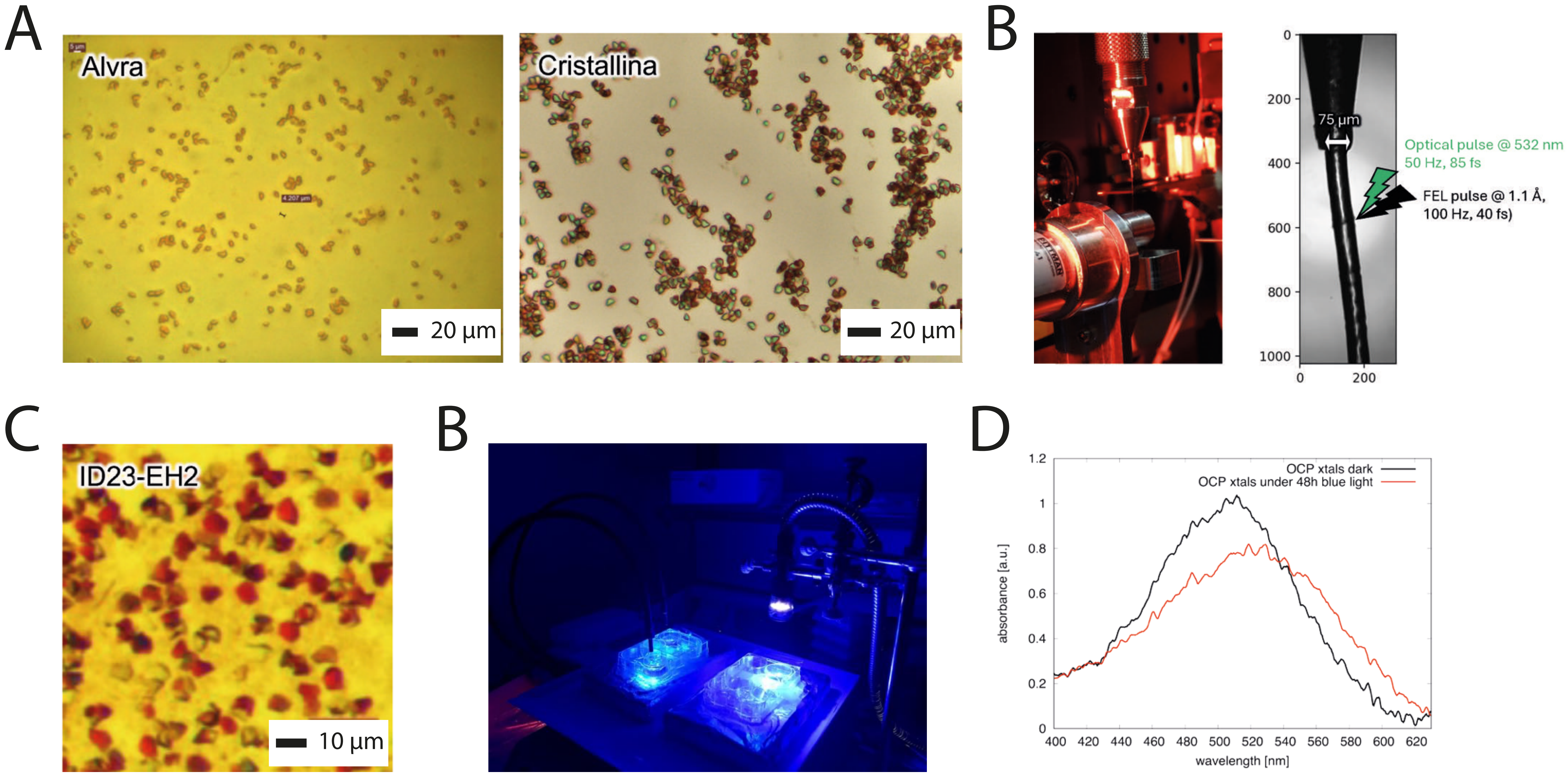


Supplementary Fig. 20

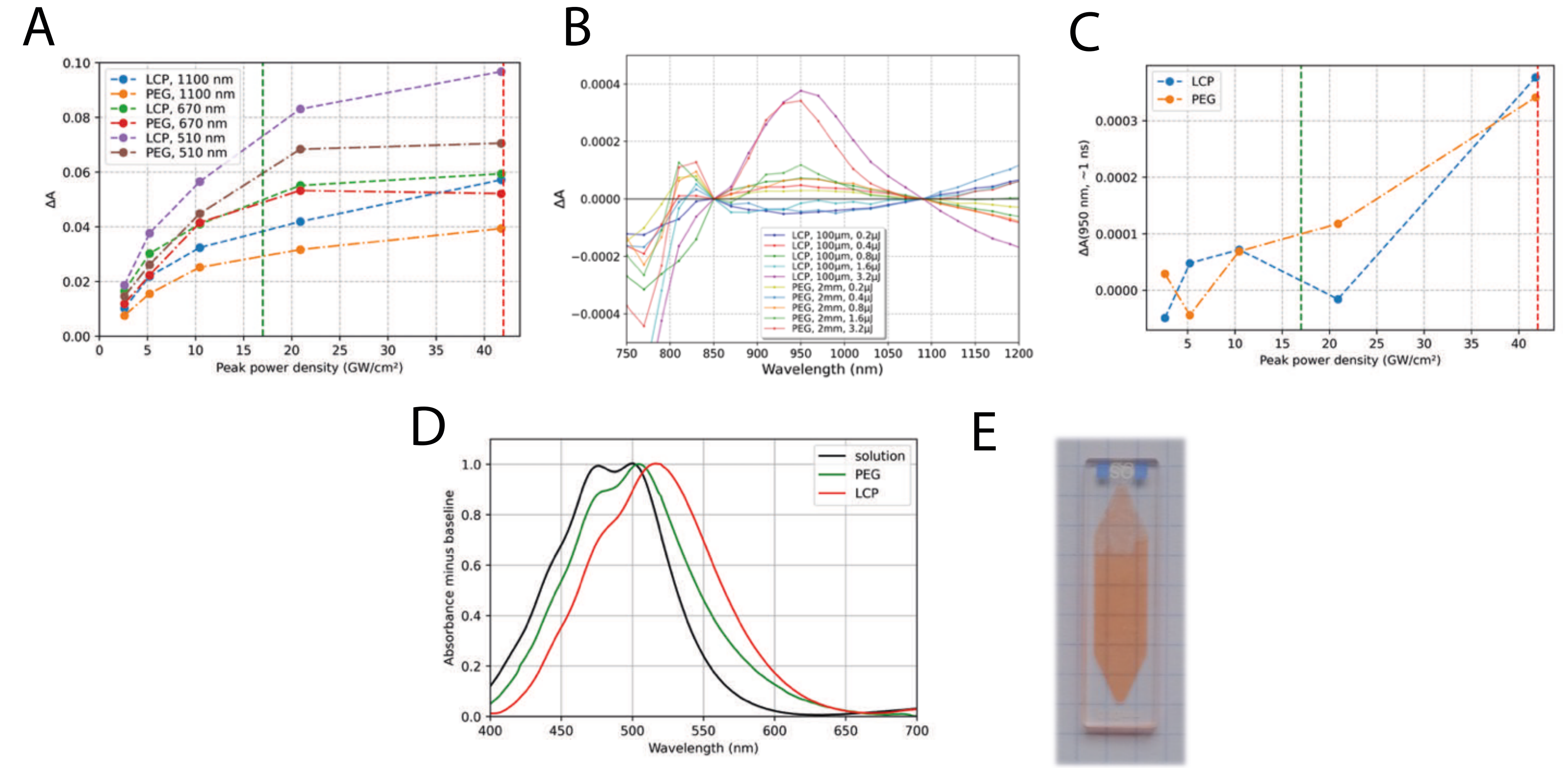


Supplementary Fig. 21

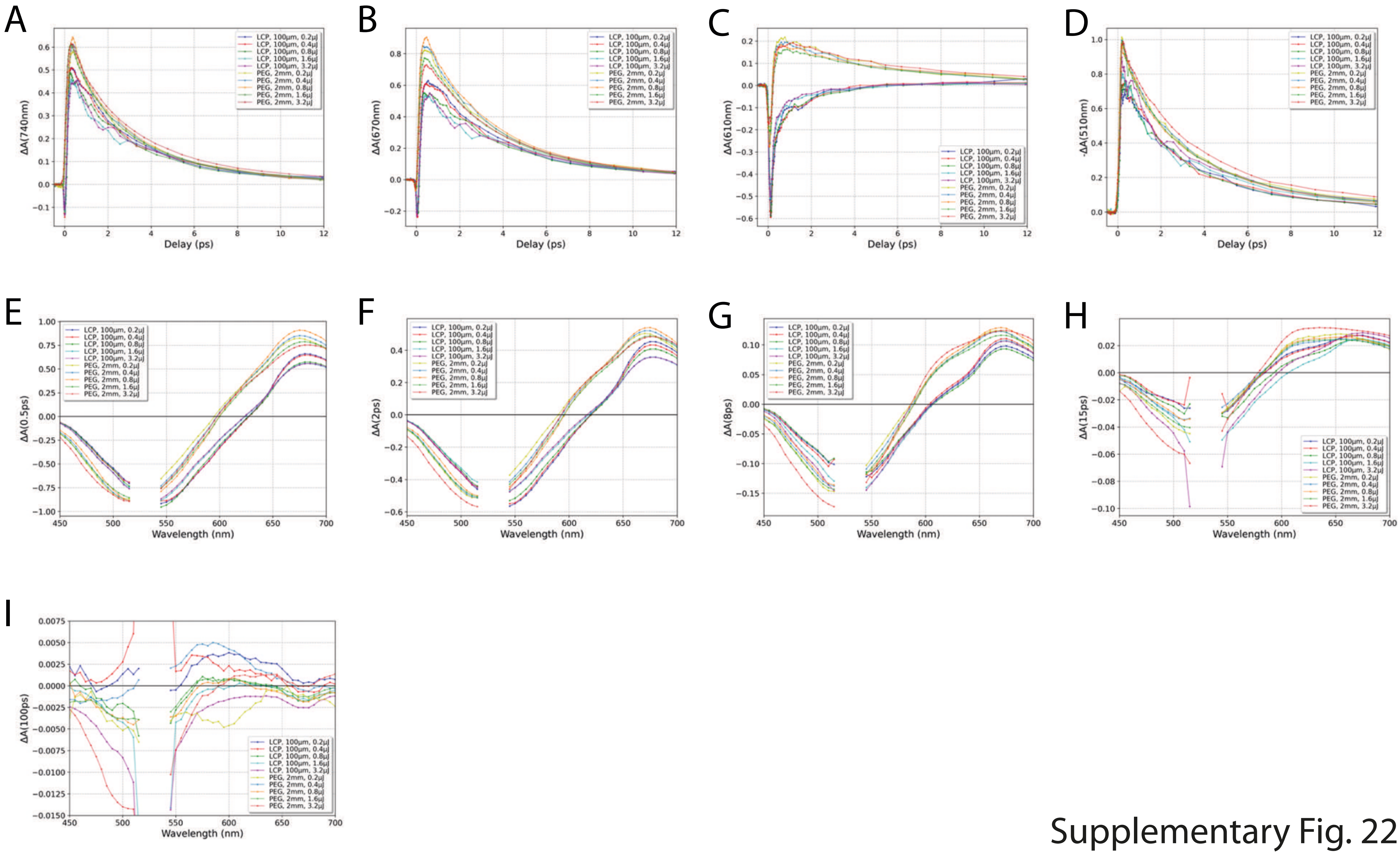


Supplementary Fig. 22

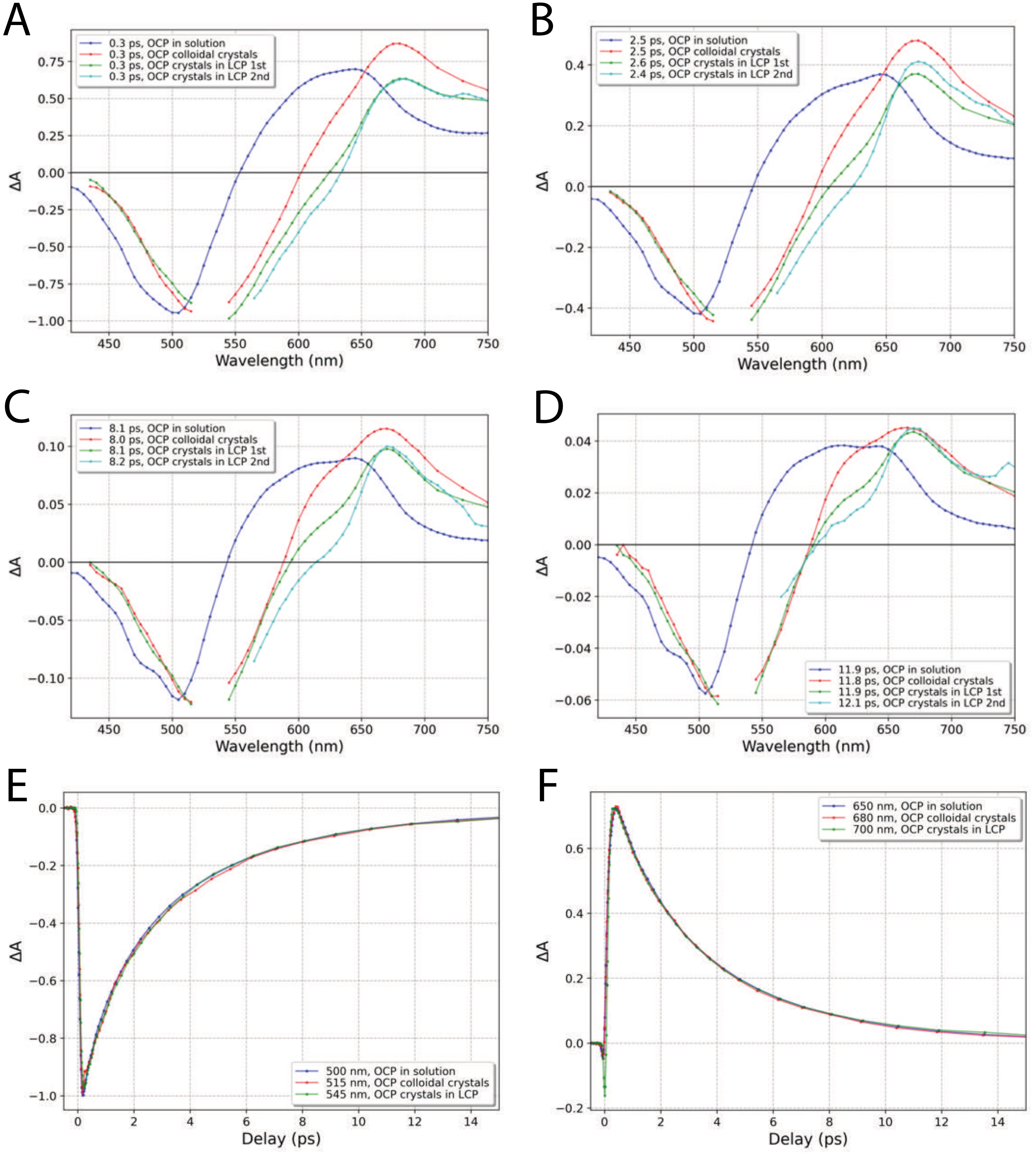


Supplementary Fig. 23

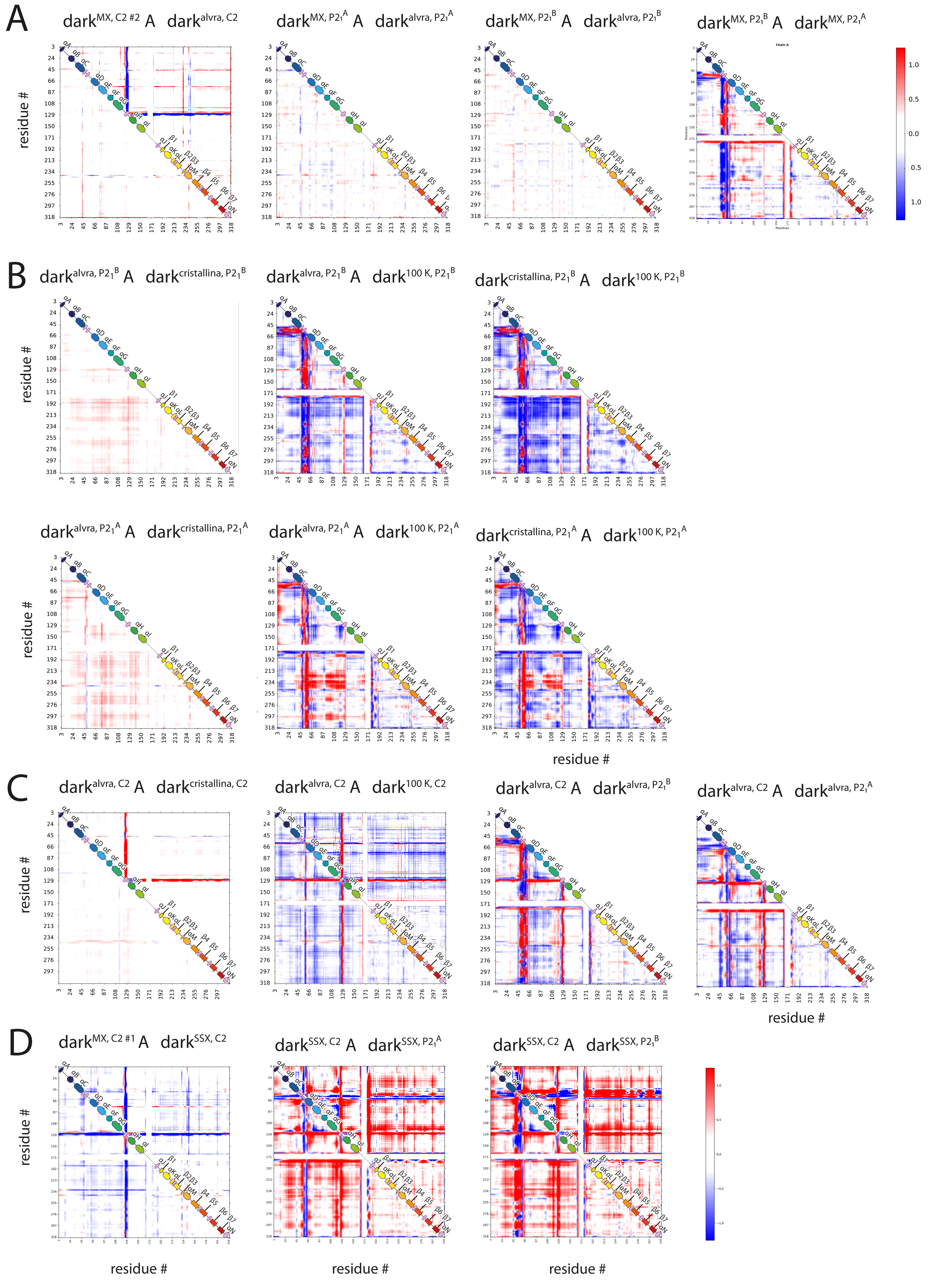


Supplementary Fig. 24

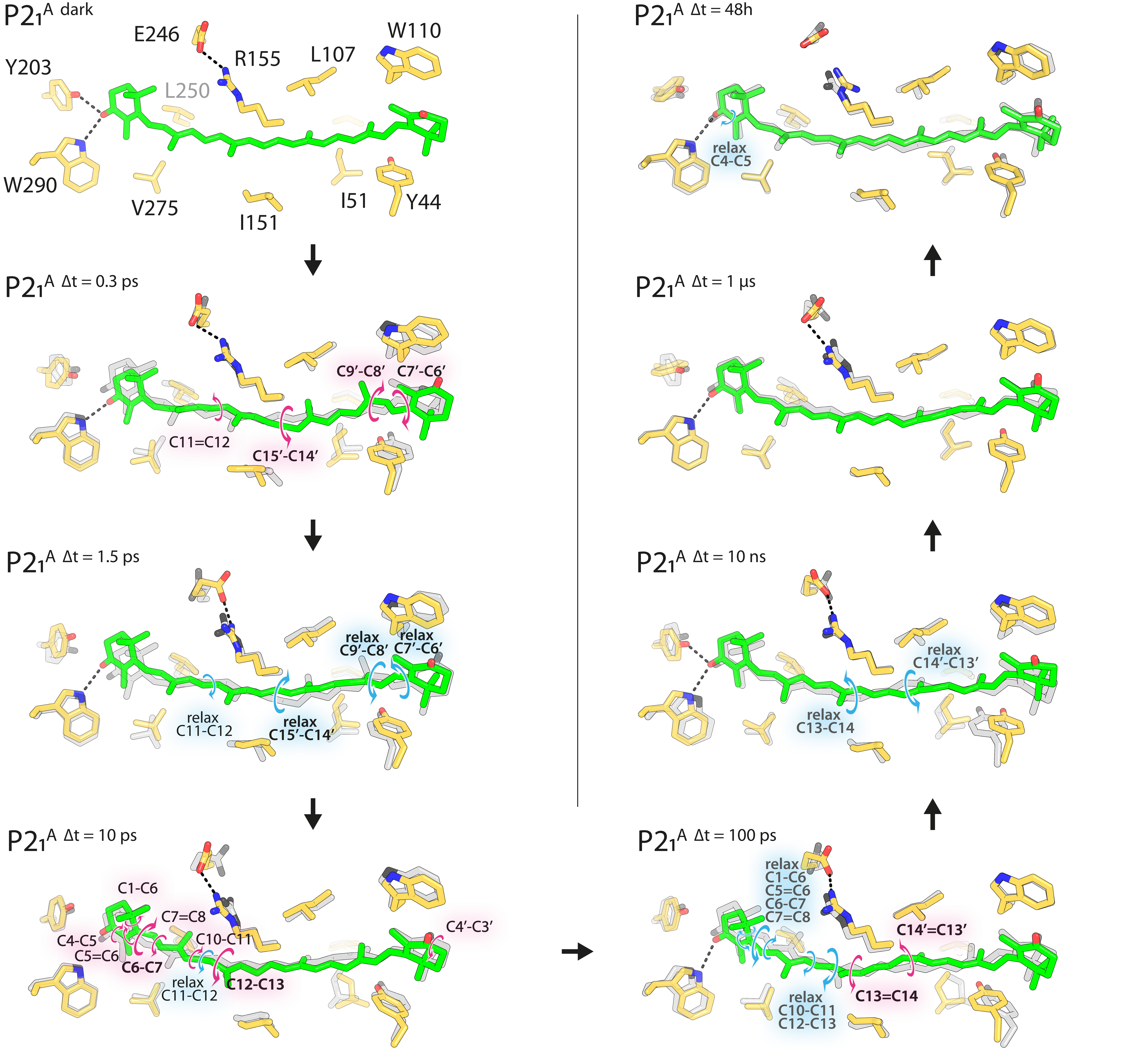


Supplementary Fig. 25

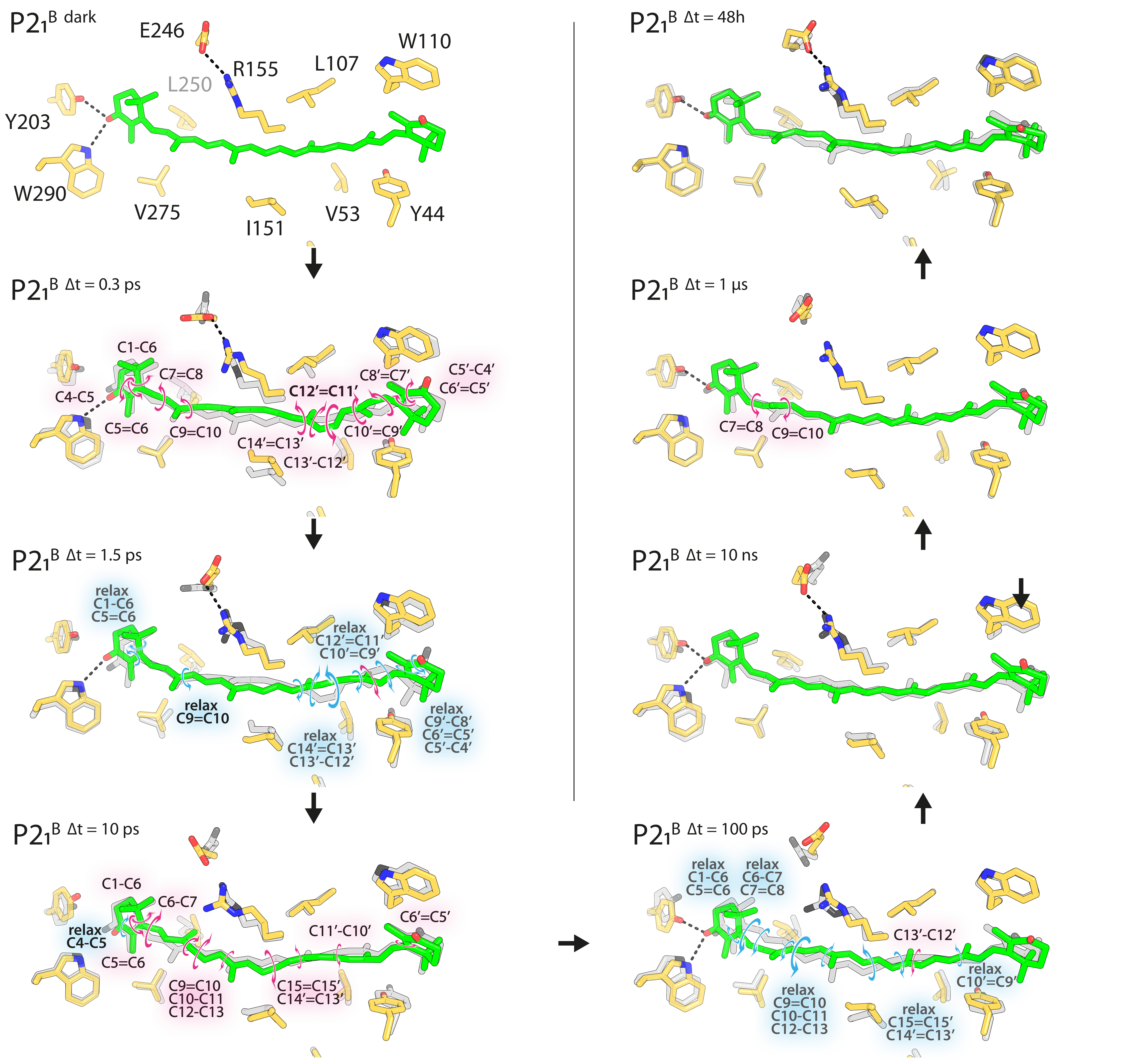


Supplementary Fig. 26

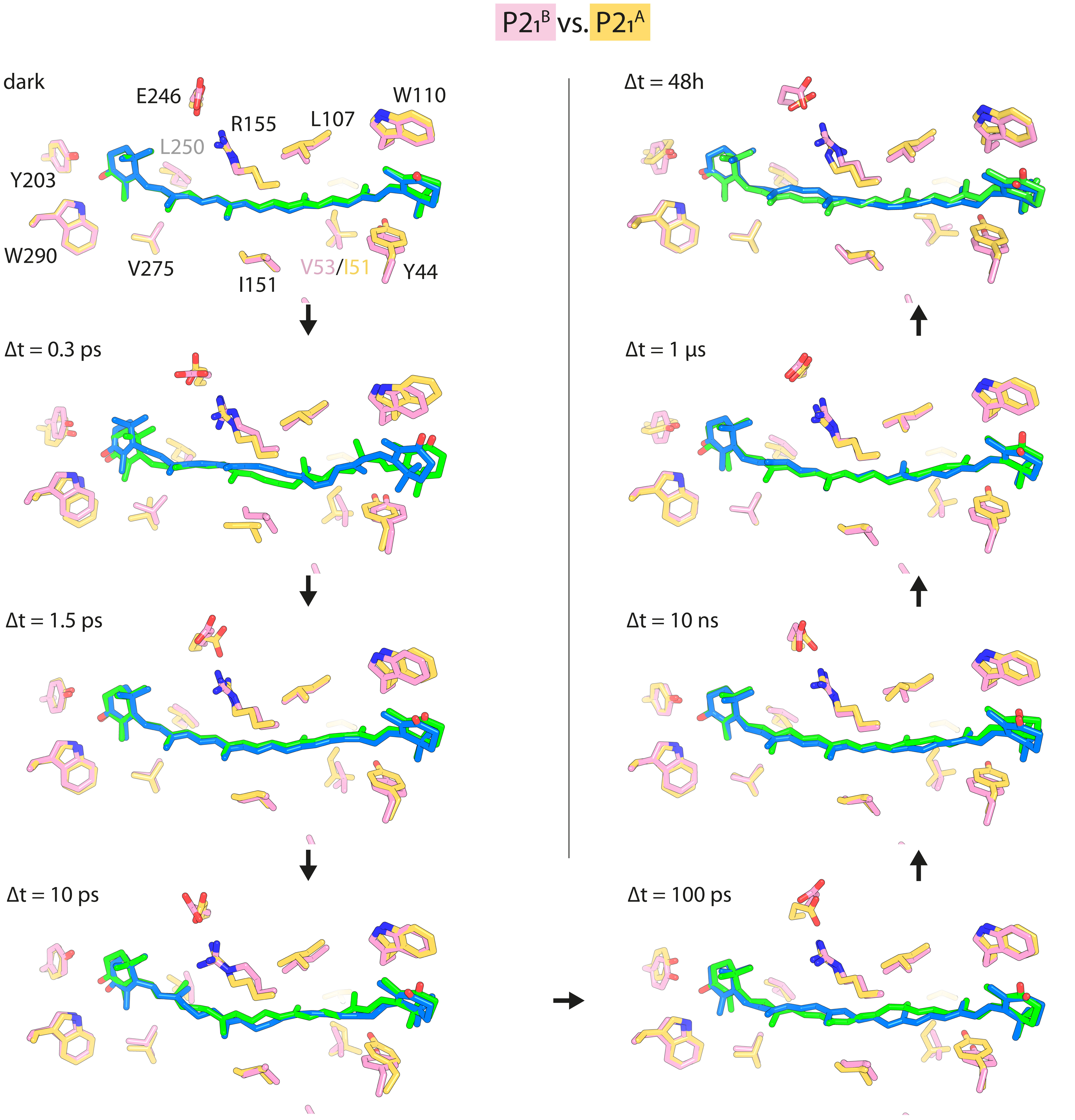


Supplementary Fig. 27

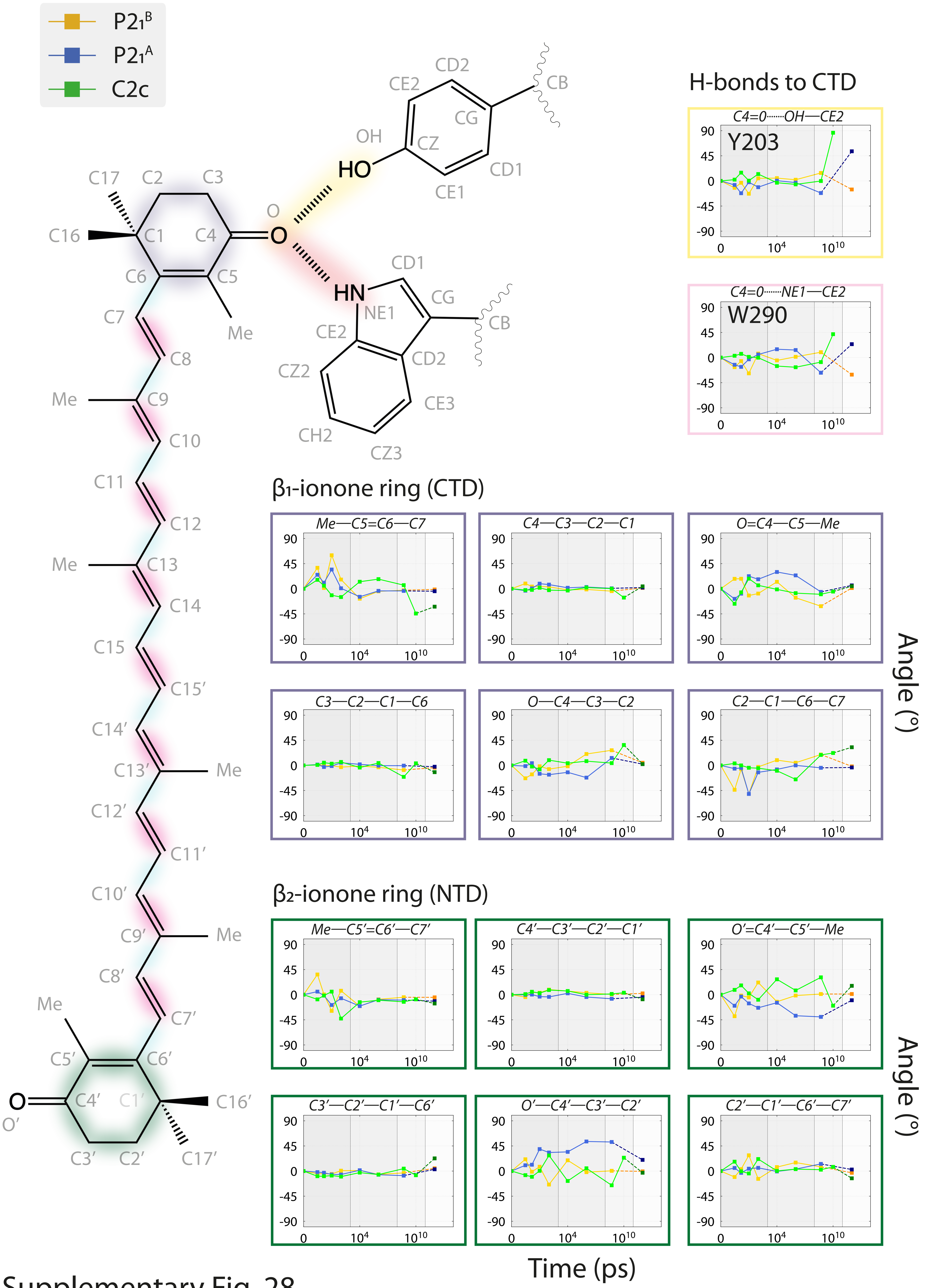


Supplementary Fig. 28

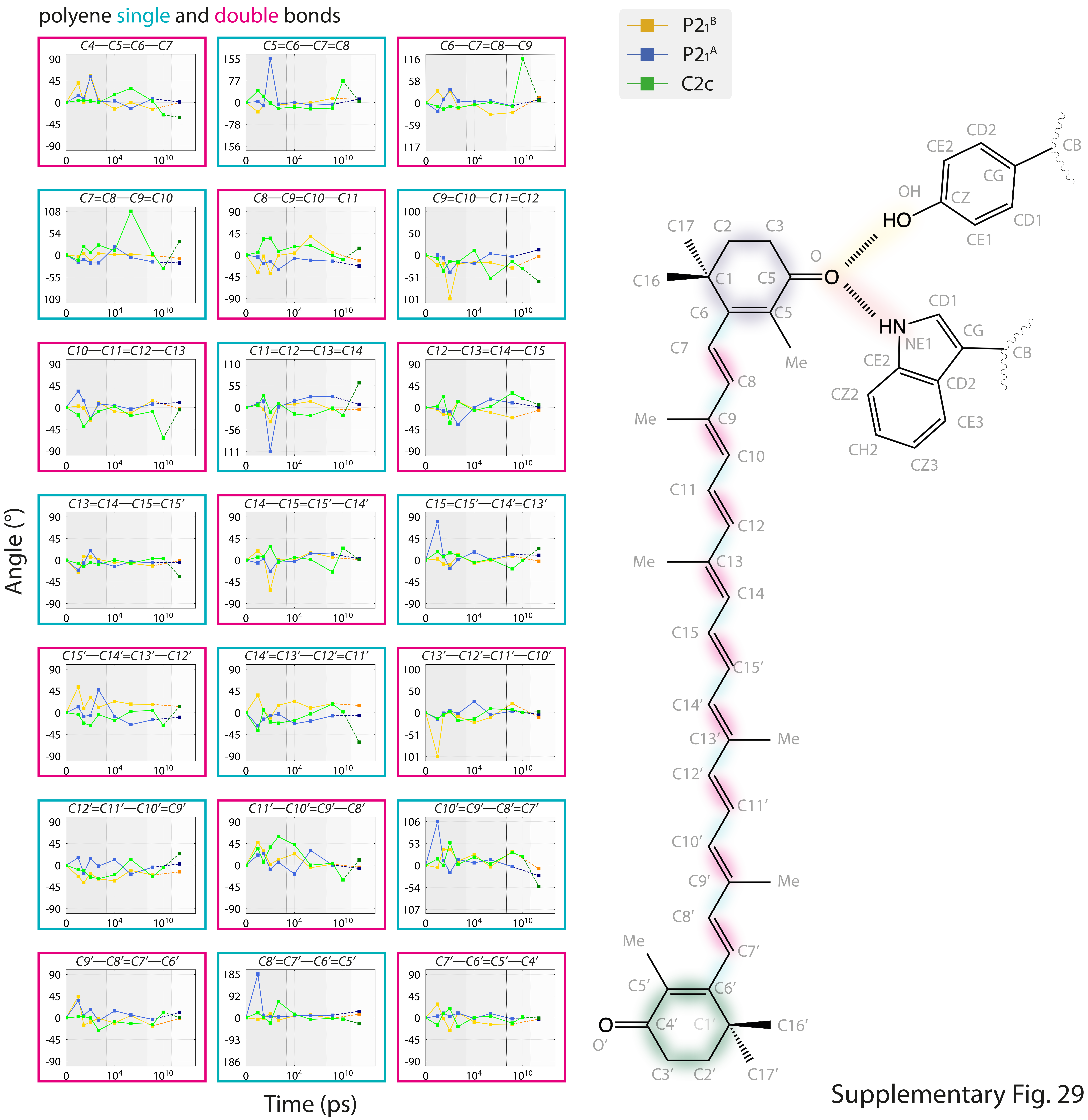


Supplementary Fig. 29

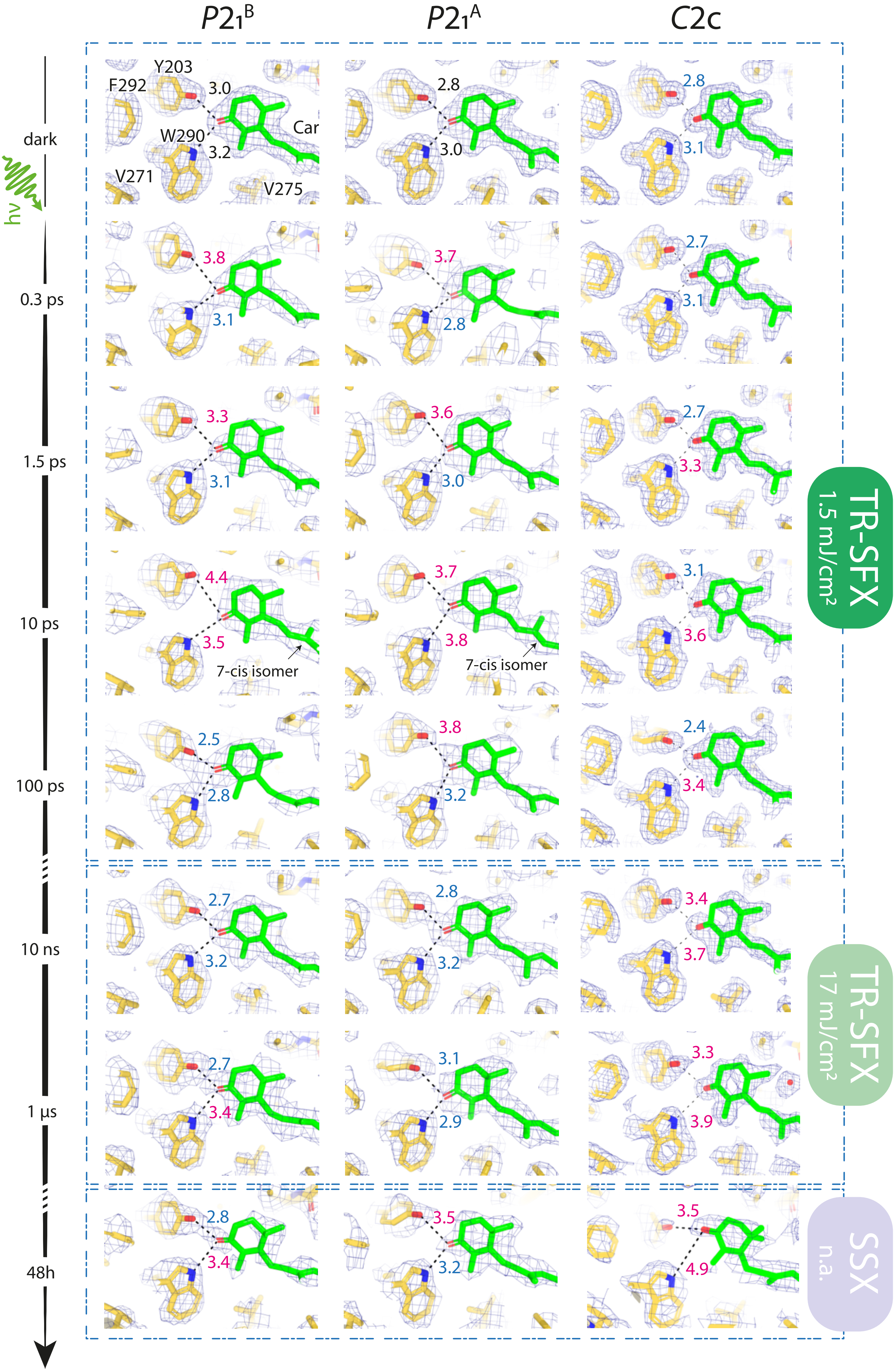


Supplementary Fig. 30

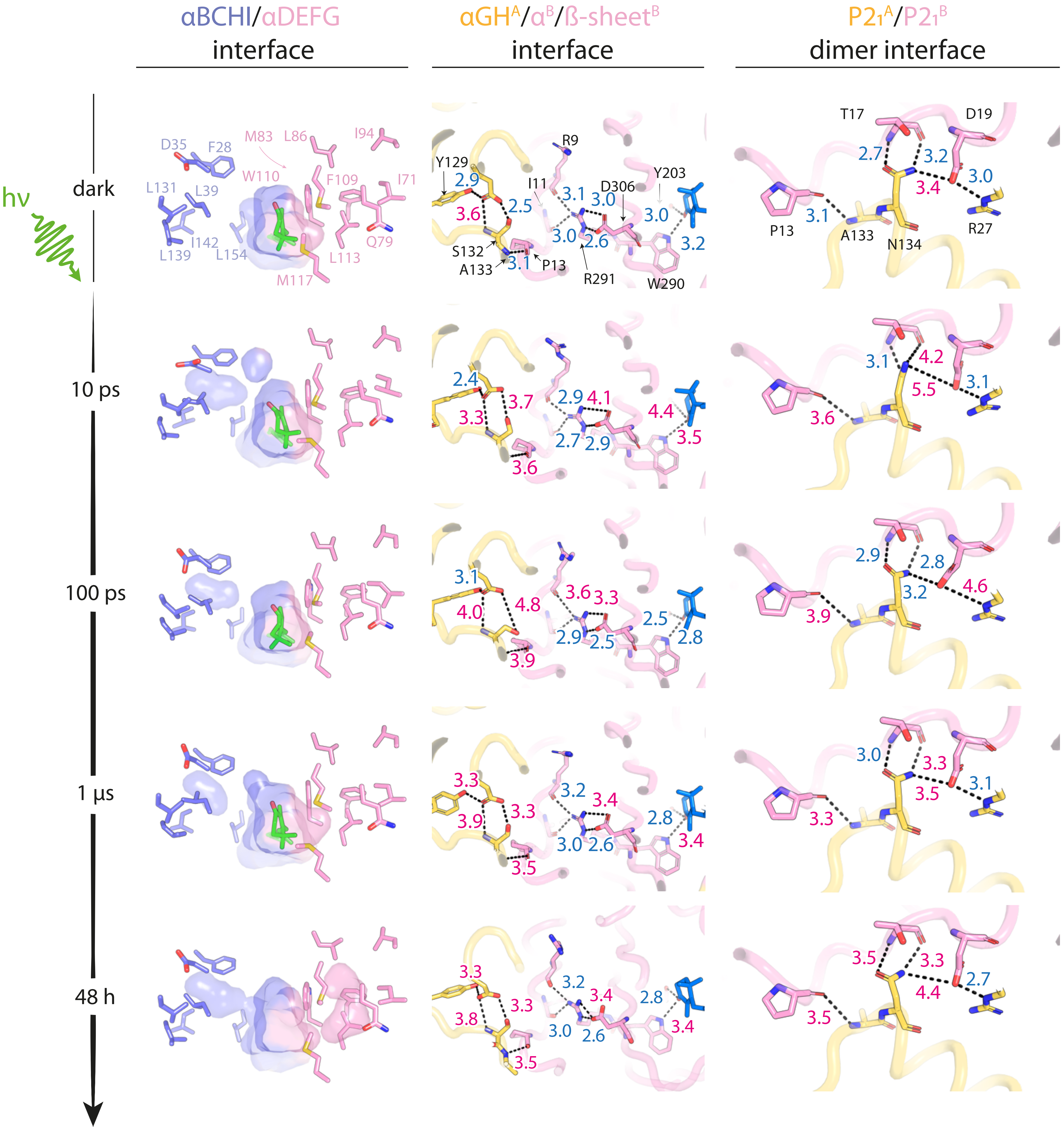


Supplementary Fig. 31

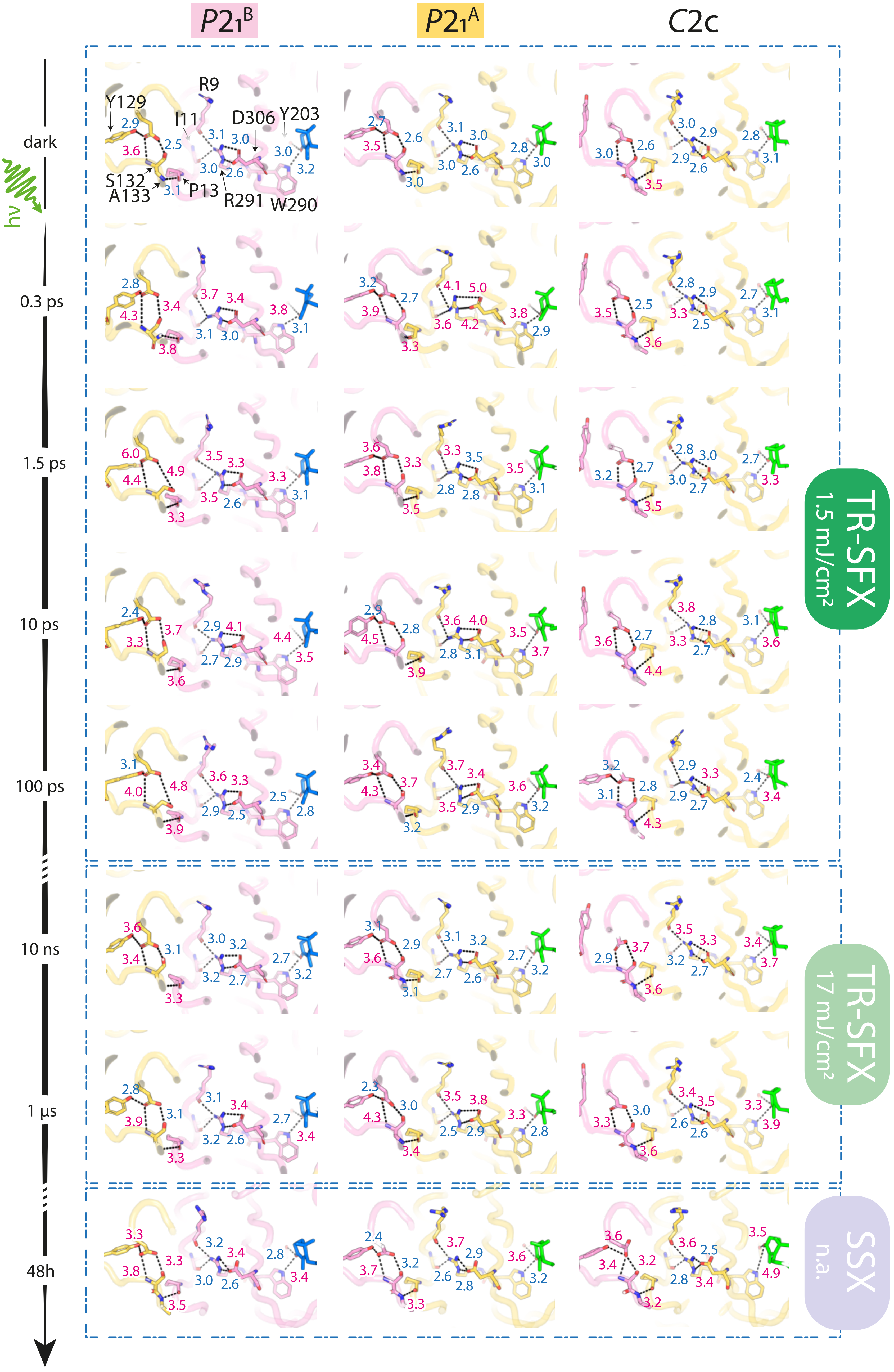


Supplementary Fig. 32

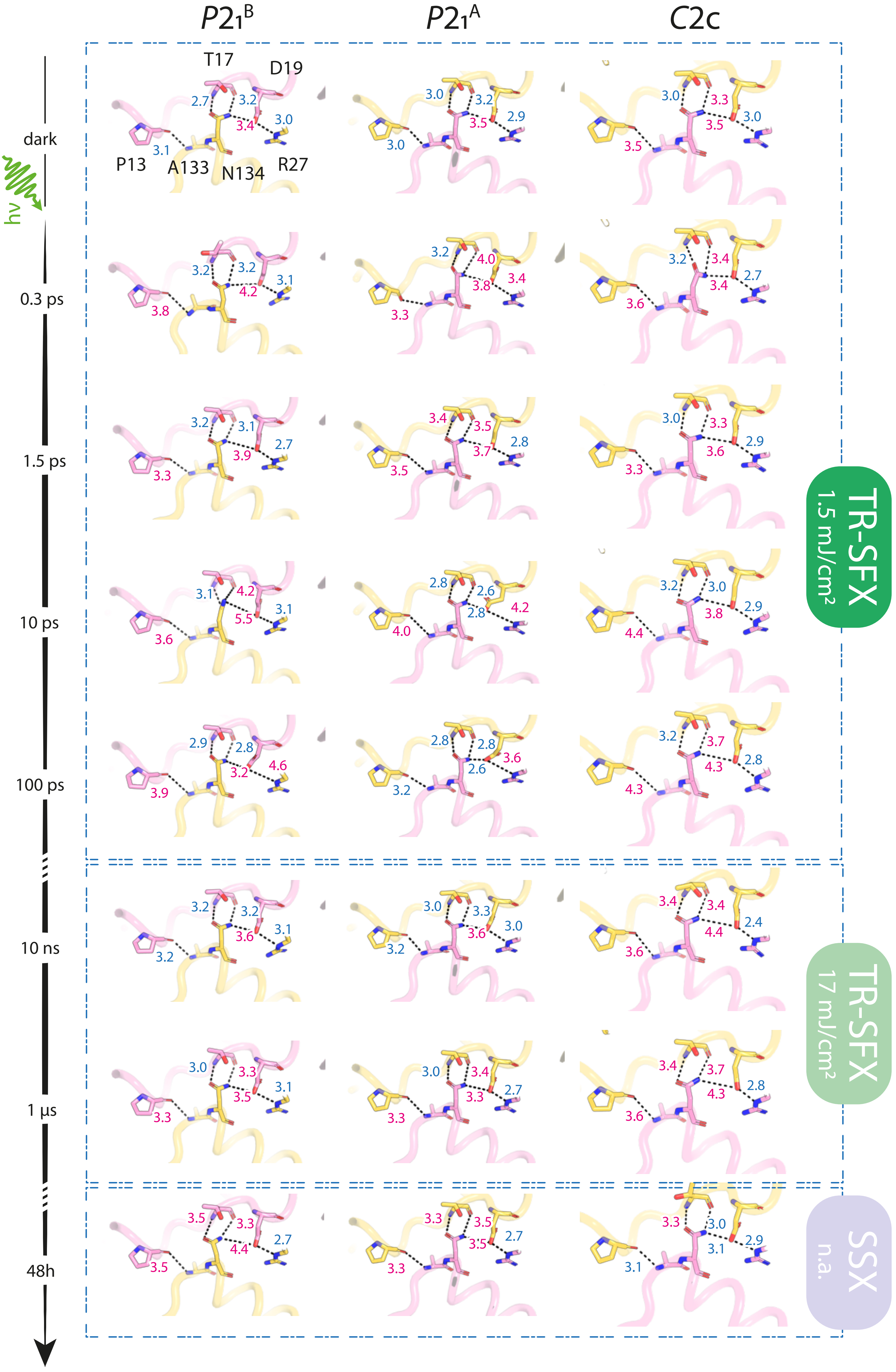


Supplementary Fig. 33

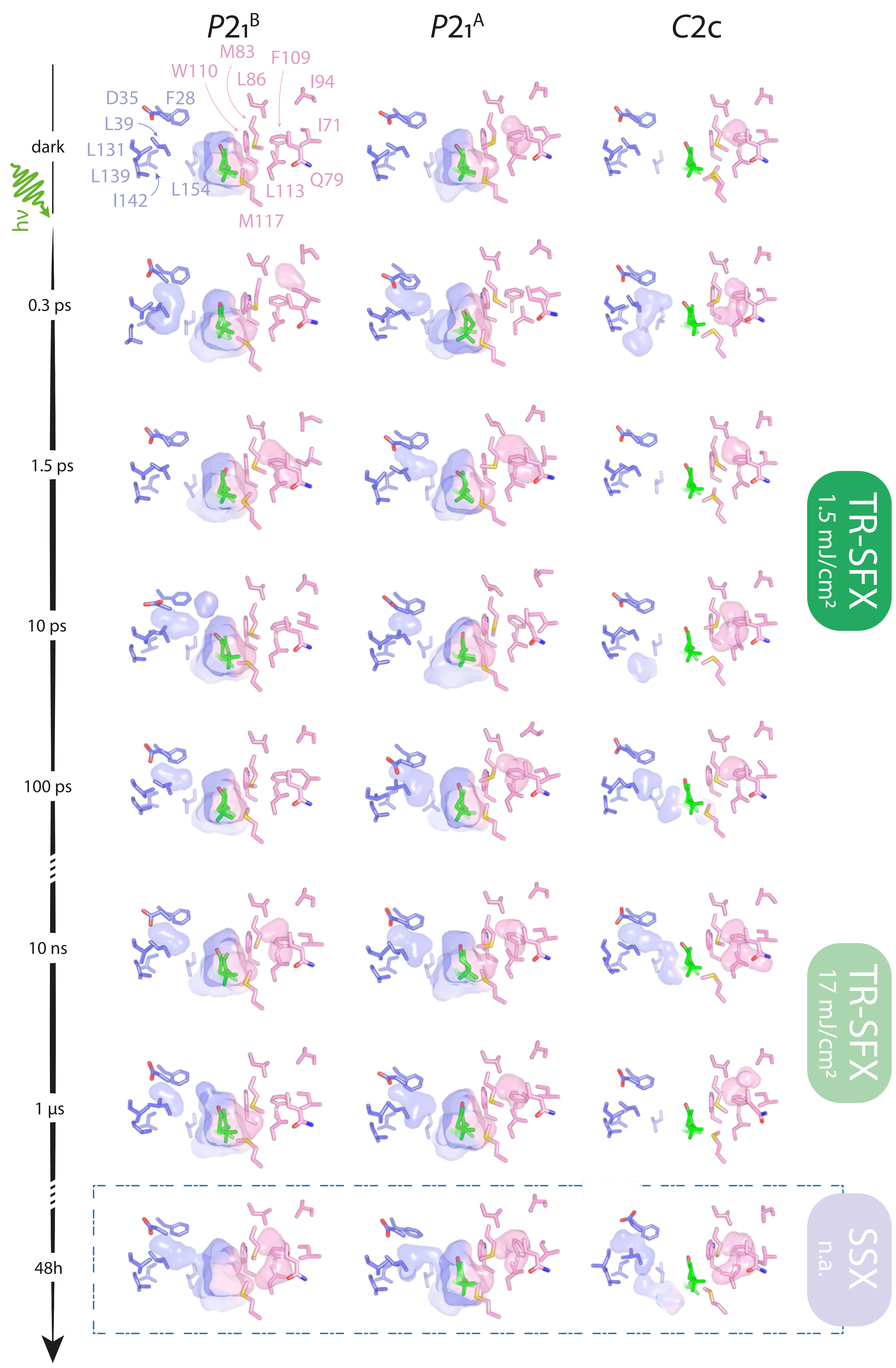


Supplementary Fig. 34

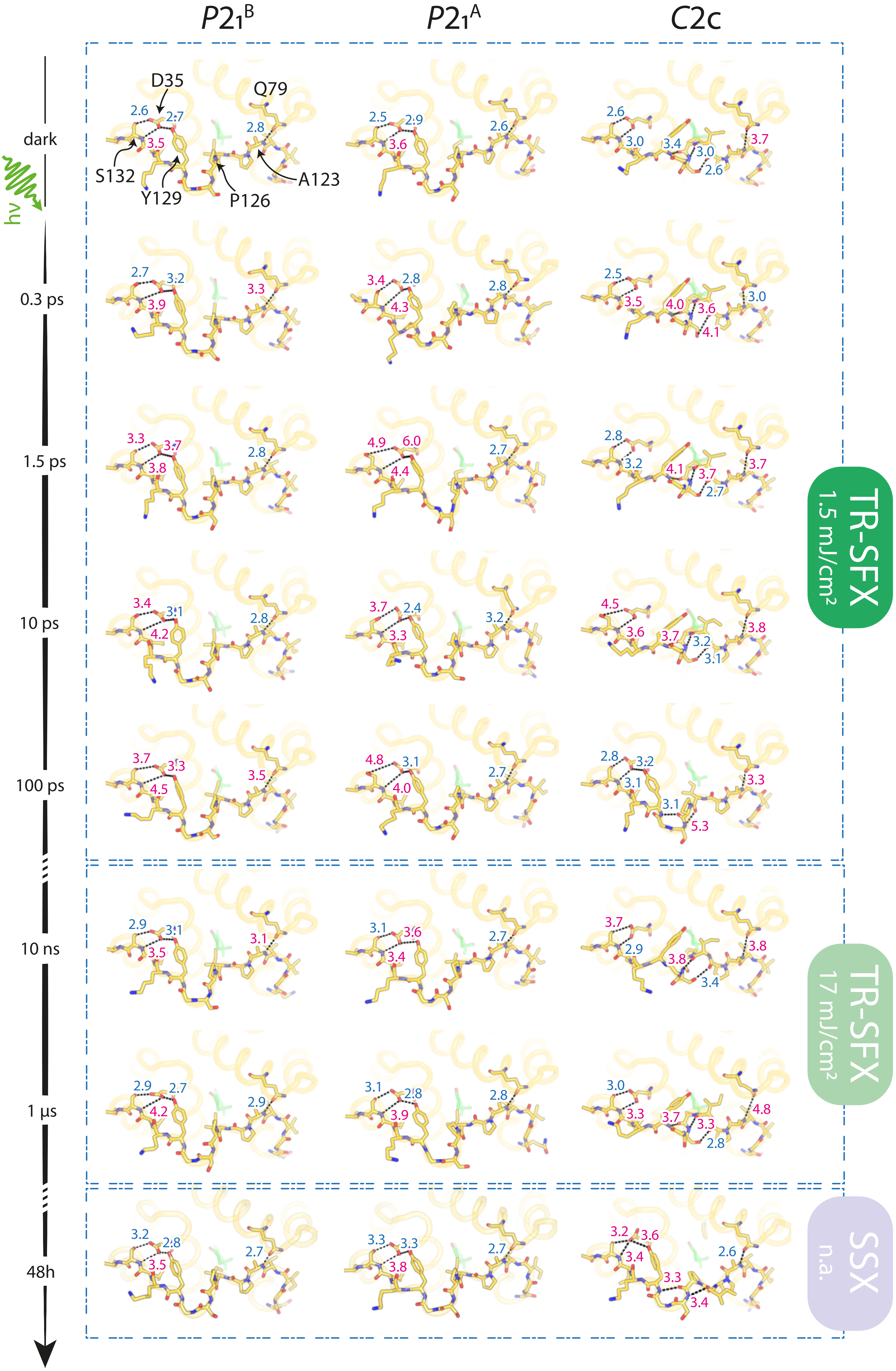


Supplementary Fig. 35

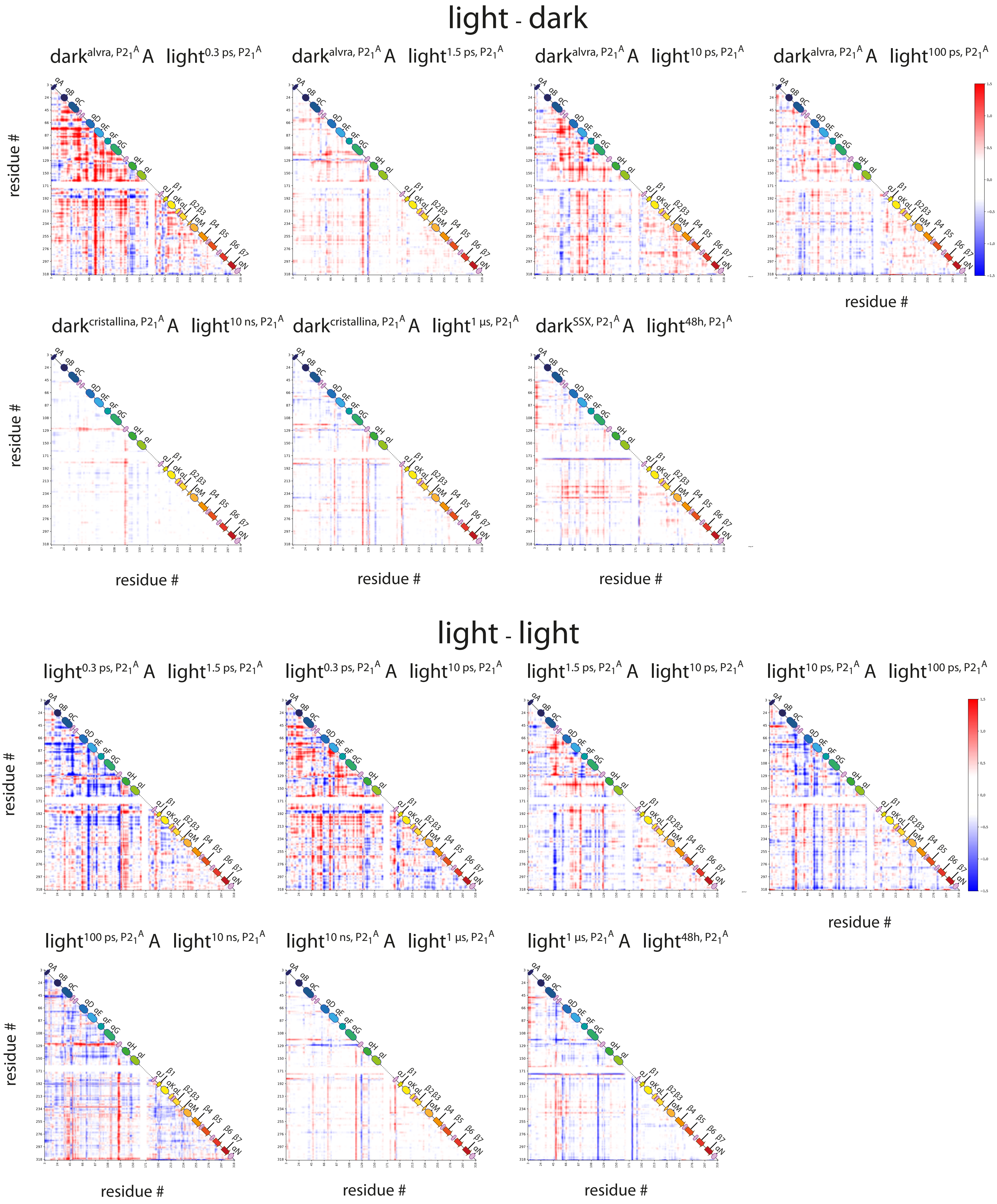


Supplementary Fig. 36

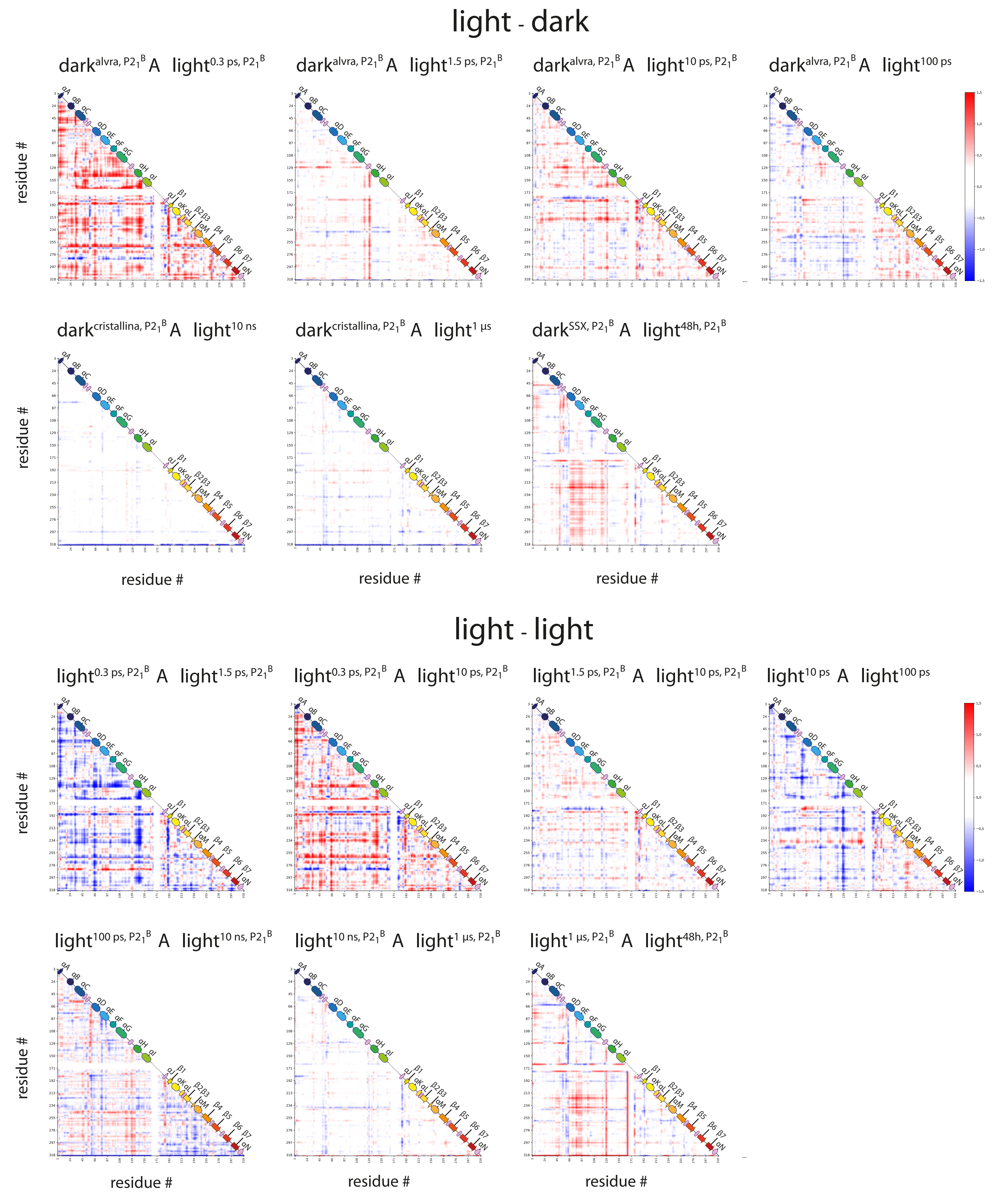


Supplementary Fig. 37

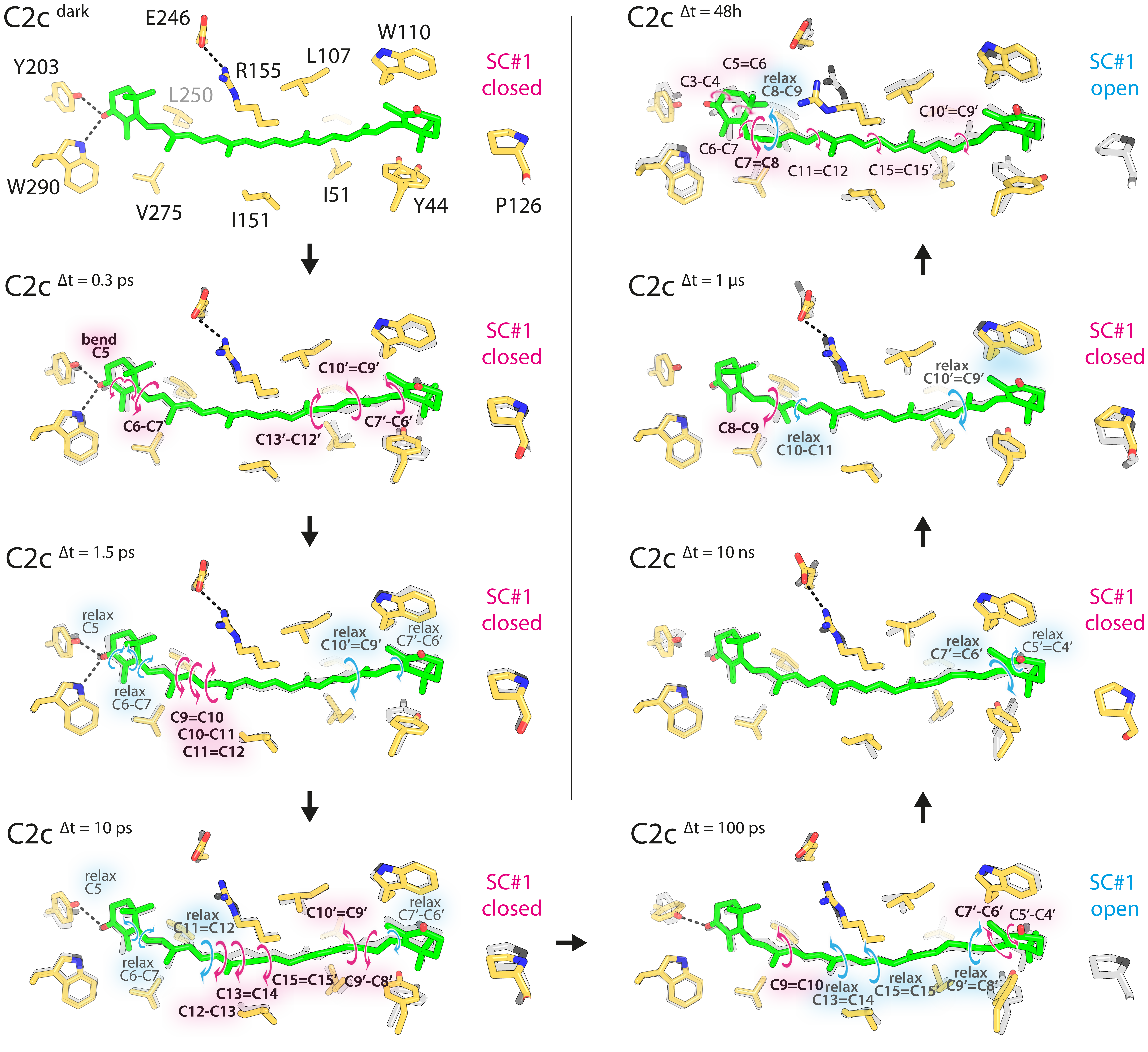


Supplementary Fig. 38

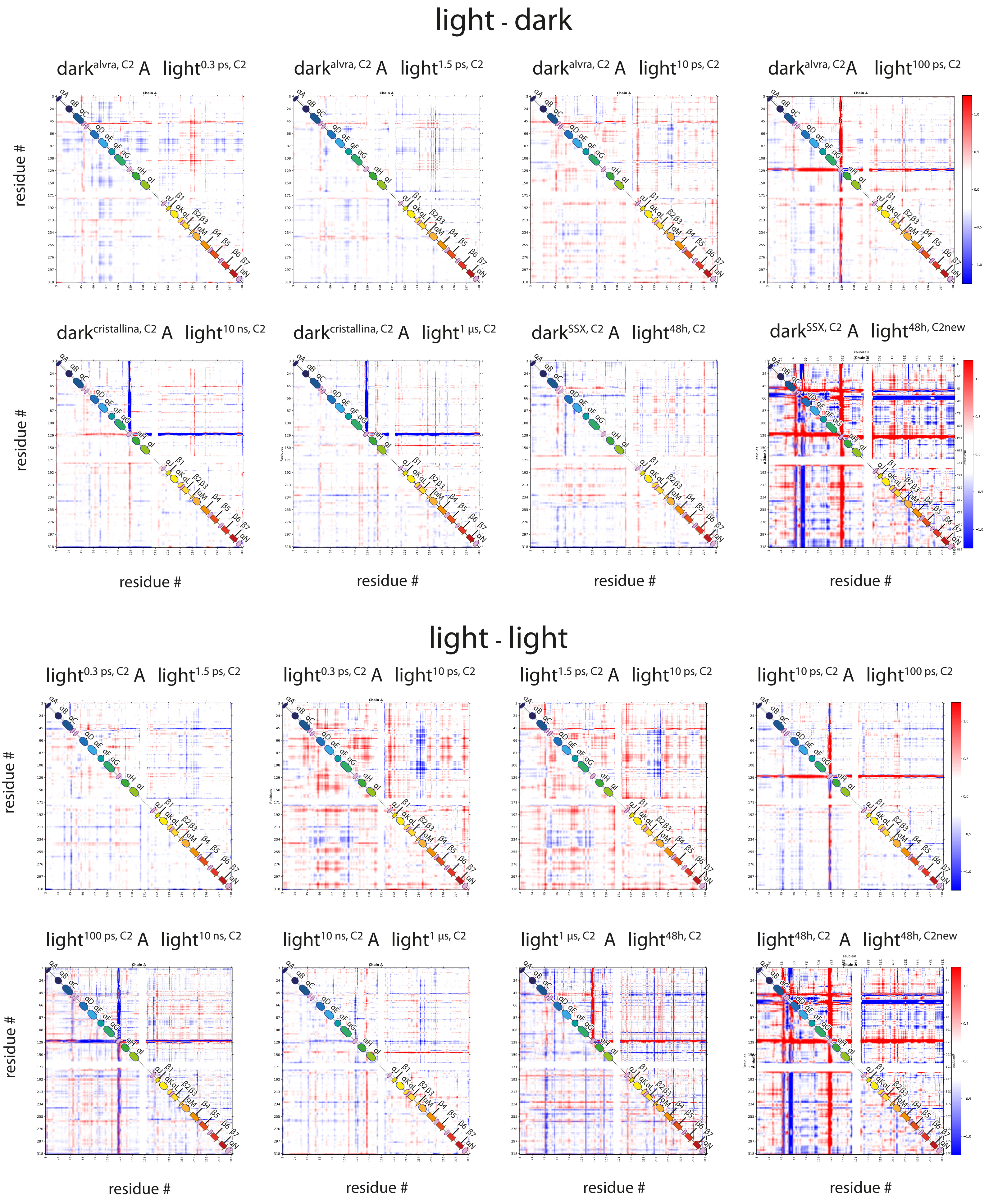


Supplementary Fig. 39

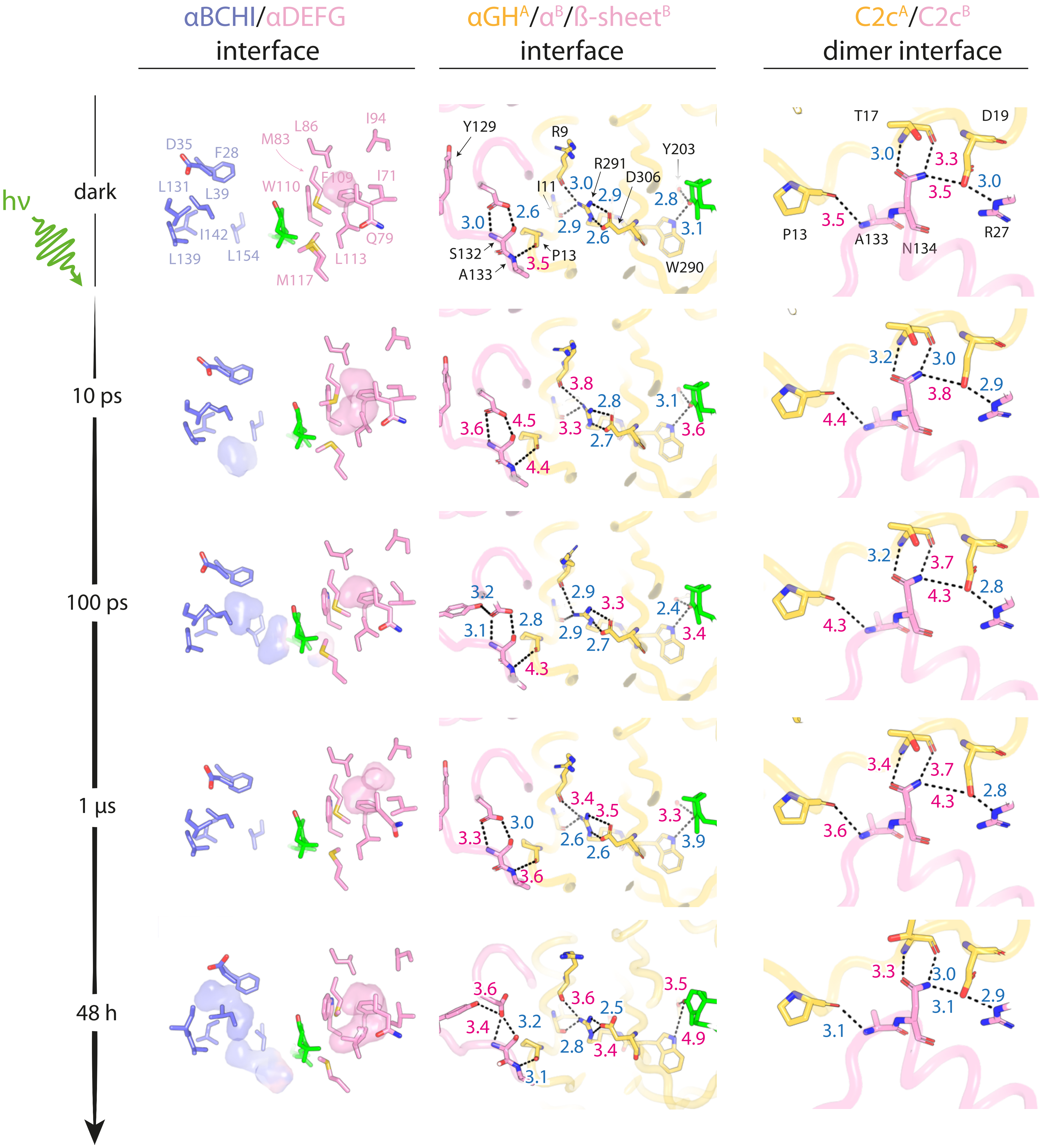


Supplementary Fig. 40

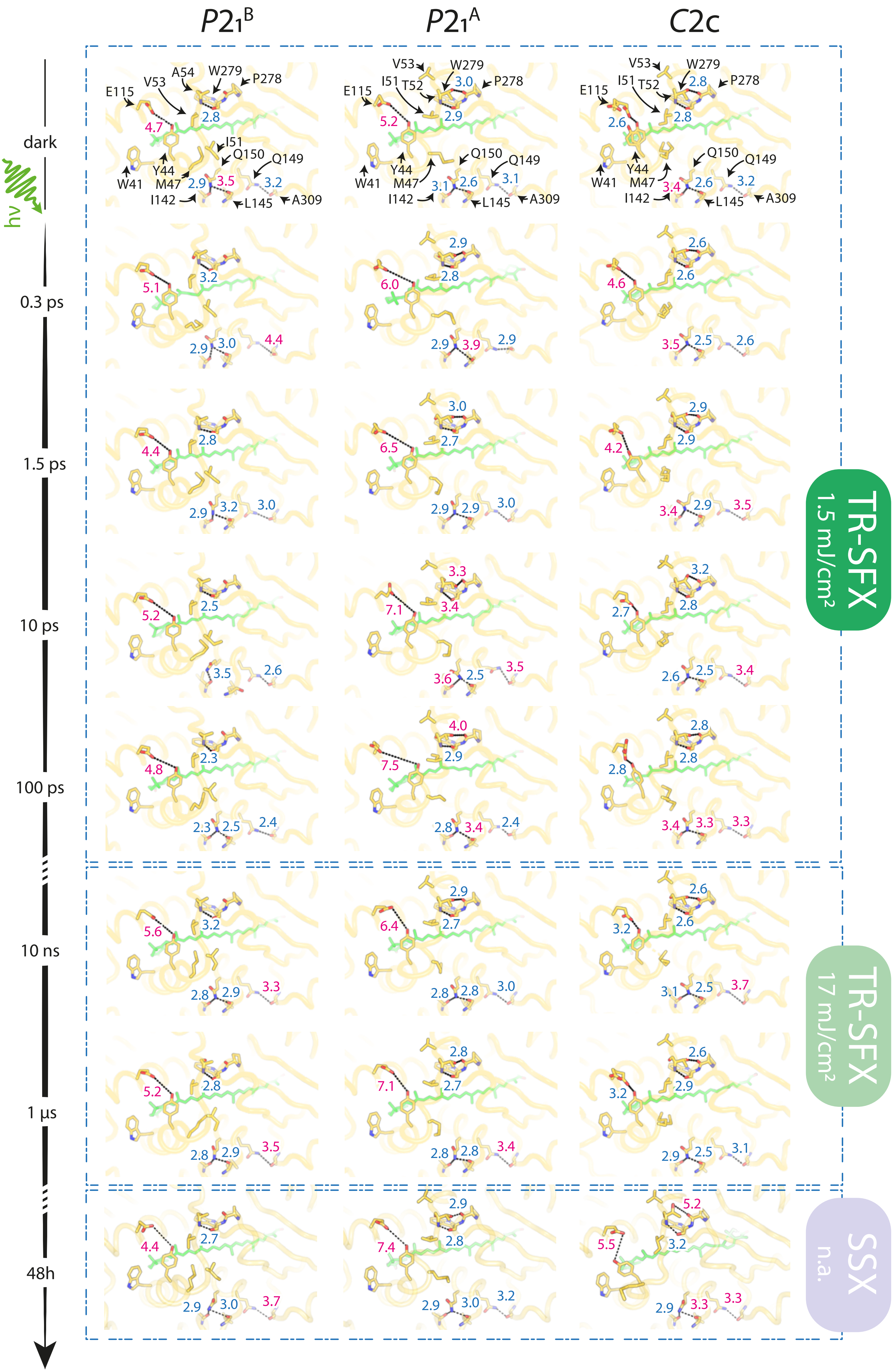

P21B
P21A
C2c
dark
hν
0.3 ps
1.5 ps
10 ps
100 ps
10 ns
1 µs
48h
E115
V53
A54
W279
P278
I51
T52
Y44
W41
M47
Q150
Q149
I142
L145
A309
TR-SFX 1.5 mJ/cm²
TR-SFX 17 mJ/cm²
SSX n.a.

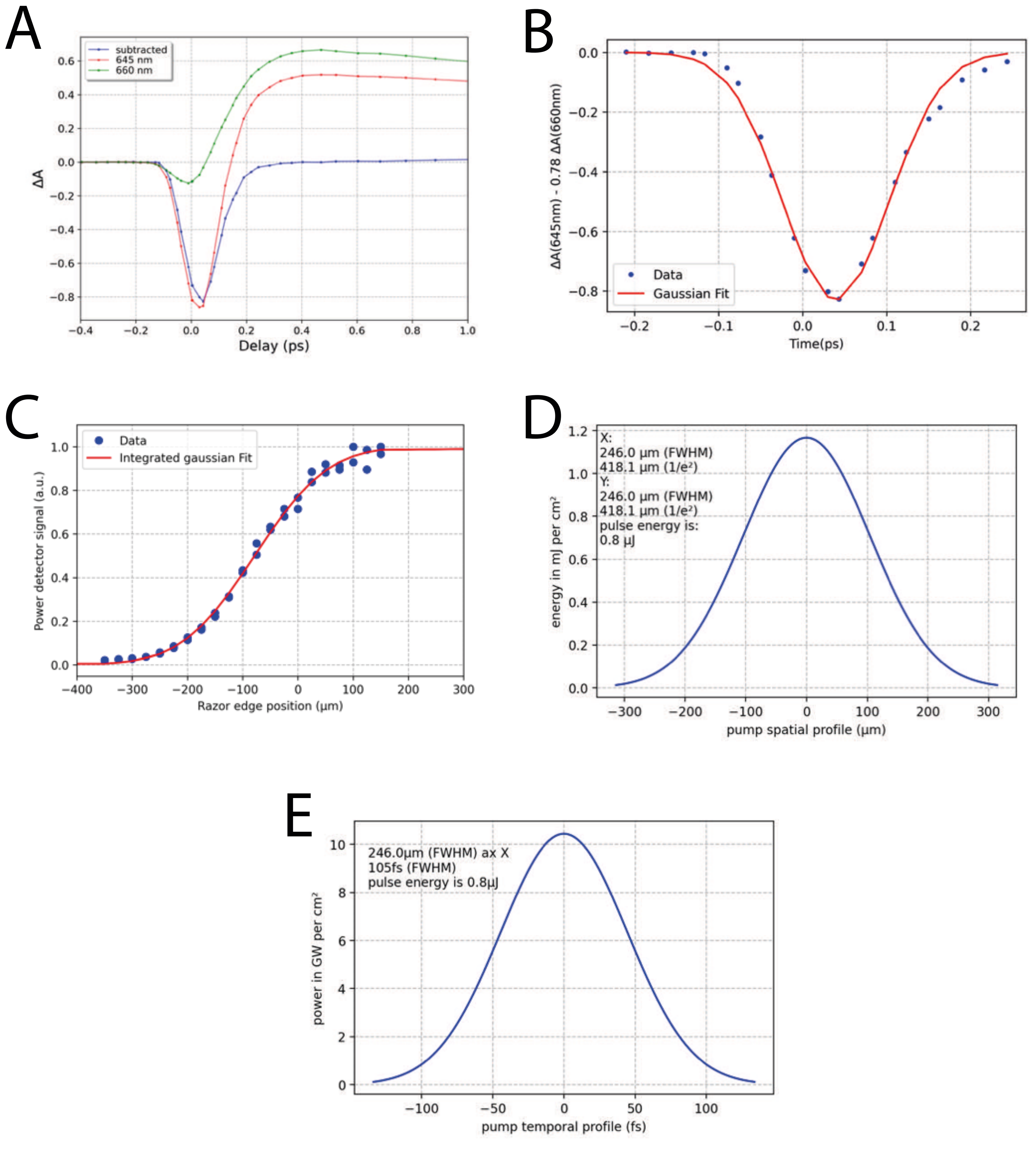


Supplementary Fig. S42
